# Gradient-based parameter optimization to determine membrane ionic current composition of human induced pluripotent stem cell-derived cardiomyocytes

**DOI:** 10.1101/2022.05.16.492203

**Authors:** Hirohiko Kohjitani, Shigeya Koda, Yukiko Himeno, Takeru Makiyama, Yuta Yamamoto, Daisuke Yoshinaga, Yimin Wuriyanghai, Asami Kashiwa, Futoshi Toyoda, Yixin Zhang, Akira Amano, Akinori Noma, Takeshi Kimura

**Affiliations:** Department of Cardiovascular Medicine, Kyoto University Graduate School of Medicine, Kyoto, Japan; Graduate School of Life Sciences, Ritsumeikan University, Kusatsu, Japan; Department Pediatrics, Kyoto University Graduate School of Medicine, Kyoto, Japan; Department of Physiology, Shiga University of Medical Science, Otsu, Japan

**Keywords:** Parameter optimization method, hiPSC-CMs, Cardiac action potential, Mathematical model, Computer simulation

## Abstract

Premature cardiac myocytes derived from human-induced pluripotent stem cells (hiPSC-CMs) show heterogeneous action potentials (APs), most probably because of different expression patterns of membrane ionic currents. We aim to develop a method of determining expression patterns of functional channels in terms of the whole-cell ionic conductances (*G_x_*) using individual spontaneous AP configurations. However, it has been suggested that apparently identical AP configurations were obtained by different sets of ionic currents in a mathematical model of cardiac membrane excitation. If so, the inverse problem of *G_x_* estimation might not be solved. We computationally tested the feasibility of the gradient-based optimization method. For realistic examination, conventional ‘cell-specific models’ were prepared by superimposing the model output of AP on each experimental AP record by the conventional manual adjustment of *G_x_*s of the baseline model. Then, *G_x_*s of 4 ~ 6 major ionic currents of the ‘cell-specific models’ were randomized within a range of ±5 ~ 15% and were used as initial parameter sets for the gradient-based automatic *G_x_*s recovery by decreasing the mean square error (MSE) between the target and model output. When plotted all data points of MSE - *G_x_* relation during the optimization, we found that the randomized population of *G_x_*s progressively converged to the original value of the cell-specific model with decreasing MSE. To confirm the absence of any other local minimum in the global search space, we mapped the MSE by randomizing *G_x_*s over a range of 0.1 ~ 10 times the control. No additional local minimum of MSE was obvious in the whole parameter space besides the global minimum of MSE at the default model parameter.

## 2. Introduction

During more than a half-century, the biophysical characteristics of ion transporting molecules (channels and ion exchangers) have been extensively analyzed, and biophysical models of each functional component have largely been detailed [1–4] (for human-induced pluripotent stem cells (hiPSC-CMs) see [5–7]). In addition, various composite cell models, including the membrane excitation, cell contraction, and the homeostasis of the intracellular ionic composition, have been developed by integrating mathematical models at molecular levels into the cardiac cell models [8–11]. These models have already been quite useful in visualizing individual currents underlying the action potential (AP) configuration under various experimental conditions in matured cardiac myocytes. However, the utility of these mathematical cell models has been limited because of the lack of extensive validation for the accuracy of the model output. This is the drawback of the subjective manual fitting used in almost all mathematical cardiac cell models so far published. A new challenge of such mechanistic models of cardiac membrane excitation might be an examination in a very different paradigm to assess if the large but continuous variety of cardiac AP configurations, for example, those recorded in the hiPSC-CMs, can be reconstructed by applying the automatic parameter optimization method to the human cardiac cell models.

The automatic parameter optimization technique has been used to determine parameters objectively in a wide range of various biological models (in cardiac electrophysiology; [12–15], in the systems pharmacology; [16–20]). Because of this utility, a large variety of improvements have been made in the area of information technology [21,22]. However, in electrophysiology, it has been suggested that different combinations of model parameters can produce APs, which are very similar[23–25] (see also [13]). It has been considered that the determination of current density at high fidelity and accuracy requires additional improvements to the optimization method in the cardiac cell model because of complex interactions among ionic currents underlying the membrane excitation (see [26], for review; [23]).

The final goal of our study is to develop an objective and accurate method of determining the current profile (that is, the expression level of functional ionic currents) underlying individual AP configurations. As a case study, we select a large variety of AP configurations in the hiPSC-CMs, which are difficult to classify into the conventional nodal-, atrial- or ventricular-types. Nevertheless, it has been clarified that the molecular bases of the ion channels expressed in the hiPSC-CMs well correspond to those in the adult cardiac myocytes (GSE154580 <u>GEO Accession viewer (nih.gov)</u>). Thus, we use the human ventricular cell model (hVC model, [11]) for the baseline model. In the present study, we computationally examine the feasibility of the basic gradient-based optimization method, pattern search (PS) algorithm [21,27,28] in the model of cardiac AP generation. We prepared a given AP configuration using each ‘cell specific model’, which was prepared by the conventional manual fitting of the hVC model to the respective experimental recordings. To assess the accuracy of the PS method of parameter optimization, this AP waveform generated by the cell-specific model was used as a target of the optimization. Then, the initial set of parameters for the optimization was prepared by uniform randomization centered around the model’s default values. The PS algorithm should return the original parameters by decreasing the error function (MSE) between the modified model output and target AP waveforms. The accuracy of optimization was definitely judged by recovering of the original values of each ionic current amplitude as the MSE progressively decreased toward zero.

## 3. Materials and Methods

### 3.1. The baseline model of hiPSC-CM membrane excitation

The baseline model of hiPSC-CMs was essentially the same as the human ventricular cell model (hVC model), which has been fully described in references [10,11] and shares many comparable characteristics with other human models so far published [8,9]. The model structure of the hVC model consists of the cell membrane with a number of ionic channel species and a few ion transporters, the sarcoplasmic reticulum equipped with the Ca^2+^ pump (SERCA), and the refined Ca^2+^ releasing units coupled with the L-type Ca^2+^ channels on the cell membrane at the nano-scale dyadic space, the contractile fibers, and the cytosolic three Ca^2+^ diffusion spaces containing several Ca^2+^-binding proteins (Fig S1). All model equations and abbreviations are in Supplemental Materials.

The source code of the present hiPSC-CM model was written in VB.Net and is available from our e-Heart website (http://www.eheartsim.com/en/downloads/).

The kinetics of the ionic currents in the baseline model were readjusted according to new experimental measurements if available in the hiPSC-CMs [29] (Fig S2). In the present study, the net membrane current (*I_tot_cell_*) is calculated as the sum of nine ion channel currents and two ion transporters (*I_NaK_* and *I_NCX_*) (Eq 1).

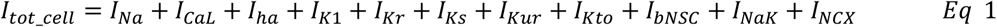

The membrane excitation of the model is generated by charging and discharging the membrane capacitance (*C_m_*) by the net ionic current (*I_tot_cell_*) across the cell membrane (Eq 2). The driving force for the ionic current is given by the potential difference between *V_m_* and the equilibrium potential (*E_x_*) (Eq 3). The net electrical conductance of the channel is changed by the dynamic changes in the open probability (*pO*) of the channel, which is mostly Vm-dependent through the *V_m_*-dependent rate constants (*α, β*) of the opening and closing conformation changes of the channel (Eqs 4 and 5).

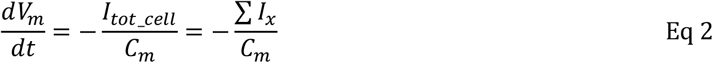

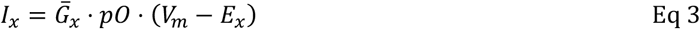

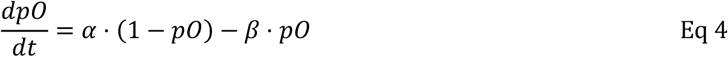

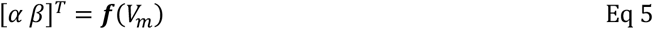

The exchange of 3Na^+^ / 2K^+^ by the Na/K pump and the 3Na^+^ / 1Ca^2+^ exchange by the NCX also generate sizeable fractions of membrane ionic current, *I_NaK_*, and *I_NCX_*, respectively. We excluded background currents of much smaller amplitude, such as *I_KACh_, I_KATP_, I_LCCa_* and *I_Cab_*, from the parameter optimization and adjusted only the non-selective background cation current (*I_bNSC_*) of significant amplitude for the sake of simplicity [30–32]. The *I_bNSC_* is re-defined in the present study as a time-independent net current, which remained after blocking all time-dependent currents.

### 3.2. The computational parameter optimization

The whole cell conductance *G_x_* of a given current system (*x*) is modified by multiplying the limiting conductance *Ḡ_x_* (Eq 3) of the baseline model by a scaling factor *sf_x_* (Eq 6) and are used for the parameter optimization.

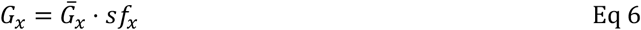

The mean square error (MSE) function (Eq 7) was used in the parameter optimization, where *V_m,a_* represents adaptive *V_m_* (the model output) generated by adjusting *sf_x_*s of the baseline model. The target *V_m,t_* represents the AP of the intact baseline model.

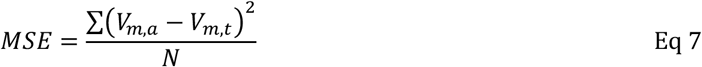

The MSE was stabilized by obtaining a quasi-stable rhythm of spontaneous APs through continuous numerical integration of the model, usually 30 ~ 100 spontaneous cycles were calculated for a new set of *sf_x_*s. The MSE was calculated within a time window. The width of this time window was adjusted according to the AP phase of interest. N is the number of digitized *V_m_* points with a time interval of 0.1 ms.

In the usual parameter optimization, the *V_m,a_* is generated by modifying the baseline model for comparison with the experimental record (*V_m,t_* = *V_m,rec_*). However, to evaluate the identifiability of the parameter optimization, a simple approach was taken in the present study. Namely, we used the manually adjusted ‘cell-specific’ model for the target (Vm,t), which was nearly identical to *V_m,rec_*. More importantly, the ‘cell-specific’ *V_m_* is totally free from extra-fluctuations (noise), which were observed in almost all AP recordings in hiPSC-CMs. In the optimization process, the initial value of each optimization parameter was prepared by randomizing the *sf_x_*s of the cell-specific model by ±5~15% at the beginning of each run of PS (*V_m,orp_*) in Eq 8 and the PS runs of several hundred were repeated. Thus, the error function is,

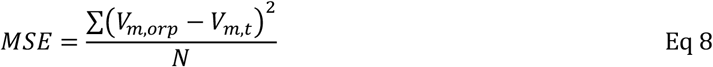

We call this optimization method ‘orp test’ in the present study.

The advantage of using a manually adjusted cell model for the optimization target is that the accuracy of parameter optimization is proved by recovering all *sf*_x_ = 1 independently from the randomized initial parameter set. Note the same approach was used in [23] in evaluating the accuracy of the parameter optimization by applying the genetic algorithm (GA) to the TNNP model of the human ventricular cell [33].

The optimization of using the randomized initial model parameters were repeated for more than 200 runs. Thus, the orp test might be classified in a ‘multi-run optimization’. The distribution of the *sf*_x_ data points obtained during all test runs was plotted in a single *sf*_x_-MSE coordinate to examine the convergence of individual *sf*_x_s with the progress of the orp test.

### 3.3. The pattern search method for the optimization

For a system showing the relatively simple gradient of MSE along the parameter axis, the gradient-based optimization methods are more efficient in general than the stochastic methods for this kind of objective function. We used one of the basic gradient-based optimization methods, the PS algorithm. The computer program code of the pattern search [34] is simple (see Supplemental Materials) and does not require derivatives of the objective function. We implemented the code into a homemade program for data analysis (in VB) to improve the method for better resolution and to save computation time.

The primary PS method uses a base and new points [27]. In brief, *sf*_x_ is coded with symbols *BP*_x_ and *NP*_x_ in the computer program, representing a base point (*BP*_x_) and a new searching point (*NP*_x_), respectively. Namely, MSE is calculated on each movement of *NP*_x_ by adding or subtracting a given step size (*stp*) *to the BP*_x_, and the search direction is decided by the smaller MSE. Then, the whole mathematical model is numerically integrated (Eqs 2, 3, 4, and 5) using *NP*_x_ to reconstruct the time course of AP (*V*_m,a_). This adjustment is conducted sequentially for each of the 4~6 selected currents in a single cycle of optimization. The cycle is repeated until no improvement in MSE is gained by a new set of *NP*_x_s. Then, the *BP*_x_ set is renewed by the new set of *NP*_x_ for the subsequent series of optimization. Simultaneously, the *stp* is reduced by a given reduction factor (*redFct* of 1/4). The individual PS run is continued until the new *stp* becomes smaller than the critical *stp* (*crtstp*), which is set to 2~10 × 10^−5^ in the present study.

### 3.4. Selection of ionic currents for the optimization

When we get a new experimental record of AP, we do not start the analysis with an automatic optimization of *G_x_* but first adjust the baseline model by conducting the conventional manual fitting. The nine ionic currents in Eq 1 in the baseline model are adjusted bit by bit to superimpose the simulated AP on the experimental one. During this step, it is important to pay attention to the influences of each *sf_x_* adjustment on the simulated AP configuration on the computer display. Thereby, one may find several key current components which should be used in the automatic parameter optimization. Usually, currents showing a relatively large magnitude of *G_x_* were selected for the automatic optimization according to Eq 2, while those which scarcely modified the simulated AP were left as default values in the baseline model.

### 3.5. Principal component analysis of the cell-specific models

When the orp test is conducted with *p* elements, it is possible to record the final point BP where the MSE is improved in the *p*-dimensional space. Suppose we represent the matrix when *n* data points are acquired as an *n* × *p* matrix *X*. In that case, we obtain a vector space based on the unit vector that maximizes the variance (first principal component: PC1) and the *p*-dimensional unit vector orthogonal to it (loadings vector ***W***(*k*) = (*w*_1_, *w*_2_ ⋯, *w_p_*). It is possible to convert each row, ***x***_(*i*)_ of the data matrix *X* into a vector of principal component scores, *t*_(*i*)_. The transformation is defined by

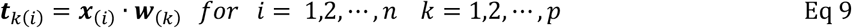

In order to maximize variance, the first weight vector w ((1)) corresponding to the first principal component thus has to satisfy,

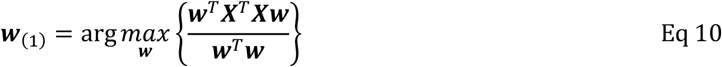

The k-th component can be found by subtracting the first (k-1)-th principal components from X

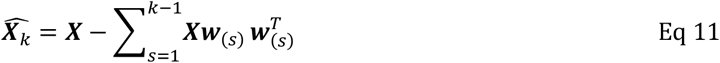

Then the weight vector is given as a vector such that the variance of the principal component scores is maximized for the new data matrix.

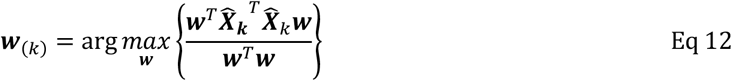

### 3.6. Membrane excitation and its cooperativity with intracellular ionic dynamics

When any of *G_x_*s is modified, the intracellular ion concentrations ([ion]_i_) change, although the variation is largely compensated for with time in intact cells through modification of the activities of both 3Na^+^/2K^+^ pump (NaK) and 3Na^+^/1Ca^2+^ exchange (NCX). In the present study, we imitated this long-term physiological homeostasis of [ion]_i_ by introducing empirical Eqs 13 and 14. These equations induced ‘negative feedback’ to the capacity (max*I_NaK_* and max*I_NCX_*) of these ion transporters. Namely, each correcting factor (*crf_x_*) was continuously scaled to modify the limiting activity of the transporters to keep the [Na^+^]_i_ or the total amount of Ca within the cell (Catot) equal to their pre-set level (*std_Nai_, std_Catot_*) with an appropriate delay (coefficients 0.3 and 0.008 in Eqs 13 and 14, respectively).

For the control of [Na^+^]_i_,

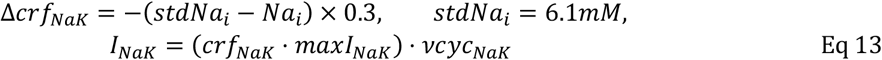

For the control of Ca_tot_,

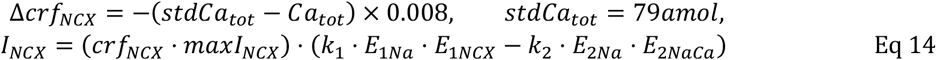

The Ca_tot_ is given by [Ca]_i_ included in the cytosolic three Ca-spaces *jnc, iz*, and *blk*, and in the sarcoplasmic reticulum *SR_up_* and *SR_rl_* in the free or bound forms, respectively.

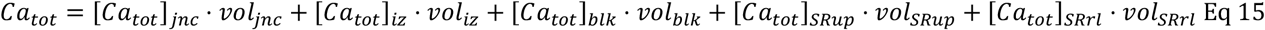

Here, the *vol* is the volume of the cellular Ca compartment (see more detail, [11]).

## 4. Results

### 4.1. Mapping the magnitude of MSE over the nine global parameter space

Parameter identifiability has been one of the central issues in the parameter optimization of biological models [14,20]. For confirmation of the identifiability of a unique set of solutions using the parameter optimization method, mapping of the MSE distribution is required over an enlarged parameter space defined by the *sf_x_* of the nine ionic currents of the baseline model. The randomization of *sf_x_* ranged from 1/10 to ~ 10 times the default values, and the calculation was performed for ~5,000,000 sets, as shown in Fig 1, where magnitudes of *log*(*MSE*) were plotted against each *sf_x_* on the abscissa.

**Fig 1.**
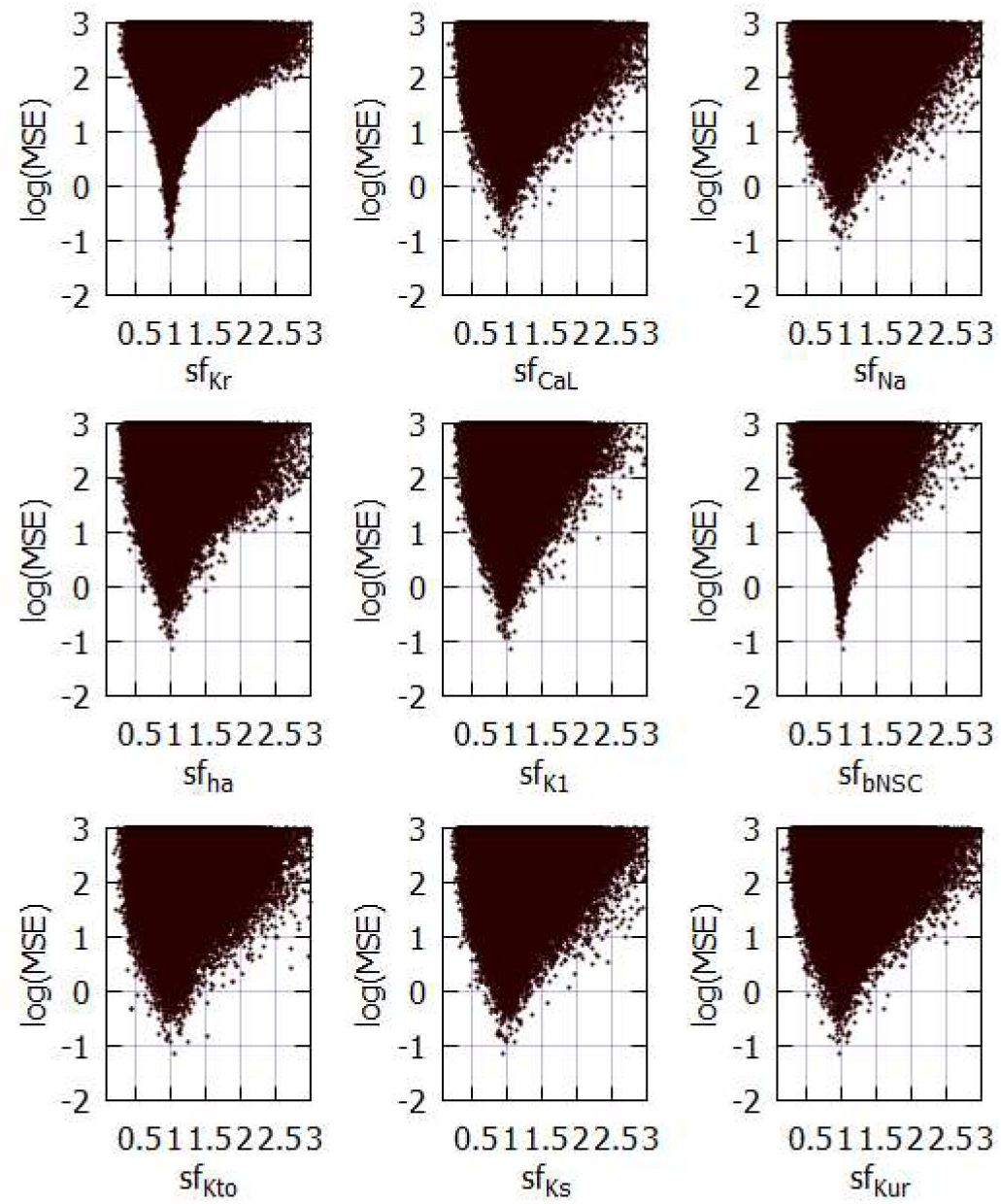
Distribution of MSE calculated between the target and the simulated APs modified by randomizing the *sf_x_* of 9 ionic currents in the coordinates of MSE-*sf_x_*. All MSE data points were plotted on the logarithmic ordinate against the linear *sf_x_*. A total of 5,141,382 points were calculated in cell model No.86 over the range of 1/10 ~ 10 times the default *sf_x_*. Since the configuration of *V_m_* records were largely unrealistic at *sfx* > 3, MSE points were cut out over *sf_x_* > 3.0. To demonstrate the sharp decrease in MSE, the data points were densely populated near the default *sf_x_*.

The data points of MSE at a given *sf_x_* include all variable combinations of the other eight *sf_x_*s. The algorithm of the PS method searches for a parameter set, which gives the minimum MSE at a given *stp* through the process of optimization. Although drawing a clear envelope curve by connecting the minimum MSEs at each *sf_x_* was difficult because of the insufficient number of data points in these graphs (Fig 1), an approximate envelope of the minimum MSEs may indicate a single global minimum of MSE located at the control *sf_x_* equals one, as typically exemplified by *I_Kr_*- and *I_bNSC_*-MSE relations. On both sides of the minimum, steep slopes of MSE/*sf_x_* are evident in all graphs. Outside this limited *sf_x_* -MSE area, the global envelope showed a gentle and monotonic upward slope toward the limit on the right side. No local minimum was observed in all of the *sf_x_* -MSE diagrams except the central sharp depression. It was concluded that the theoretical model of cardiac membrane excitation (hVC model) has only a single central sharp depression corresponding to the control model parameter.

### 4.2. The prompt necessity for a method of parameter optimization as indicated by hiPSC-CM APs

Fig 2 illustrates records of spontaneous APs (red traces) obtained in 12 experiments in the sequence of MDP (See Supplemental Materials for detail). All experimental records were superimposed with the simulated AP traces (black traces) obtained by the conventional manual fitting. In most cases, MSE of 1~6 mV^2^ remained (Eq 7) at the end of the manual fitting. This extra component of MSE might be largely attributed to slow fluctuations of *V_m_* of unknown origin in experimental recordings because the non-specific random fluctuations were quite different from the exponential gating kinetics of ion channels calculated in mathematical models. This extra-noise seriously interfered with the assessment of the accuracy of the parameter optimization of *G_x_* in the present study. Thus, APs produced by the manual adjustment (‘cell specific model’) was used as the target AP, which were completely free from the extra noise when examining the feasibility of the parameter optimization algorithm.

**Fig 2.**
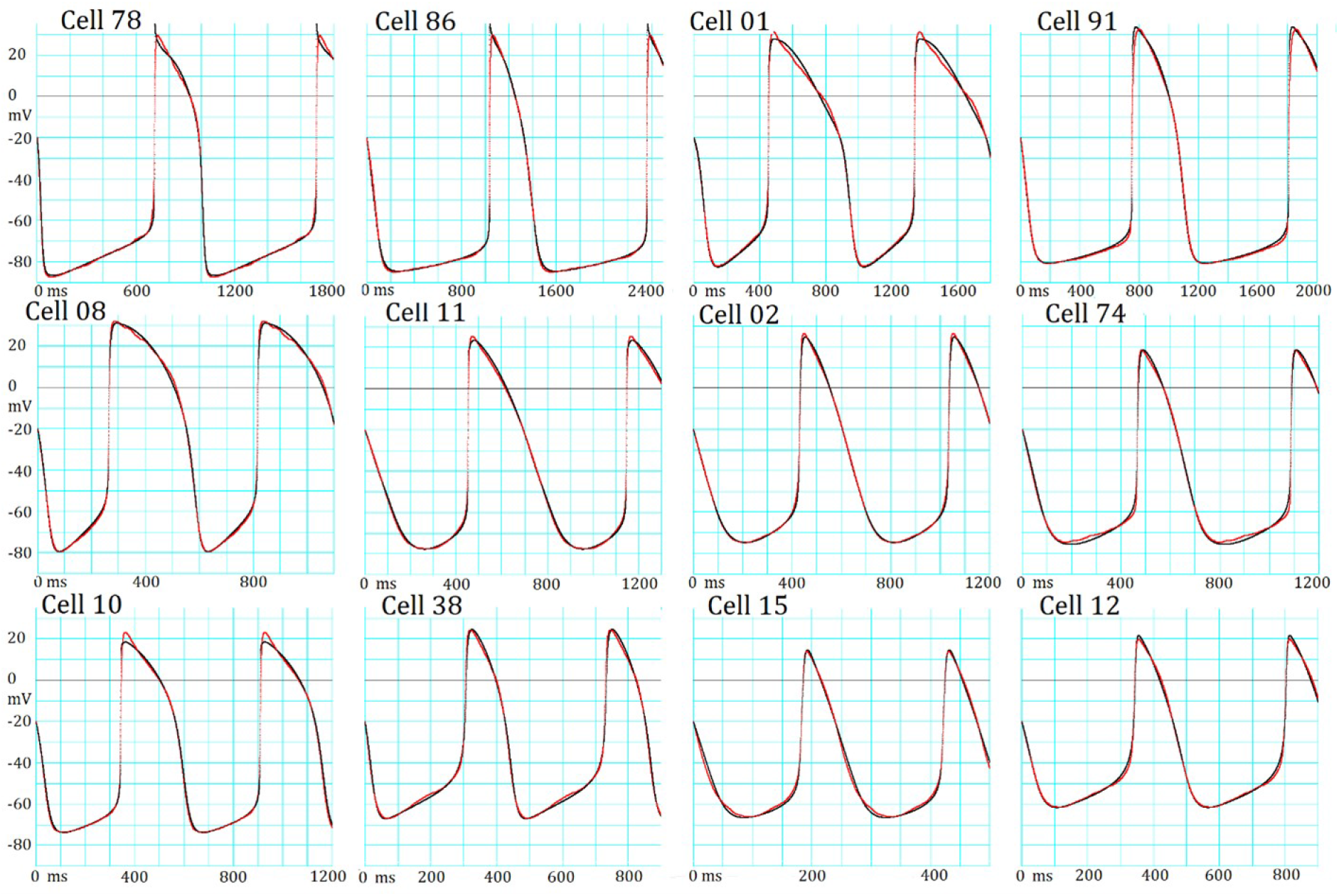
The manual fitting of variable AP configurations in 12 different hiPSC-CMs. Each panel shows the experimental record (red) superimposed by the model output (black) of the baseline model adjusted by the conventional manual fitting. At the top of each pair of AP records, the experimental cell number is presented. The extra fluctuations are obvious during the AP plateau in Cells 78, 08 and 01, while in Cells 15 and 74 during SDD. The length of abscissa is markedly different to illustrate the interval between two successive peaks of AP.

A comparison of AP configurations between these hiPSC-CMs clearly indicated that the classification of these APs into atrial-, ventricular- and nodal-types was virtually impractical, as described in [7]. On the other hand, if provided with the individual models fitted by an objective parameter optimizing tools using the baseline model (black trace), the results should be fairly straightforward not only in estimating the functional expression level of ion channels but also in clarifying the role of each current system or the ionic mechanisms in generating the AP configuration in a quantitative manner. Thus, the objective parameter optimization of the mathematical model is a vital requirement in cardiac electrophysiology.

Table 1 indicates the AP metrics; the cycle length (CL), the peak potential of the plateau (OS), the maximum diastolic potential (MDP), and the AP duration measured at −20 mV in addition to the MSE between individual experimental record and the model output fitted by manual fitting. The CL, MDP and AP were very variable among different AP recordings of cells shown in Fig 2. The cells were arranged by the sequence of MDP.

**Table 1.**
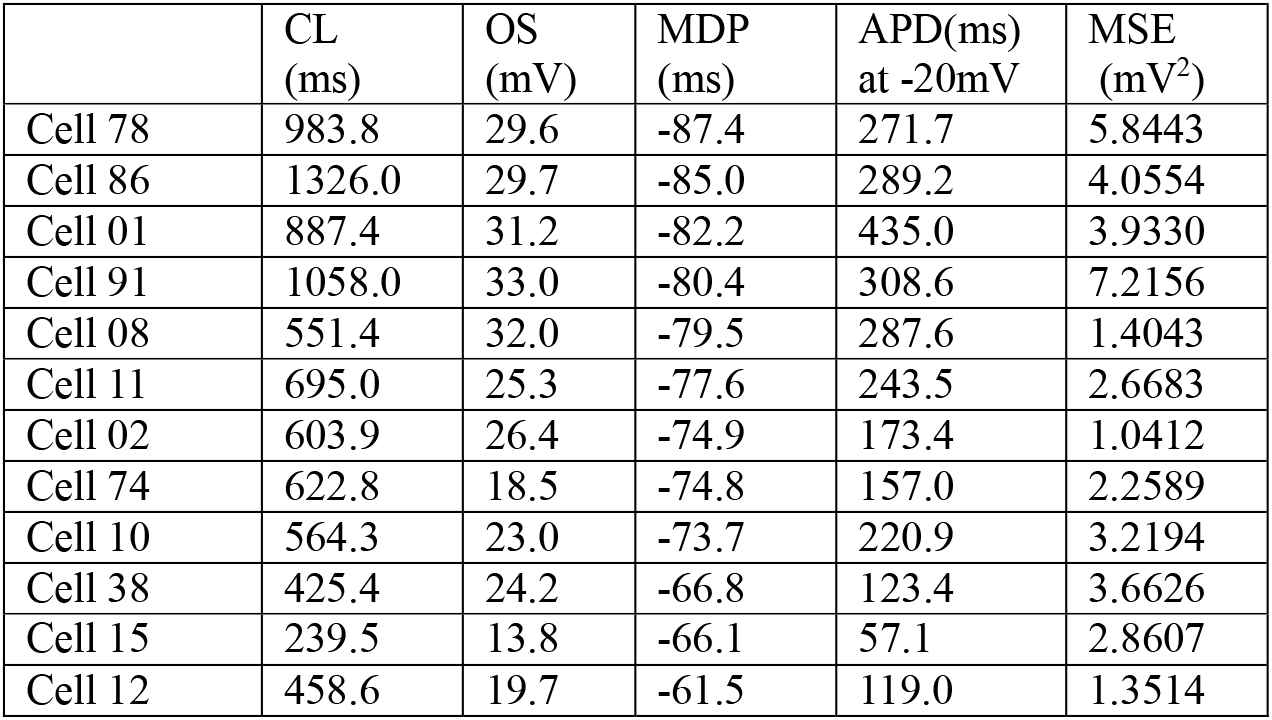
AP metrics and MSE calculated after the manual fitting of varying AP configurations in 12 different hiPSC-CMs in Fig 2. Table 1 indicates the AP metrics; the cycle length (CL), the peak potential of the plateau (OS), the maximum diastolic potential (MDP), and the AP duration measured at -20 mV in addition to the MSE between individual experimental record and the model output fitted by manual fitting. The CL, MDP and AP were very variable among different AP recordings of cells shown in Fig 2. The cells were arranged by the sequence of MDP.

|  | CL<br>(ms) | OS<br>(mV) | MDP<br>(ms) | APD(ms)<br>at -20mV | MSE<br>(mV<sup>2</sup>) |
| --- | --- | --- | --- | --- | --- |
| Cell 78 | 983.8 | 29.6 | -87.4 | 271.7 | 5.8443 |
| Cell 86 | 1326.0 | 29.7 | -85.0 | 289.2 | 4.0554 |
| Cell 01 | 887.4 | 31.2 | -82.2 | 435.0 | 3.9330 |
| Cell 91 | 1058.0 | 33.0 | -80.4 | 308.6 | 7.2156 |
| Cell 08 | 551.4 | 32.0 | -79.5 | 287.6 | 1.4043 |
| Cell 11 | 695.0 | 25.3 | -77.6 | 243.5 | 2.6683 |
| Cell 02 | 603.9 | 26.4 | -74.9 | 173.4 | 1.0412 |
| Cell 74 | 622.8 | 18.5 | -74.8 | 157.0 | 2.2589 |
| Cell 10 | 564.3 | 23.0 | -73.7 | 220.9 | 3.2194 |
| Cell 38 | 425.4 | 24.2 | -66.8 | 123.4 | 3.6626 |
| Cell 15 | 239.5 | 13.8 | -66.1 | 57.1 | 2.8607 |
| Cell 12 | 458.6 | 19.7 | -61.5 | 119.0 | 1.3514 |

The experimental study using the hiPSC-CMs was approved by the Kyoto University ethics review board (G259) and conformed to the principles of the Declaration of Helsinki.

### 4.3. Feasibility of the PS algorithm for parameter optimization of membrane excitation models

The automatic parameter optimization was applied to the model of cardiac membrane excitation in a limited number of studies (for review, see [23,26,35,36]) using various optimization methods, such as genetic algorithms. To the best of our knowledge, the principle PS algorithm has not been successfully applied to the detailed mathematic models of cardiac membrane excitation composed of both ionic channel and ion transporters models, except for the pioneering work in [12], which applied more general gradient-based optimization method to the simple ventricular cell model of Beeler and Reuter (BR model)[37].

Fig 3 shows a typical successful run of the new PS method in a hiPSC-CM, which showed an MDP of ~−85 mV. The PS parameter optimization was started after randomizing the *sf_x_s* of the major six currents, *I_Kr_, I_CaL_, I_Na_, I_ha_, I_Kl_* and *I_bNSC_*, in the manual fit model within a range of + 15% around the default values (normalized magnitude of 1). Fig 3A-1~3 compares the simulated *V_m,orp_* (black) with the target *V_m,t_* (red) at the repeat number N=1, 50 and 1167, respectively (Eq 8). The OS, APD as well as the CL of spontaneous AP were markedly different at the first cycle of AP reconstruction (Fig 3A-1). These deviations were largely decreased at the PS cycle (Fig 3A-2 *V_m_*, at N = 50), and became invisible in the final result (Fig 3A-3, N = 1167). The final individual current flow of nine current components are demonstrated in the lower panel of Fig 3A-3 (*I_m_*).

**Fig 3.**
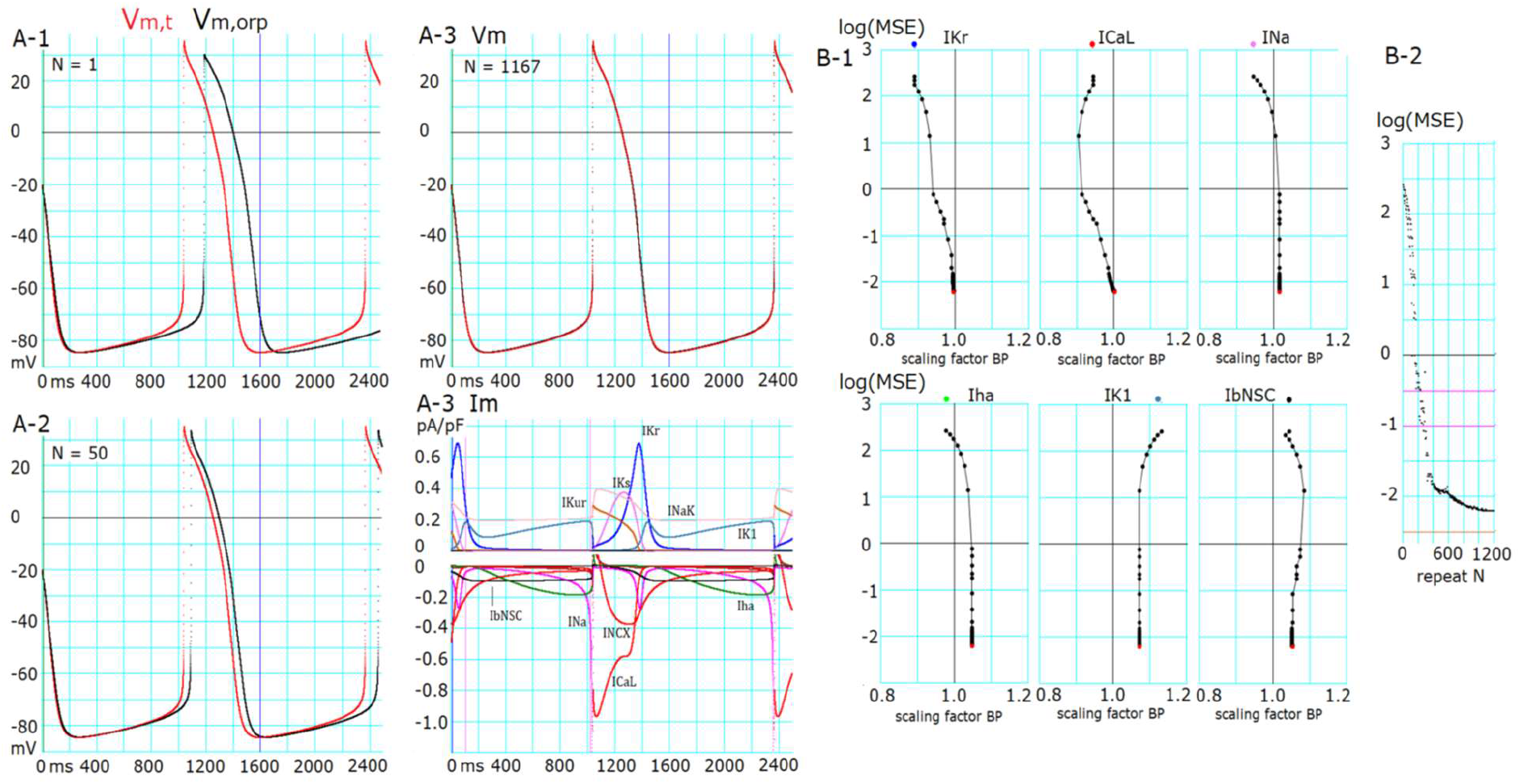
Results of the successful optimization in a cell (Cell86). (A-1) Target AP (*V_m,t_*, red) and AP generated by randomized initial *sf_x_*s (*V_m,orp_*, black). (A-2) *V_m,t_* (red) and *V_m,orp_* (black) generated after 50 cycles of adjusting BP. (A-3) *V_m_*: *V_m,t_* (red) and *V_m,orp_* (black) generated by the final *sf_x_*s. *I_m_*: corresponding time courses of each current for the finalized AP shown in A-3 *V_m_*. (B-1) Changes in *sf_x_*s vs. log(MSE) during a successful optimization process of PS. (B-2) log(MSE) of all BP points during the search process in PS. The initial values of *sf_x_*s are plotted by corresponding colors at the top of each *sf_x_*-log(MSE) graph.

The time course of decreasing *log*(*MSF*) evoked by the multi-run PS optimization is plotted for each *sf_x_* in Fig 3B-1 every time of resetting the set of base points. Fig 3B-2 shows all of the *log*(*MSF*) obtained at every adjustment by stepping individual BP points. The movement of all *sf_x_*s were synchronized to decrease *log*(*MSF*) from ~2.4 to 1 during the initial 180 cycles of decreasing *log*(*MSF*), but the search directions of BP were quite variable. The detailed adjustment of *sf_x_*s below *log*(_MSF_) < 0 was driven by adjusting *I_Kr_, I_CaL_* and *I_bNSC_* in this cell. The values of *sf_Kr_, sf_CaL_* and *sf_Na_* approached the correct value of 1, while those for *I_ha_, I_Kl_* and *I_bNSC_* remained deviated from the unit by less than 10% of the value. The explanation for the deviation of these three *sf_x_*s from the unit will be examined in the next section of the Results.

### 4.4. The six-parameter orp test successfully determined the conductance parameters of membrane excitation models

In individual runs, the PS optimization was frequently interrupted at intermediate levels during the progress of optimization and the probability of reaching *log*(*MSE*), for example, below −2, rapidly decreased with increasing extent of the randomization of the initial set of parameters. Moreover, the complementary relations between several ionic currents in determining d*V_m_*/dt might have hampered the parameter optimization. These facts indicate the requirement of statistical measures to improve the accuracy of the PS method. Fig 4 shows the results of orp tests, in which the optimization shown in Fig 3 was repeated several hundred times, and all results were plotted in a common coordinate of *log*(_*MSE*_) and individual *sf_x_*s. The population of *sf_x_* correctly converged at a single peak point very close to 1 with increasing negativity of *log*(*MSE*) for *sf_Kr_, sf_CaL_*, and *sf_Na_*, while *sf_ha_, sf_K1_*, and *sf_bNSC_* showed obvious variance. Nevertheless, they also showed a clear trend toward convergence to 1 in the average.

**Fig 4.**
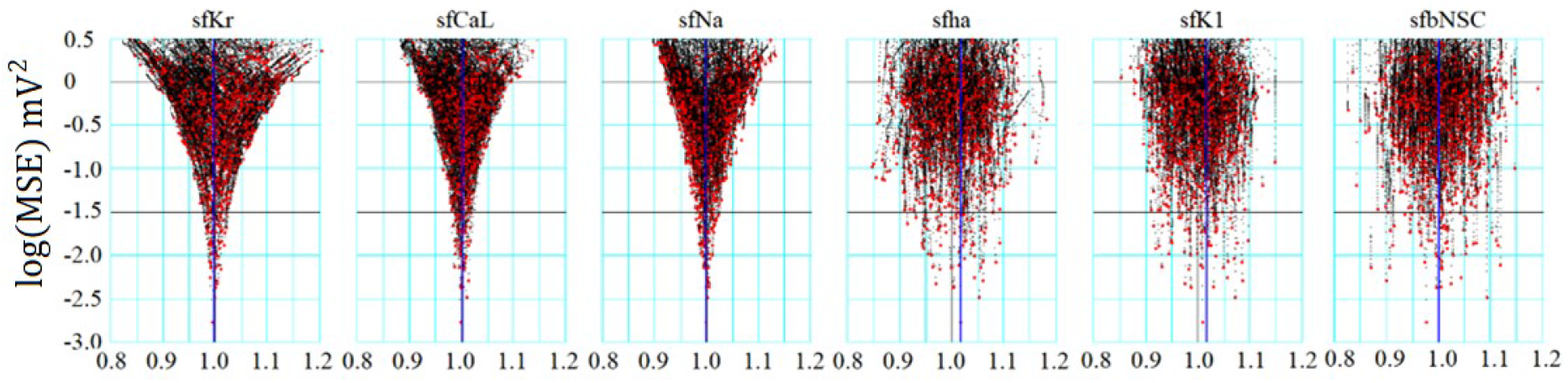
Convergence of *sf_x_* in the orp test for Cell86. The ordinate is the *log(MSE)* and the abscissa is the normalized amplitude of *sf_x_, x* stands for *Kr, CaL, Na, ha, K1*, and *bNSC*. Black points were obtained in the progress of optimization, and red ones are the final points in 829 runs of PS optimization.

Table 2 summarizes the mean of *sf_x_* determined for the top 20 runs of the PS parameter optimization in each of the 12 cells illustrated in Fig 2. The [Na^+^]_i_ as well as Catot was well controlled to the reference levels (*std_Nai_*, and *std_Catot_* in Eqs 16 and 17) of 6.1 mM and 79 amol, respectively, at the end of the parameter optimization to ensure the constant [Na^+^]_i_ as well as Ca_tot_. The mean of final *log*(*MSE*) = −2.74 indicates that the MSE was reduced by five orders of magnitude from the initial level just after the randomization by the orp test, like in the successful example shown in Fig 3B. The mean of individual *sf_x_*s were very close to 1 with a minimum standard error (SE) of mean, which were less than 1% of the mean, even for *I_K1_, I_bNSC_* and *I_ha_*, which showed weak convergence against *log*(*MSE*). These results well validate the accuracy of the parameter optimization using the multi-run PS method in all of 12 cell-specific models, which showed the large variety of spontaneous AP recorded in the hiPSC-CMs.

**Table 2.** Measurements of *sf_x_*s (mean + SE, n = 20), [Na^+^]_i_, mM and Ca_tot_ in amol in the 12 cells.

| Cell No. | log(MSE) | sfKr | sfK1 | sfCaL | sfbNSC | sfha | sfNa | sfKur | $[Na^+]_i$(mM) | $Ca_{tot}$(amol) |
| --- | --- | --- | --- | --- | --- | --- | --- | --- | --- | --- |
| 78 | -2.48321 | 1.00005 | 1.00157 | 1.00037 | 0.99460 | 1.00060 | 1.00134 |  | 6.10550 | 78.99979 |
| 91 | -2.42008 | 0.99952 | 1.00644 | 1.00063 | 1.00280 | 1.00470 | 1.00068 |  | 6.09977 | 79.00044 |
| 86 | -2.80257 | 1.00166 | 1.01394 | 1.00142 | 1.02670 | 1.00253 | 1.00702 |  | 6.09466 | 79.00008 |
| 01 | -2.79709 | 0.99871 | 1.00157 | 0.99756 | 0.99692 | 1.00054 | 0.99779 |  | 6.08973 | 78.99984 |
| 08 | -3.07432 | 0.00094 | 0.99982 | 1.00088 | 1.00041 | 0.99968 | 0.99985 |  | 6.12201 | 79.00056 |
| 11 | -2.67641 | 1.00186 | 1.00686 | 1.00129 | 0.99768 | 1.00253 | 1.01028 |  | 6.10385 | 78.99995 |
| 10 | -1.70278 | 1.00322 | 1.01081 | 1.00424 | 1.00396 |  | 0.99883 |  | 6.10968 | 79.00018 |
| 02 | -2.35441 | 1.00161 | 1.02038 | 1.00341 | 0.99815 | 1.01324 | 1.00954 |  | 6.10184 | 79.00004 |
| 74 | -2.43399 | 1.00126 | 1.01838 | 1.00308 | 0.99898 | 1.00004 | 1.00435 |  | 6.10118 | 79.99979 |
| 38 | -3.01883 | 1.00075 |  | 1.00106 | 1.00061 |  | 0.98866 | 1.00151 | 6.10530 | 78.99969 |
| 15 | -3.85992 | 1.00003 |  | 0.99894 | 0.99996 |  | 1.00015 | 0.98653 | 6.09902 | 79.00022 |
| 12 | -3.33037 | 0.99978 |  | 1.00030 | 0.99990 | 0.97587 |  | 1.00188 | 6.10012 | 79.00007 |
| Ave | -2.74617 | 0.99992 | 1.00886 | 1.00110 | 1.001723 | 0.99997 | 1.001681 | 0.99664 | 6.10272 | 79.08339 |
| SE | 0.07065 | 0.00093 | 0.010000 | 0.00164 | 0.00430 | 0.00729 | 0.00544 | 0.00690 | 0.00017 | 0.000345 |

The top 20 results obtained in the multi-run orp method were analyzed in each cell. Grand average (Ave) and SE are listed at the bottom rows.

### 4.5. Complementary relationship among *I_k1_, I_ha_* and *I_bNSC_*

Fig 5A illustrates the distribution of *sf_x_*s amplitude in the top 20 data points, where the final *sf_x_*s in individual runs were connected with lines for each run of PS in Cell 86 (Fig 2). The values of standard error (SE) of mean were quite small in the *sf_Kr_* and *sf_CaL_*, less than 1%. In contrast, *sf_ha_, sf_K1_* and *sf_bNSC_* showed evidently larger deviations. This finding is interesting since the former currents are mainly involved in determining the AP configuration and the latter group mainly in driving the relatively long-lasting SDD of approximately 1 sec in duration.

**Fig 5.**
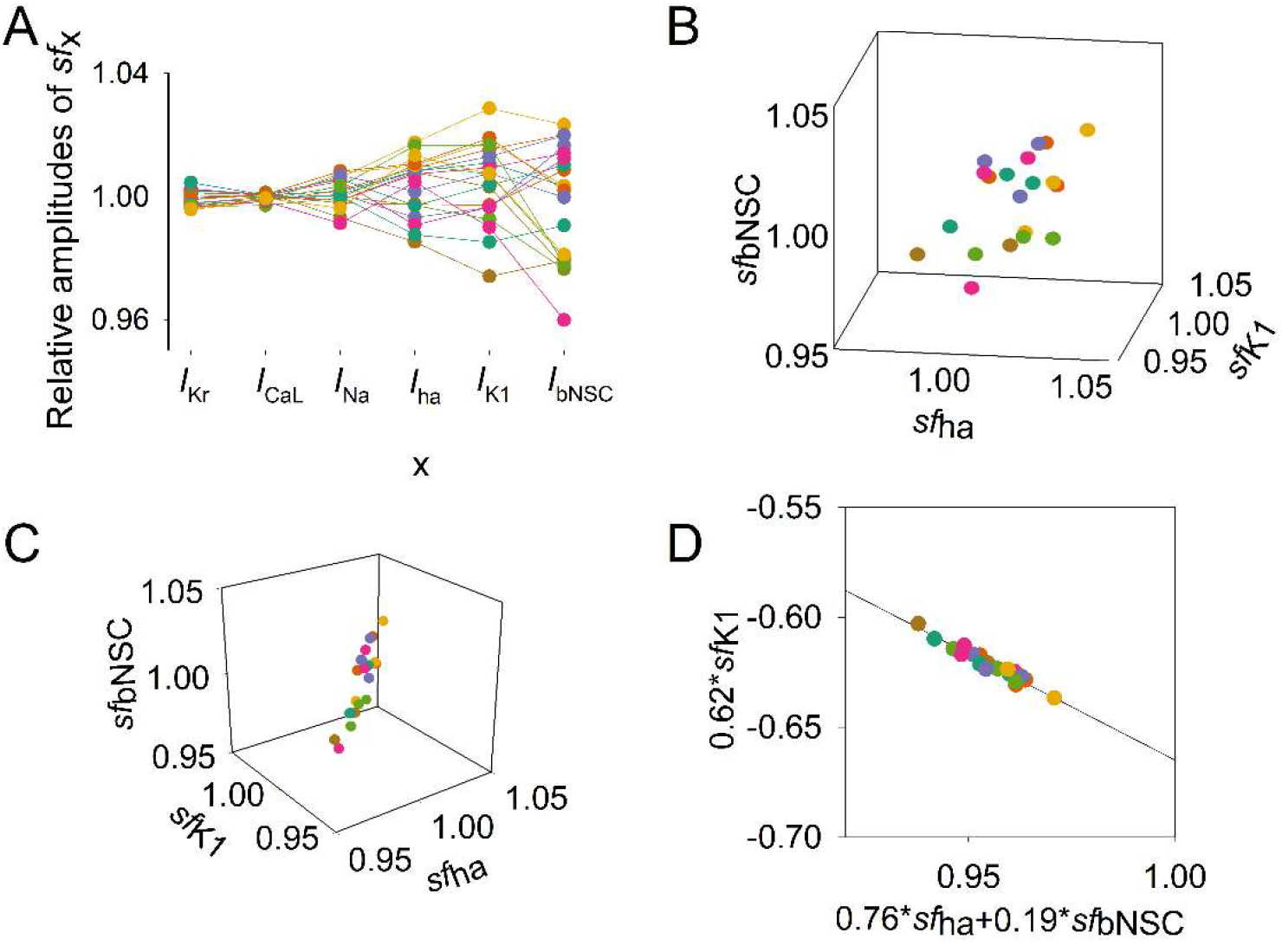
Distribution of *sf_x_* within the top 20 sets of *sf_x_*s obtained from the multi-run orp test in Cell86 in Fig 2. Data points of normalized *sf_x_* in each set were depicted in a different color. (A) amplitudes of each *sf_x_* (indicated on the abscissa) were plotted. (B) Three parameters, *sf_ha_, sf_K1_*, and *sf_bNSC_* were plotted in the 3D plot. (C) A different solid angle view of the 3D plot showed a linear correlation; see text for the plot in (D)

Thus, we analyzed the distribution of *sf_ha_, sf_K1_* and *sf_bNSC_* within the top 20 MSE. Fig 5B and C show the distribution of *sf_x_* points in the space of the three *sf_x_* dimensions. In Fig 5B, the 20 data points seemed to be dispersed randomly in the parameter space, but when the space was rotated to a specific angle, a linear distribution was observed as in Fig 5C, indicating that the points are distributed approximately on a plane surface in the 3D space. Using the multiple regression analysis, we could obtain an equation that fits the 20 data points as follows (R^2^=0.872);

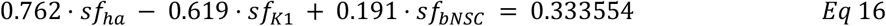

By replotting the data points in the 2D space with the abscissa for the sum of two inward-going currents (0.76 *sf_ha_* + 0.19 *sf_bNSC_*) and the ordinate for the outward current 0.62 *sf_K1_*, we obtained a regression line as shown in Fig 5D. The close correlations among the three *sf_x_*s were indicated with a quite large R^2^ of 0.941. This finding well confirms that the three currents have complementary relations with each other to give virtually identical configurations of spontaneous AP. In other words, *log*(*MSE*) remains nearly constant as far as the composition of the currents satisfies the relationship given by Eq 16.

The complementary relationship was further examined by conducting the orp test after fixing one of the two factors, *sf_K1_* or (*sf_ha_* + *sfb_NSC_*), illustrated in Fig 5B. Fig 6A shows the *log*(*MSE*) vs. *sf_K1_* relation when the (*sf_ha_* + *sf_bNSC_*) were fixed at the values obtained by the orp test. Indeed, the typical convergence of the *sf_K1_* was obtained. Alternatively, if the *sf_K1_* was fixed, the convergence was obviously improved for both *sf_ha_* and *sf_bNSC_* (Fig 6B-1, 2), but it was less sharp if compared to *sf_Kr_, sf_CaL_* and *sf_Na_* (not shown, but refer to corresponding results in Fig 4A). This finding was further explained by plotting the relationship between the *two* inward currents, *Iha* and *I_bNSC_*, as illustrated in Fig 6C. The regression line for the data points was fitted by Eq 17 with R^2^ = 0.86, supporting the complemental relationship between the two inward currents, *I_ha_* and *I_bNSC_*.

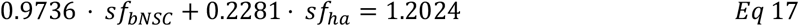

**Fig 6.**
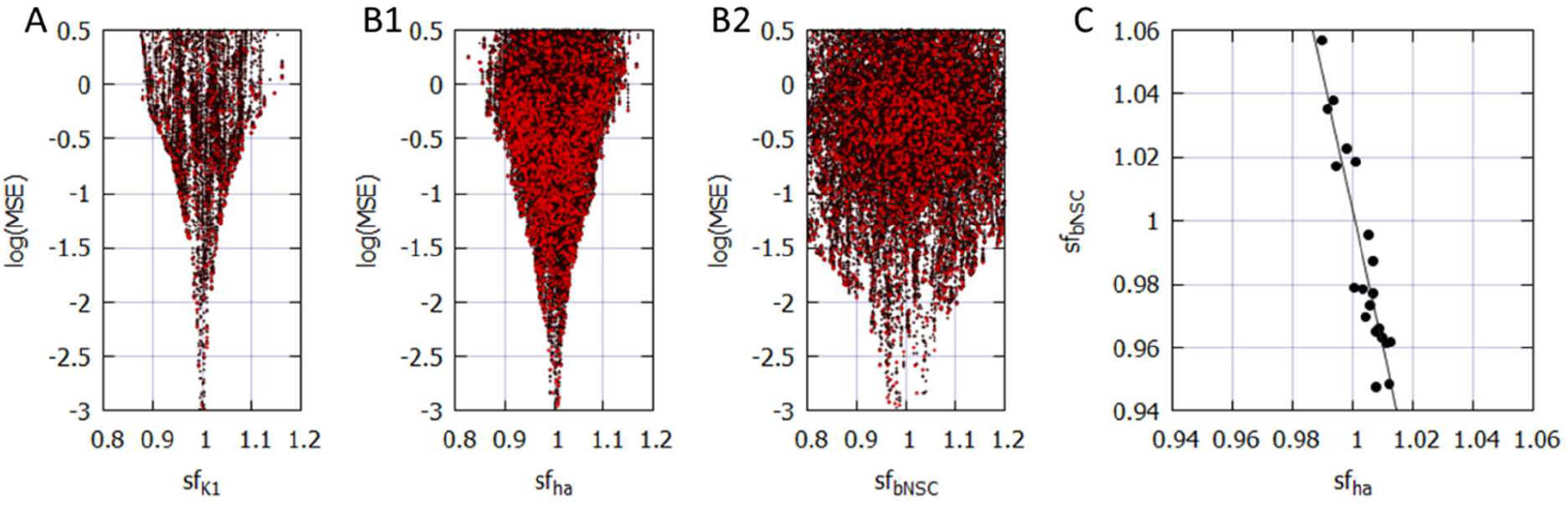
The complementary relations among *sf_K1_, sf_ha_* and *sf_bNSC_*. (A) and (B) results of the multi-run orp test. A; the perfect convergence of *sf_K1_* when *sf_ha_* and *sf_bNSC_* were fixed. (B1) improved convergence of *sf_ha_* and (B2) *sf_bNSC_* when *sf_K1_* was fixed. In these two orp tests, *sf_x_* of other currents showed quite comparable convergence as in Fig 4A. (C) the correlation between *sf_ha_* and *sf_bNSC_*.

The moderately high R^2^ indicates that the SDD is determined not only by the major *I_ha_* and *I_bNSC_ but* also by other currents, such as *I_K1_, I_Kr_*, the delayed component of *I_Na_* (*I_NaL_*) and *I_CaL_*, which were recorded during the SDD as demonstrated in Fig 3.

Essentially the same results of complementary relationship among *sf_ha_, sf_bNSC_* and *sf_K1_* were obtained in Cell 91, which also showed the long-lasting SDD with the very negative MDP as in Cell 86, as shown in Fig 2 and Table 2. The regression relation for the data points was fitted by Eqs 18 and 19 with R^2^ = 0.656 and 0.472, respectively.

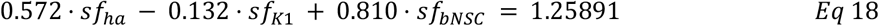

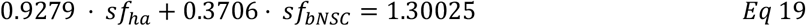

### 4.6. Principal components in the hiPSC-CM model

The PS frequently got stuck during the progress of parameter optimization and failed to reach the global minimum in the present study (Figs 4 and 6). The major cause of this interruption may most probably be attributed to the fact that *sf_x_*s were used directly as the search vector of the PS. In principle, the algorithm of PS parameter optimization gives the best performance when the parameters search is conducted in orthogonal dimensions where each dimension does not affect the adjustment of other *sf_x_* [28]. To get deeper insights, we applied the principal component (PC) analysis to the set of 6 *sf_x_*s selected in the baseline model. We performed PC analysis on the data points recorded in the vicinity of the minima (using the top 20 data).

As illustrated in Fig 7, each of the 6 PCs was not composed of a single *sf_x_* but mostly included multiple *sf_x_* sub-components. This finding indicates the inter-parameter interactions during the process of parameter optimization. For example, the changes in *sf_K1_* or *sf_bNSC_* simultaneously affect PCNo.1, 3, 6 or 1, 2, 3 PCs, respectively. Both *sf_CaL_* and *sf_Kr_* affect PCNo.4, 5. It might be concluded that the frequent interruptions of PS parameter optimization are most probably caused by the sporadic appearance of the local minima of MSE through interactions among *sf_x_*s.

**Fig 7.**
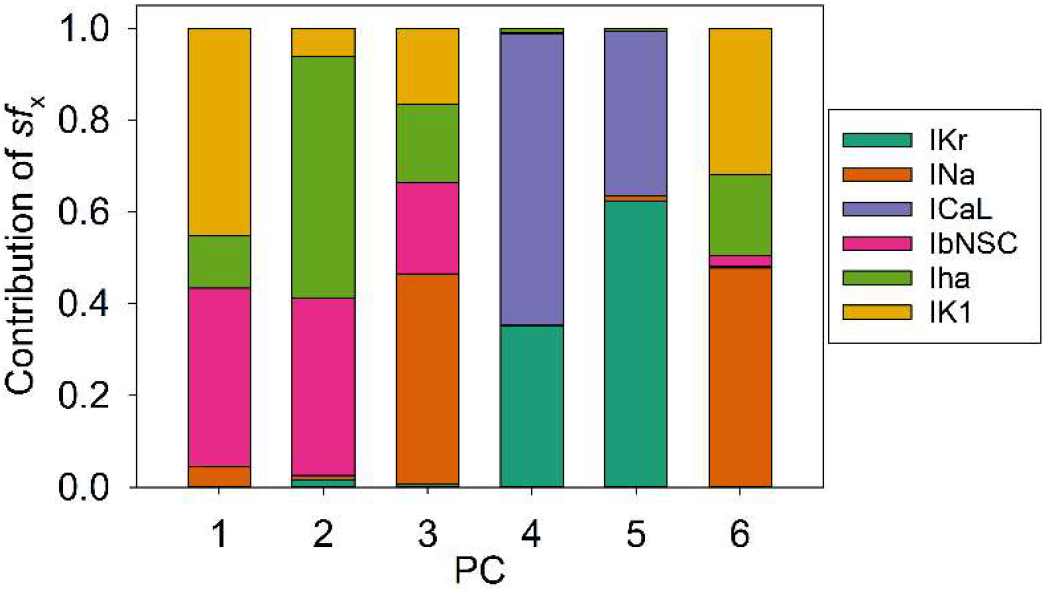
PC1~6 to describe distribution of the 6 *sf_x_*s. PC analysis was performed on the data population of the top 200 runs of the orp test as in Fig 4, which showed good optimization results (Cell 86). Each magnitude of 6 PCs was normalized to give a unit magnitude. Note each PC is composed of multiple components of ionic current, which are indicated in the Index with corresponding colors.

## 5. Discussion

New findings in the present study are listed below.

1. Mapping the MSE distribution over the enlarged parameter space was conducted by randomizing the *G_x_*s of the baseline model. It was confirmed that the baseline model has only a single sharp depression of MSE at the default *G_x_*s (Fig 1).
2. The preliminary cell-specific models were firstly prepared by the conventional manual tuning of *G_x_*s to superimpose the model output on each of twelve experimental AP recordings (Fig 2). Thereby, the parameter search space was restricted to a relatively small space to facilitate parameter optimization.
3. The *sf_x_*s of 4 ~ 6 *G_x_* parameters were initially assigned random values from a uniform distribution ranging between ± 10% of default values. The MSE was calculated between the randomized model output and the intact model AP as the target of optimization (Fig 3).
4. Plotting parameter *sf_x_* in a common *sf_x_* - MSE coordinates during each run of several hundred runs of optimization (Fig 4), we found that the *sf_x_* distribution of *I_Kr_, I_CaL_*, and *I_Na_* converged sharply to a single point with decreasing MSE, which exactly equaled the default ones. On the other hand, estimates of *sf_K1_, sf_ha_* and *sf_bNSC_* deviated slightly within a limited range around the default values in cells showing long-lasting SDD (Fig 4).
5. For statistical evaluation, the mean± SE of *sf_x_* in the top 20 estimates of MSE was calculated in individual cells (Table 2). The results of the parameter optimization in the 12 cells definitely indicated that the means of *sf_x_*s were very close to 1.00, with the SE less than 0.01 for all *G_x_*s.
6. A complementary relationship was found between *sf_K1_, sf_ha_* and *sf_bNSC_* in determining the gentle slope of long-lasting SDD in two representative cells (Fig 5). Supporting this view, the *sf_K1_* clearly focused on the unit provided that *sf_ha_* and *sf_bNSC_* were fixed and vice versa (Fig 6).
7. The six search vectors of *sf_x_* of the presented model could be replaced by the same number of theoretical PCs, and each PC was mostly composed of multiple *sf_x_*s (Fig 7). This finding definitely supports the view [12] that the complex interactions among *I_x_*s might interrupt the progress of the parameter optimization when *sf_x_*s were used as the search vector instead of using theoretical orthogonal ones.

The use of an initial randomized set of parameters was crucial in examining if an optimization method can determine unique estimates independent from the initial set of parameters, as used in the GA-based method for determining the *G_x_*s of the mathematical cardiac cell model [23]. The findings listed above well confirmed the feasibility of the PS method. Most probably, the PS method is applicable to variable mathematical models of other cell functions as well. See [26] for a more systematic review of the parameter optimization in the cardiac model development.

It has been suggested that different combinations of parameters may generate simple outputs that are very similar [12,23–25]. In the present study, this notion may be explained at least in part by the complementary relationship, for example, between the *I_K1_, I_ha_* and *I_bNSC_* in determining d*V_m_*/dt of SDD, which is a function of the total current (Eq 2, Figs 5 and 6). The gradient-based optimization method relies on the precise variation in the time course of d*V_m_*/dt induced by the time-dependent changes in individual *sf_x_*s (Eq 2). Therefore, the MSE was calculated over the whole time course of the spontaneous APs. Note, we did not use the AP metrics, which reflect only indirectly the kinetic properties of individual currents. Even with this measure of calculating the MSE, the time-dependent changes in *pO* (Eq 3) might be relatively small between two major currents, *I_K1_* and *I_ha_*, in comparison to *I_bNSC_*, which has no *V_m_*-dependent gate during the SDD as shown in the current profile Fig 3A-3. We assume that the gradient-based optimization method will be able to determine different contributions of individual currents if the optimization is conducted only within a selected time window of SDD. If MSE is calculated over multiple phases of the spontaneous AP, the influence of a particular phase on the MSE should be diluted. In our preliminary parameter optimization, this problem was partly solved by using a weighted sum for different phases of the spontaneous AP in summing up the MSE.

The small amplitude of a given current might be an additional factor in the weak convergence of *sf_x_* observed in the diagram of *sf_x_* - MSE in the orp test of optimization. If the current amplitude was much smaller in reference to the sum of all currents in determining d*V_m_*/dt (Eq 2), the resolution of the PS method would get lower. Sarkar et al. [24] demonstrated that the model output, for example, the AP plateau phase were almost superimposable when the different ratio of *G_Kr_* and *G_pK_* were used in reconstructing the model output (Figure 1 in [24]). They described that the AP metrics used for comparisons, such as APD, OS and APA seemed quite similar. It should be noted, however, that the results were obtained by applying different combinations of *sf_x_* to the same TNNP model [33]. This means that the relative amplitudes of *I_Kr_* and *I_pK_* in the TNNP model were much smaller than the major *I_CaL_* during the AP plateau, even though *I_Kr_* and *I_pK_* have totally different gating kinetics. Thus, the results of parameter optimization should be model-dependent. The same arguments will also be applied to the use of FR guinea pig model [38] in the study by Groenendaal et al. [23].

The gradient-based parameter optimization method was applied to the cardiac model of membrane excitation in [12], which analyzed the classic BR model [37]. The whole cell current in the BR model was composed of a minimum number of ionic currents, a background *I_K1_*, and three time-dependent currents; *I_Na_, I_s_*, and *I_x1_*, which were dissected from the voltage clamp experiments by applying the sucrose gap method to the multicellular preparation of ventricular tissue. The gatings of the latter three currents were formulated according to the Hodgkin-Huxley type gating kinetics, which was quite simple if compared with the recent detailed description of the ionic currents. They described that the parameter optimization was difficult if the AP configuration was used as the target of the parameter optimization, and they used the time course of the whole cell current as of the target of parameter optimization. However, the number of parameters was quite large, 63 in total, including limiting conductances and the gating kinetics. They suggested the feasibility of the parameter optimization method will be improved if provided with additional experimental data.

In the modern mathematical cardiac cell models, most ionic currents were identified by the whole-cell voltage clamp and single channel recordings in dissociated single myocyte [39] using the patch clamp technique [40] and by identifying their molecular basis of membrane protein. It has been clarified that the molecular basis of the ion channels expressed in the hiPSC-CMs is mostly identical to those in the adult cardiac myocytes rather than in the fetal heart (GSE154580 GEO Accession viewer (nih.gov)). Moreover, the gating kinetics have been much detailed to characterize the ionic currents within the cell model. In principle, the detailed characterization of individual currents should facilitate the identifiability of the model parameter but should not necessarily interfere with parameter optimization. We consider that the manual fitting of the model parameters to the AP recording by using a priori knowledge of biophysical mechanisms should largely facilitate the subsequent automatic parameter optimization. It might also be noted that the ionic currents left at the default values work as a kind of constraint to improve the identifiability of the target parameters.

After validating the automatic parameter optimization method, the final goal of our study is to find the principle of ionic mechanisms, which are applicable to the full range of variations of spontaneous AP records in both hiPSC-CMs and matured cardiomyocytes. For this purpose, we will apply the multi-run PS method to the experimental AP recordings using the initial parameter sets obtained by the conventional manual fit. The protocol of measuring the Gxs will be the same as used in the present study except for the use of experimental AP recordings in place of the output of the ‘cell-specific model’. In our preliminary analysis, the magnitude of individual model parameters obtained by the manual tuning was corrected by less than ~15% by the objective parameter optimization. Finally, the ionic mechanisms underlying the SDD of variable time courses will be analyzed in a quantitative manner, for example, by using the lead potential analysis [41], which explains changes in *V_m_* in terms of *G_x_* of individual currents.

### Limitations

In general, obvious limitations of the mathematical models of cardiac membrane excitation so far published are caused by a shortage of functional components inherent in intact cells. For example, the [ATP]i controlled by energy metabolism is a vital factor in maintaining the physiological function of ion channels as well as the active transport Na^+^/K^+^ pump [42]. Moreover, the followings are still not implemented in most models; the modulation of the ion channel activity through phosphorylation of the channel proteins, detailed modulation of the channel by the [Ca^2+^]_i_, the alterations of ion channel activity by PIP2 [43,44] and by the tension of the cell membrane through the cell volume change [45–48]. The detailed Ca^2+^ dynamics of the [Ca^2+^]_i_ are still not implemented in most of the cardiac cell models; such as the Ca^2+^ release from SR activated through the coupling of a few L-type Ca^2+^ channels with a cluster of RyRs at the dyadic junction [49], the Ca^2+^ diffusion influenced by the Ca^2+^-binding proteins [50]. To simulate the Ca^2+^-binding to troponin during the development of the contraction, it is necessary to include a dynamic model of contracting fibers [51–54]. These limitations should be thoroughly considered when pathophysiological phenomena, such as arrhythmogenesis are concerned. The scope of the present study is limited to the AP configurations of hiPSC-CMs, which were assumed to be ‘healthy’ with respect to the above concerns; for example, [ATP]_i_, [Na^+^]_i_ and Ca_tot_ were kept constant, and the standard contraction model was implemented as in the hVC model.

The parameter optimization presented in this study could be achieved in a practical way by limiting the number of unknown parameters. We determined only Gxs based on the assumption that ion channel kinetics are preserved as the same in the hiPSC-CMs as in the matured myocytes. Usually, 4~6 ionic currents were selected for the optimization. We found that the orp method could be performed simultaneously for all nine ionic currents described in Eq 1. However, the computation time was radically prolonged, and the resolution was not as high as obtained by using the modest number of parameters. We consider that the determination of the limited number of *G_x_*s is quite relevant to solving physiological problems in terms of detailed model equations for each current system.

Although *I_NCX_* and *I_NaK_* contribute sizeable fractions of the whole-cell outward and inward currents, respectively (Fig 3A-3), we excluded the scaling factors, *sf_NaK_* and *sf_NCX_* from the parameter optimization for the sake of simplicity. Instead, the possible drift of the intracellular ion concentrations was virtually fixed during the repetitive adjustment of ionic fluxes by varying *sf_x_* as shown in Table 2. The introduction of the empirical equations (Eqs 13 and 14) was quite useful to adjust the [Na^+^]_i_ and Ca_tot_ (Table 2) so that the time course as well as magnitude of *I_NCX_* remained almost constant during the parameter optimization. When influences of varying [Na^+^]_i_ and/or Ca_tot_ are examined under various experimental conditions in future, the reference levels of [Na^+^]_i_ and/or Ca_tot_ (*std_Nai_* and *std_Catot_* in Eqs 13 and 14) might be replaced by experimental measurements.

The parameter optimization method was not applied to several ionic currents. For example, it was difficult to determine the kinetics of T-type Ca^2+^ channel (*I_CaT_*; Ca_V_ 3.1) and excluded in the present study. A very fast opening and inactivation rates described in [55] suggest a complete inactivation of *I_CaT_* over the voltage range of SDD, while a sizeable magnitude of window current described in [56] suggests a much larger contribution to SDD. The kinetics of *I_CaT_* still remain to be clarified in experimental examinations. The sustained inward current, *I_st_*, is recently attributed most probably to the Ca_V_ 1.3 [57], which is activated at a more negative potential range than the activation of *I_CaL_* (Ca_V_ 1.2) [58,59]. The *I_bNSC_* was used to represent net background conductance in the present study. However, several components of the background conductance have been identified on the level of molecular basis in matured myocytes (see for review TRPM4, [60]). Experimental measurements of the current magnitude for each component are also awaited.

Gábor and Banga indicated that the multi-run method had shown good performance in certain cases, especially when high-quality first-order information is used and the parameter search space is restricted to a relatively small domain [16] (see also [19]). Indeed, the manual fitting of the parameters (Fig 1) was required to utilize the presented multi-run PS method over the restricted search space. One of the major difficulties in the manual fitting of individual Gxs arose during the SDD, where *I_Kr_, I_K1_, I_bNSC_*, and *I_ha_*, in addition to *I_NaK_* and *I_NCX_* constitute the whole-cell current (Fig 3A-3). Close inspection of the current components (Fig 3A-3), however, suggests hints of how to do with the manual fit. The transient peak of *I_Kr_* dominates the current profile during the final repolarization phase from −20 to −60 mV in all 12 hiPSC-CMs [61], since *I_CaL_* and *I_Ks_* rapidly deactivated before repolarizing to this voltage range. The *I_NaK_* and *I_NCX_* are well controlled by the extrinsic regulation in Eqs 13 and 14. Thus, the manual fitting of *sf_Kr_* is firstly applied to determine *sf_Kr_*. The MDP more negative than −70 mV is adjusted by the sum of time-dependent (*I_Kr_* + *I_K1_*) and the time-independent *I_bNSC_*. Then, *I_Kr_* is deactivated when depolarization becomes obvious after the MDP, and gradual activation of *I_ha_* and the depolarization-dependent blocking of *I_K1_* by the intracellular substances [62] take the major role in promoting the initial linear phase of SDD. Thus, the amplitude of *sf_K1_* and *sf_bNSC_* might be approximated during the initial half of SDD. The late half of SDD, including the foot of AP, namely the exponential time course of depolarization toward the rapid rising phase of AP, is mainly determined by the subthreshold *V_m_*-dependent activation of *I_Na_* (after MDP more negative than −70 mV) and/or *I_CaL_* (after MDP less negative than −65 mV). Thus, the *sf_Na_* and *sf_CaL_* are roughly determined by fitting the foot of AP and the timing of the rapid rising phase of AP. The plateau time course of AP is determined by *sf_CaL_* and the factor of Ca^2+^-mediated inactivation of *I_CaL_* (the parameter *KL*, [4]). Since the kinetics of outward currents, *I_Kur_, I_Kto_ (endo-type)*, and *I_Ks_* are quite different from that of *I_Kr_*, the plateau configuration is determined bit by bit by adjusting these currents. We failed to observe the phase 1 rapid and transient repolarization in the hiPSC-CMs (Fig 2), which is the typical sign of the absence of epicardial-type *I_Kto_*.

In hiPSC-CMs showing less negative MDP than ~-65 mV, the contribution of *I_K1_, I_Na_* and *I_ha_* should be negligibly small because *I_K1_* is nearly completely blocked by the intracellular Mg^2+^ and polyamine, *I_Na_* is inactivated, and *I_ha_* is deactivated during SDD, even if any expressed.

Nevertheless, parameter optimization might be laborious and time-consuming for those who are not familiar with the electrophysiology of the cardiac myocyte. This difficulty might be largely eased by accumulating both AP configurations and the underlying current profile obtained in the parameter optimization into a database in the future. If this database becomes available, the computer may search for several candidate APs for the initial parameter set, which is used for automatic parameter optimization.

## Supporting information

APdata

optimizing program

## Abbreviations

hiPSC-CMs: human induced pluripotent stem cell-derived cardiomyocytes
AP: action potential
MDP: the maximum diastolic potential
SDD: slow diastolic depolarization
*I*_m_: membrane current
*V*_m_: membrane voltage
orp: optimization of randomized model parameters
PS: pattern search method
BP: base point for searching minimum MSE in the Pattern Search
NP: searching point in reference to BP in the Pattern Search
MSE: mean square error between two different *V_m_* records
stp: step size to move NP
x: subscript to represent membrane current such as *I_Na_, I_CaL_, I_K1_, I_ha_, I_Kr_, I_Kur_, I_Ks_* and *I_bNSC_*

## 6. Funding and financial conflicts of interest

The authors declare that the research was conducted without any commercial or financial relationships that could be construed as a potential conflict of interest.

This work was supported by JSPS KAKENHI (Grant-in-Aid for Young Scientists) Grant Numbers JP19K17560 for HK, 16K18996 for YH, and 21K06781 for FT.

## 7. Author Contributions

HK, YH, DY, YW, AK, and TM performed the wet experiments and analyzed them. HK, SK, YH, YZ, FT, AA, and AN developed the simulation model and the parameter optimization method. HK, YH, AA and AN wrote the manuscript. All authors reviewed the manuscript. AA and TK organized the research team.

## 8. Acknowledgments

The authors would like to thank our laboratory colleagues for their valuable comments and discussions.

## Notes

http://www.eheartsim.com/en/downloads/

### Competing Interest Statement

The authors have declared no competing interest.

### Summary of Updates

small revision to correct mistakes.

