## Supplemental Materials for "Gradient-based parameter optimization to determine membrane ionic current composition of human induced pluripotent stem cell-derived cardiomyocytes"

The source code of the computer model is available at http://www.eheartsim.com.

### hiPSC-CM

#### Preparation of dissociated hiPSC-CMs and recording of spontaneous APs

201B7 and 253G1 hiPSC lines generated from a healthy individual were used in this study (Takahashi et al. 2006, Nakagawa et al. 2008). Differentiation of hiPSCs into cardiomyocytes was promoted using an embryoid body (EB) differentiating system (Yang et al. 2008). hiPSC were incubated at 37 ºC in 5% $CO_2$, 5% $O_2$, and 90% $N_2$ for the first 12 days to promote differentiation. The hiPSCs aggregated to form EBs and were cultured in suspension for 20 days. On the 20th day of culture, EBs were treated with collagenase B (Roche, Basel, Switzerland) and trypsin EDTA (Nacalai Tesque, Kyoto, Japan) and dispersed into single cells or small clusters, which were plated onto 0.1% gelatin-coated dishes. hiPSC-CMs were then maintained in a conditioned medium. The experimental study using the hiPSC-CMs was approved by the Kyoto University ethics review board (G259) and conformed to the principles of the Declaration of Helsinki.

#### Electrophysiological recordings of hiPSC-CM APs

For single-cell patch-clamp recordings, gelatin-coated glass coverslips were placed into each well of a 6-well plate, and 2 ml of DMEM/F12 containing 2% FBS and 80,000-120,000 CMs were added in each well. Spontaneous APs were recorded from beating single CM using the perforated patch-clamp technique with amphotericin B (Sigma-Aldrich) at 36 ± one ºC. Data were acquired at 20 kHz with the Multiclamp 700B amplifier (Molecular Devices, Sunnyvale, CA, USA), Digidata 1440 digitizer hardware (Molecular Devices), and pClamp 10.4 software (Molecular Devices). The glass pipettes had a resistance of 3-6 MΩ after filling them with the intracellular solution. The external solution used for AP recordings was composed of the following composition (in mM): NaCl 150, KCl 5.4, $CaCl_2$ 1.8, $MgCl_2$-$6H_2O$ 1, glucose 15, HEPES 15, and Na-pyruvate 1; pH was adjusted to 7.4 by titrating NaOH. Intracellular solution contained (in mM): KCl 150,

NaCl 5, $CaCl_2$ 2, EGTA 5, MgATP 5, HEPES 10 (pH adjusted to 7.2 with KOH), and amphotericin B 300 μg/ml.

### Abbreviations

*Table S1*. Abbreviations in model equations

| | |
|---|---|
| $V_m$ | membrane potential (mV) |
| $I_{tot\_cell}$ | total current of ion channels and exchangers (pA/pF) |
| $I_{tot\_x\_a}$ | total current of ion 'x' channels and exchangers at space 'a' (pA/pF) |
| $I_{app}$ | current applied through a patch electrode (pA/pF) |
| $E_{rev,x}$ | reversal potential of current 'x' (mV) determined from the tangential line of the $I_x$ - $V$ curve |
| $C_m$ | whole cell membrane capacitance (pF) |
| $G_x$ | conductance of current 'x' (pA /pF/mV) |
| $k, \alpha, \beta$ | rate constants ($ms^{-1}$ or $mM^{-1}ms^{-1}$) |
| $P_{I(\_X)}$ | converting factor of $GHK_X$ from mM/ms to pA/mM/pF |
| $v_{cyc\_x}$ | turnover rate of transporter 'x' ($ms^{-1}$) |
| $V_a$ | total volume of space 'a' (fL) |
| $[X_{tot}]_a$ | total concentration of substance 'X' at space 'a' (mM) |
| $[X_{free}]_a$ | concentration of free substance 'X' at space 'a' (mM) |
| $[X]_a$ | concentration of 'X' at space 'a' (mM) |
| $J_X$ | total flux of ion 'X' (attomol/ms) |
| $\frac{d[X]_a}{dt}$ | rate of change of 'X' concentration at space 'a' (mM/ms) |

### Model Parameters

*Table S2*. Physical constants

| | | |
|---|---|---|
| R | 8.3143 | C・mV/mmol/K |
| T | 310.15 | K |
| F | 96.4867 | C/mmol |

*Table S3*. Ion concentrations

| | | |
|---|---|---|
| $[K^+]_o$ | 5.4 | mM |
| $[Na^+]_o$ | 145 | mM |
| $[Ca^{2+}]_o$ | 1.8 | mM |

### GHK equation

The magnitudes of ion channel currents are described either by the Ohmic equation or by the GHK equation. In the latter case, the term to convert mM to pA (permeability times, *zF*) in the original GHK equation is represented by a lumped converting factor, P in a unit of pA/pF $mM^{-1}$, because of unknown total number of channels within a cell and single channel conductance. Then, the amplitude of current (I) for an ion X is given by,

$$I = P \cdot pO \cdot GHK_X \qquad Eq.S1$$

Where $GHK_X$ is,

$$GHK_X = \frac{Z_X F V_m}{RT} \cdot \frac{\left([X]_i - [X]_o \cdot exp\left(\frac{-Z_X F V_m}{RT}\right)\right)}{\left(1 - exp\left(\frac{-Z_X F V_m}{RT}\right)\right)} \qquad Eq.S2$$

### Nernst equation

$$E_X = \frac{RT}{z_X F} \cdot ln\left(\frac{[X]_o}{[X]_i}\right) \qquad Eq.S3$$

*Table S4.* Volume composition of cytosol in comparison with hVC model (Asakura et al., 2014, Himeno et al., 2015)

| | hiPSC-CM | hVC model |
|---|---|---|
| Input capacitance | 32 pF | 192.46 pF |
| Cell volume ($V_{cell}$) | 2510 fL | 37920 fL |
| Bulk space ($V_{blk}$) | 65% of $V_{cell}$ fL | 68% of $V_{cell}$ fL |
| Intermediate zone ($V_{iz}$) | 3.5% of $V_{cell}$ fL | 3.5% of $V_{cell}$ fL |
| Junctional space ($V_{jnc}$) | 0.8% of $V_{cell}$ fL | 0.8% of $V_{cell}$ fL |
| Total SR space ($V_{SR}$) | 1.1% of $V_{cell}$ = 27.61 fL | 6% of $V_{cell}$ fL |
| Volume of SR releasing site ($V_{SRrl}$) | 10% of $V_{SR}$ fL | 20% of $V_{SR}$ fL |
| Volume of SR releasing site ($V_{SRup}$) | 90% of $V_{SR}$ fL | 80% of $V_{SR}$ fL |

### $Ca^{2+}$ buffer

The detailed set of buffer species used in the GPB model (2010) (Grandi et al., 2010) was adopted after several simplifications as described in our previous paper (Asakura et al., 2014). In short, we deleted the myosin, $Na^{+}$ and $Mg^{2+}$ buffers, and fixed $[Mg^{2+}]$. The low affinity binding of $Ca^{2+}$ to troponin (TnCl) was replaced by a contraction model (Negroni and Lascano, 2008) and the amount of the high affinity site (TnCh) was adjusted.

#### Bulk space (blk)

$$\frac{d[CaMCa]}{dt} = k_{on\_CaM} \cdot [Ca^{2+}]_{blk} \cdot ([B_{total}CaM] - [CaMCa]) - k_{off\_CaM} \cdot [CaMCa] \quad Eq.S4$$

$$k_{off\_CaM} = 0.0238, k_{on\_CaM} = 3.4, [CaM_{tot}] = 0.024 \quad Eq.S5$$

$$\frac{d[TnChCa]}{dt} = k_{on\_TnCh} \cdot [Ca^{2+}]_{blk} \cdot ([B_{tot}TnCh] - [TnChCa]) - k_{off\_TnCh} \cdot [TnChCa] \quad Eq.S6$$

$$k_{off\_TnCh} = 0.000032, k_{on\_TnCh} = 2.37, [TnCh_{tot}] = 0.007 \quad Eq.S7$$

$$\frac{d[SRCa]}{dt} = k_{on\_SR} \cdot [Ca^{2+}]_{blk} \cdot ([B_{tot}SR] - [SRCa]) - k_{off\_SR} \cdot [SRCa] \quad Eq.S8$$

$$k_{off\_SR} = 0.006, k_{on\_SR} = 10, [SR] = 0.0171 \quad Eq.S9$$

#### Intermediate zone (iz)

$$[L_{free}]_{iz} = \frac{[B_{tot}L]_{iz}}{1 + \frac{[Ca^{2+}]_{iz}}{K_{dL\_iz}}}, [B_{tot}L] = 0.6078 \quad Eq.S10$$

$$K_{dL\_iz} = \frac{k_{off\_L\_iz}}{k_{on\_L\_iz}}, k_{off\_L\_iz} = 1.3, k_{on\_L\_iz} = 100 \quad Eq.S11$$

$$\left[H_{free}\right]_{iz} = \frac{[B_{tot}H]_{iz}}{1 + \frac{[Ca^{2+}]_{iz}}{K_{dH\_iz}}}, [B_{tot}H] = 0.2178 \qquad Eq.S12$$

$$K_{dH\_iz} = \frac{k_{off\_H\_iz}}{k_{on\_H\_iz}}, k_{off\_H\_iz} = 0.03, k_{on\_H\_iz} = 100 \qquad Eq.S13$$

$$[Ca^{2+}]_{iz} = \frac{\left[Ca_{tot}\right]_{iz}}{1 + \frac{[Lf]_{iz}}{K_{dL\_iz}} + \frac{[Hf]_{iz}}{K_{dH}}} \qquad Eq.S14$$

**Junctional space (jnc)**

$$\left[L_{free}\right]_{jnc} = \frac{[B_{tot}L]_{jnc}}{1 + \frac{[Ca^{2+}]_{jnc}}{K_{dL\_jnc}}}, [B_{tot}L] = 1.1095 \qquad Eq.S15$$

$$K_{dL\_jnc} = \frac{k_{off\_L\_jnc}}{k_{on\_L\_jnc}}, k_{off\_L\_jnc} = 1.3, k_{on\_L\_jnc} = 100 \qquad Eq.S16$$

$$\left[H_{free}\right]_{jnc} = \frac{[B_{tot}H]_{jnc}}{1 + \frac{[Ca^{2+}]_{jnc}}{K_{dH\_jnc}}}, [B_{tot}H] = 0.398 \qquad Eq.S17$$

$$K_{dH\_jnc} = \frac{k_{off\_H\_jnc}}{k_{on\_H\_jnc}}, k_{off\_H\_jnc} = 0.03, k_{on\_H\_jnc} = 100 \qquad Eq.S18$$

$$[Ca^{2+}]_{jnc} = \frac{\left[Ca_{tot}\right]_{jnc}}{1 + \frac{[Lf]_{jnc}}{K_{dL\_jnc}} + \frac{[Hf]_{jnc}}{K_{dH}}} \qquad Eq.S19$$

**Release site of the SR (SRrl)**

$$k_{off\_CSQN} = 65, k_{on\_CSQN} = 100, [B_{tot}CSQN] = 10 \qquad Eq.S20$$

$$K_{d\_CSQN\_Ca} = \frac{k_{off\_CSQN}}{k_{on\_CSQN}} \qquad Eq.S21$$

$$a = 1 \qquad Eq.S22$$

$$b = [B_{tot}CSQN] - [Ca_{tot}]_{SRrl} + K_{d\_CSQN\_Ca} \qquad Eq.S23$$

$$c = -K_{d\_CSQN\_Ca} \cdot [Ca_{tot}]_{SRrl} \qquad Eq.S24$$

$$[Ca^{2+}]_{SRrl} = \frac{-b + \sqrt{b^2 - 4ac}}{2a} \qquad Eq.S25$$

### Boundary $Ca^{2+}$ diffusion

#### $Ca^{2+}$ transfer between cytosolic compartments

$$J_{Ca\_jnciz} = G_{dCa\_jnciz} \cdot \left([Ca^{2+}]_{jnc} - [Ca^{2+}]_{iz}\right) \qquad Eq.S26$$

$$G_{dCa\_jnciz} = 32158\ (fL \cdot ms^{-1}) \qquad Eq.S27$$

$$J_{Ca\_izblk} = G_{dCa\_izblk} \cdot ([Ca^{2+}]_{iz} - [Ca^{2+}]_{blk}) \qquad Eq.S28$$

$$G_{dCa\_izblk} = 2076.1\ (fL \cdot ms^{-1}) \qquad Eq.S29$$

#### Ca2+ transfer from SR uptake site to release site

$$J_{trans\_SR} = P_{trans} \cdot \left([Ca^{2+}]_{SRup} - [Ca^{2+}]_{SRrl}\right) \qquad Eq.S30$$

$$P_{trans} = 0.017(fL \cdot ms^{-1}) \qquad Eq.S31$$

### Ion channels and transporters

#### L-type $Ca^{2+}$ current ($I_{CaL}$, LCC)

According to the scheme of (Shirokov et al., 1993) and (Ferreira et al., 1997), the same 4-state model was used for both LCCs in CaRU ($I_{CaL\_jnc}$) and for LCCs located in *blk* ($I_{CaL\_blk}$) and *iz* ($I_{CaL\_iz}$). The description of both $V_m$-dependent gate and $[Ca^{2+}]$-dependent gates in hVC model was used in the hiPSC-CM model after minor modification. The rate constants for the $V_m$-gate ($\alpha_+$ and $\alpha_-$) and $Ca^{2+}$-gate ($\varepsilon_+$ and $\varepsilon_-$) of LCC are given by *Eqs. S35, S36* and *Eqs. S37, S38*, respectively. Both activation ($\alpha_+$) and deactivation ($\alpha_-$) rates of the $V_m$-gate were described as a function of two exponential terms and adjusted to hiPSC-CM data.

$$I_{CaL\_X\_a} = f_{CaL\_a} \cdot P_{CaL\_X} \cdot GHK_{X\_a} \cdot pO_{LCC\_a} \cdot \frac{1}{1+\left(\frac{1.4}{[ATP]}\right)^3} \qquad Eq.S32$$

[ATP] was fixed to 6 mM.

### Fraction of $I_{CaL}$

$$f_{CaL\_jnc} = 0.15,\ f_{CaL\_blk} = 0.45,\ f_{CaL\_iz} = 0.40 \qquad Eq.S33$$

### Converting factors

$$P_{CaL\_Ca} = 5.068,\ P_{CaL\_Na} = 0.0000185 \cdot P_{CaL\_Ca},\ P_{CaL\_K} = 0.000367 \cdot P_{CaL\_Ca}\ (pA/pF/mM)$$

The rate constants for the $V_m$-gate,

$$v = V_m - V_{shift} \qquad \text{Eq.}S34$$

$$\alpha_+ = \frac{1}{0.763 \cdot exp\left(-\frac{v}{8.5}\right) + 0.348 \cdot exp\left(-\frac{v}{3500}\right)} \qquad Eq.S35$$

$$\alpha_- = \frac{0.5}{4.65 \cdot exp\left(\frac{v}{15}\right) + 1.363 \cdot exp\left(\frac{v}{100}\right)} \qquad Eq.S36$$

The rate constant ($\varepsilon_+$) for the $Ca^{2+}$-inactivation.

$$\varepsilon_+ = \frac{0.35 \cdot [Ca^{2+}]_{nd} \cdot \alpha_+}{T_L \cdot K_L} \qquad Eq.S37$$

The values of $T_L$ (= 147.51) and $K_L$ (=0.00396 mM) were determined by referring to the experimental measurements of steady-state inactivation. The rate of removing $Ca^{2+}$ inactivation ($\varepsilon_-$) used in hVC model was used.

$$\varepsilon_- = \frac{1}{8084 \cdot exp\left(\frac{V_m}{10}\right) + 158 \cdot exp\left(\frac{V_m}{1000}\right)} + \frac{1}{134736 \cdot exp\left(-\frac{V_m}{5}\right) + 337 \cdot exp\left(-\frac{V_m}{2000}\right)} \qquad Eq.S38$$

The composition of whole cell $I_{CaL}$.

$$I_{CaL} = (I_{CaL\_Ca\_jnc} + I_{CaL\_Na\_jnc} + I_{CaL\_K\_jnc}) + (I_{CaL\_Ca\_iz} + I_{CaL\_Na\_iz} + I_{CaL\_K\_iz}) + (I_{CaL\_Ca\_blk} + I_{CaL\_Na\_blk} + I_{CaL\_K\_blk}) \quad Eq.S39$$

### The sustained inward current (*Ist*)

In the spontaneous SA node cells, a sustained inward current was activated on depolarization to more negative potential range ($V_m$ =~ -60 mV) than the usual threshold of the L-type $Ca^{2+}$ current. The characteristics of the current was roughly similar to $I_{CaL}$, except that $I_{st}$ was resistant to the removal of $Ca^{2+}$ from the external solution and it was suggested that $I_{st}$ is most probably carried by $Na^+$ (Guo et al., 1995, Mitsuiye et al., 1999). Recently, Toyoda et al. (2017) suggested that this current is generated by Cav.1.3, which is the major subtype expressed in the SA node cells. In the present study, we calculated $I_{st}$ in the iPSC_CMs for convenience of comparing the role of $I_{st}$ between the iPSC_CMs and the matured SA node cells. If appropriate, the sum of ($I_{st}$ and the conventional $I_{CaL}$) was calculated.

$$I_{st} = I_{st,Na} + I_{st,K} \quad \text{Eq.S40}$$

$$I_{st,Na} = P_{stNa} \cdot GHK_{Na} \cdot pO \quad P_{stNa} = 0.00236\, pA/pF/mM \quad \text{Eq.S41}$$

$$I_{st,K} = P_{stK} \cdot GHK_K \cdot pO \quad P_{stK} = 0.585 \cdot P_{stNa}\, pA/pF/mM \quad \text{Eq.S42}$$

$$P_O = d \cdot f \cdot u \quad \text{Eq.S43}$$

$$(1-y) \underset{\beta}{\overset{\alpha}{\rightleftarrows}} y \qquad y = \{d, f, u\} \quad \text{Eq.S44}$$

$$\alpha_d = \frac{1}{0.15 \cdot Exp(\frac{V_m}{-11}) + 0.2 \cdot Exp(\frac{V_m}{-700})} \quad \text{Eq.S45}$$

$$\beta_d = \frac{1}{16 \cdot Exp(\frac{V_m}{8}) + 15 \cdot Exp(\frac{V_m}{50})} \quad \text{Eq.S46}$$

$$\alpha_f = \frac{1}{3100 \cdot Exp(\frac{V_m}{13}) + 700 \cdot Exp(\frac{V_m}{70})} \quad \text{Eq.S47}$$

$$\beta_f = \frac{1}{95 \cdot Exp(\frac{V_m}{-10}) + 50 \cdot Exp(\frac{V_m}{-700})} + \frac{2.5 \cdot [Ca^{2+}]_{blk}}{1 + Exp(\frac{V_m}{-5})} \quad \text{Eq. S48}$$

$$\alpha_u = \frac{1}{400000 \cdot Exp(\frac{V_m}{9}) + 60 \cdot Exp(\frac{V_m}{65})} \quad \text{Eq. S49}$$

$$\beta_u = \frac{1}{700 \cdot Exp(\frac{V_m}{-14}) + 60 \cdot Exp(\frac{V_m}{-65})} \quad \text{Eq. S50}$$

### T-type $Ca^{2+}$ current ($I_{CaT}$)

$I_{CaT}$ is assumed in the *blk* space.

$$I_{CaT} = 2 \cdot P_{CaT} \cdot GHK_{Ca} \cdot pO_{CaT},\ P_{CaT} = 9.56 \quad Eq.S51$$

$$pO_{CaT} = d \cdot f \quad Eq.S52$$

$$\alpha_d = \frac{1}{0.019 \cdot exp\left(-\frac{V_m}{5.6}\right) + 0.82 \cdot exp\left(-\frac{V_m}{250}\right)} \quad Eq.S53$$

$$\beta_d = \frac{1}{40 \cdot exp\left(\frac{V_m}{6.3}\right) + 1.5 \cdot exp\left(\frac{V_m}{10000}\right)} \quad Eq.S54$$

$$\alpha_f = \frac{1}{62000 \cdot exp\left(\frac{V_m}{10.1}\right) + 30 \cdot exp\left(\frac{V_m}{3000}\right)} \quad Eq.S55$$

$$\beta_f = \frac{1}{0.0006 \cdot exp\left(-\frac{V_m}{6.7}\right) + 1.2 \cdot exp\left(-\frac{V_m}{25}\right)} \quad Eq.S56$$

### The hyperpolarization-activated current ($I_{ha}$ or $I_f$)

In 1976, Noma and Irisawa for the first time conducted the double-microelectrode voltage clamp in a man-made small tissue preparation (0.2~0.3 mm in diameter) of the rabbit SA node tissue. They found a very slow activation time course of inward current ($I_h$) on hyperpolarization from the holding potential of -40 mV. Yanagihara and Irisawa (1980) clearly separated $I_h$ from the delayed rectifier K current by the difference in the activation range and the $Ba^{2+}$-resistant nature of $I_{ha}$. They measured the fully activated I-V relationship with the reversal potential is at -25 mV, suggesting little sensitivity to any particular ion species. They developed the Hodgkin-Huxley type kinetic model of $I_{ha}$, and suggested that $I_{ha}$ plays a significant role in keeping the pacemaker cell at a low membrane potential, but only a small role in promoting the slow diastolic depolarization because of its time constant of several seconds. Yanagihara et al., (1980) published the mathematical model of the SA node cell action potential. The detailed $I_{ha}$ model described by Maruoka et al., (1994) was used to reflect the delay in both activation and deactivation on hyper- and de-polarizations, respectively.

$$I_{ha} = I_{ha,Na} + I_{ha,K} \quad \text{Eq. S57}$$

$$I_{ha,Na} = P_{ha,Na} \cdot GHK_{Na} \cdot pO \qquad P_{ha,Na} = 0.03642 \quad pA/pF/mM \quad \text{Eq. S58}$$

$$I_{ha,K} = P_{ha,K} \cdot GHK_{K} \cdot pO \qquad P_{ha,K} = 4.244 \cdot P_{ha,Na} \quad pA/pF/mM$$

Eq. S59

$$C1 \underset{\lambda}{\overset{\mu}{\rightleftarrows}} C2 \underset{\beta}{\overset{\alpha}{\rightleftarrows}} O1 \underset{\beta}{\overset{\alpha}{\rightleftarrows}} O2 \underset{\beta}{\overset{\alpha}{\rightleftarrows}} O3 \quad \text{Eq. S60}$$

$$\alpha_{ha} = \frac{1}{3500 \cdot \exp(\frac{V_m}{16.8}) + 0.3 \cdot \exp(\frac{V_m}{400})} \quad \text{Eq. S61}$$

$$\beta_{ha} = \frac{1}{4 \cdot \exp(\frac{V_m}{-14}) + 2 \cdot \exp(\frac{V_m}{-400})} \quad \text{Eq. S62}$$

$$\mu_{ha} = \frac{1}{45000000 \cdot \exp(\frac{V_m}{8.7}) + 500 \cdot \exp(\frac{V_m}{200})} \quad \text{Eq. S63}$$

$$\lambda_{ha} = \frac{1}{10.5 \cdot \exp(\frac{V_m}{-16.4}) + 0.4 \cdot \exp(\frac{V_m}{-400})} \quad \text{Eq. S64}$$

In the steady-state, the full model is reduced to a two-state transition model of a closed state (Ct) and an open state (Ot) to obtain the steady-state open probability (pOtss)

$$Ct \underset{kb}{\overset{kf}{\rightleftarrows}} Ot \qquad \text{Eq. S65}$$

where,

$$kf = \frac{\alpha}{\frac{\lambda}{\mu}+1} \qquad kb = \frac{\beta}{1+\frac{\alpha}{\beta}+(\frac{\alpha}{\beta})^2} \qquad pOt_{SS} = \frac{kf}{kf+kb} \qquad \text{Eq. S66}$$

### Sodium current ($I_{Na}$)

The same $I_{Na}$ model as in our previous study (Asakura et al., 2014) was used, except for the amplitude parameters, $f_L$ and $P_{Na}$. $I_{Na}$ is composed of the two components, $I_{NaT}$ and $I_{NaL}$. The scheme for the state transition is shown below.

$$I_{Na} = I_{NaT} + I_{NaL} \qquad Eq.S67$$

$$f_L = \frac{I_{NaL}}{I_{NaT}+I_{NaL}} = 0.04\ or\ 0.01 \qquad Eq.S68$$

$$I_{NaT} = (1-f_L)\cdot P_{Na}\cdot(GHK_{Na}+0.18\cdot GHK_K)\cdot p(O)_{NaT} \qquad Eq.S69$$

$$I_{NaL} = f_L\cdot P_{Na}\cdot(GHK_{Na}+0.18\cdot GHK_K)\cdot p(O)_{NaL} \qquad Eq.S70$$

$$P_{Na\_Na} = 73.77825, P_{Na\_K} = 0.18\cdot P_{Na\_Na}\ (pA/pF/mM)$$

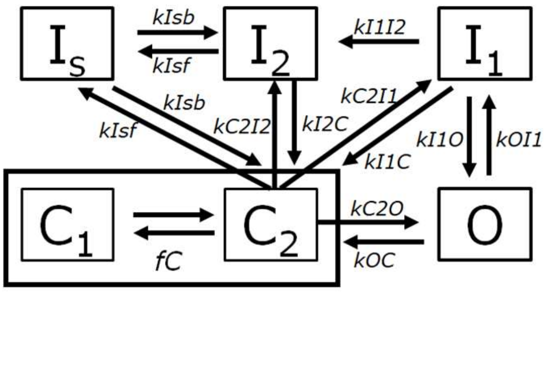

**Transient component ($I_{NaT}$)**

$$\frac{dp(O)_{NaT}}{dt} = k_{I_2O} \cdot p(I_2)_{NaT} + f_{C\_Na} \cdot k_{C_2O} \cdot p(C)_{NaT} - \left(k_{OC} + k_{OI_2}\right) \cdot p(O)_{NaT} \qquad Eq.S71$$

$$\frac{dp(I_2)_{NaT}}{dt} = f_{C\_Na} \cdot k_{C_2I_2} \cdot p(C)_{NaT} + k_{OI_2} \cdot p(O)_{NaT}$$
$$+k_{I_sb} \cdot\ p(I_s)_{NaT} - \left(k_{I_2C} + k_{I_2O} + k_{I_sf}\right) \cdot p(I_2)_{NaT} \qquad Eq.S72$$

$$\frac{dp(I_s)_{NaT}}{dt} = k_{I_sf} \cdot\ p(I_2)_{NaT} + k_{I_sf} \cdot p(C)_{NaT} - 2 \cdot k_{I_sb} \cdot\ p(I_s)_{NaT} \qquad Eq.S73$$

$$p(C)_{NaT} = 1.0 - p(I_s)_{NaT} - p(O)_{NaT} - p(I_2)_{NaT} \qquad Eq.S74$$

$$f_{C\_Na} = \frac{C_2}{C_1 + C_2} = \frac{1}{1 + exp\left(-\frac{v + 48}{7}\right)} \qquad Eq.S75$$

$$v = Vm - VshiftNa \qquad \mathrm{Eq.}\,S76$$

$$k_{C_2O} = \frac{1}{0.0025 \cdot exp\left(-\frac{v}{8.0}\right) + 0.15 \cdot exp\left(-\frac{v}{100.0}\right)} \qquad Eq.S77$$

$$k_{OC} = \frac{1}{30.0 \cdot exp\left(\frac{v}{12.0}\right) + 0.53 \cdot exp\left(\frac{v}{50.0}\right)} \qquad Eq.S78$$

$$k_{OI_2} = \frac{1}{0.0433 \cdot exp\left(-\frac{V_m}{27.0}\right) + 0.34 \cdot exp\left(-\frac{V_m}{2000.0}\right)} \qquad Eq.S79$$

$$k_{I_2O} = 0.0001312 \qquad Eq.S80$$

$$k_{C_2I_2} = \frac{0.5}{1.0 + \frac{k_{I_2O} \cdot k_{OC}}{k_{OI_2} \cdot k_{C_2O}}} \qquad Eq.S81$$

$$k_{I_sb} = \frac{1}{300000.0 \cdot exp\left(\frac{V_m}{10.0}\right) + 50000.0 \cdot exp\left(\frac{V_m}{16.0}\right)} \qquad Eq.S82$$

$$k_{I_sf} = \frac{1}{0.016 \cdot exp\left(-\frac{V_m}{9.9}\right) + 8.0 \cdot exp\left(-\frac{V_m}{45.0}\right)} \qquad Eq.S83$$

### Late component ($I_{NaL}$)

The $k_{I_1 I_2}$, $k_{OI_1}$, $k_{I_1 O}$, $k_{I_1 C}$ and $k_{C_2 I_1}$ are specific for $I_{NaL}$, and other rate constants are the same as in $I_{NaT}$.

$$\frac{dp(O)_{NaL}}{dt} = k_{I_1 O} \cdot p(I_1)_{NaL} + f_{C\_Na} \cdot k_{C_2 O} \cdot p(C)_{NaL} - \left(k_{OC} + k_{OI_1}\right) \cdot p(O)_{NaL} \qquad Eq.S84$$

$$\frac{dp(I_1)_{NaL}}{dt} = f_{C\_Na} \cdot k_{C_2 I_1} \cdot p(C)_{NaL} + k_{OI_1} \cdot p(O)_{NaL} - \left(k_{I_1 C} + k_{I_1 O} + k_{I_1 I_2}\right) \cdot p(I_1)_{NaL} \quad Eq.S85$$

$$\frac{dp(I_2)_{NaL}}{dt} = f_{C\_Na} \cdot k_{C_2 I_2} \cdot p(C)_{NaL} + k_{I_1 I_2} \cdot p(I_1)_{NaL} + k_{I_s b} \cdot\ p(I_s)_{NaL} - \left(k_{I_2 C} + k_{I_s f}\right) \cdot p(I_2)_{NaL} \qquad Eq.S86$$

$$\frac{dp(I_s)_{NaL}}{dt} = k_{I_s f} \cdot\ p(I_2)_{NaL} + k_{I_s f} \cdot p(C)_{NaL} - 2 \cdot k_{I_s b} \cdot\ p(I_s)_{NaL} \qquad Eq.S87$$

$$p(C)_{NaL} = 1.0 - p(I_s)_{NaL} - p(O)_{NaL} - p(I_1)_{NaL} - p(I_2)_{NaL} \qquad Eq.S88$$

$$k_{I_1 I_2} = 0.00534 \qquad Eq.S89$$

$$k_{OI_1} = k_{OI_2} \qquad Eq.S90$$

$$k_{I_1 O} = 0.01 \qquad Eq.S91$$

$$k_{I_1 C} = k_{I_2 C} \qquad Eq.S92$$

$$k_{C_2 I_1} = k_{C_2 I_2} \qquad Eq.S93$$

### Inward rectifier potassium current ($I_{K1}$)

The $I_{K1}$ model developed by Yan and Ishihara (Yan and Ishihara, 2005) and Ishihara and Yan, (2007) was used in hVC model after modifying several parameters (Himeno et al., 2015).

$$I_{K1} = G_{K1} \cdot (V_m - E_K) \cdot p(O)_{K1} \qquad Eq.S94$$

$$G_{K1} = \frac{0.451773 \cdot \left(\frac{[K^+]_O}{5.4}\right)^{0.4}}{1 + exp\left(-\frac{[K^+]_O - 2.2}{0.6}\right)}\ nS/pF \qquad Eq.S95$$

$$p(O)_{K1} = pO_{mo} \quad + pO_{mode2} \qquad Eq.S96$$

### Mode 1: the channel block by $Mg^{2+}$ and spermine (SPM)

The SPM-block is a time-dependent process, while the $Mg^{2+}$-block is instantaneous.

$$Pbspm \underset{\beta \cdot pOMg}{\overset{\alpha}{\rightleftarrows}} 1 - Pbspm \qquad \text{Eq. S97}$$

$$\alpha_{Mg} = 12.0 \cdot exp\left(-\frac{V_m - E_K}{40}\right) \qquad Eq.S98$$

$$\beta_{Mg} = 28.0 \cdot exp\left(-\frac{V_m - E_K}{40}\right) \cdot [Mg^{2+}]_{cyt} \qquad Eq.S99$$

$$f_O = \frac{\alpha_{Mg}}{\alpha_{Mg} + \beta_{Mg}} \qquad Eq.S100$$

$$f_B = \frac{\beta_{mg}}{\alpha_{Mg} + \beta_{Mg}} \qquad Eq.S101$$

$$pO_{Mg} = f_O \cdot f_O \cdot f_O \qquad Eq.S102$$

$$pO_{Mg} \ = 3.0 \cdot f_O \cdot f_O \cdot f_B \qquad Eq.S103$$

$$pO_{Mg2} = 3.0 \cdot f_O \cdot f_B \cdot f_B \qquad Eq.S104$$

$$pB_{Mg} \ = f_B \cdot f_B \cdot f_B \qquad Eq.S105$$

$$\alpha_{SPM} = \frac{0.17 \cdot exp\left(-0.07 \cdot \left((V_m - E_K) + 8 \cdot [Mg^{2+}]_{cyt}\right)\right)}{1.0 + 0.01 \cdot exp\left(0.12 \cdot \left((V_m - E_K) + 8 \cdot [Mg^{2+}]_{cyt}\right)\right)} \qquad Eq.S106$$

$$\beta_{SPM} = \frac{0.28 \cdot [SPM] \cdot exp\left(0.15 \cdot \left((V_m - E_K) + 8 \cdot [Mg^{2+}]_{cyt}\right)\right)}{1.0 + 0.01 \cdot exp\left(0.13 \cdot \left((V_m - E_K) + 8 \cdot [Mg^{2+}]_{cyt}\right)\right)} \qquad Eq.S107$$

$$\frac{dPb_{SPM}}{dt} = \beta_{SPM} \cdot pO_{Mg} \cdot (1 - Pb_{SPM}) - \alpha_{SPM} \cdot Pb_{SPM} \qquad Eq.S108$$

$$pO_{mode1} = 0.9 \cdot (1 - Pb_{SPM}) \cdot \left(pO_{Mg} + \frac{2}{3} pO_{Mg} \ + \frac{1}{3} pO_{Mg} \ \right) \qquad Eq.S109$$

### Mode 2: the channel block only by SPM

The channel is free from the Mg-block, and the SPM is instantaneous.

$$Pbspm \underset{Kd}{\longleftrightarrow} [SPM] \cdot (1 - Pbspm)$$

$$pO_{mo} = \frac{0.1}{1+\frac{[SPM]}{Kd}} \quad Kd = 40 \cdot exp\left(-\frac{V_m - E_K}{9.1}\right)\ mM \qquad Eq.S110$$

### Delayed rectifier K$^+$ current, fast component ($I_{Kr}$)

We installed the $I_{Kr}$ model developed by Ono and Ito (1995) (Ono and Ito, 1995), which well fitted the result of experimental $I_{Kr}$ data of hiPSC-CMs (Ma et al., 2011)..

The current amplitude is described with an Ohmic equation.

$$I_{Kr} = G_{Kr} \cdot (V_m - E_K) \cdot p(O)_{Kr} \qquad Eq.S111$$

$$G_{Kr} = 0.049644 \cdot \left(\frac{[K^+]_O}{5.4}\right)^{0.2}\ nS/pF \qquad Eq.S112$$

The open probability of the channel is described with three gating parameters, $y_1$, $y_2$, and $y_3$, each of which is calculated by a two-state transition scheme.

$$p(O)_{Kr} = (0.6 \cdot y_1 + 0.4 \cdot y_2) \cdot y_3 \qquad Eq.S113$$

$$\frac{dy_N}{dt} = \alpha_{y_N} \cdot (1.0 - y_N) - \beta_{y_N} \cdot y_N, \qquad N = 1, 2, 3 \qquad Eq.S114$$

$$\alpha_{y_1} = \frac{1}{20 \cdot exp\left(-\frac{V_m + 6}{6}\right) + 5 \cdot exp\left(-\frac{V_m + 6}{150}\right)} \qquad Eq.S115$$

$$\beta_{y_1} = \frac{1}{160 \cdot exp\left(\frac{(V_m + 6)}{28}\right) + 200 \cdot exp\left(\frac{(V_m + 6)}{1000}\right)} + \frac{1}{2500 \cdot exp\left(\frac{(Vm + 6)}{20}\right)} \qquad Eq.S116$$

$$\alpha_{y_2} = \frac{1}{200 \cdot exp\left(-\frac{(V_m + 6)}{6.5}\right) + 20 \cdot exp\left(-\frac{(V_m + 6)}{150}\right)} \qquad Eq.S117$$

$$\beta_{y_2} = \frac{1}{1600 \cdot exp\left(\frac{(V_m + 6)}{28}\right) + 2000 \cdot exp\left(\frac{(V_m + 6)}{1000}\right)} + \frac{1}{10000 \cdot exp\left(\frac{(Vm + 6)}{20}\right)} \qquad Eq.S118$$

$$\alpha_{y_3} = \frac{1}{10 \cdot exp\left(\frac{(V_m + 6)}{17}\right) + 2.5 \cdot exp\left(\frac{(V_m + 6)}{300}\right)} \qquad Eq.S119$$

$$\beta_{y_3} = \frac{1}{0.35 \cdot exp\left(-\frac{(V_m + 6)}{17}\right) + exp\left(-\frac{(V_m + 6)}{75}\right)} \qquad Eq.S120$$

**Delayed rectifier K+ current, slow component ($I_{Ks}$)**

$$I_{Ks\_K} = P_{Ks} \cdot GHK_K \cdot p(O)_{Ks}\,,\; P_{Ks} = 0.4\;(pA/pF/mM) \qquad Eq.S121$$

$$I_{Ks\_Na} = 0.04 \cdot P_{Ks} \cdot GHK_{Na} \cdot p(O)_{Ks} \qquad Eq.S122$$

$$p(O)_{Ks} = (O_v)^2 \cdot (0.99 \cdot O_C + 0.01) \qquad Eq.S123$$

**The $V_m$-dependent gate**

$$\alpha_{v_{Ks}} = \frac{1}{150 \cdot exp\left(-\frac{(V_m + 10)}{25}\right) + 900 \cdot exp\left(-\frac{(V_m + 10)}{200}\right)} \qquad Eq.S124$$

$$\beta_{v_{Ks}} = \frac{1}{1000 \cdot exp\left(\frac{(V_m + 10)}{13}\right) + 220 \cdot exp\left(\frac{(V_m + 10)}{50}\right)} \qquad Eq.S125$$

$$\frac{dO_v}{dt} = \alpha_{v_{Ks}} \cdot (1.0 - O_v) - \beta_{v_{Ks}} \cdot O_v \qquad Eq.S126$$

**The $Ca^{2+}$-dependent gate**

$$C1 \underset{\beta 3}{\overset{\alpha 3}{\rightleftarrows}} C2 \underset{\beta 2}{\overset{\alpha 2}{\rightleftarrows}} Oc$$

$$\frac{dO_c}{dt} = -\alpha_2 \cdot C_2 - \beta_2 \cdot O_c \qquad \text{Eq. S127}$$

$$\frac{dC_2}{dt} = \alpha_3 \cdot C_1 - \beta_3 \cdot C_2 - \alpha_2 \cdot C_2 + \beta_2 \cdot O_c \qquad \text{Eq. S128}$$

$$C_1 = 1.0 - C_2 - O_c \qquad \text{Eq. S129}$$

$$\alpha_3 = 0.0003, \qquad \beta_3 = 0.03 \qquad Eq.S130$$

$$\alpha_2 = 2.25 \cdot [Ca^{2+}], \qquad \beta_2 = 0.000296 \qquad Eq.S131$$

**Transient outward $K^+$ current ($I_{Kto}$)**

$$I_{Kto\_K} = P_{Kto} \cdot GHK_K \cdot p(O)_{Kto}\ , P_{Kto} = 0.01729\ (pA/pF/mM) \qquad Eq.S132$$

$$I_{Kto\_Na} = 0.09 \cdot P_{Kto} \cdot GHK_{Na} \cdot p(O)_{Kto} \qquad Eq.S40$$

$$p(O)_{Kto} = y_{1_{Kto}} \cdot y_{2_{Kto}} \qquad Eq.S41$$

$$\frac{dy_{1_{Kto}}}{dt} = \alpha_{y_1} \cdot \left(1.0 - y_{1_{Kto}}\right) - \beta_{y_1} \cdot y_{1_{Kto}}, \quad \frac{dy_{2_{Kto}}}{dt} = \alpha_{y_2} \cdot \left(1.0 - y_{2_{Kto}}\right) - \beta_{y_2} \cdot y_{2_{Kto}} \qquad Eq.S42$$

$$\alpha_{y_1} = \frac{1}{13 \cdot exp\left(-\frac{V_m}{22}\right)}, \qquad \beta_{y_1} = \frac{1}{2.1 \cdot exp\left(\frac{V_m}{90}\right)} \qquad Eq.S43$$

$$\alpha_{y_2} = \frac{0.5}{950 \cdot exp\left(\frac{V_m}{500}\right)}, \qquad \beta_{y_2} = \frac{0.5}{40 \cdot exp\left(-\frac{V_m}{9}\right) + 13 \cdot exp\left(-\frac{V_m}{1000}\right)} \qquad Eq.S44$$

**Ultra-rapid $K^+$ current ($I_{Kur}$)**

$I_{Kur}$ model in a mouse ventricular cell model (Bondarenko et al., 2004) is used.

$$I_{Kur} = G_{Kur} \cdot (a_{ur})^3 \cdot i_{ur} \cdot (V_m - E_K) \qquad Eq.S45$$

$$G_{Kur} = 0.000113 \cdot \left(1 + \frac{1}{1 + exp\left(-\frac{V_m - 30}{59}\right)}\right) nS/pF \qquad Eq.S46$$

$$\frac{da_{ur}}{dt} = \alpha_{ur} \cdot (1.0 - a_{ur}) - \beta_{ua} \cdot a_{ur}, \qquad \frac{di_{ur}}{dt} = \alpha_{ui} \cdot (1.0 - i_{ur}) - \beta_{ui} \cdot i_{ur} \qquad Eq.S47$$

$$\alpha_{ua} = \frac{1}{0.65 \cdot exp\left(-\frac{(V_m + 19)}{8.5}\right) + exp\left(-\frac{(V_m - 21)}{59}\right)}, \qquad \beta_{ua} = \frac{1}{0.65 \cdot \left(2.5 + exp\left(\frac{(V_m + 91)}{17}\right)\right)} \qquad Eq.S48$$

$$\alpha_{ui} = \frac{1}{21 + exp\left(-\frac{(V_m - 185)}{28}\right)}, \qquad \beta_{ui} = exp\left(\frac{V_m - 158}{16}\right) \qquad Eq.S49$$

### Time-independent currents

All these currents are from Takeuchi *et al.* (Takeuchi et al., 2006) as described in Asakura *et al.* (Asakura et al., 2014).

**Background Ca$^{2+}$ current ($I_{bCa}$)**

$$I_{bCa\_a} = P_{bCa\_a} \cdot 2 \cdot GHK_{Ca}, \qquad a = (blk, iz) \qquad Eq.S50$$

$$P_{bCa} = 0.00125 \ (pA/pF/mM) \qquad Eq.S51$$

**Background non-selective cation current ($I_{bNSC}$)**

$$I_{bNSC\_X} = P_{bNSC_X} \cdot GHK_X, \qquad X = (K, Na) \qquad Eq.S52$$

$$P_{bNSC\_Na} = 0.000182875, P_{bNSC\_K} = 0.4 \cdot P_{bNSC\_Na}(pA/pF/mM) \qquad Eq.S53$$

$$I_{bNSC} = I_{bNSC\_K} + I_{bNSC\_Na} \qquad Eq.S54$$

**Calcium-activated background cation current ($I_{l(Ca)}$)**

$$p(O)_a = \frac{1.0}{1.0 + \left(\frac{0.0012}{[Ca^{2+}]_a}\right)^3} \qquad Eq.S148$$

$$I_{l(Ca)\_X\_a} = P_{l(Ca)\_X\_a} \cdot f_{l(Ca)\_X\_a} \cdot GHK_X \cdot p(O)_a, \qquad X = (Na, K), \qquad a = (blk, iz) \quad Eq.S149$$

$$P_{l(Ca)\_Na} = 0.01375 \ (pA/pF/mM) \qquad Eq.S150$$

$$P_{l(Ca)\_K} = P_{l(Ca)\_Na} \ (pA/pF/mM) \qquad Eq.S151$$

***Fraction of $I_{l(Ca)}$***

$$f_{l(Ca)\_iz} = 0.1, f_{l(Ca)\_blk} = 0.9 \qquad Eq.S152$$

$$I_{l(Ca)} = I_{l(Ca)\_Na\_iz} + I_{l(Ca)\_K\_iz} + I_{l(Ca)\_Na\_blk} + I_{l(Ca)\_K\_blk} \qquad Eq.S153$$

**ATP-sensitive potassium current ($I_{KATP}$)**

$$p(O)_{KATP} = \frac{0.8}{1.0 + \left(\frac{[ATP]_{cyt}}{0.1}\right)^2} \qquad Eq.S154$$

$$\chi_{KATP} = 0.0236 \cdot ([K^+]_o)^{0.24} \qquad Eq.S155$$

$$I_{KATP} = G_{KATP} \cdot (V_m - E_K) \cdot p(O)_{KATP} \cdot \chi_{KATP} \qquad Eq.S156$$

$$G_{KATP} = 18.75 \qquad Eq.S157$$

### $Na^+/K^+$ pump current ($I_{NaK}$)

The $Na^+/K^+$ pump model developed by Oka *et al.* (Oka et al., 2010) on the framework of Smith and Crampin (Smith and Crampin, 2004) was used after adjusting the amplitude as indicated in the manuscript in Eq. 13.

### $Na^+/Ca^{2+}$ exchange current ($I_{NCX}$)

The NCX model developed by Takeuchi *et al.* (Takeuchi et al., 2006) was used after adjusting the amplitude as indicated in the manuscript in Eq. 14.

### CaRU

The model of CaRU described in the hVC model was used. The model structure of CaRU is shown in Fig. S1. Derivation of the instantaneous $[Ca^{2+}]_{nd}$, which is sensed by both LCC and RyRs for inactivation and activation, respectively, was originally given in Hinch 2004 and modified by Himeno et al. 2015. In short, the volume of *nd* was assumed to be virtually zero so that the $[Ca^{2+}]_{nd}$ can be given by the instantaneous equation Eq. S158,

$$[Ca]_{nd} = \frac{[Ca]_{jnc} + \frac{g_R}{g_d}\cdot[Ca]_{SRrl} + \frac{g_L}{g_d}\cdot\frac{\delta V\cdot e^{-\delta V}}{1-e^{-\delta V}}\cdot[Ca]_o}{(1+\frac{g_R}{g_d}+\frac{g_L}{g_d}\cdot\frac{\delta V\cdot e^{-\delta V}}{1-e^{-\delta V}})} \qquad \text{Eq. S158}$$

where $Ca^{2+}$ fluxes through LCC and RyR, and diffusion from *nd* to *jnc* ($J_L$, $J_R$ and $J_D$) were given as Eqs. S134-136.

$$J_L = g_L\cdot\frac{\delta V\cdot e^{-\delta V}}{1-e^{-\delta V}}\cdot([Ca]_o - [Ca]_{nd}) \qquad \text{Eq. S159}$$

$$J_R = g_R\cdot([Ca]_{SRrl} - [Ca]_{nd}) \qquad \text{Eq. S160}$$

$$J_D = g_D\cdot([Ca]_{nd} - [Ca]_{jnc})$$

Eq. S161

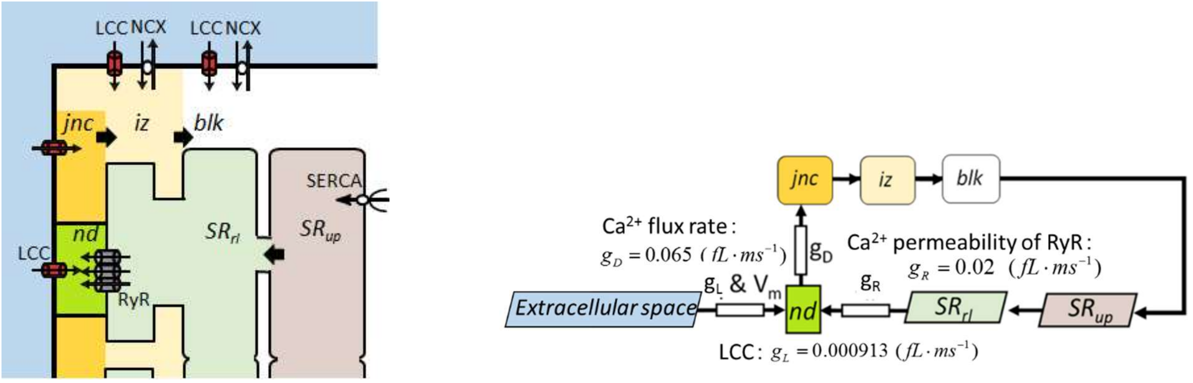

*Fig. S1 Model structure of CaRU in relation to three $Ca^{2+}$ diffusion compartments (left) and direction of $Ca^{2+}$ diffusion (right) within the cell model. The L-type $Ca^{2+}$ channel (LCC) and NCX are located on the sarcolemma, SERCA and RyRs are on the SR membrane. A single CaRU consists of a hypothetical LCC and a couplon (a cluster of RyRs) in the junctional cleft (filled with lime green), and individual CaRUs are spatially separated from its neighbors by jnc. The inset on the right shows schematic presentation of the diffusion pathway of $Ca^{2+}$ from the $Ca^{2+}$ sources (SR or extracellular space) to the sink (nd) and then to the cytoplasm (jnc, iz and blk). $J_L$, $J_R$ and $g_D$ represent permeability of single LCC and RyR, and $Ca^{2+}$ flux rate from nd to jnc, respectively.*

### Sarcoplasmic reticulum $Ca^{2+}$ pump (SERCA) current ($J_{SERCA}$)

The three-state model developed by Tran *et al.* (Tran et al., 2009) was used after several minor modifications as described in Asakura *et al.* (Asakura et al., 2014). The limiting amplitude of $J_{SERCA}$, $Amp_{SERCA}$, was modified.

### Rate of change in the membrane potential and ion concentrations

### Membrane potential

$$\frac{dV_m}{dt} = -(I_{tot\_cell} + I_{app}) \qquad Eq.S162$$

$$I_{tot\_cell} = I_{tot\_Ca} + I_{tot\_Na} + I_{tot\_K} \qquad Eq.S163$$

$$I_{tot\_Ca} = I_{tot\_Ca\_jnc} + I_{tot\_Ca\_iz} + I_{tot\_Ca\_blk} \qquad Eq.S164$$

$$I_{tot\_Ca\_jnc} = I_{CaL\_Ca\_LR} + I_{CaL\_Ca\_L0} \qquad Eq.S165$$

$$I_{tot\_Ca\_iz} = I_{CaL\_Ca\_iz} + I_{NCX\_Ca\_iz} + I_{Cab\_iz} \qquad Eq.S166$$

$$I_{tot\_Ca\_blk} = I_{CaL\_Ca\_blk} + I_{CaT} + I_{Cab\_blk} + I_{NCX\_Ca\_blk} \qquad Eq.S167$$

$$I_{tot\_Na} = \left(I_{CaL\_Na\_jnc} + I_{CaL\_Na\_iz} + I_{CaL\_Na\_blk}\right) + \left(I_{NCX\_Na\_iz} + I_{NCX\_Na\_blk}\right) + \left(I_{Ks\_Na\_iz} + I_{Ks\_Na\_blk}\right) + I_{NaT\_Na} + I_{NaL\_Na} + I_{NaK\_Na} + I_{Kto\_Na} + I_{bNSC\_Na} + \left(I_{LCCa\_Na\_iz} + I_{LCCa\_Na\_blk}\right) + I_{st\_Na} + I_{ha\_Na} \qquad Eq.S168$$

$$I_{tot\_K} = \left(I_{CaL\_K\_jnc} + I_{CaL\_K\_iz} + I_{CaL\_K\_blk}\right) + I_{NaT\_K} + I_{NaL\_K} + I_{K1} + I_{Kur} + I_{Kpl} + I_{Kr} + \left(I_{Ks\_K\_iz} + I_{Ks\_K\_blk}\right) + I_{Kto\_K} + I_{KATP\_K\_cyt} + I_{bNSC\_K} + \left(I_{LCCa\_K\_iz} + I_{LCCa\_K\_blk}\right) + I_{NaK\_K} + I_{KAC} + I_{st\_K} + I_{ha\_K} \qquad Eq.S169$$

### Ion concentrations

$$\frac{d[Ca^{2+}_{total}]_{jnc}}{dt} = -\frac{I_{tot\_Ca\_jnc} \cdot C_m}{V_{jnc} \cdot 2 \cdot F} + \frac{J_{Ca\_rel}}{V_{jnc}} - \frac{J_{Ca\_jnciz}}{V_{jnc}} \qquad Eq.S170$$

$$\frac{d[Ca^{2+}_{total}]_{iz}}{dt} = -\frac{I_{tot\_Ca\_iz} \cdot C_m}{V_{iz} \cdot 2 \cdot F} + \frac{J_{Ca\_jnciz}}{V_{iz}} - \frac{J_{Ca\_izblk}}{V_{iz}} \qquad Eq.S171$$

$$\frac{d[Ca^{2+}_{total}]_{blk}}{dt} = -\frac{I_{tot\_Ca\_blk} \cdot C_m}{V_{blk} \cdot 2 \cdot F} + \frac{J_{Ca\_izblk}}{V_{blk}} - \frac{J_{Ca\_SERCA}}{V_{blk}} \qquad Eq.S172$$

$$\frac{d[Ca^{2+}]_{SRup}}{dt} = \frac{J_{Ca\_SERCA}}{V_{SRup}} - \frac{J_{trans\_SR}}{V_{SRup}} \qquad Eq.S173$$

$$\frac{d[Ca^{2+}_{total}]_{SRrl}}{dt} = \frac{J_{trans\_SR}}{V_{SRrl}} - \frac{J_{rel\_SR}}{V_{SRrl}} \qquad Eq.S174$$

$$\frac{d[Na^{+}]_i}{dt} = -\frac{I_{tot\_Na} \cdot C_m}{V_{cyt} \cdot F} \qquad Eq.S175$$

$$\frac{d[K^{+}]_i}{dt} = -\frac{(I_{tot\_K} + I_{app}) \cdot C_m}{V_{cyt} \cdot F} \qquad Eq.S176$$

### Contraction

The original model of Negroni and Lascano (Negroni and Lascano, 2008) was used. The magnitude of $F_b$ is given in a unit of mN·mm$^{-2}$. The binding of $Ca^{2+}$ to a troponin system (TS) having 3 $Ca^{2+}$ binding sites (given in μM) was included in the equation of determining the concentration of free $Ca^{2+}$ in the bulk compartment.

$$[Ca^{2+}]_{blk} = [Ca^{2+}_{total}]_{blk} - \left([CaMCa] + [TnChCa] + [SRCa] + \frac{3 \cdot ([TSCa_3] + [TSCa_3^{\sim}] + [TSCa_3^{*}])}{1000}\right)$$

$$Eq.S177$$

### Fitting the hiPSC_CM model to ion channel data of hiPSC-CMs

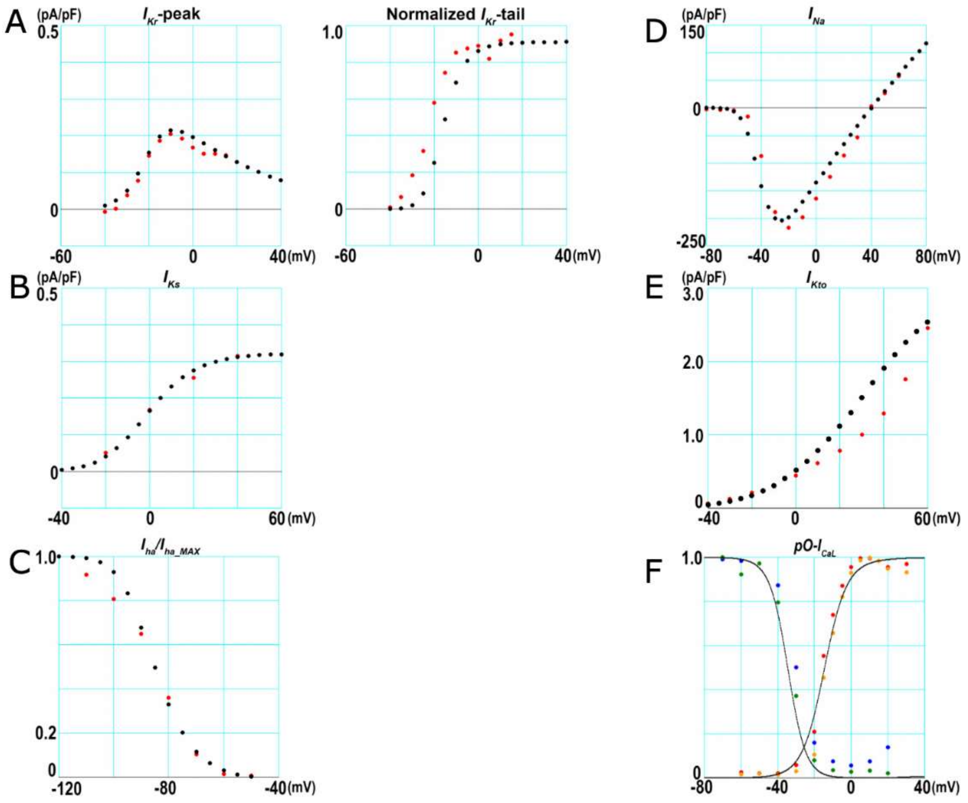

*Fig. S2 Comparison of previously reported experimental results with mathematical models fitted to them. hiPSC-CMs mathematical model was constructed on the basis of hVC model and modified for experimental data from hiPSC-CMs. A, B, C, D, and E: Red points show experimental data from Ma et al., 2011, and black points exhibit fitting results of mathematical modeling. F: Red and blue points show experimental data from Ma et al., 2011. Orange and green points in F are our original experimental results. Black traces exhibit fitting results of mathematical modeling.*

### Pattern Search Algorithm

Pseudo code of the pattern search algorithm used in the research is shown in Fig. S3.

```
PatternSearch():
  Stp = 1/100;
  RedFct = 1/4;
  EPS = 1.0E+37;
  Eval = 0;
  for (i = 0; i < NumPara; i++) {
    NP[i] = BP[i] = RandomBP();
  }

  MIN = MSE = givenEquation(NP);
  Eval++;

  do {
    Fails = EXPLORER();
    while (Fails != NumPara) {
      for (i = 0; i < NumPara; i++) {
        Advance[i] = NP[i] - BP[i];
      }
      do {
        for (i = 0; i < NumPara; i++) {
              BP[i] = NP[i];
          NP[i] = BP[i] + Advance[i];
          if (Advance[i] * Stp[i] <= 0) {
            Stp[i] = -Stp[i];
          }
        }
        If Eval > EvalMax {
          return;
        }
        MinSto = MIN;
        MIN = MSE = givenEquation(NP);
        Fails = EXPLORE();
        if (MIN <= MinSto) {
          for (i = 1; i < NumPara; i++) {
            Advance[i] = NP[i] - BP[i];
          }
          maxAdvance = max(Advance[]);
        }
      } while ((MIN <= MinSto) && (maxAdvance > EPS));
      MIN = MinSto;
      for (i = 0; i < NumPara; i++) {
        NP[i] = BP[i];
      }
      Fails = EXPLORE();
    }
    Stp = Stp * RedFct;
  } while (Stp / RedFct > CrtStp);
}
```

```
EXPLORE() {
  Fails = 0;
  for (i = 0; i < NumPara; i++) {
    HOME = NP[i];
    NP[i] = HOME + Stp;
    MSEp = givenEquation(NP);
    NP[i] = HOME - Stp;
    MSEn = givenEquation(NP);
    minMSE = min(MSEp, MSEn);
    if (minMSE < MSE) {
      if (MSEp < MSEn) {
        NP[i] = HOME + Stp;
      } else {
        NP[i] = HME - Stp;
      }
      MSE = minMSE;
    } else {
      NP[i] = HOME;
      Fails++;
    }
  }
  return Fails;
}
```

*Fig. S3 Pseudo code of the pattern search algorithm explained in Sec. 3.3. Function EXPLORER() searches for the set of* $sf_x$ *that gives smaller MSE by evaluating* $sf_x \pm stp$ *for each* $sf_x$*s. Function givenEquation() calculates MSE with* $sf_x$ *given by the variable NP[].*

### Results on less negative cell (Cell 38)

#### Mapping the magnitude of MSE over the 9 global parameter space

Global random test of nine parameters for a cell with MDP less than -75 mV (Cell 86) was shown in the main manuscript. Here results of same random test for a cell with MDP higher than -75 mV (Cell 38) is shown in Fig S4. Similar with Cell 86, single peak was observed for all the selected currents, and no other local solution was found in this global range.

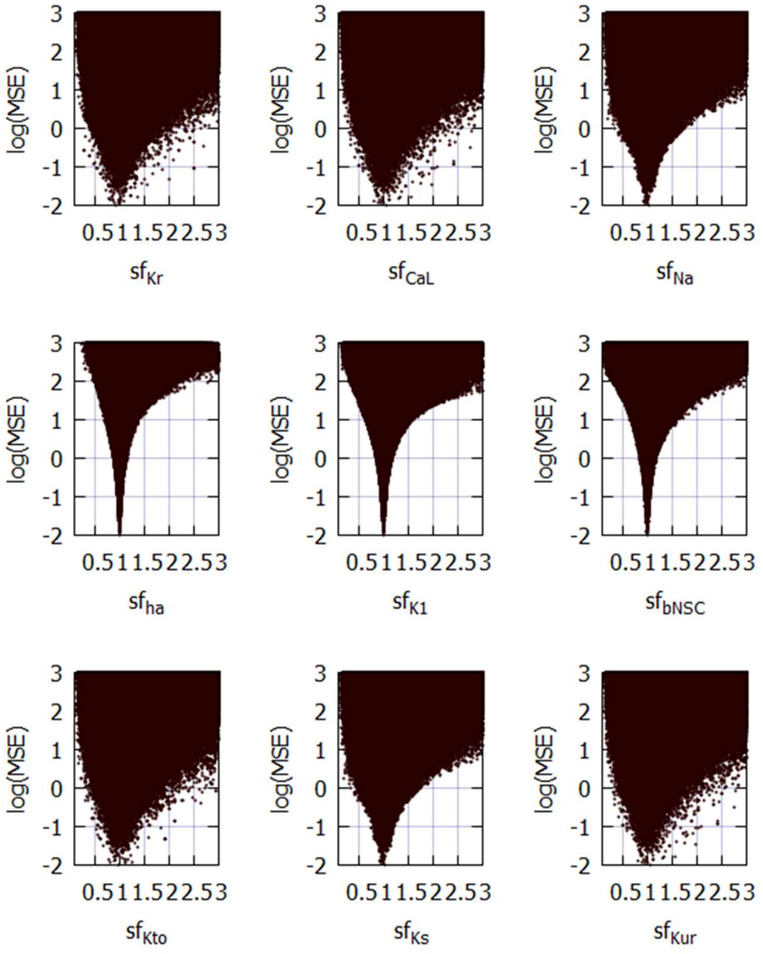

Fig. S4. Nine-parameter random test of cell-38 for the MSE in the $sf_x$-space. The relative value of each current component is plotted on the x-axis, and the log(MSE) is plotted on the y-axis. 37,070,869 points were plotted. The random range for setting the initial value of $sf_x$ were put to the target value of 0.1-10.

**The four-parameter orp test**

As shown in the nine parameter orp test, significant currents are $I_{Kr}$, $I_{CaL}$, $I_{Kur}$ and $I_{bNSC}$ in Cell 38, which can be selected from the physiological consideration. Results of this four parameter orp test is shown in Fig S5. We can find clear convergence of selected four currents.

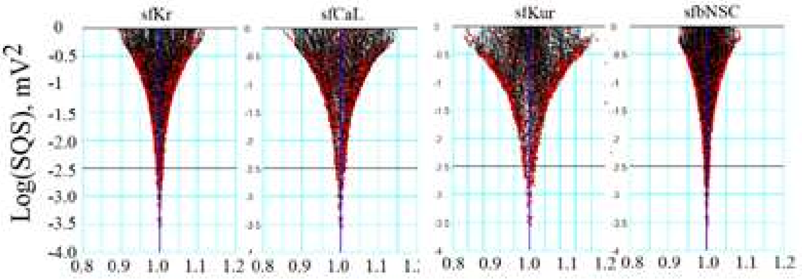

*Fig. S5. Convergence of $sf_x$ in the orp test for Cell38. The ordinate is the Log(MSE) and the abscissa the normalized amplitude of $sf_x$; x stands for $I_{Kr}$, $I_{CaL}$, $I_{Kur}$ and $I_{bNSC}$. Points indicate $sf_x$ obtained in 838 runs of PS optimization.*

### Nomenclature of the ionic currents

*Table S5 Molecular determinants for each current and their expression in iPS-CMs hiPS, human iPS cell line (253G1); hiPS-CM, human iPS cell-derived cardiomyocytes; FH, fetal human heart tissues; AH, adult human heart tissues. RNA expression profiles in hiPS, hiPS-CM, FH and AH were obtained from the Gene Expression Omnibus (GEO) public database (accession: GSE154580* GEO Accession viewer (nih.gov)*). * This is presumed by recent studies (Toyoda et al., 2017; Toyoda et al., 2018).*

| Current | | Protein | Gene | RNA expression (#GSE154580) | | | |
|---|---|---|---|---|---|---|---|
| | | | | hiPS | hiPS-CM | FH | AH |
| $I_{Kr}$ | rapid component of delayed rectifier $K^+$ current | $K_v$11.1 | *KCNH2* | 29.50 | 244.41 | 60.75 | 116.85 |
| $I_{K1}$ | inward rectifier $K^+$ current | $K_{ir}$2.1 | *KCNJ2* | 1.18 | 6.16 | 69.03 | 48.00 |
| $I_{CaL}$ | L-type $Ca^{2+}$ current | $Ca_v$1.2<br>$Ca_v$1.3 | *CACNA1C*<br>*CACNA1D* | 0.00<br>0.34 | 6.58<br>3.37 | 3.02<br>1.39 | 2.84<br>0.23 |
| $I_{bNSC}$ | background nonselective cation current | unidentified | unidentified | | | | |
| $I_{ha}$ | hyperpolarization activated currenIt | HCN4<br>HCN1<br>HCN2 | *HCN4*<br>*HCN1*<br>*HCN2* | 54.48<br>7.19<br>2.27 | 491.55<br>17.02<br>11.16 | 19.97<br>0.85<br>0.16 | 28.13<br>1.32<br>10.61 |
| $I_{Ks}$ | slow component of delayed rectifier $K^+$ current | $K_v$7.1<br>(+KCNE1) | *KCNQ1*<br>(+*KCNE1*) | 4.16<br>(0.00) | 86.48<br>(0.19) | 60.75<br>(0.93) | 53.6<br>(4.55) |
| $I_{Kto}$ | transient outward $K^+$ current | $K_v$4.3<br>$K_v$1.4 | *KCND3*<br>*KCNA4* | 0.00<br>0.08 | 0.00<br>8.45 | 0.23<br>4.80 | 0.28<br>10.97 |
| $I_{Kur}$ | ultra-rapid component of rectifier $K^+$ current | $K_v$1.5 | *KCNA5* | 0.33 | 0.56 | 3.25 | 6.60 |
| $I_{Na}$ | sum of voltage-gated $Na^+$ current in transient and late modes ($I_{NaT}$ + $I_{NaL}$) | $Na_v$1.5 | *SCN5A* | 15.00 | 388.97 | 58.42 | 274.9 |
| $I_{CaT}$ | T-type $Ca^{2+}$ current | $Ca_v$3.2<br>$Ca_v$3.1 | *CACNA1H*<br>*CACNA1G* | 26.98<br>0.00 | 1.77<br>0.64 | 113.76<br>0.00 | 4.30<br>0.00 |
| $I_{st}$ | sustained inward current | $Ca_v$1.3* | *CACNA1D* | 0.34 | 3.37 | 1.39 | 0.23 |
| $I_{NaK}$ | $Na^+$/$K^+$ pump current | Na,K-ATPase<br>α subunit<br>β subunit | *ATP1A1*<br>*ATP1B1* | 603.19<br>80.03 | 1522.44<br>601.05 | 631.26<br>480.27 | 562.90<br>539.60 |
| $I_{NCX}$ | $Na^+$/$Ca^{2+}$ exchanger current | NCX1 | *SLC8A1* | 0.14 | 1509.25 | 266.76 | 560.11 |
