## Supplementary material for "Gradient-based parameter optimization to determine membrane ionic current composition of human induced pluripotent stem cell-derived cardiomyocytes": optimizing program: ComboBoxControl.vb

```
Public Class ComboBoxControl

    Private filePath As String = ""
    Private fileName As String = ""

    'コンストラクタ
    Public Sub New(ByVal id As String, ByVal name As String)
        filePath = id
        fileName = name
    End Sub

    '実際の値
    Public ReadOnly Property FullPath() As String
        Get
            Return filePath
        End Get
    End Property

    '表示名称
    '（このプロパティはこのサンプルでは使わないのでなくても良い）
    Public ReadOnly Property Name() As String
        Get
            Return fileName
        End Get
    End Property

    'オーバーライドしたメソッド
    'これがコンボボックスに表示される
    Public Overrides Function ToString() As String
        Return fileName
    End Function
End Class
```
