## Supplementary material for "Gradient-based parameter optimization to determine membrane ionic current composition of human induced pluripotent stem cell-derived cardiomyocytes": optimizing program: CSVReadAndWriteE15.vb

```
Imports System.IO
Imports System.Reflection

''' <summary>
''' CSVファイルへの書き込みと読み込みを行います。
''' </summary>
''' <remarks></remarks>
Public Class CSVReadAndWriteE15

    ' Refactored by E.Yoshimoto in Feb. 2018.

    ''' <summary>
    ''' テキストファイルのフォーマットを指定します。
    ''' </summary>
    Private Shared SHIFT_JIS As System.Text.Encoding = System.Text.Encoding.GetEncoding("Shift_JIS")

    ''' <summary>
    ''' 数値のフォーマットを指定します。
    ''' </summary>
    ''' <remarks></remarks>
    Private Shared FORMAT As String = "E15"

    ' 複数ファイルの書き込みに対応するリストです。
    Private Shared listOfFiles As New List(Of String)
    Private Shared listOfCounters As New List(Of Integer)

    ''' <summary>
    ''' コンストラクタを不可視にします。
    ''' </summary>
    Private Sub New()
    End Sub

    ''' <summary>
    ''' ファイル名を指定してdataArrayの内容を時系列を含めてCSVファイルに追記します。
    ''' charttime = 0 のとき、ファイルを新規作成します。
    ''' </summary>
    Public Shared Sub CreateNewCSV(filepath As String, data As Object,
                                   Optional ByVal names() As String = Nothing,
                                   Optional ByVal checkNames As Boolean = True)

        ' データが空の場合、メッセージを出して終了します。
        If data Is Nothing Then
            MessageBox.Show("データがありません。")
            Exit Sub
        End If

        ' names() の設定が不正の場合、メッセージを出して終了します。
        If Not names Is Nothing AndAlso checkNames AndAlso Not AllNamesExist(data, names) Then
            Exit Sub
        End If

        ' ファイルのリストを参照し、カウンタの値を取得します。
        Dim fileIndex As Integer = listOfFiles.IndexOf(filepath)

        ' カウンタの値を初期化します。
        If fileIndex < 0 Then
            listOfFiles.Add(filepath)
            fileIndex = listOfFiles.Count - 1
            listOfCounters.Add(0)
        Else
            listOfCounters(fileIndex) = 0
        End If

        ' 出力用StreamWriterを定義します。
        Dim writer As StreamWriter = Nothing

        Try
            ' StreamWriterをインスタンス化します。
            writer = New StreamWriter(filepath, False, SHIFT_JIS)

            ' ヘッダを書き込み、フィールド情報を取得します。
            WriteCSVHeader(writer, data, names)

        Catch e As Exception
            ' 例外発生時にエラーメッセージを表示します。
            MessageBox.Show("ファイルの書き込みに失敗しました。" & Environment.NewLine & e.Message)

        Finally
            ' ファイルを閉じます。
            If Not writer Is Nothing Then
                writer.Close()
            End If
        End Try

    End Sub

    ''' <summary>
    ''' ファイル名を指定してdataArrayの内容を時系列を含めてCSVファイルに追記します。
    ''' charttime = 0 のとき、ファイルを新規作成します。
    ''' </summary>
    Public Shared Sub WriteCSV(filepath As String, chartTime As Double, data As Object, interval As Integer,
                               Optional ByVal names() As String = Nothing,
                               Optional ByVal checkNames As Boolean = True)

        ' データが空の場合、メッセージを出して終了します。
        If data Is Nothing Then
            MessageBox.Show("データがありません。")
            Exit Sub
        End If

        ' names() の設定が不正の場合、メッセージを出して終了します。
        If Not names Is Nothing AndAlso checkNames AndAlso Not AllNamesExist(data, names) Then
            Exit Sub
        End If

        ' ファイルのリストを参照し、カウンタの値を取得します。
        Dim fileIndex As Integer = listOfFiles.IndexOf(filepath)
        Dim counter As Integer

        ' カウンタの値を設定します。
        If fileIndex < 0 Then
            listOfFiles.Add(filepath)
            fileIndex = listOfFiles.Count - 1
            listOfCounters.Add(0)
            counter = 0
        Else
            counter = listOfCounters(fileIndex)
        End If

        ' 出力用StreamWriterを定義します。
        Dim writer As StreamWriter = Nothing

        Try
            ' 計算開始時のみファイルを新規作成し、ヘッダを書き込みます。
            If chartTime = 0 Then
                ' StreamWriterをインスタンス化します。
                writer = New StreamWriter(filepath, False, SHIFT_JIS)
                ' ヘッダを書き込み、フィールド情報を取得します。
                WriteCSVHeader(writer, data, names)
                ' カウンタをリセットします。
                counter = 0
            Else
                ' StreamWriterを追記モードでインスタンス化します。
                writer = New StreamWriter(filepath, True, SHIFT_JIS)
            End If

            ' カウンタが0のとき出力します。
            If counter = 0 Then
                ' データ1行を書き込みます。
                WriteCSVDataLine(writer, chartTime, data, names)
            End If

            ' カウンタの更新を行います。
            If counter >= interval - 1 Then
                ' カウンタをリセットします。
                counter = 0
            Else
                ' カウンタを更新します。
                counter += 1
            End If

            ' リスト内のカウンタの値を更新します。
            listOfCounters(fileIndex) = counter

        Catch e As Exception
            ' 例外発生時にエラーメッセージを表示します。
            MessageBox.Show("ファイルの書き込みに失敗しました。" & Environment.NewLine & e.Message)

        Finally
            ' ファイルを閉じます。
            If Not writer Is Nothing Then
                writer.Close()
            End If
        End Try

    End Sub

    ''' <summary>
    ''' 変数名を指定する配列の中身にオブジェクトの変数名が含まれているかどうかを確認します。
    ''' </summary>
    Private Shared Function AllNamesExist(data As Object, names() As String) As Boolean

        ' 要素のTypeを取得します。
        Dim t As Type = data.GetType()

        ' Publicフィールド名を取得します。
        Dim fields As FieldInfo()
        If t.IsArray Then
            fields = t.GetElementType().GetFields()
        Else
            fields = t.GetFields()
        End If

        ' フィールド名のリストを作成します。
        Dim fieldNames As New HashSet(Of String)
        For Each field As FieldInfo In fields
            fieldNames.Add(field.Name)
        Next

        ' fieldNames 内に存在しない変数名が names() 内に存在する場合、エラーを発生させます。
        Dim errorNames As List(Of String) = New List(Of String)
        For Each name As String In names
            If Not fieldNames.Contains(name) Then
                errorNames.Add(name)
            End If
        Next

        If errorNames.Count > 0 Then

            ' エラーメッセージを作成します。
            Dim message As String = "以下の変数を出力できませんでした。出力する変数の名前を確認してください。"

            For Each errorName As String In errorNames
                message &= """" & errorName & """, "
            Next

            message = message.Substring(0, message.Length - 2)

            ' メッセージを表示します。
            MessageBox.Show(message)

        End If

        Return (errorNames.Count = 0)

    End Function

    ''' <summary>
    ''' CSVファイルのヘッダを出力します。
    ''' </summary>
    Private Shared Sub WriteCSVHeader(writer As StreamWriter, data As Object, names() As String)

        ' 先頭に時刻を出力します。
        writer.Write("""time"",")

        ' 要素のTypeを取得します。
        Dim t As Type = data.GetType()

        ' Publicフィールド名を取得します。
        Dim fields As FieldInfo()
        If t.IsArray Then
            fields = t.GetElementType().GetFields()
        Else
            fields = t.GetFields()
        End If

        If t.IsArray Then
            ' 配列データを一行に出力します。
            Dim dataArray As Array = DirectCast(data, Array)
            For j As Integer = 0 To dataArray.GetUpperBound(0)
                For Each field As FieldInfo In fields

                    ' 値型と文字列型のみ出力します。
                    If field.FieldType.IsValueType OrElse field.FieldType Is GetType(String) Then

                        ' 定数を除外します。
                        If Not field.IsLiteral AndAlso Not field.IsInitOnly Then

                            ' 変数名が指定されている場合には対応する変数のみ出力します。
                            Dim fieldName = field.Name
                            If names Is Nothing OrElse names.Contains(fieldName) Then
                                ' 変数名を出力します。
                                writer.Write("""" & fieldName & "(" & j & ")"",")
                            End If

                        End If

                    End If

                Next
            Next
        Else
            ' データを一行に出力します。
            For Each field As FieldInfo In fields

                ' 値型と文字列型のみ出力します。
                If field.FieldType.IsValueType OrElse field.FieldType Is GetType(String) Then

                    ' 定数を除外します。
                    If Not field.IsLiteral AndAlso Not field.IsInitOnly Then

                        ' 変数名が指定されている場合には対応する変数のみ出力します。
                        Dim fieldName = field.Name
                        If names Is Nothing OrElse names.Contains(fieldName) Then
                            ' 変数名を出力します。
                            writer.Write("""" & fieldName & """,")
                        End If

                    End If

                End If

            Next
        End If

        ' 改行します。
        writer.WriteLine()

    End Sub

    ''' <summary>
    ''' CSVファイルのデータ1行を出力します。
    ''' </summary>
    Private Shared Sub WriteCSVDataLine(writer As StreamWriter, charttime As Double, data As Object, names() As String)

        ' 要素のTypeを取得します。
        Dim t As Type = data.GetType()

        ' Publicフィールド名を取得します。
        Dim fields As FieldInfo()
        If t.IsArray Then
            fields = t.GetElementType().GetFields()
        Else
            fields = t.GetFields()
        End If

        ' 時刻を出力します。
        writer.Write("""" & charttime.ToString(FORMAT) & """,")

        If data.GetType().IsArray Then
            ' 配列データを一行に出力します。
            Dim dataArray As Array = DirectCast(data, Array)
            For j As Integer = 0 To dataArray.GetUpperBound(0)
                For Each field As FieldInfo In fields

                    ' 定数を除外します。
                    If Not field.IsLiteral AndAlso Not field.IsInitOnly Then

                        ' 変数名が指定されている場合には対応する変数のみ出力します。
                        Dim fieldName = field.Name
                        If names Is Nothing OrElse names.Contains(fieldName) Then

                            ' 値型と文字列型のみ出力します。
                            If field.FieldType Is GetType(String) OrElse
                                field.FieldType Is GetType(Boolean) Then
                                ' 値を出力します。
                                writer.Write("""" & field.GetValue(dataArray(j)) & """,")
                            ElseIf field.FieldType.IsValueType Then
                                ' 値を出力します。
                                writer.Write("""" & CDbl(field.GetValue(dataArray(j))).ToString(FORMAT) & """,")
                            End If

                        End If

                    End If

                Next
            Next
        Else
            ' 各データを出力します。
            For Each field As FieldInfo In fields

                ' 定数を除外します。
                If Not field.IsLiteral AndAlso Not field.IsInitOnly Then

                    ' 変数名が指定されている場合には対応する変数のみ出力します。
                    Dim fieldName = field.Name
                    If names Is Nothing OrElse names.Contains(fieldName) Then

                        ' 値型と文字列型のみ出力します。
                        If field.FieldType Is GetType(String) OrElse
                            field.FieldType Is GetType(Boolean) Then
                            ' 値を出力します。
                            writer.Write("""" & field.GetValue(data) & """,")
                        ElseIf field.FieldType.IsValueType Then
                            ' 値を出力します。
                            writer.Write("""" & CDbl(field.GetValue(data)).ToString(FORMAT) & """,")
                        End If

                    End If

                End If

            Next
        End If

        ' 改行します。
        writer.WriteLine()

    End Sub

    ''' <summary>
    ''' CSVファイルを読み込み、値を代入します。
    ''' </summary>
    Public Overloads Shared Sub ReadCSV(filepath As String, data As Object, Optional names() As String = Nothing)

        ' ファイルがないときは何もせず終了します。
        If Not File.Exists(filepath) Then
            Exit Sub
        End If

        ' インスタンス化されていない場合、メッセージを出して終了します。
        If data Is Nothing Then
            MessageBox.Show("配列が空、または要素がインスタンス化されていません。")
            Exit Sub
        End If

        ' 入力用TextFieldParserを定義します。
        Dim reader As FileIO.TextFieldParser = Nothing

        Try
            ' TextFieldParserをインスタンス化します。
            reader = New FileIO.TextFieldParser(filepath, SHIFT_JIS)

            ' 区切り文字「,」を設定します。
            reader.TextFieldType = FileIO.FieldType.Delimited
            reader.SetDelimiters(",")

            ' Typeを取得します。
            Dim t As Type = data.GetType()

            ' Publicフィールド名を取得します。
            Dim fields As FieldInfo() = t.GetFields()

            ' 1行目の変数名を読み込みます。
            Dim header As String() = reader.ReadFields()

            ' 各要素とCSVデータとの対応付けを行います。
            Dim index(header.GetUpperBound(0)) As Integer
            For i As Integer = 0 To header.GetUpperBound(0)
                index(i) = -1
            Next
            For i As Integer = 1 To header.GetUpperBound(0)
                For j As Integer = 0 To fields.GetUpperBound(0)
                    If header(i) = fields(j).Name Then
                        index(i) = j
                        Exit For
                    End If
                Next
            Next

            ' ファイルを最終行まで読み込みます。
            While Not reader.EndOfData

                ' データを1行読み込みます。
                Dim row As String() = reader.ReadFields()

                ' 各フィールドのデータを取得し、代入します。 
                For i As Integer = 1 To row.GetUpperBound(0)

                    ' 空のフィールドをスキップします。
                    If index(i) >= 0 AndAlso Not row(i) = String.Empty Then

                        ' 参照フィールドを取得します。
                        Dim field As FieldInfo = fields(index(i))

                        ' 変数名が指定されている場合には対応する変数のみ読み込みます。
                        If names Is Nothing OrElse names.Contains(field.Name) Then

                            ' 変数の値を代入します。
                            SetValue(field, row(i), data)

                        End If

                    End If

                Next

            End While

        Catch e As Exception
            ' 例外発生時にエラーメッセージを表示します。
            MessageBox.Show("ファイルの読み込みに失敗しました。" & Environment.NewLine & e.Message)

        Finally
            ' ファイルを閉じます。
            If Not reader Is Nothing Then
                reader.Close()
            End If
        End Try

    End Sub

    ''' <summary>
    ''' CSVファイルを読み込み、値を配列に代入します。
    ''' </summary>
    Public Overloads Shared Sub ReadCSV(filepath As String, dataArray As Object(), Optional names() As String = Nothing)

        ' ファイルがないときは何もせず終了します。
        If Not File.Exists(filepath) Then
            Exit Sub
        End If

        ' 配列が空、またはインスタンス化されていない場合、メッセージを出して終了します。
        If dataArray.Length = 0 OrElse dataArray(0) Is Nothing Then
            MessageBox.Show("配列が空、または要素がインスタンス化されていません。")
            Exit Sub
        End If

        ' 入力用TextFieldParserを定義します。
        Dim reader As FileIO.TextFieldParser = Nothing

        Try
            ' TextFieldParserをインスタンス化します。
            reader = New FileIO.TextFieldParser(filepath, SHIFT_JIS)

            ' 区切り文字「,」を設定します。
            reader.TextFieldType = FileIO.FieldType.Delimited
            reader.SetDelimiters(",")

            ' 配列の要素のTypeを取得します。
            Dim t As Type = dataArray.GetType().GetElementType()

            ' Publicフィールド名を取得します。
            Dim fields As FieldInfo() = t.GetFields()

            ' 1行目の変数名を読み込みます。
            Dim header As String() = reader.ReadFields()

            ' 1細胞単位のデータ長を算出します。
            Dim length As Integer = Array.IndexOf(header, header(1).Substring(0, header(1).Length - 3) & "(1)") - 1
            If length < 0 Then length = header.Length - 1

            Dim index(header.GetUpperBound(0)) As Integer
            For i As Integer = 0 To length
                index(i) = -1
            Next
            For i As Integer = 1 To length
                For j As Integer = 0 To fields.GetUpperBound(0)
                    If header(i) = fields(j).Name & "(0)" Then
                        index(i) = j
                        Exit For
                    End If
                Next
            Next

            ' すべての要素の対応付けを行います。
            For i As Integer = length + 1 To header.GetUpperBound(0)
                index(i) = index(i - length)
            Next

            ' ファイルを最終行まで読み込みます。
            While Not reader.EndOfData

                ' データを1行読み込みます。
                Dim row As String() = reader.ReadFields()

                ' 各フィールドのデータを取得し、代入します。 
                For i As Integer = 1 To row.GetUpperBound(0)

                    ' 空のフィールドをスキップします。
                    If index(i) >= 0 AndAlso Not row(i) = String.Empty Then

                        ' 参照フィールドを取得します。
                        Dim field As FieldInfo = fields(index(i))

                        ' 変数名が指定されている場合には対応する変数のみ読み込みます。
                        If names Is Nothing OrElse names.Contains(field.Name) Then

                            ' 変数の値を代入します。
                            SetValue(field, row(i), dataArray((i - 1) \ length))

                        End If

                    End If

                Next

            End While

        Catch e As Exception
            ' 例外発生時にエラーメッセージを表示します。
            MessageBox.Show("ファイルの読み込みに失敗しました。" & Environment.NewLine & e.Message)

        Finally
            ' ファイルを閉じます。
            If Not reader Is Nothing Then
                reader.Close()
            End If
        End Try

    End Sub

    ''' <summary>
    ''' オブジェクトのフィールドへ値を代入します。
    ''' </summary>
    Private Shared Sub SetValue(field As FieldInfo, valueString As String, obj As Object)
        If field.FieldType Is GetType(Double) Then
            ' Double型への変換を行います。
            Dim value As Double = Double.Parse(valueString)
            field.SetValue(obj, value)
        ElseIf field.FieldType Is GetType(Integer) Then
            ' Integer型への変換を行います。
            Dim value As Integer = Integer.Parse(valueString)
            field.SetValue(obj, CInt(value))
        ElseIf field.FieldType Is GetType(Decimal) Then
            ' Decimal型への変換を行います。
            Dim value As Decimal = Decimal.Parse(valueString)
            field.SetValue(obj, CDec(value))
        ElseIf field.FieldType Is GetType(Long) Then
            ' Long型への変換を行います。
            Dim value As Long = Long.Parse(valueString)
            field.SetValue(obj, CLng(value))
        ElseIf field.FieldType Is GetType(Boolean) Then
            ' Boolean型への変換を行います。
            Dim value As Boolean = Boolean.Parse(valueString)
            field.SetValue(obj, CBool(value))
        ElseIf field.FieldType Is GetType(String) Then
            ' String型で読み込みます。
            field.SetValue(obj, valueString)
        Else
            Throw New IOException("変換対象外の型が含まれています。")
        End If
    End Sub

End Class
```
