## Supplementary material for "Gradient-based parameter optimization to determine membrane ionic current composition of human induced pluripotent stem cell-derived cardiomyocytes": optimizing program: Graph2018.vb

```
Public Class Graph2018
    Inherits PictureBox
    ' Created by CY Cha in October 2008, revised by AN in November and by E.Santos in April 2009
    ' Refactored by E.Yoshimoto in Feb. 2018.

    Public g As Graphics                '   'g' is used as Graphics 

    Private minX As Double            'minimum value on the abscissa
    Private maxX As Double           'maximum value on the abscissa
    Private Xorigin As Double         'X address to give the minimum value

    Private minY As Double           'minimum value on the ordinate
    Private maxY As Double          'maximum value on the ordinate
    Private Yorigin As Double        'Y address to give the minimum value

    Private GOriginX As Integer        'X address of the left upper corner of the graph
    Private GOriginY As Integer        'Y address of the left upper corner of the graph

    Private GWidthX As Integer        'width of the graph
    Private GWidthY As Integer        'height of the graph

    Private UnitX As Double              'scaling factor
    Private UnitY As Double              'scaling factor

    Private PBorigin As PictureBox
    Private flushTimer As New Timer
    Private frameRate As Double = 30

    ''' <summary>
    ''' Default instance of Graph
    ''' </summary>
    Public Sub New()
        PBorigin = Me
        BackColor = Color.White
    End Sub

    ''' <summary>
    '''  Make instance of  Graph
    ''' </summary>
    Public Sub New(ByRef PB As PictureBox, ByVal min_X As Double, ByVal max_X As Double, ByVal min_Y As Double, ByVal max_Y As Double)
        PBorigin = PB
        Initialize(min_X, max_X, min_Y, max_Y)
    End Sub

    ''' <summary>
    ''' Initialize Graph
    ''' </summary>
    Public Sub Initialize(ByVal min_X As Double, ByVal max_X As Double, ByVal min_Y As Double, ByVal max_Y As Double, Optional ByVal antiAlias As Boolean = True)

        PBorigin.Image = New Bitmap(PBorigin.ClientRectangle.Width, PBorigin.ClientRectangle.Height)

        flushTimer.Interval = 1000 / frameRate

        AddHandler flushTimer.Tick, AddressOf flushBuffer

        g = Graphics.FromImage(PBorigin.Image)
        If antiAlias Then
            g.SmoothingMode = Drawing2D.SmoothingMode.AntiAlias
            g.TextRenderingHint = Drawing.Text.TextRenderingHint.AntiAlias
        End If

        GOriginX = PBorigin.Width * 0.1                  'define the X address of the origin 
        GOriginY = PBorigin.Height * 0.9                  'define the Y address of the origin

        GWidthX = PBorigin.Width * 0.85                'define the width of the graph
        GWidthY = PBorigin.Height * 0.85               'define the height of the graph

        minX = min_X                                       'give the minimum value to the left end
        maxX = max_X                                    'give the maximum value to the right end

        minY = min_Y                                        'give the minimum value to the bottom end
        maxY = max_Y                                      'give the maximum value to the top end


        UnitX = GWidthX / (maxX - minX)        'scaling constant on X axis
        UnitY = -GWidthY / (maxY - minY)       'scaling constant on Y axis

    End Sub

    Public Sub DrawAxis(ByVal IntervalX As Double, ByVal IntervalY As Double, ByVal gTitle As String, ByVal FontSize As Integer, Optional ByVal formatString As String = "###0.###")

        Dim AxisPen As Pen = New Pen(Color.Cyan, 1)
        Dim blackpen As Pen = New Pen(Color.Black, 1)
        Dim MyFont As Font = New Font("Arial", FontSize)
        Dim MyBrush As Brush = New SolidBrush(Color.Black)
        Dim label As String

        '***************************************************************************************************
        Dim gX1, gY1, gX2, gY2 As Integer                'coordinate of x and y
        Dim valueX As Double
        Dim valueY As Double

        'g.Clear(Color.White)              'Clear the picture box using a white color
        valueX = minX

        Do                                                                        'vertical line
            gX1 = GOriginX + UnitX * (valueX - minX)
            gY1 = GOriginY

            gX2 = gX1
            gY2 = GOriginY + UnitY * (maxY - minY)

            g.DrawLine(AxisPen, gX1, gY1, gX2, gY2)

            'label = CStr(valueX)
            label = Format(valueX, formatString)                           'corrected by AN on 2019,0512                
            g.DrawString(label, MyFont, MyBrush, gX1 - FontSize, gY1 + CInt(FontSize * 0.5))

            valueX = valueX + IntervalX
        Loop While valueX <= maxX 'この数値まで繰り返し

        valueY = minY
        Do                                                                          'horizontal line
            gX1 = GOriginX
            gY1 = GOriginY + UnitY * (valueY - minY)

            gX2 = GOriginX + UnitX * (maxX - minX)
            gY2 = gY1

            g.DrawLine(AxisPen, gX1, gY1, gX2, gY2)

            label = Format(valueY, formatString)
            g.DrawString(label, MyFont, MyBrush, gX1 - 3 * FontSize, gY1 - CInt(FontSize * 0.5))

            valueY = valueY + IntervalY
        Loop While valueY <= maxY
        valueY = 0
        gY1 = GOriginY + UnitY * (valueY - minY)
        gY2 = gY1
        g.DrawLine(blackpen, gX1, gY1, gX2, gY2)


        gX1 = GOriginX
        gY1 = GOriginY + UnitY * (maxY - minY)
        g.DrawString(gTitle, MyFont, MyBrush, gX1, gY1 - CInt(FontSize * 0.5))

        flushTimer.Enabled = True

    End Sub

    Public Overloads Sub PlotLine(ByVal valueXL As Double, ByVal valueYL As Double, ByVal valueXR As Double, ByVal valueYR As Double, ByVal color As Pen)
        Dim gXL, gYL As Integer
        Dim gXR, gYR As Integer

        Try
            gXL = GOriginX + UnitX * (valueXL - minX)
            gYL = GOriginY + UnitY * (valueYL - minY)
            gXR = GOriginX + UnitX * (valueXR - minX)
            gYR = GOriginY + UnitY * (valueYR - minY)
            g.DrawLine(color, gXL, gYL, gXR, gYR)

            flushTimer.Enabled = True
        Catch exc As Exception
        End Try

    End Sub
    Public Overloads Sub PlotLine(ByVal valueXL As Double, ByVal valueYL As Double, ByVal valueXR As Double, ByVal valueYR As Double, ByVal color As Color, ByVal width As Integer)
        PlotLine(valueXL, valueYL, valueXR, valueYR, New Pen(color, width))
    End Sub

    Public Overloads Sub PlotPoint(ByVal valueX As Double, ByVal valueY As Double, ByVal color As Pen, Optional ByVal width As Integer = -1)
        Dim gX1, gY1 As Integer
        'revised on 150509
        Try
            gX1 = GOriginX + UnitX * (valueX - minX) - color.Width / 2
            gY1 = GOriginY + UnitY * (valueY - minY) - color.Width / 2

            If width <= 0 Then
                g.FillEllipse(New SolidBrush(color.Color), gX1, gY1, color.Width, color.Width)            '丸を書く
            Else
                g.DrawEllipse(New Pen(color.Color, width), gX1, gY1, color.Width, color.Width)
            End If

            flushTimer.Enabled = True

        Catch exc As Exception
        End Try

    End Sub
    Public Overloads Sub PlotPoint(ByVal valueX As Double, ByVal valueY As Double, ByVal color As Color, ByVal pointWidth As Integer, Optional ByVal width As Integer = -1)
        PlotPoint(valueX, valueY, New Pen(color, pointWidth), width)
    End Sub

    Public Overloads Sub PlotFillPoint(ByVal valueX As Double, ByVal valueY As Double, ByVal color As SolidBrush, ByVal width As Integer)
        Dim gX1, gY1 As Integer

        Try
            If width > 1 AndAlso width Mod 2 = 1 Then
                width = width - 1
            End If

            gX1 = GOriginX + UnitX * (valueX - minX) - width / 2
            gY1 = GOriginY + UnitY * (valueY - minY) - width / 2
            g.FillEllipse(color, gX1, gY1, width, width)

            flushTimer.Enabled = True
        Catch exc As Exception
        End Try

    End Sub
    Public Overloads Sub PlotFillPoint(ByVal valueX As Double, ByVal valueY As Double, ByVal color As Color, ByVal width As Integer)
        PlotFillPoint(valueX, valueY, New SolidBrush(color), width)
    End Sub

    Public Overloads Sub PlotRectangle(ByVal color As Pen, ByVal valueX As Double, ByVal valueY As Double, ByVal width As Double, ByVal height As Double)
        Dim gX1, gY1, gW, gH As Integer

        Try
            gX1 = GOriginX + UnitX * (valueX - minX)
            gY1 = GOriginY + UnitY * (valueY - minY)
            gW = UnitX * width
            gH = -UnitY * height
            g.DrawRectangle(color, gX1, gY1, gW, gH)                      'rectangleを書く

            flushTimer.Enabled = True
        Catch exc As Exception
        End Try

    End Sub
    Public Overloads Sub PlotRectangle(ByVal color As Color, ByVal penWidth As Integer, ByVal valueX As Double, ByVal valueY As Double, ByVal width As Double, ByVal height As Double)
        PlotRectangle(New Pen(color, penWidth), valueX, valueY, width, height)
    End Sub

    Public Sub Clear()
        g.Clear(color.White)
        flushTimer.Enabled = True
    End Sub

    Public Overloads Sub LogPlotPoint(ByVal valueX As Double, ByVal valueY As Double, ByVal minimumValue As Double, ByVal color As Pen)
        Dim gX1, gY1 As Integer

        gX1 = GOriginX + UnitX * (valueX - minX)
        gY1 = GOriginY + UnitY * Math.Log10(valueY / minimumValue)
        g.DrawEllipse(color, gX1, gY1, 1, 1) '楕円を書く

        flushTimer.Enabled = True

    End Sub
    Public Overloads Sub LogPlotPoint(ByVal valueX As Double, ByVal valueY As Double, ByVal minimumValue As Double, ByVal color As Color, ByVal width As Integer)
        LogPlotPoint(valueX, valueY, minimumValue, New Pen(color, width))
    End Sub

    Public Overloads Sub PlotVerticalLine(ByVal valueX As Double, ByVal minY1 As Double, ByVal maxY1 As Double, ByVal color As Pen)
        Dim gX1 As Integer
        Dim gY1 As Integer
        Dim gX2 As Integer
        Dim gY2 As Integer

        gX1 = GOriginX + UnitX * (valueX - minX)
        gY1 = GOriginY + UnitY * (maxY - minY)
        gX2 = GOriginX + UnitX * (valueX - minX)
        gY2 = GOriginY + UnitY * (minY - minY)
        g.DrawLine(color, gX1, gY1, gX2, gY2)

        g.DrawEllipse(color, gX1, gY1, 3, 3)

        flushTimer.Enabled = True
    End Sub
    Public Overloads Sub PlotVerticalLine(ByVal valueX As Double, ByVal minY1 As Double, ByVal maxY1 As Double, ByVal color As Color, ByVal width As Integer)
        PlotVerticalLine(valueX, minY1, maxY1, New Pen(color, width))
    End Sub

    ''' <summary>
    ''' 色の配列と太さを指定して Pen の配列を生成する関数です。PlotArea で使用します。
    ''' </summary>
    Public Overloads Function CreatePensArray(ByVal colorArray() As Color, ByVal width As Integer) As Pen()

        Dim penArray(colorArray.Length - 1) As Pen

        For i As Integer = 0 To colorArray.Length - 1
            penArray(i) = New Pen(colorArray(i), width)
        Next

        Return penArray
    End Function

    ''' <summary>
    ''' 色の配列と太さの配列を指定して Pen の配列を生成する関数です。PlotArea で使用します。
    ''' </summary>
    Public Overloads Function CreatePensArray(ByVal colorArray() As Color, ByVal widthArray() As Integer) As Pen()

        Dim penArray(colorArray.Length - 1) As Pen

        For i As Integer = 0 To colorArray.Length - 1
            penArray(i) = New Pen(colorArray(i), widthArray(i))
        Next

        Return penArray
    End Function

    Public Overloads Sub PlotArea(ByVal valueX As Double, ByVal currentsArray() As Double, ByVal pensArray() As Pen)
        plotArea(valueX, currentsArray, pensArray, False, 0)
    End Sub
    Public Overloads Sub PlotArea(ByVal valueX As Double, ByVal currentsArray() As Double, ByVal pensArray() As Pen, ByVal total As Boolean)
        PlotArea(valueX, currentsArray, pensArray, True, total)
    End Sub
    Public Overloads Sub PlotArea(ByVal valueX As Double, ByVal currentsArray() As Double, ByVal pensArray() As Pen, ByVal percent As Boolean, ByVal total As Double)
        Dim gX1, gY1, gY2, pos, neg As Double

        gX1 = GOriginX + UnitX * (valueX - minX)

        'If we want to plot the graph in % and no total is specified we sum all the currents
        If percent And total = 0 Then
            For i = 0 To currentsArray.Length - 1
                total += Math.Abs(currentsArray(i))
            Next
        End If


        For i = 0 To currentsArray.Length - 1
            If (Math.Sign(currentsArray(i)) = 1) Then
                'Positive
                gY1 = GOriginY + UnitY * (pos - minY)
                If percent Then
                    'If we are plotting in percentage
                    pos = pos + currentsArray(i) / total * 100
                Else
                    'If we are plotting in absoulte value
                    pos = pos + currentsArray(i)
                End If
                gY2 = GOriginY + UnitY * (pos - minY)
            Else
                'Negative
                gY1 = GOriginY + UnitY * (neg - minY)
                If percent Then
                    neg = neg + currentsArray(i) / total * 100
                Else
                    neg = neg + currentsArray(i)
                End If
                gY2 = GOriginY + UnitY * (neg - minY)
            End If
            g.DrawLine(pensArray(i), CInt(gX1), CInt(gY1), CInt(gX1), CInt(gY2))
        Next
        flushTimer.Enabled = True
    End Sub

    Private Sub flushBuffer(ByVal Sender As Object, ByVal e As EventArgs)
        PBorigin.Invalidate()
        flushTimer.Enabled = False
    End Sub

    Public Class GradientPen
        Public minGradient As Double = 1000000
        Public maxGradient As Double = -1000000
        Dim colorValue As Double
        Dim gradientPen As Pen

        Public Sub New(ByVal width As Integer)
            gradientPen = New Pen(Drawing.Color.Red, width)
        End Sub

        Public Overloads Function Pen(ByVal gradientVariable As Double)
            If gradientVariable >= maxGradient Then
                maxGradient = gradientVariable
            ElseIf gradientVariable < minGradient Then
                minGradient = gradientVariable
            End If

            colorValue = Math.Round(((gradientVariable - minGradient) / (maxGradient - minGradient)) * 255)

            If colorValue > 255 Then
                colorValue = 255
            ElseIf colorValue < 0 Then
                colorValue = 0
            End If

            gradientPen.Color = Drawing.Color.FromArgb(colorValue, 0, 255 - colorValue)
            Return gradientPen

        End Function

        Public Overloads Function PenRedBlue(ByVal gradientVariable As Double, ByVal min As Double, ByVal max As Double)
            colorValue = Math.Round(((gradientVariable - min) / (max - min)) * 255)

            If colorValue > 255 Then
                colorValue = 255
            ElseIf colorValue < 0 Then
                colorValue = 0
            End If

            gradientPen.Color = Drawing.Color.FromArgb(colorValue, 0, 255 - colorValue)

            Return gradientPen

        End Function

        Public Function PenGreenBlue(ByVal gradientVariable As Double, ByVal min As Double, ByVal max As Double)
            colorValue = Math.Round(((gradientVariable - min) / (max - min)) * 255)

            If colorValue > 255 Then
                colorValue = 255
            ElseIf colorValue < 0 Then
                colorValue = 0
            End If

            gradientPen.Color = Drawing.Color.FromArgb(0, colorValue, 255 - colorValue)

            Return gradientPen

        End Function

        Public Sub startCalibration()
            minGradient = 1000000
            maxGradient = -1000000
        End Sub
    End Class

End Class
```
