## Supplementary material for "Gradient-based parameter optimization to determine membrane ionic current composition of human induced pluripotent stem cell-derived cardiomyocytes": optimizing program: IPSmodel1022.vb

```
Imports System.Math
Imports System.IO

Public Class IPSmodel1022

    Public constraints As Boolean = False                       'if constraints are used or not in the Pattern Search. Under control usage, no constraints.

    '********** Physical Contents **********
    Public Const R As Double = 8.3143                                   ' mV C mmol-1 K-1     Gas constant   
    Public Const Faraday As Double = 96.4867                       'C mmol-1                   Faraday const.
    Public Const tempK As Double = 310.15                                'K
    Public Const RTF As Double = R * tempK / Faraday
    Public Const RTF2 As Double = R * tempK / Faraday / 2

    '********** Cell size and related Parameters **********
    Public Cm As Double = 32              '177     '138.1 * ratioCm              'pF

    '********************************   External ion concentrations ************************
    Public Ko As Double = 5.4    'External K   concentration,   [K]o
    Public Nao As Double = 145    'External Na concentration,   [Na]o
    Public Cao As Double = 1.8  ' External Ca concentration,   [Ca]o

    Public newKo As Double = 5.4         'new External K   concentration,   [K]o
    Public newNao As Double = 145        'new External Na concentration,   [Na]o
    Public newCao As Double = 1.8        'new External Ca concentration,   [Ca]o

    '*********** Heart Rate **********************
    Public HR As Double                'heart rate   min-1
    '*****        ******
    Public Const Vol_cell As Double = 2510                    'fL
    '*******************
    Public Vol_blk As Double = 0.65 * Vol_cell                               'fL            the same as in the V cell model
    Public Vol_iz As Double = 0.035 * Vol_cell                     '0.04 * Vol_cell       'fL      the same as in the V cell model
    Public Vol_jnc As Double = 0.008 * Vol_cell                 '0.011    fL        the same as in the V cell model
    Public Vol_cyt As Double = Vol_jnc + Vol_iz + Vol_blk             'fL           the same as in the V cell model
    '************** total volume of cytosolic ionic space = 69.3 % of the whole cell volume  .  The rest is mostly occupied by mitochondria
    '*********** total SR volume = Vol_SRrl + Vol_SRup = 1.1% of the whole cell volume *******************
    Public Vol_SRt As Double = 0.011 * Vol_cell           'fL  ref. 0.06 in the ventricular cell model
    Public Vol_SRrl As Double = 0.1 * Vol_SRt              'fL       smaller than in the V cell model
    Public Vol_SRup As Double = 0.9 * Vol_SRt             'fL       larger than  in the V cell model
    '****************** parameters used to include the mitochondria model ****************
    '********** Fraction of Currents jnc, iz, blk **********
    Public Frc_iz As Double = 0.1         'fractional  distribution at the iz myoplasm
    Public Frc_blk As Double = 0.9       'fractional  distribution at the iz myoplasm

    Public FrcCaL_jnc As Double = 0.15                     'fractional distribution;分画散布 of ICaL at the junctional myoplasm
    Public FrcCaL_iz As Double = 0.45                       '0.4fractional distribution of ICaL at the iz myoplasm
    Public FrcCaL_blk As Double = 0.4                       '0.35fractional distribution of ICaL at the blk myoplasm

    Public FrcNCX_jnc = 0                                                        'fractional distribution of NCX at the junctional myoplasm
    Public FrcNCX_iz = 0.1                                            'fractional distribution of NCX at the iz myoplasm
    Public FrcNCX_blk = 0.9                                          'fractional distribution of NCX at the blk myoplasm

    '********** Membrane permeability

    Dim maxIPMCA As Double = 0.1615

    '************* SR設定 *********************
    Public ampISERCA As Double = 1512 * 0.7         '50400 * 0.03 * 0.6        '0.1 in case of using SERCA_TC    
    Public Ptrans As Double = 0.068235 / 4               '0.017            '22.745  * 0.005    
    Public PRyR As Double = 2.12 * 2                        '* 1.3     3532 * 0.03 * 0.01             '3532 * 0.0001 0.004    

    Public po_baseRyR As Double = 0.000075       '0.000075 * SRsaFactor / 300leak is also scaled for the SA Cell

    '********** Membrane potential **************************************************************************************************************
    Public Vm As Double               'mV, membrane potential
    Public tgVm As Double           'mv, the experimental Vm 
    Public baseVshift As Double   'mV, correcting  experimental recording of Vm
    Public ExtraVshift As Double 'mV, adjustable component for the correcting  experimental recording of Vm

    Public dVmdt As Double          'dy/dt
    Public Itot_Ca As Double         'total current carried by Calcium ion
    Public Itot_Na As Double         'total current carried by Sodium ion
    Public Itot_K As Double            'total current carried by Pottasium ion
    Public Itot_cell As Double         'total current

    '*********  cytosolic Ion Concentrations    *******************************
    Public ItotCa_jnc As Double
    Public ItotCa_iz As Double
    Public ItotCa_blk As Double

    Public Catot_jnc As Double
    Public Catot_iz As Double
    Public Catot_blk As Double

    Public P1_Catot_blk As Double
    Public P1_Catot_jnc As Double
    Public P1_Catot_iz As Double

    Public Cafree_jnc As Double            'free Ca in the junctional space
    Public Cafree_iz As Double             'free Ca in the iz space
    Public Cafree_blk As Double            'free Ca in the bulk space

    Public dCatot_jncdt As Double
    Public dCatot_izdt As Double
    Public dCatot_blkdt As Double

    Public ItotNa_cyt As Double

    Public Nai As Double                'no compartmentation
    Public dNaidt As Double
    'Public stdNai As Double = 6.2      '6.2 mM for AP showing Long plateau, 6.4 mM for AP showing shorter plateau 

    Public ItotK_cyt As Double      'no compartmentation

    Public Ki As Double
    Public dKidt As Double

    '*************** [Ca] within SR  ***********************
    Public Ca_SRup As Double
    Public dCa_SRupdt As Double

    Public Ca_SRrl As Double
    Public Catot_SRrl As Double
    Public dCato_SRrldt As Double

    '********** Nernst Potential  **********
    Public ECa_jnc As Double
    Public ECa_iz As Double
    Public ECa_blk As Double
    Public ENa As Double
    Public EK As Double

    '********** Reversal Potential defined in VL analysis  **********
    Public ErevCa_LR As Double               'mV
    Public ErevCa_L0 As Double               'mV

    Public ErevCa_jnc As Double
    Public ErevCa_iz As Double
    Public ErevCa_blk As Double
    Public ErevNa As Double
    Public ErevK As Double

    '********** GHK equation **********
    Public GHKCa_LR As Double             'GHK equation for Ca
    Public GHKCa_L0 As Double              'GHK equation for Ca

    Public GHKCa_jnc As Double             'GHK equation for Ca
    Public GHKCa_iz As Double                  'GHK equation for Ca
    Public GHKCa_blk As Double               'GHK equation for Ca
    Public GHKNa As Double
    Public GHKK As Double
    '   *****************   VL analysis of GHK components ************
    Public dGHKCa_LR As Double             'nS,  slope conductance of GHK equation for Ca   
    Public dGHKCa_L0 As Double               'nS,  slope conductance of GHK equation for Ca

    Public dGHKCa_jnc As Double             'nS,  slope conductance of GHK equation for Ca   
    Public dGHKCa_iz As Double               'nS,  slope conductance of GHK equation for Ca
    Public dGHKCa_blk As Double             'nS, slope conductance of GHK equation for Ca
    Public dGHKNa As Double                    'nS, slope conductance of GHK equation for Na
    Public dGHKK As Double                     'nS, slope conductance of GHK equation for K

    '********** mitochondria   15 variables **********
    Public ATPt_cyt As Double = 6            'mM ATP concentration
    Public ADPt_cyt As Double             'mM  ADP concentration
    Public Pi_cyt As Double                   'mM cytosol Pi concentration
    Public PCr_cyt As Double               'mM PCr concentration
    Public H_cyt As Double = 0.0001             '0038276520463351       'mM cytosol H concentration   Constant at present 

    Public ATPt_mit As Double             'mM  matrix ATP concentration
    Public Pi_mit As Double                  'mM  matrix Pi concentration
    Public NADH_mit As Double           'mM  matrix NADH concentration
    Public H_mit As Double                   'mM  matrix H concentration
    Public K_mit As Double                   'mM matrix K concentration
    Public UQr_mit As Double               'mM matrix UQH concentration
    Public ctCrd_mit As Double              'mM matrix cytcrome concentration
    Public O2_mit As Double = 0.24       'mM O2 concentration   This is constant in the present model.
    Public dpsi As Double                          'mV  matrix membrane potential

    Public dATPt_cytdt As Double             'mM/ms,  rate of cytosol ATP concentration
    Public dADPt_cytdt As Double             'mM/ms,  rate of cytosol ADP concentration
    Public dPi_cytdt As Double                   'mM/ms,  rate of cytosol Pi concentration
    Public dPCr_cytdt As Double               'mM/ms,  rate of cytosol PCr concentration
    'Public dH_cytdt As Double                   'mM/ms,  rate of cytosol H concentration

    Public dATPt_mitdt As Double             'mM/ms,  rate of matrix ATP concentration
    Public dPi_mitdt As Double                  'mM/ms,  rate of matrix Pi concentration
    Public dNADH_mitdt As Double           'mM/ms,  rate of matrix NADH concentration
    Public dH_mitdt As Double                   'mM/ms,  rate of matrix H concentration
    Public dK_mitdt As Double                   'mM/ms,  rate of matrix K concentration
    Public dUQr_mitdt As Double               'mM/ms,  rate of matrix UQH concentration
    Public dctCrd_mitdt As Double              'mM/ms,  rate of matrix cytcrome concentration
    'Public dO2_mitdt As Double                 'mM/ms,  rate of O2 concentration

    Public ddpsidt As Double                       'mV/ms, rate of matrix membrane potential

    '******************************** MODEL CONSTANTS  ********************************
    Public pKa_Pi As Double = 6.8                                                'unitless, pK for Pi      
    Public AXPSUM_mit As Double = 16.26                         'mM     
    Public NADt_mit As Double = 2.79                                  'mM     
    Public UQt_mit As Double = 1.35                                    'mM    
    Public cytCt_mit As Double = 0.27                                  'mM    
    Public cytAt_mit As Double = 0.135                                 'mM    
    Public Mg_mit As Double = 0.38                                      'mM          corrected from 0.3 mM by AN on 24 May
    Public KdADP_mit As Double = 0.282
    Public kdATP_mit As Double = 0.017
    Public EmNAD0_mit As Double = -320                           'mV   
    Public EmUQ0_mit As Double = 40                                 'mV   
    Public EmctC0_mit As Double = 250                             'mV   
    Public EmctA0_mit As Double = 540                              'mV  

    Public nA As Double = 2.5                                                 'unitless    
    Public NADbuf_mit As Double = 5                                   'unitless,   buffering capacity coefficient 

    Public AXPSUM_cyt As Double = 6.7                              'mM    
    Public CrSUM_cyt As Double = 25                                   'mM    
    Public Mg_cyt As Double = 0.8                                             'mM   
    Public KdATP_cyt As Double = 0.024
    Public KdADP_cyt As Double = 0.347

    '********** technical variables **********
    Public fixzero As Double = 1
    Public fixzero0 As Double = 1

    Public Nvar As Double = 71           'number of time-dependent variables  
    Public Mery(Nvar) As Double                   'memory array
    Public dydt(Nvar) As Double             'memory array
    Public Carry(100) As Double             'used to bring data to Form1

    Public EndAddress As Long

    Public isIVcurve As Boolean            'true  ;  specify for calculation of  I-V curve
    Public fixCa As Boolean = False                 'fix [Ca] in all compartments, default = false

    Public Denominator As Double                 'avoid denominator = 0
    Public dt As Double = 0.05                       ' msec, 

    Public NP(20) As Double                           'for the pattern search()
    Public BP1 As Double        'for the csvFileWrite ***** csvFile の最初の１行はIxsfの初期値である
    Public BP2 As Double        'for the csvFileWrite ***** csvFile の最初の１行はIxsfの初期値である
    Public BP3 As Double        'for the csvFileWrite ***** csvFile の最初の１行はIxsfの初期値である
    Public BP4 As Double        'for the csvFileWrite ***** csvFile の最初の１行はIxsfの初期値である
    Public BP5 As Double        'for the csvFileWrite ***** csvFile の最初の１行はIxsfの初期値である
    Public BP6 As Double        'for the csvFileWrite ***** csvFile の最初の１行はIxsfの初期値である
    Public BP7 As Double        'for the csvFileWrite ***** csvFile の最初の１行はIxsfの初期値である

    Public Eval As Double        'for the csvFileWrite ***** csvFile の最初の１行はIxsfの初期値である
    Public CR2 As Double        'countR2,    for the csvFileWrite 

    Public PScycle As Double        'for the csvFileWrite ***** csvFile の最初の１行はIxsfの初期値である
    Public LogMIN As Double = 3        'for the csvFileWrite ***** csvFile の最初の１行はIxsfの初期値である


    Public Sub ModelCell(ByVal sweepTime As Double)

        extConcentrations()                                       'Change in the ext ion concentrations

        substrateConcentrations()                              'substrate concentrations

        CaHomeostasis()                                            'Ion concentrations and  Buffer reactions in compartments  

        CaNanoDomain()                                            'determine [Ca]nd at various combinations

        MembraneCurrentOrg()                         'calculate the ionic currents
        '**********  If constraints = False Then
        '**********    MembraneCurrentOrg()                         'calculate the ionic currents
        '**********  ElseIf constraints = True Then
        '**********   MembraneCurrentSimple()        'calculate using pOen(), which is predetermined by the given time course of Vm
        '***********End If

        SRCadynamics()   'revised in October 2020, so that all Vm dependent rate of LCC is calculated in the new calculateICaLVmgate()

        MembranePotential(sweepTime)                 'calculate the membrane fluxes

        ContractionNL2008(Cafree_blk)                     'Negroni & Lascano 2008

        dIonConcdt()                                                     'rate of change in ion concentrations
        totalCai = Catot_jnc * Vol_jnc + Catot_iz * Vol_iz + Catot_blk * Vol_blk + Ca_SRup * Vol_SRup + Catot_SRrl * Vol_SRrl

        'EnergyMetabolism()                                         'Energy balance
        'EulerMethod(dt)                                          'integrate ODE using Euler Method,  renew the value in y(x)

    End Sub

    Public Sub EulerIntegrationStd(ByVal dt As Double)                                 'Euler method

        ''*****************  Vm  ***********************
        Vm = Vm + dVmdt * dt
        If Vm > -0.0000001 And Vm < 0.0000001 Then Vm = 0.0000001

        '********* Nai, Ki *************************
        Nai = Nai + dNaidt * dt * fixzero0
        Ki = Ki + dKidt * dt * fixzero0

        '********** Catot_blk and Ca-buffers ***************
        Catot_blk = Catot_blk + dCatot_blkdt * dt            'total Ca within blk space

        CaMCa = CaMCa + dCaMCadt * dt  '
        TnChCa = TnChCa + dTnChCadt * dt   '
        SRCa = SRCa + dSRCadt * dt         '

        '********** Catot_iz and Ca-buffers ***************
        Catot_iz = Catot_iz + dCatot_izdt * dt                  'total Ca within iz space
        'LCa_iz = LCa_iz + dLCa_izdt * dt                        'calculate rate of change in LCa_iz
        'HCa_iz = HCa_iz + dHCa_izdt * dt                        'calculate rate of change in LCa_iz

        '********** Catot_jnc and Ca-buffers ***************
        Catot_jnc = Catot_jnc + dCatot_jncdt * dt                  'total Ca within jnc space
        'LCa_jnc = LCa_jnc + dLCa_jncdt * dt                        'calculate rate of change in LCa_iz
        'HCa_jnc = HCa_jnc + dHCa_jncdt * dt                        'calculate rate of change in LCa_iz

        Ca_SRup = Ca_SRup + dCa_SRupdt * dt             'free Ca within SRup
        Catot_SRrl = Catot_SRrl + dCato_SRrldt * dt        'free Ca within SRrl

        '**************  IRyR -Hinch model,   Full number of 8 states model    dyadic space ******************************
        Yooo = Yooo + dYooodt * dt
        Yooc = Yooc + dYoocdt * dt

        Ycoo = Ycoo + dYcoodt * dt
        Ycoc = Ycoc + dYcocdt * dt

        Ycco = Ycco + dYccodt * dt
        'Yccc = Yccc + dYcccdt * dt     'calculate from the mass concervation

        Yoco = Yoco + dYocodt * dt
        Yocc = Yocc + dYoccdt * dt

        '******************  4 state model; ICaL state transition in the iz and blk spaces *************
        us_CaL = us_CaL + dus_CaLdt * dt                    'ultraslow gate of ICaL

        Yco_iz = Yco_iz + dYco_izdt * dt
        Yoc_iz = Yoc_iz + dYoc_izdt * dt
        Yoo_iz = Yoo_iz + dYoo_izdt * dt

        Yco_blk = Yco_blk + dYco_blkdt * dt
        Yoc_blk = Yoc_blk + dYoc_blkdt * dt
        Yoo_blk = Yoo_blk + dYoo_blkdt * dt

        '****************** INa model *******************
        'pO_TM = pO_TM + dO_TMdt * dt                         'gating of INa_TM
        'I2_TM = I2_TM + dI2_TMdt * dt                       'gating of INa_TM
        'Is_TM = Is_TM + dIs_TMdt * dt                         'gating of INa_TM

        'pO_LM = pO_LM + dO_LMdt * dt                                                      'gating of INa_LM
        'I1_LM = I1_LM + dI1_LMdt * dt                                                    'gating of INa_LM
        'I2_LM = I2_LM + dI2_LMdt * dt * fixzero_INaLslowgate             'gating of INa_LM
        'Is_LM = Is_LM + dIs_LMdt * dt * fixzero_INaLslowgate              'gating of INa_LM

        O_TM = O_TM + d12_TM - d23_TM                          'gating of INa_TM
        I2_TM = I2_TM + d23_TM - d31_TM - d34_TM                       'gating of INa_TM
        Is_TM = Is_TM + d34_TM - d41_TM                         'gating of INa_TM

        O_LM = O_LM + d12_LM - d23_LM                                                    'gating of INa_LM
        I1_LM = I1_LM + d23_LM - d31_LM - d34_LM * fixzero_INaLslowgate                                                   'gating of INa_LM
        I2_LM = I2_LM + d34_LM - d41_LM * fixzero_INaLslowgate - d45_LM           'gating of INa_LM
        Is_LM = Is_LM + d45_LM - d51_LM * fixzero_INaLslowgate              'gating of INa_LM

        '********* ICaT   *************
        d_CaT = d_CaT + dd_CaTdt * dt
        f_CaT = f_CaT + df_CaTdt * dt

        '*********** Ist  ************
        d_st = d_st + dd_stdt * dt
        f_st = f_st + df_stdt * dt
        us_st = us_st + dus_stdt * dt

        '********** Iha ************
        C1_ha = C1_ha + dC1_hadt * dt
        C2_ha = C2_ha + dC2_hadt * dt             'dy/dt * dt
        O1_ha = O1_ha + dO1_hadt * dt             'dy/dt * dt
        O2_ha = O2_ha + dO2_hadt * dt             'dy/dt * dt

        '*********** IK1 ***********
        'O_IK1 = O_IK1 + dO_IK1dt * dt
        Pbspm = Pbspm + dPbspmdt * dt

        '************ IKr  ***********
        y1_IKr = y1_IKr + dy1_IKrdt * dt
        y2_IKr = y2_IKr + dy2_IKrdt * dt
        y3_IKr = y3_IKr + dy3_IKrdt * dt

        '************ IKur  ***********
        ua = ua + duadt * dt
        ui = ui + duidt * dt

        '********** IKs  ***********
        Ov_IKs = Ov_IKs + dOv_IKsdt * dt                   'voltage-gate

        C2IKs_iz = C2IKs_iz + dC2IKs_izdt * dt            'nrs compartment, Calcium gate with C1 - C2 - Oc states
        OcIKs_iz = OcIKs_iz + dOcIKs_izdt * dt            'nrs compartment, Calcium gate with C1 - C2 - Oc states

        C2IKs_blk = C2IKs_blk + dC2IKs_blkdt * dt      'blk compartment, Calcium gate with C1 - C2 - Oc states
        OcIKs_blk = OcIKs_blk + dOcIKs_blkdt * dt      'blk compartment, Calcium gate with C1 - C2 - Oc states

        '*********** Kto  *********
        y1_IKto = y1_IKto + dy1_IKtodt * dt
        y2_IKto = y2_IKto + dy2_IKtodt * dt

        '********** IKACh ******
        O_ACh = O_ACh + dO_AChdt * dt

        '****NCX Na/Ca exchange, and Na/K pump ****
        E1NCX_iz = E1NCX_iz + dE1NCX_izdt * dt
        I1NCX_iz = I1NCX_iz + dI1NCX_izdt * dt
        I2NCX_iz = I2NCX_iz + dI2NCX_izdt * dt

        E1NCX_blk = E1NCX_blk + dE1NCX_blkdt * dt
        I1NCX_blk = I1NCX_blk + dI1NCX_blkdt * dt
        I2NCX_blk = I2NCX_blk + dI2NCX_blkdt * dt

        '****************** NaK pump  *************************************************
        P1_6_NaK = P1_6_NaK + dP1_6_NaKdt * dt
        P7_NaK = P7_NaK + dP7_NaKdt * dt
        P8_13_NaK = P8_13_NaK + dP8_13_NaKdt * dt

        ' ******** NL model of Contraction  *************
        TSCa3 = TSCa3 + d12 - d23
        TSCa3W = TSCa3W + d23 - d34
        TSCa3S = TSCa3S + d34 - d45
        TSS = TSS + d45 - d56
        TSW = TSW + d56 - d61

        hw = hw + dhwdt * dt                   'change in hw and hs

        hp = hp + dhpdt * dt                     'change in hw and hs
        Dim hSL As Double = halfSL       'dammy to be remove

    End Sub


    Public DA(20, 43) As Double         'for VL analysis

    Public VL As Double                   'mV, Lead potential (n)
    Public oldVL As Double                   'mV, Lead potential (n-1)
    Public dVLdt As Double                    'dVL/dt

    Dim Gtot As Double                          ' nS,  Whole cell conductance for all cations      Gtot(n+1)   
    Dim dV As Double = 0.2                    'mV, half interval to measure the slope conductance

    Public sumGx As Double                  'total G
    Public SCtbdVLdt As Double               'summator of contributions of individual current components to dVL/dt
    Public SCtbVL As Double                     'summator of contributions of individual current components to VL
    Dim Ctb As Double
    Dim CtbEx As Double                        'contribution attributable to changes in Ex
    Public absVL As Double
    Public absdVLdt As Double
    Dim numX As Double = 42                    'number of VL components
    Public INa1 As Double
    Public Ist1 As Double
    Public Iha1 As Double
    Public IcaT1 As Double
    Public ICaL1 As Double
    Public IbCa1 As Double
    Public IK11 As Double
    Public IKr1 As Double
    Public IKs1 As Double
    Public Ito1 As Double
    Public IKur1 As Double
    Public IKAch1 As Double
    Public IKpl1 As Double
    Public IbNSC1 As Double
    Public ILCCa1 As Double
    Public IKATP1 As Double
    Public INaK1 As Double
    Public INCX1 As Double

    Public INa2 As Double
    Public Ist2 As Double
    Public Iha2 As Double
    Public IcaT2 As Double
    Public ICaL2 As Double
    Public IbCa2 As Double
    Public IK12 As Double
    Public IKr2 As Double
    Public IKs2 As Double
    Public Ito2 As Double
    Public IKur2 As Double
    Public IKAch2 As Double
    Public IKpl2 As Double
    Public IbNSC2 As Double
    Public ILCCa2 As Double
    Public IKATP2 As Double
    Public INaK2 As Double
    Public INCX2 As Double

    Public s1 As Double
    Public s2 As Double
    Public s3 As Double
    Public s4 As Double
    Public s5 As Double
    Public s6 As Double
    Public s7 As Double
    Public s8 As Double
    Public s9 As Double
    Public s10 As Double
    Public s11 As Double
    Public s12 As Double
    Public s13 As Double
    Public s14 As Double
    Public s15 As Double
    Public s16 As Double
    Public s17 As Double
    Public s18 As Double
    Public s19 As Double
    Public s20 As Double
    Public s21 As Double
    Public s22 As Double
    Public s23 As Double
    Public s24 As Double
    Public s25 As Double
    Public s26 As Double
    Public s27 As Double
    Public s28 As Double
    Public s29 As Double
    Public s30 As Double
    Public s31 As Double
    Public s32 As Double
    Public s33 As Double
    Public s34 As Double
    Public s35 As Double
    Public s36 As Double
    Public s37 As Double
    Public s38 As Double
    Public s39 As Double
    Public s40 As Double
    Public s41 As Double
    Public s42 As Double
    Public sall As Double

    Public c1 As Double
    Public c2 As Double
    Public c3 As Double
    Public c4 As Double
    Public c5 As Double
    Public c6 As Double
    Public c7 As Double
    Public c8 As Double
    Public c9 As Double
    Public c10 As Double
    Public c11 As Double
    Public c12 As Double
    Public c13 As Double
    Public c14 As Double
    Public c15 As Double
    Public c16 As Double
    Public c17 As Double
    Public c18 As Double
    Public c19 As Double
    Public c20 As Double
    Public c21 As Double
    Public c22 As Double
    Public c23 As Double
    Public c24 As Double
    Public c25 As Double
    Public c26 As Double
    Public c27 As Double
    Public c28 As Double
    Public c29 As Double
    Public c30 As Double
    Public c31 As Double
    Public c32 As Double
    Public c33 As Double
    Public c34 As Double
    Public c35 As Double
    Public c36 As Double
    Public c37 As Double
    Public c38 As Double
    Public c39 As Double
    Public c40 As Double
    Public c41 As Double
    Public c42 As Double
    Public csall As Double

    Public Sub calculateLeadP()     'Lead potential analysis based on the linearlized whole cell ion conductances
        Dim I As Integer
        For I = 1 To numX
            DA(3, I) = DA(2, I)                  'copy the Gion(n) as Gion(n-1)
            DA(5, I) = DA(4, I)                  'copy the Ex(n) as Ex(n-1)
        Next
        'DA(3, 35) = DA(1, 35)                 'copy the IPMCA_iz(n) as IPMCA(n-1)
        'DA(3, 36) = DA(1, 36)                 'copy the IPMCA_blk as IPMCA(n-1)

        oldVL = VL                      'copy Lead Potential to obtain dVL/dt directly if necessary

        'FillDarray()                                 'fill the VL sheet (DAR(1,I)) including IPMCA  

        '*********** Calculate Gx, VL,  and finally VL  *****************
        sumGx = 0                                                 'prepare the summator of Gx
        Dim sumGxEx As Double = 0                     'prepare the summator of GxEx

        For I = 1 To numX                                            'Firstly, determine the sumGx of individual ionic current
            sumGx += DA(2, I)                                'Sum of Gx
        Next I

        For I = 1 To numX                                                      'contribution to VL
            DA(7, I) = DA(2, I) * DA(4, I) / sumGx          'Gx*Ex / sumGx      
            sumGxEx += DA(7, I)                                      'Sum of contributions of ion channels to VL
        Next I
        '******************** ion exchangers *********************
        'DA(7, 35) = -DA(1, 35) / sumGx
        'sumGxEx += DA(7, 35)                            'Sum of contributions of PMCA to VL
        'DA(7, 36) = -DA(1, 36) / sumGx
        'sumGxEx += DA(7, 36)                            'Sum of contributions of PMCA to VL

        VL = sumGxEx                                    'Lead Potential

        '***********      contributions of individual current component to dVL/dt (mV/msec) ; Rate of VL change *********
        SCtbdVLdt = 0                        'summator of dVL/dt
        For I = 1 To numX                         'contribution of Ion channels to VL based on the analytical solution dVL/dt
            DA(6, I) = ((DA(2, I) - DA(3, I)) * (DA(4, I) - VL) + DA(2, I) * (DA(4, I) - DA(5, I))) / sumGx / dt  'mV/ms
            SCtbdVLdt += DA(6, I)   'mV/ms
            'DA(9, I) = (DA(2, I) * (DA(4, I) - DA(5, I))) / sumGx / dt      'contribution attributed to a change in Ex 
            'DA(8, I) = DA(9, I) / Ctb
        Next I

        'DA(6, 35) = -(DA(1, 35) - DA(3, 35)) / sumGx / dt      'IPMCA_iz
        'SCtbdVLdt += DA(6, 35)
        'DA(6, 36) = -(DA(1, 36) - DA(3, 36)) / sumGx / dt      'IPMCA_blk
        'SCtbdVLdt += DA(6, 36)

        dVLdt = (VL - oldVL) / dt       'ms/mV    direct calculation of dVL/dt

        '******* summate values of elements for individual currents **************** 
        DA(11, 2) = DA(7, 1) + DA(7, 2) + DA(7, 3) + DA(7, 4)                          'INa elements of VL
        DA(11, 4) = DA(7, 5) + DA(7, 6)                         'Ist elements of VL
        DA(11, 6) = DA(7, 7) + DA(7, 8)                          'Iha elements of VL
        DA(11, 7) = DA(7, 9)                                          'ICaT element of VL
        DA(11, 17) = DA(7, 10) + DA(7, 11) + DA(7, 12) + DA(7, 13) + DA(7, 14) + DA(7, 15) + DA(7, 16) + DA(7, 17) + DA(7, 18) + DA(7, 19)   'ICaL elements of VL
        DA(11, 19) = DA(7, 20) + DA(7, 21)                          'IbCa elements of VL
        DA(11, 20) = DA(7, 22)                                          'IK1 element of VL
        DA(11, 21) = DA(7, 23)                                          'IKr element of VL
        DA(11, 25) = DA(7, 24) + DA(7, 25) + DA(7, 26) + DA(7, 27)     'IKr elements of VL
        DA(11, 26) = DA(7, 28)                                          'IKACh element of VL
        DA(11, 27) = DA(7, 29)                                          'IKpl element of VL
        DA(11, 29) = DA(7, 30) + DA(7, 31)                          'ItoK elements of VL
        DA(11, 31) = DA(7, 32) + DA(7, 33)                          'IbNSC elements of VL
        DA(11, 35) = DA(7, 34) + DA(7, 35) + DA(7, 36) + DA(7, 37)     'ILCCa elements of VL
        DA(11, 36) = DA(7, 38)                                          'IKATP element of VL
        DA(11, 38) = DA(7, 39) + DA(7, 40)                          'INaCa elements of VL
        DA(11, 39) = DA(7, 41)                                          'IKATP element of VL
        DA(11, 40) = DA(7, 42)                                          'IKur element of VL

        DA(12, 2) = DA(6, 1) + DA(6, 2) + DA(6, 3) + DA(6, 4)                           'INa elements of dVL/dt
        DA(12, 4) = DA(6, 5) + DA(6, 6)                          'Ist elements of dVL/dt
        DA(12, 6) = DA(6, 7) + DA(6, 8)                           'Iha elements of dVL/dt
        DA(12, 7) = DA(6, 9)                                          'ICaT element of dVL/dt
        DA(12, 17) = DA(6, 10) + DA(6, 11) + DA(6, 12) + DA(6, 13) + DA(6, 14) + DA(6, 15) + DA(6, 16) + DA(6, 17) + DA(6, 18) + DA(6, 19)   'ICaL elements of dVL/dt
        DA(12, 19) = DA(6, 20) + DA(6, 21)                           'IbCa elements of dVL/dt
        DA(12, 20) = DA(6, 22)                                          'IK1 element of dVL/dt
        DA(12, 21) = DA(6, 23)                                          'IKr element of dVL/dt
        DA(12, 25) = DA(6, 24) + DA(6, 25) + DA(6, 26) + DA(6, 27)     'IKr elements of dVL/dt
        DA(12, 26) = DA(6, 28)                                          'IKACh element of dVL/dt
        DA(12, 27) = DA(6, 29)                                          'IKpl element of dVL/dt
        DA(12, 29) = DA(6, 30) + DA(6, 31)                           'ItoK elements of dVL/dt
        DA(12, 31) = DA(6, 32) + DA(6, 33)                           'IbNSC elements of dVL/dt
        DA(12, 35) = DA(6, 34) + DA(6, 35) + DA(6, 36) + DA(6, 37)     'ILCCa elements of dVL/dt
        DA(12, 36) = DA(6, 38)                                          'IKATP element of dVL/dt
        DA(12, 38) = DA(6, 39) + DA(6, 40)                           'INaCa elements of dVL/dt
        DA(12, 39) = DA(6, 41)                                          'IKATP element of dVL/dt
        DA(12, 40) = DA(6, 42)                                          'IKur element of dVL/dt

        DA(14, 2) = Math.Abs(DA(11, 2))
        DA(14, 4) = Math.Abs(DA(11, 4))
        DA(14, 6) = Math.Abs(DA(11, 6))
        DA(14, 7) = Math.Abs(DA(11, 7))
        DA(14, 17) = Math.Abs(DA(11, 17))
        DA(14, 19) = Math.Abs(DA(11, 19))
        DA(14, 20) = Math.Abs(DA(11, 20))
        DA(14, 21) = Math.Abs(DA(11, 21))
        DA(14, 25) = Math.Abs(DA(11, 25))
        DA(14, 26) = Math.Abs(DA(11, 26))
        DA(14, 27) = Math.Abs(DA(11, 27))
        DA(14, 29) = Math.Abs(DA(11, 29))
        DA(14, 31) = Math.Abs(DA(11, 31))
        DA(14, 35) = Math.Abs(DA(11, 35))
        DA(14, 36) = Math.Abs(DA(11, 36))
        DA(14, 38) = Math.Abs(DA(11, 38))
        DA(14, 39) = Math.Abs(DA(11, 39))
        DA(14, 40) = Math.Abs(DA(11, 40))
        absVL = DA(14, 2) + DA(14, 4) + DA(14, 6) + DA(14, 7) + DA(14, 17) + DA(14, 19) + DA(14, 20) + DA(14, 21) + DA(14, 25) + DA(14, 26) + DA(14, 27) + DA(14, 29) + DA(14, 31) + DA(14, 35) + DA(14, 36) + DA(14, 38) + DA(14, 39) + DA(14, 40)

        DA(15, 2) = Math.Abs(DA(12, 2))
        DA(15, 4) = Math.Abs(DA(12, 4))
        DA(15, 6) = Math.Abs(DA(12, 6))
        DA(15, 7) = Math.Abs(DA(12, 7))
        DA(15, 17) = Math.Abs(DA(12, 17))
        DA(15, 19) = Math.Abs(DA(12, 19))
        DA(15, 20) = Math.Abs(DA(12, 20))
        DA(15, 21) = Math.Abs(DA(12, 21))
        DA(15, 25) = Math.Abs(DA(12, 25))
        DA(15, 26) = Math.Abs(DA(12, 26))
        DA(15, 27) = Math.Abs(DA(12, 27))
        DA(15, 29) = Math.Abs(DA(12, 29))
        DA(15, 31) = Math.Abs(DA(12, 31))
        DA(15, 35) = Math.Abs(DA(12, 35))
        DA(15, 36) = Math.Abs(DA(12, 36))
        DA(15, 38) = Math.Abs(DA(12, 38))
        DA(15, 39) = Math.Abs(DA(12, 39))
        DA(15, 40) = Math.Abs(DA(12, 40))
        absdVLdt = DA(15, 2) + DA(15, 4) + DA(15, 6) + DA(15, 7) + DA(15, 17) + DA(15, 19) + DA(15, 20) + DA(15, 21) + DA(15, 25) + DA(15, 26) + DA(15, 27) + DA(15, 29) + DA(15, 31) + DA(15, 35) + DA(15, 36) + DA(15, 38) + DA(15, 39) + DA(15, 40)

        INa1 = DA(11, 2)                    'INa elements of VL
        Ist1 = DA(11, 4)                        'Ist elements of VL
        Iha1 = DA(11, 6)                  'Iha elements of VL
        IcaT1 = DA(11, 7)                                       'ICaT element of VL
        ICaL1 = DA(11, 17) 'ICaL elements of VL
        IbCa1 = DA(11, 19)                    'IbCa elements of VL
        IK11 = DA(11, 20)                                       'IK1 element of VL
        IKr1 = DA(11, 21)                                    'IKr element of VL
        IKs1 = DA(11, 25)     'IKr elements of VL
        IKAch1 = DA(11, 26)                                        'IKACh element of VL
        IKpl1 = DA(11, 27)                                        'IKpl element of VL
        Ito1 = DA(11, 29)                           'ItoK elements of VL
        IbNSC1 = DA(11, 31)                           'IbNSC elements of VL
        ILCCa1 = DA(11, 35)     'ILCCa elements of VL
        IKATP1 = DA(11, 36)                                      'IKATP element of VL
        INCX1 = DA(11, 38)                           'INaCa elements of VL
        INaK1 = DA(11, 39)                                         'IKATP element of VL
        IKur1 = DA(11, 40)


        s1 = (DA(2, 1))
        s2 = (DA(2, 2))
        s3 = (DA(2, 3))
        s4 = (DA(2, 4))
        s5 = (DA(2, 5))
        s6 = (DA(2, 6))
        s7 = (DA(2, 7))
        s8 = (DA(2, 8))
        s9 = (DA(2, 9))
        s10 = (DA(2, 10))
        s11 = (DA(2, 11))
        s12 = (DA(2, 12))
        s13 = (DA(2, 13))
        s14 = (DA(2, 14))
        s15 = (DA(2, 15))
        s16 = (DA(2, 16))
        s17 = (DA(2, 17))
        s18 = (DA(2, 18))
        s19 = (DA(2, 19))
        s20 = (DA(2, 20))
        s21 = (DA(2, 21))
        s22 = (DA(2, 22))
        s23 = (DA(2, 23))
        s24 = (DA(2, 24))
        s25 = (DA(2, 25))
        s26 = (DA(2, 26))
        s27 = (DA(2, 27))
        s28 = (DA(2, 28))
        s29 = (DA(2, 29))
        s30 = (DA(2, 30))
        s31 = (DA(2, 31))
        s32 = (DA(2, 32))
        s33 = (DA(2, 33))
        s34 = (DA(2, 34))
        s35 = (DA(2, 35))
        s36 = (DA(2, 36))
        s37 = (DA(2, 37))
        s38 = (DA(2, 38))
        s39 = (DA(2, 39))
        s40 = (DA(2, 40))
        s41 = (DA(2, 41))
        s42 = (DA(2, 42))
        sall = sumGx

        c1 = (DA(2, 1)) * (DA(4, 1))
        c2 = (DA(2, 2)) * (DA(4, 2))
        c3 = (DA(2, 3)) * (DA(4, 3))
        c4 = (DA(2, 4)) * (DA(4, 4))
        c5 = (DA(2, 5)) * (DA(4, 5))
        c6 = (DA(2, 6)) * (DA(4, 6))
        c7 = (DA(2, 7)) * (DA(4, 7))
        c8 = (DA(2, 8)) * (DA(4, 8))
        c9 = (DA(2, 9)) * (DA(4, 9))
        c10 = (DA(2, 10)) * (DA(4, 10))
        c11 = (DA(2, 11)) * (DA(4, 11))
        c12 = (DA(2, 12)) * (DA(4, 12))
        c13 = (DA(2, 13)) * (DA(4, 13))
        c14 = (DA(2, 14)) * (DA(4, 14))
        c15 = (DA(2, 15)) * (DA(4, 15))
        c16 = (DA(2, 16)) * (DA(4, 16))
        c17 = (DA(2, 17)) * (DA(4, 17))
        c18 = (DA(2, 18)) * (DA(4, 18))
        c19 = (DA(2, 19)) * (DA(4, 19))
        c20 = (DA(2, 20)) * (DA(4, 20))
        c21 = (DA(2, 21)) * (DA(4, 21))
        c22 = (DA(2, 22)) * (DA(4, 22))
        c23 = (DA(2, 23)) * (DA(4, 23))
        c24 = (DA(2, 24)) * (DA(4, 24))
        c25 = (DA(2, 25)) * (DA(4, 25))
        c26 = (DA(2, 26)) * (DA(4, 26))
        c27 = (DA(2, 27)) * (DA(4, 27))
        c28 = (DA(2, 28)) * (DA(4, 28))
        c29 = (DA(2, 29)) * (DA(4, 29))
        c30 = (DA(2, 30)) * (DA(4, 30))
        c31 = (DA(2, 31)) * (DA(4, 31))
        c32 = (DA(2, 32)) * (DA(4, 32))
        c33 = (DA(2, 33)) * (DA(4, 33))
        c34 = (DA(2, 34)) * (DA(4, 34))
        c35 = (DA(2, 35)) * (DA(4, 35))
        c36 = (DA(2, 36)) * (DA(4, 36))
        c37 = (DA(2, 37)) * (DA(4, 37))
        c38 = (DA(2, 38)) * (DA(4, 38))
        c39 = (DA(2, 39)) * (DA(4, 39))
        c40 = (DA(2, 40)) * (DA(4, 40))
        c41 = (DA(2, 41)) * (DA(4, 41))
        c42 = (DA(2, 42)) * (DA(4, 42))
        csall = sumGxEx

    End Sub

    '********** Injection current **********
    Public clampMode As String = "Iclamp"         'always Current Clamp Mode            'decide if V clamp or I clamp

    Public Iapl_blk As Double = 0                'stimulus current applied from external device, assumed to be carried by K+
    Public Samplitude As Double = 0         'pA     current amplitude 
    Public ItestOn As Double = 25000            'start time of external stimulation
    Public ItestOff As Double = 25500             'end time of external stimulation
    Public SInterval As Double               'ms,   stimulus interval
    Public VtestInterval As Double = 5        'mV    interval of voltage clamp test pulse

    Public Sub Iclampsub(ByRef t As Double)
        Iapl_blk = 0
        If t >= ItestOn And t < ItestOff Then
            Iapl_blk = Samplitude
        ElseIf t >= (ItestOn + 15000) And t < (ItestOff + 15000) Then
            Iapl_blk = Samplitude
        Else
            Iapl_blk = 0
        End If
    End Sub

    Public Vhold As Double = -50
    Public Vtest As Double
    Public VtestOn As Double = 50 '= 10
    Public VtestOff As Double = 450

    Public Sub Vclampsub(ByRef t As Double, ByVal dt As Double, ByVal vtest As Double)
        Dim FBgain As Double = 1 / (100 * dt)     'feedback gain is determined by the (x*dt)
        If t >= VtestOn And t < VtestOff Then
            Iapl_blk = -Itot_cell - (vtest - Vm) * FBgain
        Else
            Iapl_blk = -Itot_cell - (Vhold - Vm) * FBgain
        End If

    End Sub

    Public Sub MembraneCurrentOrg()
        NernstGHK()

        calculateINarate()
        calculateINamag(GHKNa, GHKK, INaT_Na_cyt, INaT_K_cyt, INaL_Na_cyt, INaL_K_cyt)

        INaT = INaT_Na_cyt + INaT_K_cyt
        INaL = INaL_Na_cyt + INaL_K_cyt
        INa = INaT + INaL
        'calculateINa(Vm, GHKNa, GHKK, INa, INa_Na, INa_K)   'calculate INa


        '(1)'******    LCC    ********  Vm-dependent kinetics ******4 状態モデル
        calculateICaLslowGate()                           'calculate the ulstraslow inactivation

        calculateICaLVmgate(kcoVm, kocVm)          'Vm-dependent gate  for both 4 state and 8 state model (RyR)

        calculateICaLEta(kccco, kocoo)                     'Vm-dependent gate  kocoo, kccco,     

        '(2)********** LCC *************** Ca-dependent kinetics *******　4 状態モデル

        'calculateICaL4stateCaGate(Cafree_iz, Ycc_iz, Yco_iz, Yoo_iz, Yoc_iz, kcocc_iz, koooc_iz, dYco_izdt, dYoo_izdt, dYoc_izdt)
        calculateICaLCagate(Ca_loc_iz, koooc_iz)      'forward rate, when LCC is open
        calculateICaLCagate(Cafree_iz, kcocc_iz)      'forward rate, when LCC is still not open

        'calculateICaL4stateCaGate(Cafree_blk, Ycc_blk, Yco_blk, Yoo_blk, Yoc_blk, kcocc_blk, koooc_blk, dYco_blkdt, dYoo_blkdt, dYoc_blkdt)
        calculateICaLCagate(Ca_loc_blk, koooc_blk)      'forward rate, when LCC is open
        calculateICaLCagate(Cafree_blk, kcocc_blk)      'forward rate, when LCC is still not open

        izblkLCC_dydt()                'calculate both iz- and blk-spaces


        calculateICaL4stateAmp(FrcCaL_iz, Yoo_iz, GHKCa_iz, GHKNa, GHKK, ICaLCa_iz, ICaLNa_iz, ICaLK_iz)
        calculateICaL4stateAmp(FrcCaL_blk, Yoo_blk, GHKCa_blk, GHKNa, GHKK, ICaLCa_blk, ICaLNa_blk, ICaLK_blk)

        ICaL = ICaLCa_LR_jnc + ICaLCa_L0_jnc + ICaLNa_jnc + ICaLK_jnc + ICaLCa_iz + ICaLNa_iz + ICaLK_iz + ICaLCa_blk + ICaLNa_blk + ICaLK_blk
        ICaL_jnc = ICaLCa_LR_jnc + ICaLCa_L0_jnc + ICaLNa_jnc + ICaLK_jnc
        ICaL_iz = ICaLCa_iz + ICaLNa_iz + ICaLK_iz
        ICaL_blk = ICaLCa_blk + ICaLNa_blk + ICaLK_blk

        ''******************* ICaT ****************************************
        calculateICaT(GHKCa_blk, ICaT_blk)            'calculate ICaT
        ICaT = ICaT_blk

        calculateIst(Cafree_blk)           'calculate Ist

        calculateIha()           'calculate Iha

        calculateIK1(EK, Pbspm, dPbspmdt)
        IK1 = IK1_K_cyt

        calculateIKur()

        calculateIKr()
        IKr = IKr_K_cyt

        VgateIKs(Ov_IKs, dOv_IKsdt)           ''IKs voltage gate
        calculateIKs(Cafree_iz, Frc_iz, Ov_IKs, C2IKs_iz, OcIKs_iz, dOcIKs_izdt, dC2IKs_izdt, pO_IKs_iz)
        IKs_K_iz = IKs_K                                          'define IKs_K_iz
        IKs_Na_iz = IKs_Na                                      'define IKs_Na_iz
        IKs_iz = (IKs_K_iz + IKs_Na_iz)

        calculateIKs(Cafree_blk, Frc_blk, Ov_IKs, C2IKs_blk, OcIKs_blk, dOcIKs_blkdt, dC2IKs_blkdt, pO_IKs_blk)
        IKs_K_blk = IKs_K                                              'define IKs_K_blk
        IKs_Na_blk = IKs_Na                                          'define IKs_Na_blk
        IKs_blk = (IKs_K_blk + IKs_Na_blk)

        IKs = IKs_iz + IKs_blk

        calculateIKtoEndo(IKto_K_cyt, IKto_Na_cyt, y1_IKto, y2_IKto, dy1_IKtodt, dy2_IKtodt)
        IKto = IKto_K_cyt + IKto_Na_cyt

        calculateIKACh()                        'calculate IKACH
        calculateIKpl()                      ' IK plateau
        calculateIbNSC(GHKNa, GHKK)         'non-selective cation current

        calculateIbCa(GHKCa_blk, IbCa_blk)  'background Ca current in blk-space
        calculateIbCa(GHKCa_iz, IbCa_iz)      'background Ca current in iz-space
        IbCa = IbCa_blk + IbCa_iz                   'sum of the two background Ca current 


        calculateIL_Ca(Frc_iz, Cafree_iz, ILCCa_Na_iz, ILCCa_K_iz, ILCCa_iz)             'Ca-dependent non-selective cation current in iz-space
        calculateIL_Ca(Frc_blk, Cafree_blk, ILCCa_Na_blk, ILCCa_K_blk, ILCCa_blk)  'Ca-dependent non-selective cation current in blk-space
        ILCCa = ILCCa_Na_iz + ILCCa_K_iz + ILCCa_Na_blk + ILCCa_K_blk                         'sum of IL_Ca for calculation of dVmdt
        ILCCa_iz = ILCCa_Na_iz + ILCCa_K_iz
        ILCCa_blk = ILCCa_Na_blk + ILCCa_K_blk

        calculateIKATP()                       'calculate IKATP

        calculateNaK(Nai, Ki, MgATP_cyt, MgADP_cyt, INaK, INaK_Na_cyt, INaK_K_cyt, dATP_NaK, P1_6_NaK, P7_NaK, P8_13_NaK, dP1_6_NaKdt, dP7_NaKdt, dP8_13_NaKdt)

        calculateNCX(FrcNCX_iz, Nai, Cafree_iz, INCX_iz, INCXNa_iz, INCXCa_iz, E1NCX_iz, I1NCX_iz, I2NCX_iz, dE1NCX_izdt, dI1NCX_izdt, dI2NCX_izdt)
        INCXNa_iz = 3 * INCX_iz
        INCXCa_iz = -2 * INCX_iz

        calculateNCX(FrcNCX_blk, Nai, Cafree_blk, INCX_blk, INCXNa_blk, INCXCa_blk, E1NCX_blk, I1NCX_blk, I2NCX_blk, dE1NCX_blkdt, dI1NCX_blkdt, dI2NCX_blkdt)           'calculate  INaCa
        INCXNa_blk = 3 * INCX_blk
        INCXCa_blk = -2 * INCX_blk

        INCX = INCXNa_iz + INCXCa_iz + INCXNa_blk + INCXCa_blk


    End Sub

    'Public Sub MembraneCurrentSimple()
    '    NernstGHK()
    '    '*********** calculateINarate() ***************
    '    d12_TM = 0.0
    '    d23_TM = 0.0
    '    d31_TM = 0.0
    '    d34_TM = 0.0
    '    d41_TM = 0.0

    '    d12_LM = 0.0
    '    d23_LM = 0.0
    '    d31_LM = 0.0
    '    d34_LM = 0.0
    '    d41_LM = 0.0
    '    d45_LM = 0.0
    '    d51_LM = 0.0

    '    calculateINamag(GHKNa, GHKK, INaT_Na_cyt, INaT_K_cyt, INaL_Na_cyt, INaL_K_cyt)


    '    INaT = INaT_Na_cyt + INaT_K_cyt
    '    INaL = INaL_Na_cyt + INaL_K_cyt
    '    INa = INaT + INaL

    '    ''******    ICaL    ********  ICaL *****************************
    '    dus_CaLdt = 0.0
    '    'calculateICaLslowGate()                           'calculate the ulstraslow inactivation

    '    'calculateICaLVmgateSimple()          'Vm-dependent gate  for both 4 state and 8 state model (RyR)
    '    'calculateICaLVmgate(kcoVm, kocVm)           'Vm-dependent gate  for both 4 state and 8 state model (RyR)
    '    '                 kcooo, kooco, kccoc, koccc, kcooo,       kYcooYooo, kYoooYcoo, kYcocYooc, kYoocYcoc, YccoYoco, kYocoYcco, kYcccYocc, kYoccYccc

    '    '************************** LCC  Ca gate ***********************************
    '    calculateICaLCagate(Ca_loc_iz, koooc_iz)      'forward rate, when LCC is open
    '    calculateICaLCagate(Cafree_iz, kcocc_iz)      'forward rate, when LCC is still not open

    '    calculateICaLCagate(Ca_loc_blk, koooc_blk)      'forward rate, when LCC is open
    '    calculateICaLCagate(Cafree_blk, kcocc_blk)      'forward rate, when LCC is still not open

    '    'calculateICaLEtaSimple()          'Vm-dependent gate  kocoo, kccco,       kYcccYcoc, kYoccYooc, kYccoYcoo, kYocoYooo
    '    'calculateICaLEta(kccco, kocoo)  'Vm-dependent gate  in both membrane kocoo, kccco, and CaRU   kYcccYcoc, kYoccYooc, kYccoYcoo, kYocoYooo

    '    izblkLCC_dydt()                'calculate ICaL-gate in both iz- and blk-spaces

    '    '************************** LCC  ICaL amplitude  ***********************************
    '    calculateICaL4stateAmp(FrcCaL_iz, Yoo_iz, GHKCa_iz, GHKNa, GHKK, ICaLCa_iz, ICaLNa_iz, ICaLK_iz)
    '    calculateICaL4stateAmp(FrcCaL_blk, Yoo_blk, GHKCa_blk, GHKNa, GHKK, ICaLCa_blk, ICaLNa_blk, ICaLK_blk)   'in the blk-space

    '    ICaL_jnc = ICaLCa_LR_jnc + ICaLCa_L0_jnc + ICaLNa_jnc + ICaLK_jnc
    '    ICaL_iz = ICaLCa_iz + ICaLNa_iz + ICaLK_iz
    '    ICaL_blk = ICaLCa_blk + ICaLNa_blk + ICaLK_blk

    '    ICaL = ICaLCa_LR_jnc + ICaLCa_L0_jnc + ICaLNa_jnc + ICaLK_jnc + ICaLCa_iz + ICaLNa_iz + ICaLK_iz + ICaLCa_blk + ICaLNa_blk + ICaLK_blk
    '    ''******************************************************************************************************

    '    calculateICaTSimple()            'calculate ICaT
    '    ICaT = ICaT_blk

    '    calculateIstSimple(tgVm)           'calculate Ist
    '    'calculateIst(Cafree_blk)           'calculate Ist

    '    calculateIhasimple()                     'simplified

    '    calculateIKrSimple()                      'simplified
    '    IKr = IKr_K_cyt

    '    calculateIK1Simple()                       'simplified
    '    IK1 = IK1_K_cyt

    '    calculateIKurSimple()                      'simplified

    '    calculateIKtoSimple()                      'simplified
    '    IKto = IKto_K_cyt + IKto_Na_cyt

    '    '*********** Ca dependent currents ************************************
    '    'VgateIKs(Vm, Ov_IKs, dOv_IKsdt)     ''IKs     simplified
    '    calculateIKsSimple(Cafree_iz, Frc_iz, Ov_IKs, C2IKs_iz, OcIKs_iz, dOcIKs_izdt, dC2IKs_izdt, pO_IKs_iz)
    '    IKs_K_iz = IKs_K                                          'define IKs_K_iz
    '    IKs_Na_iz = IKs_Na                                      'define IKs_Na_iz
    '    IKs_iz = (IKs_K_iz + IKs_Na_iz)

    '    calculateIKsSimple(Cafree_blk, Frc_blk, Ov_IKs, C2IKs_blk, OcIKs_blk, dOcIKs_blkdt, dC2IKs_blkdt, pO_IKs_blk)
    '    IKs_K_blk = IKs_K                                              'define IKs_K_blk
    '    IKs_Na_blk = IKs_Na                                          'define IKs_Na_blk
    '    IKs_blk = (IKs_K_blk + IKs_Na_blk)

    '    IKs = IKs_iz + IKs_blk

    '    '************** background currents *****************************************
    '    calculateIbNSC(GHKNa, GHKK)         'non-selective cation current

    '    calculateIKACh()                        'calculate IKACH
    '    calculateIKpl()                      ' IK plateau

    '    calculateIbCa(GHKCa_blk, IbCa_blk)  'background Ca current in blk-space
    '    calculateIbCa(GHKCa_iz, IbCa_iz)      'background Ca current in iz-space
    '    IbCa = IbCa_blk + IbCa_iz                   'sum of the two background Ca current 

    '    calculateIL_Ca(Frc_iz, Cafree_iz, ILCCa_Na_iz, ILCCa_K_iz, ILCCa_iz)             'Ca-dependent non-selective cation current in iz-space
    '    calculateIL_Ca(Frc_blk, Cafree_blk, ILCCa_Na_blk, ILCCa_K_blk, ILCCa_blk)  'Ca-dependent non-selective cation current in blk-space
    '    ILCCa = ILCCa_Na_iz + ILCCa_K_iz + ILCCa_Na_blk + ILCCa_K_blk                         'sum of IL_Ca for calculation of dVmdt
    '    ILCCa_iz = ILCCa_Na_iz + ILCCa_K_iz
    '    ILCCa_blk = ILCCa_Na_blk + ILCCa_K_blk

    '    calculateIKATP()                       'calculate IKATP

    '    '**************** ion transporters ************************************************
    '    calculateNaK(Nai, Ki, MgATP_cyt, MgADP_cyt, INaK, INaK_Na_cyt, INaK_K_cyt, dATP_NaK, P1_6_NaK, P7_NaK, P8_13_NaK, dP1_6_NaKdt, dP7_NaKdt, dP8_13_NaKdt)

    '    calculateNCX(FrcNCX_iz, Nai, Cafree_iz, INCX_iz, INCXNa_iz, INCXCa_iz, E1NCX_iz, I1NCX_iz, I2NCX_iz, dE1NCX_izdt, dI1NCX_izdt, dI2NCX_izdt)
    '    INCXNa_iz = 3 * INCX_iz
    '    INCXCa_iz = -2 * INCX_iz

    '    calculateNCX(FrcNCX_blk, Nai, Cafree_blk, INCX_blk, INCXNa_blk, INCXCa_blk, E1NCX_blk, I1NCX_blk, I2NCX_blk, dE1NCX_blkdt, dI1NCX_blkdt, dI2NCX_blkdt)           'calculate  INaCa
    '    INCXNa_blk = 3 * INCX_blk
    '    INCXCa_blk = -2 * INCX_blk

    '    INCX = INCXNa_iz + INCXCa_iz + INCXNa_blk + INCXCa_blk

    'End Sub


    Public IsMClamp As Boolean

    Public Sub MembranePotential(ByVal sweeptime As Double)
        ItotCa_jnc = ICaLCa_LR_jnc + ICaLCa_L0_jnc
        ItotCa_iz = ICaLCa_iz + IbCa_iz + INCXCa_iz       '+ IPMCA_Ca_iz 
        ItotCa_blk = ICaLCa_blk + IbCa_blk + ICaT + INCXCa_blk    '+ IPMCA_Ca_blk 

        ItotNa_cyt = (ICaLNa_jnc + ICaLNa_iz + ICaLNa_blk) + (INCXNa_iz + INCXNa_blk) + (IKs_Na_iz + IKs_Na_blk) _
         + INaT_Na_cyt + INaL_Na_cyt + INaK_Na_cyt + IKto_Na_cyt + IbNSC_Na + ILCCa_Na_iz + ILCCa_Na_blk + Ist_Na + Iha_Na

        ItotK_cyt = (ICaLK_jnc + ICaLK_iz + ICaLK_blk) + IK1 + IKur + IKr + (IKs_K_iz + IKs_K_blk) _
           + INaT_K_cyt + INaL_K_cyt + IKto_K_cyt + IKpl + INaK_K_cyt + IKATP + IbNSC_K + ILCCa_K_iz + ILCCa_K_blk + IKACh + Ist_K + Iha_K

        '*************************************************************************************************************
        Itot_Ca = ItotCa_jnc + ItotCa_iz + ItotCa_blk
        Itot_Na = ItotNa_cyt
        Itot_K = ItotK_cyt

        Itot_cell = Itot_Ca + Itot_Na + Itot_K

        dVmdt = -Itot_cell - Iapl_blk

    End Sub

    Public Sub dIonConcdt()
        dCatot_jncdt = (-ItotCa_jnc * Cm / (2 * Faraday) + Jrel_SR - JCa_jnciz) / Vol_jnc
        dCatot_izdt = (-ItotCa_iz * Cm / (2 * Faraday) + JCa_jnciz - JCa_izblk) / Vol_iz

        dCatot_blkdt = (-ItotCa_blk * Cm / (2 * Faraday) - JSERCA + JCa_izblk) / Vol_blk '- CaTropC     'buffer generates Ca flux.

        dNaidt = -Itot_Na * Cm / (Vol_cyt * Faraday)

        dKidt = -(Itot_K + Iapl_blk) * Cm / (Vol_cyt * Faraday)   ' - (vKuni - vKHex) / Vol_cyt                'including Mitochondria

        dCa_SRupdt = (JSERCA - Jtrans_SR) / Vol_SRup                                                     'both fluxes refer to [Ca]up
        dCato_SRrldt = (Jtrans_SR - Jrel_SR) / Vol_SRrl

    End Sub

    Dim BathDiffusion As Double = 0.0005
    Dim pipetteDiffusion As Double = 0.0002
    Public Sub extConcentrations()
        Cao = Cao + BathDiffusion * dt * (newCao - Cao)
        Nao = Nao + BathDiffusion * dt * (newNao - Nao)
        Ko = Ko + BathDiffusion * dt * (newKo - Ko)
        ACh = ACh + BathDiffusion * dt * (newACh - ACh)

    End Sub

    Public Sub CaHomeostasis()              'Ion concentrations
        CaBuffblk()
        CaBuffiz_Inst()                      'instantaneous equilibrium
        CaBuffjnc_Inst()                    'instantaneous equilibrium
        CaBuffSR()
        CaBoundaryDiffusion()          'Ca diffusion among the three Ca-spaces; jnc-, iz- and blk-spaces
    End Sub

    Public Sub SRCadynamics()
        '***********' unit : amole / ms  *********************************************************
        TransferCaSR(Ca_SRup, Ca_SRrl, Jtrans_SR)

        'If constraints = False Then
        CaRU_Rate()             'calculate the Vm- and Ca-dependent rates of state transition in CaRU
        'Else
        'CaRU_RateSimple()             'calculate the Vm- and Ca-dependent rates of state transition in CaRU
        'End If

        CaRU_dydt()               '********** prepare dy/dt for each transition step ***

        CaRU_IonFluxes(Vm, FrcCaL_jnc, GHKNa, GHKK, ICaLCa_LR_jnc, ICaLCa_L0_jnc, ICaLNa_jnc, ICaLK_jnc, Jrel_SR)

        calculateSERCA(Cafree_blk, Ca_SRup, JSERCA, dATP_SERCA)       'Tran Crampin model

    End Sub


    '********** SR fluxe: RyR-kinetics, SR Ca pump, SR Ca leak **********
    Public Jrel_SR As Double
    Public JSERCA As Double
    Public JSERCA_TC As Double            'Tran Crampin model is used
    Public Jleak_SR As Double
    Public Jtrans_SR As Double                 'Transfer of Ca from uptake site to release site
    Public freezeSR As Double = 1

    Private Sub TransferCaSR(ByVal Ca_SRup As Double, ByVal Ca_SRrl As Double, ByRef Jtrans_SR As Double)
        Jtrans_SR = Ptrans * (Ca_SRup - Ca_SRrl) * freezeSR
    End Sub

    ''''''''''''''**********' ISERCA Hilgemann Model *  Now, this is not used in this SA node model ******************
    Public E1_ISERCA As Double
    Public dE1_ISERCAdt As Double
    Public dATP_SERCA As Double
    Public dATP_SERCA_TC As Double


    Public SERCAmag As Double = 1                           'scaling factor of Vmax in NCX


    Public Sub calculateSERCA(ByVal Cai As Double, ByVal Casr As Double, ByRef JSERCA As Double, ByRef dATP_SERCA As Double)
        'SERCA model of Tran K, Smith NP, Loiselle DS, Crampin EJ. Biophys J. 2009 Mar 4;96(5):2029-42

        Dim Hill As Double = 1.7
        Dim Kd_Cai As Double = 0.0027            '0.0017    mM
        Dim Kd_Casr As Double = 1.378                'mM

        Dim a1_plus As Double = 25900 * MgATP_cyt
        Dim a2_plus As Double = 2540 / (1 + Math.Pow(Kd_Cai / Cai, Hill))         '0.0025     k2_plus * Math.Pow(Cai / Kd_Cai, 2) / (Math.Pow(Cai / Kd_Cai, 2) + 0.0000284)
        Dim a3_plus As Double = 5.35 / (1 + Math.Pow(Casr / Kd_Casr, Hill))            '19.522 / (1.3812 * (1 + Math.Pow(Casr / Kd_Casr, 2)) + 2.2676)

        Dim a1_minus As Double = 0.1972 / (1 + Math.Pow(Cai / Kd_Cai, Hill))                 '0.0025    k1_minus * 0.0000028 / (Math.Pow(Cai / Kd_Cai, 2) + 0.0000284)
        Dim a2_minus As Double = 25435 * MgADP_cyt / (1 + Math.Pow(Kd_Casr / Casr, Hill))    'k2_minus * MgADP * Math.Pow(Casr / Kd_Casr, 2) * 1.3812 / (1.3812 * (1 + Math.Pow(Casr / Kd_Casr, 2)) + 2.2676)
        Dim a3_minus As Double = 149 * Pifree_cyt

        Dim denomi As Double = a2_plus * a3_plus + a1_minus * a3_plus + a1_minus * a2_minus + a1_plus * a3_plus + a2_minus * a1_plus + a2_minus * a3_minus + a1_plus * a2_plus + a3_minus * a1_minus + a3_minus * a2_plus
        Dim numer As Double = a1_plus * a2_plus * a3_plus - a1_minus * a2_minus * a3_minus
        Dim Vcyc As Double = numer / denomi * 6.86

        JSERCA = (SERCAmag * ampISERCA * Vcyc) / (2 * Faraday) * freezeSR / 1000                           'Vol_sr   amole / ms
        dATP_SERCA = JSERCA / 2                       'stoichiometry = 2Ca / 1ATP      amole / ms

    End Sub

    '*******************************************************************
    Public Sub NernstGHK()
        GHKNernst(ECa_jnc, ErevCa_jnc, GHKCa_jnc, dGHKCa_jnc, Cafree_jnc, Cao, 2)   'Cafree_jnc
        GHKNernst(ECa_jnc, ErevCa_LR, GHKCa_LR, dGHKCa_LR, Cafree_iz, Cao, 2)     'Cand_LR
        GHKNernst(ECa_jnc, ErevCa_L0, GHKCa_L0, dGHKCa_L0, Cafree_iz, Cao, 2)       'Cand_L0
        GHKNernst(ECa_iz, ErevCa_iz, GHKCa_iz, dGHKCa_iz, Cafree_iz, Cao, 2)
        GHKNernst(ECa_blk, ErevCa_blk, GHKCa_blk, dGHKCa_blk, Cafree_blk, Cao, 2)

        GHKNernst(ENa, ErevNa, GHKNa, dGHKNa, Nai, Nao, 1)
        GHKNernst(EK, ErevK, GHKK, dGHKK, Ki, Ko, 1)
    End Sub

    Public Sub GHKNernst(ByRef ES As Double, ByRef Erev As Double, ByRef GHKS As Double, ByRef dGHKS As Double, ByVal Si As Double, ByVal Se As Double, ByVal z As Double)
        ES = RTF / z * Log(Se / Si)

        If Vm > -0.00001 And Vm < 0.00001 Then Vm = 0.000001
        Dim B As Double = z * Vm / RTF
        Dim A As Double = Exp(-B)

        GHKS = B * (Si - Se * A) / (1 - A)      'GHK equation removed for P * zF
        dGHKS = ((Si - Se * A) / (1 - A) + (Se - Si) * B * A / (1 - A) ^ 2) / (RTF / z)
        Erev = Vm - GHKS / dGHKS
    End Sub

    '**********************  Myoplasmic Calcium Buffers ********************************************

    Public JCaB_blk As Double

    '********** Buffer  parameters  in blk **********
    Public newBAPTA As Double = 0                      'mM, default = 0        concentration in the pipette solution

    Public BtotBAPTA_blk As Double = 0                      'default = 0
    Const konBAPTA As Double = 940 / 10            'ms-1              approximation by decreasing rate ten-fold
    Const koffBAPTA As Double = 0.11938 / 10               ' mM-1, ms-1,   approximation by decreasing rate ten-fold
    Public BAPTACa_blk As Double                        'mM
    Public BAPTAf_blk As Double                            'mM 

    Public CaMf As Double
    Public TnChf As Double
    Public SRf As Double

    Public CaMCa As Double
    Public dCaMCadt As Double
    Public P1_dCaMCadt As Double

    Const BtotCaM As Double = 0.024                   '0.024     0.05           'mM
    Const KoffCaM As Double = 0.238 / 10               'ms^-1                      decreased to avoid oscillation
    Const KonCaM As Double = 34 / 10                    'ms^-1mM^-1          decreased to avoid oscillation
    Const KdCaM As Double = KoffCaM / KonCaM           'mM, dissociation constant

    Public TnChCa As Double
    Public dTnChCadt As Double
    Public P1_dTnChCadt As Double

    Const BtotTnCh As Double = 0.007                                 '0.12        '0.14         'mM
    Const KonTnCh As Double = 2.37                        'ms^-1mM^-1
    Const KoffTnCh As Double = 0.000032               'ms^-1

    Public SRCa As Double
    Public dSRCadt As Double
    Public P1_dSRCadt As Double

    Const BtotSR As Double = 0.0171                  '19 * 0.0009           'mM
    Const KonSR As Double = 100 / 10                      'ms^-1mM^-1          decreased to avoid oscillation
    Const KoffSR As Double = 0.06 / 10                     'ms^-1                    decreased to avoid oscillation
    Const KdSR As Double = KoffSR / KonSR           'mM, dissociation constant

    Private Sub CaBuffblk()
        Cafree_blk = Catot_blk - (CaMCa + TnChCa + SRCa) - nCaBS * (TSCa3 + TSCa3W + TSCa3S) / 1000            'mM, calculate free Ca concentration

        dCaMCadt = KonCaM * Cafree_blk * (BtotCaM - CaMCa) - KoffCaM * CaMCa          'mM, calculate rate of change in CaM-Ca
        dTnChCadt = KonTnCh * Cafree_blk * (BtotTnCh - TnChCa) - KoffTnCh * TnChCa  'caluclate rate of change in TnCh-Ca
        dSRCadt = KonSR * Cafree_blk * (BtotSR - SRCa) - KoffSR * SRCa                             'calculate rate of change in SR-Ca

    End Sub


    '*********************** Instantaneous Ca buffers ************************************

    '********** Buffer  parameters  in iz **********
    Public JCaB_iz As Double         '=SLLsl + SLHsl

    Const BtotL_iz As Double = 0.6078          '0.0374 * Vol_blk / Vol_iz        'mM
    Const koffL_iz As Double = 1.3                            'ms^-1    sarcolemmal Ca buffer with a low affinity
    Const KonL_iz As Double = 100                             'ms^-1mM^-1   sarcolemmal Ca buffer with a low affinity
    Dim KdL_iz As Double = koffL_iz / KonL_iz                   'dissociation constant of the low affinity buffer in the iz/jnc space

    Const BtotH_iz As Double = 0.2178          '0.0134 * Vol_blk / Vol_iz       'mM
    Const KoffH_iz As Double = 0.03                           'ms^-1
    Const KonH_iz As Double = 100                             'ms^-1mM^-1
    Dim KdH_iz As Double = KoffH_iz / KonH_iz                'dissociation constant of the high affinity buffer in the iz/jnc space

    Public LCa_iz As Double                'bound form
    Public HCa_iz As Double                 'bound form
    Public Lf_iz As Double                       'free buffer with a low affinity
    Public Hf_iz As Double                      'free buffer with a high affinity

    Private Sub CaBuffiz_Inst()                  'This is used to calculate the instantaneous equilibrium of binding reactions.
        Lf_iz = BtotL_iz - LCa_iz                       'free buffer having a low affinity
        Hf_iz = BtotH_iz - HCa_iz                      'free buffer having a high affinity
        'Catot_iz = Cafree_iz + LCa_iz + HCa_iz      '+ BAPTACa_iz         'total Ca within sl

        Dim N As Long = 0                                                                  'counter of the Do Loop
        Do
            Cafree_iz = Catot_iz / (1 + Lf_iz / KdL_iz + Hf_iz / KdH_iz)     ' + BAPTAf_iz / KdBAPTA)      'calculate free Ca

            Lf_iz = BtotL_iz / (1 + (Cafree_iz / KdL_iz))                            'Calculate free buffer with a low affinity
            Hf_iz = BtotH_iz / (1 + (Cafree_iz / KdH_iz))                          'calculate free buffer with a high affinity
            N = N + 1
        Loop While N < 10                                                              '10 cycles seems to be good enough.

        LCa_iz = BtotL_iz - Lf_iz                    'Ca-bound buffer having a low affinity 
        HCa_iz = BtotH_iz - Hf_iz                  'Ca-bound buffer having a high affinity 

    End Sub

    '********** Buffer  parameters  in jnc **********

    Public JCaB_jnc As Double       '

    Const BtotL_jnc As Double = 1.1095             ''0.00046 * Vol_blk / Vol_jnc      'mM   Note, the on- and off-rates are the same as in sl
    Public LCa_jnc As Double

    Const BtotH_jnc As Double = 0.398              ' '0.000165 * Vol_blk / Vol_jnc    'mM
    Public HCa_jnc As Double

    Dim Lf_jnc As Double
    Dim Hf_jnc As Double

    Private Sub CaBuffjnc_Inst()                   'This is used to calculate the instantaneous equilibrium of binding reactions.
        Lf_jnc = BtotL_jnc - LCa_jnc                      'free buffer with a low affinity;
        Hf_jnc = BtotH_jnc - HCa_jnc                    'free buffer with a high affinity
        'Catot_jnc = Cafree_jnc + LCa_jnc + HCa_jnc        'total Ca within sl

        Dim N As Double = 0
        Do
            Cafree_jnc = Catot_jnc / (1 + Lf_jnc / KdL_iz + Hf_jnc / KdH_iz)      'calculate free Ca using the same on off rates

            Lf_jnc = BtotL_jnc / (1 + (Cafree_jnc / KdL_iz))                                                       'Calculate free buffer with a low affinity
            Hf_jnc = BtotH_jnc / (1 + (Cafree_jnc / KdH_iz))                                                     'calculate free buffer with a high affinity
            N = N + 1
        Loop While N < 10                                                              '10 cycles seems to be good enough.
        LCa_jnc = BtotL_jnc - Lf_jnc                    'Ca-bound buffer with a low affinity buffer
        HCa_jnc = BtotH_jnc - Hf_jnc                  'Ca-bound buffer with a high affinity buffer
    End Sub


    '********** Buffer  parameters  in SR **********
    Const Btot_Csqn As Double = 10 '3
    Const KoffCsqn As Double = 65                            'ms^-1
    Const KonCsqn As Double = 100                            'ms^-1mM^-1

    Private Sub CaBuffSR()

        Dim KdCsqnCa As Double = KoffCsqn / KonCsqn
        Dim aa As Double = 1
        Dim bb As Double = Btot_Csqn - Catot_SRrl + KdCsqnCa                 '(Btotal - Catotal + KD)
        Dim cc As Double = -KdCsqnCa * Catot_SRrl                                     'KD * Catotal
        Ca_SRrl = (-bb + Sqrt(bb * bb - 4 * aa * cc)) / (2 * aa)                 'solution of second-order equation

    End Sub

    '********** Diffusion between spaces **********
    Public Const GCa_jnciz As Double = 459.4 * 20 * 3.5        '32158 * ratioCm * ratioVolume   
    Public Const GCa_izblk As Double = 2076.1                      '2076.1 * ratioCm * ratioVolume    

    Public JCa_jnciz As Double
    Public JCa_izblk As Double
    Public A As Double
    Public B As Double

    Private Sub CaBoundaryDiffusion()
        JCa_jnciz = GCa_jnciz * (Cafree_jnc - Cafree_iz)
        A = Cafree_jnc - Cafree_iz

        JCa_izblk = GCa_izblk * (Cafree_iz - Cafree_blk)
        B = (Cafree_iz - Cafree_blk)

    End Sub

    '********* INaK **********************
    Public maxINaK As Double = 68.68 * 1.2              '25.1779 * 1.24 * 2.2            ' 

    Public INaKmag As Double = 2   '0.952     ' = 0.85 * 1.12           '*1.12, 1.15, etc    'revised on 180918 by AN. a scaling factor of Na/K pump
    'Public crf_NaK As Double = 1                         'To keep Nai close to stdNai, the INaKmag is scaled by crf_NaK,   INaKmag * crf_NaK

    Public INaK As Double
    Public INaK_Na_cyt As Double
    Public INaK_K_cyt As Double

    Public P1_6_NaK As Double
    Public P7_NaK As Double
    Public P8_13_NaK As Double
    Public P14_15_NaK As Double

    Public dP1_6_NaKdt As Double
    Public dP7_NaKdt As Double
    Public dP8_13_NaKdt As Double

    Public dATP_NaK As Double

    Public k1p_NaK As Double = 0.72        '
    Public k1m_NaK As Double = 0.08      '
    Public k2p_NaK As Double = 0.08       '
    Public k2m_NaK As Double = 0.008    '
    Public k3p_NaK As Double = 4             '
    Public k3m_NaK As Double = 8000     '
    Public k4p_NaK As Double = 0.3         '
    Public k4m_NaK As Double = 0.2        '

    Public KdKe0_NaK As Double = 0.8         '
    Public KdKi0_NaK As Double = 18.8        '
    Public KdNae0_NaK As Double = 26.8        '
    Public KdNai0_NaK As Double = 5              '
    Public KdMgATP_NaK As Double = 0.6     '

    Public stoiNa_NaK As Double = 3                  '
    Public stoiK_NaK As Double = -2                   '
    Public dV_Nae_NaK As Double = 0.44             '
    Public dV_Nai_NaK As Double = -0.14            '
    Public dV_Ke_NaK As Double = 0.23            '
    Public dV_Ki_NaK As Double = -0.14            '

    Public fVm_NaK As Double
    Public KdNao_NaK As Double
    Public KdNai_NaK As Double
    Public KdKo_NaK As Double
    Public KdKi_NaK As Double


    Public Sub calculateNaK(ByVal Nai As Double, ByVal Ki As Double, ByVal MgATP_cyt As Double, ByVal MgADP_cyt As Double, ByRef INaK As Double, ByRef INaK_Na As Double, ByRef INaK_K As Double, ByRef dNATP_NaK As Double,
                      ByVal P1_6 As Double, ByVal P7 As Double, ByVal P8_13 As Double, ByRef dP1_6dt As Double, ByRef dP7dt As Double, ByRef dP8_13dt As Double)

        fVm_NaK = Faraday * Vm / (R * tempK)                     'VF/R/T
        KdNao_NaK = KdNae0_NaK * Exp(dV_Nae_NaK * fVm_NaK)     'relative Kd 
        KdNai_NaK = KdNai0_NaK * Exp(dV_Nai_NaK * fVm_NaK)       'relative Kd 
        KdKo_NaK = KdKe0_NaK * Exp(dV_Ke_NaK * fVm_NaK)           'relative Kd 
        KdKi_NaK = KdKi0_NaK * Exp(dV_Ki_NaK * fVm_NaK)             'relative Kd 

        Dim Nai_ As Double = Nai / KdNai_NaK                 'relative concentration referring to the dissociation constant 
        Dim Nae_ As Double = Nao / KdNao_NaK            'relative concentration referring to the dissociation constant
        Dim Ki_ As Double = Ki / KdKi_NaK                       'relative concentration referring to the dissociation constant
        Dim Ke_ As Double = Ko / KdKo_NaK                 'relative concentration referring to the dissociation constant

        Dim a1f_NaK As Double = (k1p_NaK * (Nai_ ^ 3.0)) / (((1 + Nai_) ^ 3) + ((1 + Ki_) ^ 2) - 1)
        Dim a2f_NaK As Double = k2p_NaK
        Dim a3f_NaK As Double = k3p_NaK * (Ke_ ^ 2) / (((1 + Nae_) ^ 3) + ((1 + Ke_) ^ 2) - 1)
        Dim a4f_NaK As Double = k4p_NaK * MgATP_cyt / KdMgATP_NaK / (1 + MgATP_cyt / KdMgATP_NaK)

        ' transition rates in the backward direction
        Dim a1b_NaK As Double = k1m_NaK * MgADP_cyt
        Dim a2b_NaK As Double = k2m_NaK * (Nae_ ^ 3) / (((1 + Nae_) ^ 3) + ((1 + Ke_) ^ 2) - 1)
        Dim a3b_NaK As Double = k3m_NaK * Pifree_cyt * H_cyt / (1 + MgATP_cyt / KdMgATP_NaK)
        Dim a4b_NaK As Double = k4m_NaK * (Ki_ ^ 2) / (((1 + Nai_) ^ 3) + ((1 + Ki_) ^ 2) - 1)

        Dim P14_15 As Double = (1 - P1_6 - P7 - P8_13)

        Dim vstep1_NaK As Double = (a2f_NaK * P7 - a2b_NaK * P8_13)
        '************* dy/dt for the state transitions *************************************
        dP1_6dt = (-a1f_NaK * P1_6 + a1b_NaK * P7 + a4f_NaK * P14_15 - a4b_NaK * P1_6)
        dP7dt = (-a2f_NaK * P7 + a2b_NaK * P8_13 + a1f_NaK * P1_6 - a1b_NaK * P7)
        dP8_13dt = (-a3f_NaK * P8_13 + a3b_NaK * P14_15 + a2f_NaK * P7 - a2b_NaK * P8_13)

        '******************* magnitude of INaK **********************************************
        Dim vcyc_NaK As Double = vstep1_NaK                                                    'any step when calculating the steady state distribution
        INaK = maxINaK * vcyc_NaK * INaKmag * crf_NaK    ' current  density (pA/pF)
        INaK_Na = stoiNa_NaK * INaK        ' calculate ion flux
        INaK_K = stoiK_NaK * INaK

        dNATP_NaK = INaK * Cm / Faraday           'amole / ms

    End Sub


    Public Sub calculateNaKSimple(ByVal v As Double, ByVal Nai As Double, ByVal Ki As Double, ByVal MgATP_cyt As Double, ByVal MgADP_cyt As Double, ByRef INaK As Double, ByRef INaK_Na As Double, ByRef INaK_K As Double, ByRef dNATP_NaK As Double,
                      ByVal P1_6 As Double, ByVal P7 As Double, ByVal P8_13 As Double, ByRef dP1_6dt As Double, ByRef dP7dt As Double, ByRef dP8_13dt As Double)

        fVm_NaK = Faraday * v / (R * tempK)                     'VF/R/T
        KdNao_NaK = KdNae0_NaK * Exp(dV_Nae_NaK * fVm_NaK)     'relative Kd 
        KdNai_NaK = KdNai0_NaK * Exp(dV_Nai_NaK * fVm_NaK)       'relative Kd 
        KdKo_NaK = KdKe0_NaK * Exp(dV_Ke_NaK * fVm_NaK)           'relative Kd 
        KdKi_NaK = KdKi0_NaK * Exp(dV_Ki_NaK * fVm_NaK)             'relative Kd 

        Dim Nai_ As Double = Nai / KdNai_NaK                 'relative concentration referring to the dissociation constant 
        Dim Nae_ As Double = Nao / KdNao_NaK            'relative concentration referring to the dissociation constant
        Dim Ki_ As Double = Ki / KdKi_NaK                       'relative concentration referring to the dissociation constant
        Dim Ke_ As Double = Ko / KdKo_NaK                 'relative concentration referring to the dissociation constant

        'Dim MgATP_ As Double = ATPMg / kd_MgATP  'relative concentration of ATP referring to the dissociation constant
        'Dim Kicongen As Double = 0                                 'used only when inclucing Cs+ to replace K+ 
        'Dim Cseffect As Double = Kicongen * (2 * Ki_ + 3 * Nai_ + 1)        'used only when inclucing Cs+ to replace K+ 
        ' transition rates in the forward direction

        Dim a1f_NaK As Double = (k1p_NaK * (Nai_ ^ 3.0)) / (((1 + Nai_) ^ 3) + ((1 + Ki_) ^ 2) - 1)
        Dim a2f_NaK As Double = k2p_NaK
        Dim a3f_NaK As Double = k3p_NaK * (Ke_ ^ 2) / (((1 + Nae_) ^ 3) + ((1 + Ke_) ^ 2) - 1)
        Dim a4f_NaK As Double = k4p_NaK * MgATP_cyt / KdMgATP_NaK / (1 + MgATP_cyt / KdMgATP_NaK)

        ' transition rates in the backward direction
        Dim a1b_NaK As Double = k1m_NaK * MgADP_cyt
        Dim a2b_NaK As Double = k2m_NaK * (Nae_ ^ 3) / (((1 + Nae_) ^ 3) + ((1 + Ke_) ^ 2) - 1)
        Dim a3b_NaK As Double = k3m_NaK * Pifree_cyt * H_cyt / (1 + MgATP_cyt / KdMgATP_NaK)
        Dim a4b_NaK As Double = k4m_NaK * (Ki_ ^ 2) / (((1 + Nai_) ^ 3) + ((1 + Ki_) ^ 2) - 1)

        Dim P14_15 As Double = (1 - P1_6 - P7 - P8_13)

        Dim vstep1_NaK As Double = (a2f_NaK * P7 - a2b_NaK * P8_13)
        '************* dy/dt for the state transitions *************************************
        dP1_6dt = (-a1f_NaK * P1_6 + a1b_NaK * P7 + a4f_NaK * P14_15 - a4b_NaK * P1_6)
        dP7dt = (-a2f_NaK * P7 + a2b_NaK * P8_13 + a1f_NaK * P1_6 - a1b_NaK * P7)
        dP8_13dt = (-a3f_NaK * P8_13 + a3b_NaK * P14_15 + a2f_NaK * P7 - a2b_NaK * P8_13)

        '******************* magnitude of INaK **********************************************
        Dim vcyc_NaK As Double = vstep1_NaK                                                    'any step when calculating the steady state distribution
        INaK = maxINaK * vcyc_NaK * INaKmag * crf_NaK    ' current  density (pA/pF)
        INaK_Na = stoiNa_NaK * INaK        ' calculate ion flux
        INaK_K = stoiK_NaK * INaK

        dNATP_NaK = INaK * Cm / Faraday           'amole / ms

    End Sub

    Public orgcrf_NaK As Double = 0.24
    Public crf_NaK As Double
    'Public stdNai As Double = 6.4      'N-like cells  6.2132        '6.1856            'mM,   optimum Nai obtained in the SS of the model
    Public stdNai As Double = 6.1      'V-like cells 6.2132        '6.1856            'mM,   optimum Nai obtained in the SS of the model
    Public slope As Double = 1.2

    Public Sub RegulatemaxINaK()
        'crf_NaK = orgcrf_NaK - (stdNai - Nai) * slope  ' 0.6      
        'If crf_NaK < 0.01 Then crf_NaK = 0.01 'avoid a negative crf_NaK      less sensitive

        crf_NaK = crf_NaK - (stdNai - Nai) * 0.3    ' N-like cells   0.3   provides much finer regulation
    End Sub


    Public orgcrf_NCX As Double = 1
    Public crf_NCX As Double
    Public stdtotCai As Double = 79       'atto mole  10^-18 mole
    Public totalCai As Double                 'atto mole  10^-18 mole

    Public wholeCai As Double

    Public Sub RegulateCaitotal()
        'crf_NCX = orgcrf_NCX - (stdtotCai - totalCai) * 0.008  '0.02
        crf_NCX = crf_NCX - (stdtotCai - totalCai) * 0.008  '0.02
    End Sub


    ''''''''*****************************    INCX   **********************************************
    Public maxINCX As Double = 29 * 1.2    '61.06 * 0.5 * 0.95        'NCX 
    '************** parameters for the Ca- and Na-dependent inactivation *************************
    Dim a1off_NCX As Double = 0.0015
    Dim a1on_NCX As Double = 0.002
    Dim b1off_NCX As Double = 0.0000005
    Dim b1on_NCX As Double = 0.0012
    Dim a2off_NCX As Double = 0.02
    Dim a2on_NCX As Double = 0.00006
    Dim b2off_NCX As Double = 0.0002
    Dim b2on_NCX As Double = 0.18
    '************** parameters for the Ca- and Na-binding to transport sites *************************
    Dim KmNao_NCX As Double = 87.5                        'mM
    Dim KmNai_NCX As Double = 20.74854
    Dim KmCao_NCX As Double = 1.38
    Dim KmCai_NCX As Double = 0.0184
    Dim Kmact_NCX As Double = 0.004
    Dim nHNa_NCX As Double = 3
    Dim partition_NCX As Double = 0.32


    Public INCX As Double

    '********** iz ***********************
    Public INCX_iz As Double
    Public INCXNa_iz As Double
    Public INCXCa_iz As Double

    Public I1NCX_iz As Double
    Public I2NCX_iz As Double
    Public E1NCX_iz As Double
    Public E2NCX_iz As Double

    Public dE1NCX_izdt As Double
    Public dI1NCX_izdt As Double
    Public dI2NCX_izdt As Double

    '************** blk *******************
    Public INCX_blk As Double
    Public INCXNa_blk As Double
    Public INCXCa_blk As Double

    Public I1NCX_blk As Double
    Public I2NCX_blk As Double
    Public E1NCX_blk As Double
    Public E2NCX_blk As Double

    Public dE1NCX_blkdt As Double
    Public dI1NCX_blkdt As Double
    Public dI2NCX_blkdt As Double

    'Public fNaCa As Double = 1
    Public INCXmag As Double = 2    '1.3
    'Public crfNCX As Double = 1

    Public E2Na As Double
    Public E2Ca As Double

    Public E1Na As Double
    Public E1Ca As Double

    Public fCaina As Double

    Public alpha1 As Double
    Public beta1 As Double

    Public alpha2 As Double
    Public beta2 As Double

    Public k1_NCX As Double
    Public k2_NCX As Double
    Public k3_NCX As Double = 1.0
    Public k4_NCX As Double = 1.0

    Public alphaE As Double
    Public betaE As Double


    Public Sub calculateNCX(ByVal frc As Double, ByVal Nai As Double, ByVal Cai As Double, ByRef INCX As Double, ByRef INCX_Na As Double, ByRef INCX_Ca As Double,
                           ByVal E1_NCX As Double, ByVal I1_NCX As Double, ByVal I2_NCX As Double, ByRef dE1_NCXdt As Double, ByRef dI1_NCXdt As Double, ByRef dI2_NCXdt As Double)

        Dim E2_NCX As Double = 1 - E1_NCX - I1_NCX - I2_NCX        'probability of each conformation state

        '************** kinetics of NCX,  Vm-independent part *******************************************************
        E2Na = 1.0 / (1.0 + ((KmNao_NCX / Nao) ^ nHNa_NCX) * (1.0 + Cao / KmCao_NCX))    'Na binding probability
        E2Ca = 1.0 / (1.0 + (KmCao_NCX / Cao) * (1.0 + ((Nao / KmNao_NCX) ^ nHNa_NCX)))   'Ca binding probability

        E1Na = 1.0 / (1.0 + ((KmNai_NCX / Nai) ^ nHNa_NCX) * (1 + Cai / KmCai_NCX))        'Na binding probability
        E1Ca = 1.0 / (1.0 + (KmCai_NCX / Cai) * (1.0 + ((Nai / KmNai_NCX) ^ nHNa_NCX)))   'Ca binding probability

        fCaina = Cai / (Cai + Kmact_NCX)          'Ca-dependency of the inactivation step

        ' I1 inactivation gate (Na-dependent, Vm-independent)
        alpha1 = (E1Na * (fCaina * a1on_NCX + (1 - fCaina) * a1off_NCX))   'rate from E1 to I1
        beta1 = (fCaina * b1on_NCX + (1 - fCaina) * b1off_NCX)                    'rate from I1 to E1

        ' I2 inactivation gate (Ca-dependent, Vm-independent)
        alpha2 = (fCaina * a2on_NCX + (1 - fCaina) * a2off_NCX)              'rate from E1 to I2
        beta2 = (fCaina * b2on_NCX + (1 - fCaina) * b2off_NCX)                 'rate from I2 to E1

        '**************  Vm-dependent part *******************************************************
        k1_NCX = Exp(partition_NCX * Vm / (R * tempK / Faraday))                            'voltage-dependent rate
        k2_NCX = Exp((partition_NCX - 1.0) * Vm / (R * tempK / Faraday))                  'voltage-dependent rate

        alphaE = k2_NCX * E2Na + k4_NCX * E2Ca                  'overall transition rate from E2 to E1, Vm-dependent
        betaE = k1_NCX * E1Na + k3_NCX * E1Ca                    'overall transition rate from E1 to E2, Vm-dependent


        '**********************  dy/dt for the Euler Integration *******************
        'If isIVcurve <> True Then                       'when no IV-curve is calculated
        dE1_NCXdt = E2_NCX * alphaE + I1_NCX * beta1 + I2_NCX * beta2 - E1_NCX * (betaE + alpha1 + alpha2)
        dI1_NCXdt = E1_NCX * alpha1 - I1_NCX * beta1
        dI2_NCXdt = E1_NCX * alpha2 - I2_NCX * beta2
        'End If

        '************** Amplitude  of NCX flux *******************************************************
        'INCX = frc * maxINCX * INCXmag * fNaCa * (k1_NCX * E1Na * E1_NCX - k2_NCX * E2Na * E2_NCX)
        INCX = frc * maxINCX * INCXmag * (k1_NCX * E1Na * E1_NCX - k2_NCX * E2Na * E2_NCX) * crf_NCX
        INCX_Na = 3 * INCX
        INCX_Ca = -2 * INCX

    End Sub


    '***** IKpl *********
    Public orgP_Kpl As Double = 0.0028 * 0.1             '
    Public P_Kpl As Double                '
    Public IKpl As Double
    Public IKplmag As Double = 1

    Public Sub calculateIKpl()
        ''******** Ko-dependence of IK plateau channel   *****************
        Dim Vshift As Double = 38 '3
        Dim Vslope As Double = 30 '13

        '******** Ko-dependence of IK plateau channel   *****************
        P_Kpl = orgP_Kpl * IKplmag * (Ko / 5.4) ^ 0.16 / Cm        '* 1.5   current density  K Component,  conductance increases with higher Ke.

        If Vm = -Vshift Then                                                '******* outward rectifier *****
            IKpl = P_Kpl * GHKK * 13.0077
        Else
            IKpl = P_Kpl * GHKK * (Vm + Vshift) / (1 - Math.Exp(-(Vm + Vshift) / Vslope))
        End If
    End Sub

    '*****Ibnsc**********

    Public IbNSC As Double                      '=ILNSCNa + ILNSCK
    Public IbNSC_Na As Double              '= PILNSCNa * NaCF   Na component of the Non-selective Cation current
    Public IbNSC_K As Double                 '= PILNSCK * KCF
    Public P_bnscNa As Double = 0.000182875     '=0.0532 * 1.1 / 10 / Cm       pA/mM  background Na current
    Public P_bnscK As Double = P_bnscNa * 0.4            'pA/mM  background K current     'leak K component
    Public IbNSCmag As Double = 1            '1.3

    Public Sub calculateIbNSC(ByVal NaCF As Double, ByVal KCF As Double)

        IbNSC_Na = P_bnscNa * IbNSCmag * NaCF * IbNSCsf
        IbNSC_K = P_bnscK * IbNSCmag * KCF * IbNSCsf
        IbNSC = IbNSC_Na + IbNSC_K

    End Sub

    'Public Sub calculateIbNSCSimple(ByVal NaCF As Double, ByVal KCF As Double)

    '    IbNSC_Na = P_bnscNa * IbNSCmag * NaCF * IbNSCsf
    '    IbNSC_K = P_bnscK * IbNSCmag * KCF * IbNSCsf
    '    IbNSC = IbNSC_Na + IbNSC_K

    'End Sub


    '********** IbCa **********
    Public IbCa As Double                      '= PbCa * CaCF  '(Vmembrane - ECa)
    Public IbCa_iz As Double
    Public IbCa_blk As Double

    Public P_bCa As Double = 0.00125       '0.004 / Cm * 10 = 0.00125          current density               2.75   backgrund Ca conductance

    Public Sub calculateIbCa(ByVal CaCF As Double, ByRef IbCa As Double)
        IbCa = 2 * P_bCa * CaCF
    End Sub

    '****** ILCa *********
    'Public ILCCa_Na As Double                 'Na component of Leak cation current induced by the intracellular Ca Ehara, Noma & Ono, 1988.
    'Public ILCCa_K As Double                   'K component of Leak cation current induced by the intracellular Ca Ehara, Noma & Ono, 1988.
    Public ILCCa As Double                     'Leak cation current induced by the intracellular Ca Ehara, Noma & Ono, 1988.

    Public ILCCa_iz As Double
    Public ILCCa_blk As Double
    Public ILCCa_Na_iz As Double
    Public ILCCa_K_iz As Double
    Public ILCCa_Na_blk As Double
    Public ILCCa_K_blk As Double

    Public PLCCa As Double = 0.01375            '=0.44 / Cm     pA/mM   conductance for the ILCCa
    Public Sub calculateIL_Ca(ByVal frc As Double, ByVal ca As Double, ByRef ILCCa_Na As Double, ByRef ILCCa_K As Double, ByRef ILCCa As Double)
        ILCCa_Na = frc * PLCCa * GHKNa / (1 + (0.0012 / ca) ^ 3)    'Leak cation current induced by the intracellular Ca Ehara, Noma & Ono, 1988.
        ILCCa_K = frc * PLCCa * GHKK / (1 + (0.0012 / ca) ^ 3)      'Leak cation current induced by the intracellular Ca Ehara, Noma & Ono, 1988.
        ILCCa = ILCCa_Na + ILCCa_K
    End Sub

    Public ICaTInf As Double
    Public ICaT As Single
    Public ICaT_blk As Double
    Public ICaT_iz As Double
    Public ICaTmag As Double = 1                       'scaling factor for the ICaT magnitude

    Public PCaT As Double = 306 / Cm                   'T-type Ca channel amplitude
    Public dalfa_CaT As Double
    Public dbeta_CaT As Double
    Public falfa_CaT As Double
    Public fbeta_CaT As Double

    Public d_CaT As Double            'ICaT
    Public dd_CaTdt As Double
    Public f_CaT As Double
    Public df_CaTdt As Double
    Public pO_CaT As Double

    '***** ICaT **********
    Public Sub calculateICaT(ByVal CaCF As Double, ByRef ICaT As Double)
        dalfa_CaT = 1 / (0.019 * Math.Exp(Vm / -5.6) + 0.82 * Math.Exp(Vm / -250))
        dbeta_CaT = 1 / (40 * Math.Exp(Vm / 6.3) + 1.5 * Math.Exp(Vm / 10000))
        falfa_CaT = 1 / (62000 * Math.Exp(Vm / 10.1) + 30 * Math.Exp(Vm / 3000))
        fbeta_CaT = 1 / (0.0006 * Math.Exp(Vm / -6.7) + 1.2 * Math.Exp(Vm / -25))

        dd_CaTdt = dalfa_CaT * (1 - d_CaT) - dbeta_CaT * d_CaT
        df_CaTdt = falfa_CaT * (1 - f_CaT) - fbeta_CaT * f_CaT

        pO_CaT = d_CaT * f_CaT
        ICaTInf = 2 * PCaT * CaCF * ICaTmag * ICaTsf        '= 2 * 大きさ * Caの流束
        ICaT = ICaTInf * pO_CaT
    End Sub

    Public Sub calculateICaTSimple()
        'dalfa_CaT = 1 / (0.019 * Math.Exp(Vm / -5.6) + 0.82 * Math.Exp(Vm / -250))
        'dbeta_CaT = 1 / (40 * Math.Exp(Vm / 6.3) + 1.5 * Math.Exp(Vm / 10000))
        'falfa_CaT = 1 / (62000 * Math.Exp(Vm / 10.1) + 30 * Math.Exp(Vm / 3000))
        'fbeta_CaT = 1 / (0.0006 * Math.Exp(Vm / -6.7) + 1.2 * Math.Exp(Vm / -25))

        'dd_CaTdt = dalfa_CaT * (1 - d_CaT) - dbeta_CaT * d_CaT
        'df_CaTdt = falfa_CaT * (1 - f_CaT) - fbeta_CaT * f_CaT
        dd_CaTdt = 0.0
        df_CaTdt = 0.0


        'pO_CaT = d_CaT * f_CaT

        'pO_CaT = pO_CaT(I)
        ICaTInf = 2 * PCaT * GHKCa_blk * ICaTmag         '= 2 * 大きさ * Caの流束
        ICaT = ICaTInf * pO_CaT
    End Sub

    '***** Ist **********
    Public Ist As Double                      'Ist = IstNa + IstK
    Public Ist_Na As Double                    'Na component
    Public Ist_K As Double                     'K component
    Public IstNainf As Double
    Public PstNa As Double                     'Na conductance
    Public Pstmag As Double = 1        'relative  conductance
    Public IstKinf As Single
    Public RstK As Double = 0.585                  'K conductance
    Public RstKmag As Double = 1                  'K conductance
    Public originalPstNa As Double = 0.00236          'Na conductance


    Public vShift_st As Double = 0           ' to shift voltage gate
    'Public Ist_Mag As Double = 1             'scaling factor for Ist magnitude
    Public alfad_st As Double
    Public betad_st As Double
    Public alfaf_st As Double
    Public betaf_st As Double
    Public alfaus_st As Double
    Public betaus_st As Double

    Public d_st As Double                   'Ist
    Public f_st As Double
    Public us_st As Double
    Public dd_stdt As Double
    Public df_stdt As Double
    Public dus_stdt As Double

    Public pdus_Ist As Double          '(d * us)
    Public pO_Ist As Double


    Public Sub calculateIst(ByVal Caf_cyt As Double)
        '***********************   Ist  ***********************************
        alfad_st = 1 / (0.15 * Math.Exp((Vm - vShift_st) / -11) + 0.2 * Math.Exp((Vm - vShift_st) / -700))            'from rat data, alpha, activation parameter for IST
        betad_st = 1 / (16 * Math.Exp((Vm - vShift_st) / 8) + 15 * Math.Exp((Vm - vShift_st) / 50))                        'from rat data, beta
        dd_stdt = alfad_st * (1 - d_st) - betad_st * d_st

        alfaf_st = 1 / (3100 * Math.Exp((Vm - vShift_st) / 13) + 700 * Math.Exp((Vm - vShift_st) / 70))             'from rat data, alpha, activation parameter for IST
        betaf_st = 1 / (95 * Math.Exp((Vm - vShift_st) / -10) + 50 * Math.Exp((Vm - vShift_st) / -700)) + Caf_cyt * 2.5 / (1 + Math.Exp(Vm / -5))                       'from rat data, beta
        df_stdt = alfaf_st * (1 - f_st) - betaf_st * f_st

        alfaus_st = 1 / (400000 * Math.Exp(Vm / 9) + 60 * Math.Exp(Vm / 65))
        betaus_st = 1 / (700 * Math.Exp(Vm / -14) + 60 * Math.Exp(Vm / -65))
        dus_stdt = alfaus_st * (1 - us_st) - betaus_st * us_st

        IstNainf = originalPstNa * GHKNa * Pstmag * Istsf
        IstKinf = RstK * originalPstNa * GHKK * Pstmag * Istsf

        pO_Ist = d_st * f_st * us_st
        Ist_Na = IstNainf * pO_Ist
        Ist_K = IstKinf * pO_Ist
        Ist = Ist_Na + Ist_K
    End Sub


    Public Sub calculateIstSimple(ByVal tgVm As Double)

        'dd_stdt = 0.0
        '************ Vm-gate   Ist (constraints; prepared by using the command potential )  ***********************************
        alfad_st = 1 / (0.15 * Math.Exp((Vm - vShift_st) / -11) + 0.2 * Math.Exp((Vm - vShift_st) / -700))            'from rat data, alpha, activation parameter for IST
        betad_st = 1 / (16 * Math.Exp((Vm - vShift_st) / 8) + 15 * Math.Exp((Vm - vShift_st) / 50))                        'from rat data, beta
        dd_stdt = alfad_st * (1 - d_st) - betad_st * d_st

        '****** calculate f_st using the expVm(I) ****************** 
        alfaf_st = 1 / (3100 * Math.Exp((tgVm - vShift_st) / 13) + 700 * Math.Exp((tgVm - vShift_st) / 70))             'from rat data, alpha, activation parameter for IST
        betaf_st = 1 / (95 * Math.Exp((tgVm - vShift_st) / -10) + 50 * Math.Exp((tgVm - vShift_st) / -700)) + Cafree_blk * 2.5 / (1 + Math.Exp(tgVm / -5))                       'from rat data, beta
        df_stdt = alfaf_st * (1 - f_st) - betaf_st * f_st

        dus_stdt = 0.0
        'alfaus_st = 1 / (400000 * Math.Exp(Vm / 9) + 60 * Math.Exp(Vm / 65))         '
        'betaus_st = 1 / (700 * Math.Exp(Vm / -14) + 60 * Math.Exp(Vm / -65))

        IstNainf = originalPstNa * GHKNa * Pstmag * Istsf
        IstKinf = RstK * originalPstNa * GHKK * Pstmag * Istsf

        pO_Ist = us_st * d_st * f_st

        Ist_Na = IstNainf * pO_Ist
        Ist_K = IstKinf * pO_Ist
        Ist = Ist_Na + Ist_K
    End Sub


    '***** Iha ***
    Public Iha As Double                      'total = IhaK + IhaNa
    Public Iha_K As Double                     'K component of Iha
    Public Iha_Na As Double                    'Na component of Iha
    Public originalPNaha As Double = 0.007284 * 5   '0.1821 * 0.01 * 4             'Amplitude factor for the Na comp. Iha
    Public originalPKha As Double = originalPNaha * 4.244     '0.77286 * 0.01 * 4                    'Amplitude factor for the K comp. Iha

    Public PKha As Double                    'Amplitude factor for the K comp. Iha
    Public PNaha As Double         'Amplitude factor for the Na comp. Iha

    Public alfaf_ha As Double
    Public betaf_ha As Double                    'closing rate
    Public myuf_ha As Double                     'activation rate from C1 to C2
    Public Lamdaf_ha As Single

    Public vshift_ha As Double = -10                  'voltage shift of the Iha kinetics.
    Public Ihamag As Double = 1         '0.45                   'scaling factor of the Iha amplitude
    Public C1_ha As Double              'Iha
    Public C2_ha As Double
    Public O1_ha As Double          'open state1
    Public O2_ha As Double          'open state2
    Public dC1_hadt As Double
    Public dC2_hadt As Double
    Public dO1_hadt As Double
    Public dO2_hadt As Double

    Public pO_ha As Double            '= O1_ha + O2_ha


    Public Sub calculateIha()
        '************* Iha  Kinetics of 5 state transition;  C1 <--> C2 <--> O1 <--> O2 <--> O3 ******************
        Dim v As Double = Vm - vshift_ha
        alfaf_ha = 1 / (3500 * Math.Exp(v / 16.8) + 0.3 * Math.Exp(v / 400))                     'slightly faster than Maruoka 1994
        betaf_ha = 1 / (4 * Math.Exp(v / -14) + 2 * Math.Exp(v / -400))
        myuf_ha = 1 / (45000000 * Math.Exp(v / 8.7) + 500 * Math.Exp(v / 200))
        Lamdaf_ha = 1 / (10.5 * Math.Exp(v / -16.4) + 0.4 * Math.Exp(v / -400))

        dC1_hadt = Lamdaf_ha * C2_ha - myuf_ha * C1_ha                                                         'C1 state of If
        dC2_hadt = myuf_ha * C1_ha + betaf_ha * O1_ha - (Lamdaf_ha + alfaf_ha) * C2_ha       'C2 state of if channel
        dO1_hadt = alfaf_ha * C2_ha + betaf_ha * O2_ha - (betaf_ha + alfaf_ha) * O1_ha            'O1 state
        dO2_hadt = alfaf_ha * O1_ha + betaf_ha * (1 - C1_ha - C2_ha - O1_ha - O2_ha) - (alfaf_ha + betaf_ha) * O2_ha  'O2 state

        '************* Iha  Amplitude  ******************
        pO_ha = 1 - C1_ha - C2_ha                              ' = O1_ha + O2_ha + O3_ha
        Iha_K = originalPKha * Ihamag * GHKK * pO_ha * Ihasf
        Iha_Na = originalPNaha * Ihamag * GHKNa * pO_ha * Ihasf

        Iha = Iha_Na + Iha_K

    End Sub

    'Public Sub calculateIhaSimple()
    '    '************* Iha  Kinetics of 5 state transition;  C1 <--> C2 <--> O1 <--> O2 <--> O3 ******************
    '    'Dim v As Double = Vm - vshift_ha
    '    'alfaf_ha = 1 / (3500 * Math.Exp(v / 16.8) + 0.3 * Math.Exp(v / 400))                                        'slightly faster than Maruoka 1994
    '    'betaf_ha = 1 / (4 * Math.Exp(v / -14) + 2 * Math.Exp(v / -400))
    '    'myuf_ha = 1 / (45000000 * Math.Exp(v / 8.7) + 500 * Math.Exp(v / 200))
    '    'Lamdaf_ha = 1 / (10.5 * Math.Exp(v / -16.4) + 0.4 * Math.Exp(v / -400))

    '    'dC1_hadt = Lamdaf_ha * C2_ha - myuf_ha * C1_ha                                                         'C1 state of If
    '    'dC2_hadt = myuf_ha * C1_ha + betaf_ha * O1_ha - (Lamdaf_ha + alfaf_ha) * C2_ha       'C2 state of if channel
    '    'dO1_hadt = alfaf_ha * C2_ha + betaf_ha * O2_ha - (betaf_ha + alfaf_ha) * O1_ha            'O1 state
    '    'dO2_hadt = alfaf_ha * O1_ha + betaf_ha * (1 - C1_ha - C2_ha - O1_ha - O2_ha) - (alfaf_ha + betaf_ha) * O2_ha  'O2 state

    '    ''************* Iha  Amplitude  ******************
    '    'pO_ha = 1 - C1_ha - C2_ha                              ' = O1_ha + O2_ha + O3_ha

    '    'pO_ha = pOIha(I)                                                   'use the fixed pOIha()

    '    Iha_K = originalPKha * Ihamag * GHKK * pO_ha * Ihasf
    '    Iha_Na = originalPNaha * Ihamag * GHKNa * pO_ha * Ihasf

    '    Iha = Iha_Na + Iha_K

    'End Sub


    '*** Ik1***
    Public IK1 As Single
    Public GIK1 As Double = 0.1                   '0.1

    Public myu_Mg As Double                 'M^-1,msec^-1  single channel data maximum slope
    Public lamda_Mg As Double              '120, 'adjust half block = -50 mV
    Public fracB As Double                       'fraction of Mg-blocked
    Public fracO As Double                       'fraction of Mg-unblocked component
    Public bkspm_K1 As Double
    Public aKspm_K1 As Double

    Public O_IK1 As Double                 'IK1
    Public dO_IK1dt As Double

    Public SSIK1kinetics As Boolean = False
    Public IK1mag As Double = 1            '1.1 * 0.9936                    'revised on 180918 by AN
    Public IKrmag As Double = 1           '1.1 * 1.0176                    'revised on 180918 by AN

    Public t0IKrmag As Double                                        'under the assumption of slow exchange of a test solution in experiments
    Public targetIKrmag As Double                                  'under the assumption of slow exchange of a test solution in experiments


    Public orgGKur As Double = 0.05 * 0.02 * 0.113      '0.000113
    Public GKur As Double

    Public aua As Double
    Public bua As Double
    Public aui As Double
    Public bui As Double
    Public tua As Double
    Public ua As Double
    Public tui As Double
    Public ui As Double
    Public IKur As Double
    Public duadt As Double
    Public duidt As Double
    Public uim As Double
    Public yui As Double
    Public y2ui As Double
    Public ui_o As Double
    Public IKurmag As Double = 1
    Public ua_sift As Double = 9

    Public pO_IKur As Double

    Public Sub calculateIKur()

        'Dim Vm_u As Double = Vm + ua_sift
        aua = 1 / (0.65 * (Math.Exp(-(Vm + 19) / 8.5) + Math.Exp(-(Vm - 21) / 59)))
        bua = 1 / (0.65 * (2.5 + Math.Exp((Vm + 91) / 17)))
        'aua = 1 / (0.65 * (Math.Exp(-(Vm_u + 10) / 8.5) + Math.Exp(-(Vm_u - 30) / 59)))
        'bua = 1 / (0.65 * (2.5 + Math.Exp((Vm_u + 82) / 17)))

        aui = 1 / (21 + Math.Exp(-(Vm - 185) / 28))
        bui = Math.Exp((Vm - 158) / 16)

        duadt = 3 * aua * (1 - ua) - 3 * bua * ua
        duidt = 3 * aui * (1 - ui) - 3 * bui * ui
        pO_IKur = ua ^ 3 * ui

        GKur = orgGKur * IKurmag * IKursf
        IKur = GKur * (1 + 1 / (1 + Math.Exp(-(Vm - 30) / 59))) * pO_IKur * (Vm - EK)
    End Sub


    Public Sub calculateIKurSimple()
        'Dim Vm_u As Double = Vm + ua_sift
        'aua = 1 / (0.65 * (Math.Exp(-(Vm_u + 10) / 8.5) + Math.Exp(-(Vm_u - 30) / 59)))
        'bua = 1 / (0.65 * (2.5 + Math.Exp((Vm_u + 82) / 17)))

        'aui = 1 / (21 + Math.Exp(-(Vm - 185) / 28))
        'bui = Math.Exp((Vm - 158) / 16)

        'duadt = 3 * aua * (1 - ua) - 3 * bua * ua
        'duidt = 3 * aui * (1 - ui) - 3 * bui * ui

        'pO_ur = ua ^ 3 * ui                                         'pOpen of IKur
        duadt = 0
        duidt = 0

        GKur = orgGKur * IKurmag * IKursf
        IKur = GKur * (1 + 1 / (1 + Math.Exp(-(Vm - 30) / 59))) * pO_IKur * (Vm - EK)
    End Sub


    Public oGK1 As Double = 1.07565 * 0.42     '0.451773    nS, conductance of IK1
    Public GK1 As Double = oGK1      'nS, default value of conductance of IK1
    Public IK1_K_cyt As Double
    Public Pbspm As Double
    Public ssPbspm As Double        'steady state probability of spm block   a optional variable
    Public dPbspmdt As Double
    Public pO_IK1 As Double
    Public pO_mode1 As Double       'pOpen of mode1
    Public pO_mode2 As Double       'pOpen of mode2

    Public fK_IK1 As Double
    Public poMg1 As Double
    Public poMg2 As Double
    Public po_Mggate As Double
    Public pOpen_Mggate As Double       'pOpen of the whole Mg gate

    Public alphaSPM As Double
    Public betaSPM As Double
    Public alphaMg As Double
    Public betaMg As Double

    Public LimitGK1 As Double                 'instantaneous limitting GK1

    Public SPM As Double = 5                              'uM, default concentration of spermine
    Public frac_mode1 As Double = 0.9

    Public Sub calculateIK1(ByVal EK As Double, ByVal Pbspm As Double, ByRef dPbspmdt As Double)
        '
        'Dim SPM As Double = 5              '0.005 * 1000                                 ' 
        '************** instantaneous Mg-block *************************
        alphaMg = 12.0 * Exp(-0.025 * (Vm - EK))                                            ' 
        betaMg = 28 * Mg_cyt * Exp(0.025 * (Vm - EK))                                 ' 
        Dim fracO As Double = alphaMg / (alphaMg + betaMg)                  ' 
        Dim fracB As Double = betaMg / (alphaMg + betaMg)                    ' 
        poMg2 = 3.0 * fracO * fracB * fracB                                                  ' 
        poMg1 = 3.0 * fracO * fracO * fracB                                                  '
        po_Mggate = fracO * fracO * fracO
        pOpen_Mggate = po_Mggate + 2 / 3 * poMg1 + 1 / 3 * poMg2          'net Mg_block

        '************** spermin-block *************************
        alphaSPM = 0.17 * Exp(-0.07 * ((Vm - EK) + 8 * Mg_cyt)) / (1.0 + 0.01 * Exp(0.12 * ((Vm - EK) + 8 * Mg_cyt)))               '
        betaSPM = 0.28 * SPM * Exp(0.15 * ((Vm - EK) + 8 * Mg_cyt)) / (1.0 + 0.01 * Exp(0.13 * ((Vm - EK) + 8 * Mg_cyt)))      '
        dPbspmdt = betaSPM * po_Mggate * (1 - Pbspm) - alphaSPM * Pbspm                                                                               '

        ssPbspm = (betaSPM * po_Mggate) / (alphaSPM + betaSPM * po_Mggate)
        '**************** IK1 amplitude **************************************************
        'If SSIK1kinetics = True Then pO_mode1 = frac_mode1 * (1 - ssPbspm) * pOpen_Mggate '
        fK_IK1 = (Ko / 5.4) ^ 0.4 / (1.0 + Exp(-(Ko - 2.2) / 0.6))          'dependency on Kout
        'fK_IK1 = 1.07565 * (Ko / 5.4) ^ 0.4 / (1.0 + Exp(-(Ko - 2.2) / 0.6))          'dependency on Kout

        LimitGK1 = GK1 * fK_IK1 * IK1mag * IK1sf

        pO_mode2 = (1.0 - frac_mode1) / (1.0 + SPM / (40.0 * Exp(-(Vm - EK) / 9.1)))          'instantaneous mode2                                                                            '
        pO_mode1 = frac_mode1 * (1 - Pbspm) * pOpen_Mggate                                                         '

        pO_IK1 = (pO_mode1 + pO_mode2)        'Two populations of IK1 channel in respect to the spermine and Mg block

        IK1_K_cyt = LimitGK1 * (Vm - EK) * pO_IK1                   '

    End Sub

    Public Sub calculateIK1Simple()
        '************** instantaneous Mg-block *************************
        'alphaMg = 12.0 * Exp(-0.025 * (v - EK))                                            ' 
        'betaMg = 28 * Mg_cyt * Exp(0.025 * (v - EK))                                 ' 
        'Dim fracO As Double = alphaMg / (alphaMg + betaMg)                  ' 
        'Dim fracB As Double = betaMg / (alphaMg + betaMg)                    ' 
        'poMg2 = 3.0 * fracO * fracB * fracB                                                  ' 
        'poMg1 = 3.0 * fracO * fracO * fracB                                                  '
        'po_Mggate = fracO * fracO * fracO
        'pOpen_Mggate = po_Mggate + 2 / 3 * poMg1 + 1 / 3 * poMg2          'net Mg_block

        ''************** spermin-block *************************
        'alphaSPM = 0.17 * Exp(-0.07 * ((v - EK) + 8 * Mg_cyt)) / (1.0 + 0.01 * Exp(0.12 * ((v - EK) + 8 * Mg_cyt)))               '
        'betaSPM = 0.28 * SPM * Exp(0.15 * ((v - EK) + 8 * Mg_cyt)) / (1.0 + 0.01 * Exp(0.13 * ((v - EK) + 8 * Mg_cyt)))      '
        'dPbspmdt = betaSPM * po_Mggate * (1 - Pbspm) - alphaSPM * Pbspm                                                                               '

        'ssPbspm = (betaSPM * po_Mggate) / (alphaSPM + betaSPM * po_Mggate)
        '**************** IK1 amplitude **************************************************
        'If SSIK1kinetics = True Then pO_mode1 = frac_mode1 * (1 - ssPbspm) * pOpen_Mggate '

        'pO_mode2 = (1.0 - frac_mode1) / (1.0 + SPM / (40.0 * Exp(-(v - EK) / 9.1)))          'instantaneous mode2                                                                            '
        'pO_mode1 = frac_mode1 * (1 - Pbspm) * pOpen_Mggate                                                         '

        'pO_IK1 = pOIK1(I)      'Two populations of IK1 channel in respect to the spermine and Mg block

        dPbspmdt = 0

        fK_IK1 = 1.07565 * (Ko / 5.4) ^ 0.4 / (1.0 + Exp(-(Ko - 2.2) / 0.6))          'dependency on Kout
        LimitGK1 = GK1 * fK_IK1 * IK1mag * IK1sf

        IK1_K_cyt = LimitGK1 * (Vm - EK) * pO_IK1                   '

    End Sub

    Public Sub calculateIK1ss(ByVal v As Double, ByVal EK As Double, ByVal Pbspm As Double)
        Dim frac_mode1 As Double = 0.9                                     '
        'Dim SPM As Double = 5          '0.005 * 1000                                 ' 
        '************** Mg-block *************************
        alphaMg = 12.0 * Exp(-0.025 * (v - EK))                                            ' 
        betaMg = 28 * Mg_cyt * Exp(0.025 * (v - EK))                                 ' 
        Dim fracO As Double = alphaMg / (alphaMg + betaMg)                  ' 
        Dim fracB As Double = betaMg / (alphaMg + betaMg)                    ' 
        poMg2 = 3.0 * fracO * fracB * fracB                                                  ' 
        poMg1 = 3.0 * fracO * fracO * fracB                                                  '
        po_Mggate = fracO * fracO * fracO                                                   '
        '************** spermin-block *************************
        alphaSPM = 0.17 * Exp(-0.07 * ((v - EK) + 8 * Mg_cyt)) / (1.0 + 0.01 * Exp(0.12 * ((v - EK) + 8 * Mg_cyt)))               '
        betaSPM = 0.28 * SPM * Exp(0.15 * ((v - EK) + 8 * Mg_cyt)) / (1.0 + 0.01 * Exp(0.13 * ((v - EK) + 8 * Mg_cyt)))      '
        dPbspmdt = betaSPM * po_Mggate * (1 - Pbspm) - alphaSPM * Pbspm                                                                               '

        ssPbspm = (betaSPM * po_Mggate) / (alphaSPM + betaSPM * po_Mggate)
        '**************** IK1 amplitude **************************************************
        pO_mode1 = frac_mode1 * (1 - ssPbspm) * (po_Mggate + 2 / 3 * poMg1 + 1 / 3 * poMg2) '
        pO_mode2 = (1.0 - frac_mode1) / (1.0 + SPM / (40.0 * Exp(-(v - EK) / 9.1)))                                                                               '

        pO_IK1 = (pO_mode1 + pO_mode2)                                        '

        fK_IK1 = 1.07565375693257 * (Ko / 5.4) ^ 0.4 / (1.0 + Exp(-(Ko - 2.2) / 0.6))          '

        IK1_K_cyt = GK1 * IK1mag * (v - EK) * fK_IK1 * pO_IK1                   '

    End Sub

    '********** IKsKyoto **********
    Dim b2_IKs As Double = 0.000296 '* fixzero
    Dim a3_IKs As Double = 0.0003 '* fixzero
    Dim b3_IKs As Double = 0.03 '* fixzero

    Public IKs As Double
    Public IKs_iz As Double
    Public IKs_blk As Double

    Public IKs_K_iz As Double
    Public IKs_Na_iz As Double
    Public IKs_K_blk As Double
    Public IKs_Na_blk As Double

    Public pO_IKs_iz As Double
    Public pO_IKs_blk As Double

    Public Ov_IKs As Double                'voltage gate with  two states , common for both iz and blk compartments, 
    Public dOv_IKsdt As Double

    Public C2IKs_iz As Double             'iz compartment, Calcium gate with C1 - C2 - Oc states
    Public OcIKs_iz As Double             'iz compartment, Calcium gate with C1 - C2 - Oc states
    Public dOcIKs_izdt As Double        'iz compartment, Calcium gate with C1 - C2 - Oc states
    Public dC2IKs_izdt As Double        'iz compartment, Calcium gate with C1 - C2 - Oc states

    Public C2IKs_blk As Double              'blk compartment, Calcium gate with C1 - C2 - Oc states
    Public OcIKs_blk As Double              'blk compartment, Calcium gate with C1 - C2 - Oc states
    Public dOcIKs_blkdt As Double        'blk compartment, Calcium gate with C1 - C2 - Oc states
    Public dC2IKs_blkdt As Double        'blk compartment, Calcium gate with C1 - C2 - Oc states

    Public PKs As Double = 0.4                               '0.002782
    Public perNa_IKs As Double = 0.04 * PKs

    Public a1_IKs As Double
    Public b1_IKs As Double
    Public a1_IKso As Double
    Public b1_IKso As Double
    Public vsift_IKs As Double = -10                       '-5   '-5   -10  -15

    Public IKs_K As Double
    Public IKs_Na As Double
    Public IKsmag As Double = 1          '1 * 1.0229                           'revised by AN on 180918

    Public Sub VgateIKs(ByVal Ov As Double, ByRef dOvdt As Double)
        a1_IKs = 1.0 / (150.0 * Exp(-(Vm - vsift_IKs) / 25.0) + 900.0 * Exp(-(Vm - vsift_IKs) / 200.0))
        b1_IKs = 1.0 / (1000.0 * Exp((Vm - vsift_IKs) / 13.0) + 220.0 * Exp((Vm - vsift_IKs) / 50.0))
        dOvdt = a1_IKs * (1.0 - Ov) - b1_IKs * Ov                          'rate of change of the Vm-dependent gate pOpen
    End Sub

    Private Sub calculateIKs(ByVal ca As Double, ByVal fraction As Double, ByVal Ov_IKs As Double, ByVal C2_IKs As Double, ByVal Oc_IKs As Double, ByRef dOc_IKsdt As Double, ByRef dC2_IKsdt As Double, ByRef pO_IKs As Double)
        Dim C1_IKs As Double = 1 - C2_IKs - Oc_IKs
        Dim a2_IKs As Double = ca * 2.25                                     '* fixzero

        dOc_IKsdt = a2_IKs * C2_IKs - b2_IKs * Oc_IKs                                                                 'Calcium gate
        dC2_IKsdt = a3_IKs * C1_IKs - b3_IKs * C2_IKs - a2_IKs * C2_IKs + b2_IKs * Oc_IKs     'Calcium gate

        pO_IKs = (Ov_IKs ^ 2) * (0.99 * Oc_IKs + 0.01)           'open probability of the IKs_gate

        IKs_Na = fraction * perNa_IKs * GHKNa * pO_IKs * IKsmag * IKssf
        IKs_K = fraction * PKs * GHKK * pO_IKs * IKsmag * IKssf

    End Sub


    'Private Sub calculateIKsSimple(ByVal ca As Double, ByVal fraction As Double, ByVal Ov_IKs As Double, ByVal C2_IKs As Double, ByVal Oc_IKs As Double, ByRef dOc_IKsdt As Double, ByRef dC2_IKsdt As Double, ByRef pO_IKs As Double)
    '    dOv_IKsdt = 0.0                  'fix the Vm-dependent gate

    '    Dim C1_IKs As Double = 1 - C2_IKs - Oc_IKs
    '    Dim a2_IKs As Double = ca * 2.25                                     '* fixzero

    '    dOc_IKsdt = a2_IKs * C2_IKs - b2_IKs * Oc_IKs                                                                 'Calcium gate
    '    dC2_IKsdt = a3_IKs * C1_IKs - b3_IKs * C2_IKs - a2_IKs * C2_IKs + b2_IKs * Oc_IKs     'Calcium gate

    '    pO_IKs = (Ov_IKs ^ 2) * (0.99 * Oc_IKs + 0.01)           'open probability of the IKs_gate

    '    IKs_Na = fraction * perNa_IKs * GHKNa * pO_IKs * IKsmag * IKssf
    '    IKs_K = fraction * PKs * GHKK * pO_IKs * IKsmag * IKssf

    'End Sub

    '********** IKto ***********
    Public IKto As Double
    Public IKto_Na_cyt As Double
    Public IKto_K_cyt As Double
    Public IKto_K As Double
    Public IKto_Na As Double

    Public y1_IKto As Double     'fast inactivation parameter  (57)
    Public y2_IKto As Double     'slow inactivation parameter (58)
    Public dy1_IKtodt As Double
    Public dy2_IKtodt As Double
    Public a1_IKto As Double
    Public b1_IKto As Double
    Public a2_IKto As Double
    Public b2_IKto As Double

    Public pO_IKto As Double

    Public PKtoEndo As Double = 0.0133 * 1.3           '0.01729
    Public pNa_IKto As Double = PKtoEndo * 0.09

    Public IKtomag As Double = 1
    'Public IKto_orgmag As Double = 1.3


    Public Sub calculateIKtoEndo(ByRef IKto_K As Double, ByRef IKto_Na As Double, ByRef y1_IKto As Double, ByRef y2_IKto As Double, ByRef dy1_IKtodt As Double, ByRef dy2_IKtodt As Double)

        a1_IKto = 1.0 / (13 * Exp(-Vm / 22.0))   'activation gate
        b1_IKto = 1.0 / (2.1 * Exp(Vm / 90.0))   'activation gate

        a2_IKto = 1.0 * 0.5 / (950 * Exp(Vm / 500)) '* 0.6                                                'inactivation gate   modified on 201204
        b2_IKto = 1.0 * 0.5 / (40.0 * Exp(-Vm / 9.0) + 13.0 * Exp(-Vm / 1000.0)) '* 0.6    'inactivation gate   modified on 201204

        dy1_IKtodt = (a1_IKto * (1.0 - y1_IKto) - b1_IKto * y1_IKto)
        dy2_IKtodt = (a2_IKto * (1.0 - y2_IKto) - b2_IKto * y2_IKto)

        pO_IKto = y1_IKto * y2_IKto                                       'version of a complete inactivation

        '*************** Current Amplitude ***************************************
        IKto_Na = pNa_IKto * GHKNa * IKtomag * pO_IKto * IKtosf
        IKto_K = PKtoEndo * GHKK * IKtomag * pO_IKto * IKtosf

    End Sub


    'Public Sub calculateIKtoSimple()
    '    'a1_IKto = 1.0 / (13 * Exp(-v / 22.0))   'activation gate
    '    'b1_IKto = 1.0 / (2.1 * Exp(v / 90.0))   'activation gate

    '    'a2_IKto = 1.0 / (950 * Exp(v / 500)) '* 0.6                                                'inactivation gate
    '    'b2_IKto = 1.0 / (40.0 * Exp(-v / 9.0) + 13.0 * Exp(-v / 1000.0)) '* 0.6    'inactivation gate

    '    'dy1_IKtodt = (a1_IKto * (1.0 - y1_IKto) - b1_IKto * y1_IKto)
    '    'dy2_IKtodt = (a2_IKto * (1.0 - y2_IKto) - b2_IKto * y2_IKto)

    '    'pO_IKto = y1_IKto * y2_IKto                                       'version of a complete inactivation

    '    'pO_IKto = pOIKto(I)

    '    dy1_IKtodt = 0
    '    dy2_IKtodt = 0
    '    '************************************** Current Amplitude ***************************************
    '    IKto_Na = pNa_IKto * GHKNa * pO_IKto * IKtomag * IKtosf
    '    IKto_K = PKtoEndo * GHKK * pO_IKto * IKtomag * IKtosf


    'End Sub
    '****** IKACh ********
    Public IKACh As Double
    Public ACh As Single = 0
    Public newACh As Single = 0
    Public GKACh As Double = 0.135 * 1.5               'current density

    Public O_ACh As Double
    Public dO_AChdt As Double
    Public alfa_ACh As Double
    Public beta_ACh As Double

    Public Sub calculateIKACh()
        '************* ACH CURRENT  *****************
        If ACh = 0 Then
            alfa_ACh = 0.0002475                             'spontaneous opening   /msec"
        Else
            alfa_ACh = 0.01232 / (1 + 0.003 / ACh) + 0.0002
        End If
        beta_ACh = 0.01 * Math.Exp(0.0133 * (Vm + 40))           'beta of IKACh

        dO_AChdt = alfa_ACh * (1 - O_ACh) - beta_ACh * (O_ACh)

        IKACh = GKACh * (Vm - EK) / (1 + Math.Exp((Vm + 20) / 20)) * O_ACh

    End Sub

    '''''***************************************** IKr
    Public IKr As Double
    Public IKr_K_cyt As Double

    Public y1_IKr As Double
    Public y2_IKr As Double
    Public y3_IKr As Double
    Public dy1_IKrdt As Double
    Public dy2_IKrdt As Double
    Public dy3_IKrdt As Double
    Public pO_IKr As Double
    Public pOv_IKs As Double

    Public a1_IKr As Double
    Public b1_IKr As Double
    Public a2_IKr As Double
    Public b2_IKr As Double
    Public a3_IKr As Double
    Public b3_IKr As Double


    'Public scaleIKr As Double = 1                         'scaling factor for IKr amplitude, defalt = 1
    Public oGIKr As Double = 0.03546 * 1.4           '0.049644     'nS, 
    Public GIKr As Double = oGIKr
    Public vsift_IKr2 As Double = 5
    Public vsift_IKr1 As Double = 6                'mV,   voltage shift
    Public limitGIKr As Double                         'pA/pF, fully activated IKr

    'Public pOIKr(50000) As Double

    Public Sub calculateIKr()
        'the fast component of activation
        a1_IKr = 1.0 / (20.0 * Exp(-(Vm + vsift_IKr1) / 6) + 5.0 * Exp(-(Vm + vsift_IKr1) / 150.0))
        b1_IKr = 1.0 / (160.0 * Exp((Vm + vsift_IKr1) / 28) + 200.0 * Exp((Vm + vsift_IKr1) / 1000.0)) + 1.0 / (2500.0 * Exp((Vm + vsift_IKr1) / 20))
        'the slow component of activation
        a2_IKr = 1.0 / (200.0 * Exp(-(Vm + vsift_IKr1) / 6.5) + 20.0 * Exp(-(Vm + vsift_IKr1) / 150.0))            '* fixzero
        b2_IKr = 1.0 / (1600.0 * Exp((Vm + vsift_IKr1) / 28) + 2000.0 * Exp((Vm + vsift_IKr1) / 1000.0)) + 1.0 / (10000.0 * Exp((Vm + vsift_IKr1) / 20))              '* fixzero

        ' inactivation gate (fast gate)
        a3_IKr = 1.0 / (10 * Exp((Vm + vsift_IKr1) / 17.0) + 2.5 * Exp((Vm + vsift_IKr1) / 300.0))                      '* fixzero
        b3_IKr = 1.0 / (0.35 * Exp(-(Vm + vsift_IKr1) / 17) + 1.0 * Exp(-(Vm + vsift_IKr1) / 75.0))                '* fixzero

        dy1_IKrdt = a1_IKr * (1.0 - y1_IKr) - b1_IKr * y1_IKr                                 'differential equation of y1
        dy2_IKrdt = (a2_IKr * (1.0 - y2_IKr) - b2_IKr * y2_IKr)                               'differential equation of y2
        dy3_IKrdt = a3_IKr * (1.0 - y3_IKr) - b3_IKr * y3_IKr                                 'differential equation of y3

        pO_IKr = (0.6 * y1_IKr + 0.4 * y2_IKr) * y3_IKr                                      'determine pOpen from the previous step

        limitGIKr = GIKr * (Ko / 5.4) ^ 0.2 * IKrmag * IKrsf
        IKr_K_cyt = limitGIKr * (Vm - EK) * pO_IKr
    End Sub

    'Public Sub calculateIKrSimple()
    '    ''the fast component of activation
    '    'a1_IKr = 1.0 / (20.0 * Exp(-(v + vsift_IKr1) / 6) + 5.0 * Exp(-(v + vsift_IKr1) / 150.0))
    '    'b1_IKr = 1.0 / (160.0 * Exp((v + vsift_IKr1) / 28) + 200.0 * Exp((v + vsift_IKr1) / 1000.0)) + 1.0 / (2500.0 * Exp((v + vsift_IKr1) / 20))
    '    ''the slow component of activation
    '    'a2_IKr = 1.0 / (200.0 * Exp(-(v + vsift_IKr1) / 6.5) + 20.0 * Exp(-(v + vsift_IKr1) / 150.0))            '* fixzero
    '    'b2_IKr = 1.0 / (1600.0 * Exp((v + vsift_IKr1) / 28) + 2000.0 * Exp((v + vsift_IKr1) / 1000.0)) + 1.0 / (10000.0 * Exp((v + vsift_IKr1) / 20))              '* fixzero

    '    '' inactivation gate (fast gate)
    '    'a3_IKr = 1.0 / (10 * Exp((v + vsift_IKr1) / 17.0) + 2.5 * Exp((v + vsift_IKr1) / 300.0))                      '* fixzero
    '    'b3_IKr = 1.0 / (0.35 * Exp(-(v + vsift_IKr1) / 17) + 1.0 * Exp(-(v + vsift_IKr1) / 75.0))                '* fixzero

    '    'dy1_IKrdt = a1_IKr * (1.0 - y1_IKr) - b1_IKr * y1_IKr                                 'differential equation of y1
    '    'dy2_IKrdt = (a2_IKr * (1.0 - y2_IKr) - b2_IKr * y2_IKr)                               'differential equation of y2
    '    'dy3_IKrdt = a3_IKr * (1.0 - y3_IKr) - b3_IKr * y3_IKr                                 'differential equation of y3

    '    dy1_IKrdt = 0.0                                'differential equation of y1
    '    dy2_IKrdt = 0.0                               'differential equation of y2
    '    dy3_IKrdt = 0.0                               'differential equation of y3

    '    'pO_IKr = (0.6 * y1_IKr + 0.4 * y2_IKr) * y3_IKr                                      'determine pOpen from the previous step

    '    'pO_IKr = pOIKr(I)

    '    limitGIKr = GIKr * (Ko / 5.4) ^ 0.2 * IKrmag * IKrsf
    '    IKr_K_cyt = limitGIKr * (Vm - EK) * pO_IKr

    'End Sub


    '*** IKATP ***
    Public IKATP As Double                 '##################   IKATP    ###################
    Public N_KATP As Double = 600 / Cm             'Current density normalized by the cell capacitance Cm    ref. Kakei et al 1985
    Public rKATP As Double                    'single channel conductance (pS)
    Public PoK_ATP As Double = 0.8        'maximum open probability of KATP channel
    Public GKATP As Double                  'net conductance 

    Public Sub calculateIKATP()
        '*********************** KATP currents ******************************
        rKATP = 23.6 * Ko ^ 0.24 / 1000                'conductance of a single ATP sensitive K channel
        GKATP = rKATP * N_KATP                       '  
        IKATP = GKATP * (Vm - EK) / (1 + (MgATP_cyt / 0.1) ^ 2)
    End Sub

    '''***************************************' INa   *****************************************
    Public INa As Double                          'pA, current amplitude at the time of observation
    Public sclI1I2 As Double = 1                                             'scaling factor for the slow inactivation rate of INaL
    '********* inward current amplitudes **************  current density **************

    Public INaT As Double                         'pA, current amplitude
    Public INaL As Double                         'pA, current amplitude
    Public INaLratio As Double = 0.08       '0.131    '0.175 * 0.75        '0.9  0.75
    Public PNa As Double = 32.43 * 2.275           '73.77825                 '
    Public relativePK_INa As Double = 0.18

    Public INaT_Na_cyt As Double             'pA, amplitude of Na component
    Public INaT_K_cyt As Double               'pA, amplitude of K component
    Public INaL_Na_cyt As Double             'pA, amplitude of Na component
    Public INaL_K_cyt As Double               'pA, amplitude of K component

    Public fC_INa As Double

    Public pO_INaT As Double
    Public pO_INaL As Double
    Public C_TM As Double             'transient mode of INa       pC_TM
    Public O_TM As Double             'transient mode of INa       pO_TM
    Public I2_TM As Double            'transient mode of INa       pI2_TM
    Public Is_TM As Double             'transient mode of INa       pIs_TM

    'Public us
    Public dO_TMdt As Double             'transient mode of INa   dpO_TMdt
    Public dI2_TMdt As Double             'transient mode of INa   dpI2_TMdt
    Public dIs_TMdt As Double             'transient mode of INa   dpIs_TMdt

    Public C_LM As Double             'LSM mode of INa
    Public O_LM As Double             'LSM mode of INa
    Public I1_LM As Double             'LSM mode of INa
    Public I2_LM As Double             'LSM mode of INa
    Public Is_LM As Double             'LSM mode of INa
    '   Public Iinact_LM As Double        'Is_LSM  used in calculating the steady state I-V

    Public fixedIs_LM As Double

    Public dO_LMdt As Double
    Public dI1_LMdt As Double
    Public dI2_LMdt As Double
    Public dIs_LMdt As Double

    Public kC2O_INa, kOC_INa, kOI2_TM, kI2O_TM, kC2I2_TM, kI2C_TM, kIsb_TM, kIsf_TM As Double
    Public kI1I2_LM, kI2I1_LM, kOI1_LM, kI1O_LM, kI1C_LM, kC2I1_LM, kC2I2_LM, kI2C_LM, kIsb_LM, kIsf_LM As Double
    Public fixzero_INaLslowgate As Double = 1

    Public orgVshiftNa As Double = 4
    Public VshiftNa As Double
    Public VshiftNasf As Double = 1


    Public d12_TM As Double
    Public d23_TM As Double
    Public d31_TM As Double
    Public d34_TM As Double
    Public d41_TM As Double

    Public d12_LM As Double
    Public d23_LM As Double
    Public d31_LM As Double
    Public d34_LM As Double
    Public d41_LM As Double
    Public d45_LM As Double
    Public d51_LM As Double

    Public Sub calculateINarate()
        '************************************ pure Voltage-dependent gating rate from the HuVEC model ******************************************************* 
        VshiftNa = orgVshiftNa * VshiftNasf

        Dim v As Double = Vm - VshiftNa     'applied only to the activation rates

        kC2O_INa = 1.0 / (0.0025 * Math.Exp(-v / 8.0) + 0.15 * Math.Exp(-v / 100.0))           '
        kOC_INa = 1.0 / (30.0 * Math.Exp(v / 12.0) + 0.53 * Math.Exp(v / 50.0))                    '

        fC_INa = 1 / (1 + Math.Exp(-(v + 48) / 7))                               '

        v = Vm                                                  'applied to the other rates
        kOI2_TM = 1.0 / (0.0433 * Math.Exp(-v / 27) + 0.34 * Math.Exp(-v / 2000.0))           '
        kI2O_TM = 0.0001312                                                                                  '

        kC2I2_TM = 0.5 / (1.0 + kI2O_TM * kOC_INa / kOI2_TM / kC2O_INa)           '
        kI2C_TM = 0.5 - kC2I2_TM                                                                                      '

        kIsb_TM = 1.0 / (300000 * Math.Exp(v / 10) + 50000.0 * Math.Exp(v / 16.0))             '     
        kIsf_TM = 1.0 / (0.016 * Math.Exp(-v / 9.9) + 8 * Math.Exp(-v / 45.0))                          '

        '*******************  Calculate state transitions of TM ***************************************************************
        C_TM = 1.0 - Is_TM - O_TM - I2_TM                          '' 


        'dO_TMdt = I2_TM * kI2O_TM + fC_INa * C_TM * kC2O_INa - pO_TM * (kOC_INa + kOI2_TM)                                                             '
        'dI2_TMdt = fC_INa * C_TM * kC2I2_TM + pO_TM * kOI2_TM + Is_TM * kIsb_TM - I2_TM * (kI2C_TM + kI2O_TM + kIsf_TM)    '
        'dIs_TMdt = I2_TM * kIsf_TM + C_TM * kIsf_TM - Is_TM * 2 * kIsb_TM
        '
        deltaprobability(d12_TM, C_TM, O_TM, fC_INa * kC2O_INa, kOC_INa, dt)
        deltaprobability(d23_TM, O_TM, I2_TM, kOI2_TM, kI2O_TM, dt)
        deltaprobability(d31_TM, I2_TM, C_TM, kI2C_TM, fC_INa * kC2I2_TM, dt)
        deltaprobability(d34_TM, I2_TM, Is_TM, kIsf_TM, kIsb_TM, dt)
        deltaprobability(d41_TM, Is_TM, C_TM, kIsb_TM, kIsf_TM, dt)

        '****************  determine the gating of LSM *********** using common C1-C2-O gating with the transient mode ****************************
        kI1I2_LM = 0.00534 * sclI1I2 * fixzero_INaLslowgate                                   '
        kI2I1_LM = kI1I2_LM / 1000
        kOI1_LM = kOI2_TM                                                                                '
        kI1O_LM = 0.01                                                                                         '
        kI1C_LM = kI2C_TM                                                                               '
        kC2I1_LM = kC2I2_TM                                                                          '

        kC2I2_LM = kC2I2_TM * fixzero_INaLslowgate                 '
        kI2C_LM = kI2C_TM * fixzero_INaLslowgate                      '

        kIsb_LM = 1.0 / (300000 * Exp(v / 10) + 50000.0 * Exp(v / 16.0)) * fixzero_INaLslowgate   ' set #1    
        kIsf_LM = 1.0 / (0.016 * Exp(-v / 9.9) + 8 * Exp(-v / 45)) * fixzero_INaLslowgate

        '****************  Calculate state transitions of LSM ***************************************
        C_LM = 1.0 - Is_LM - O_LM - I1_LM - I2_LM

        'dO_LMdt = I1_LM * kI1O_LM + C_LM * fC_INa * kC2O_INa - pO_LM * (kOC_INa + kOI1_LM)                                        '
        'dI1_LMdt = pO_LM * kOI1_LM + C_LM * fC_INa * kC2I1_LM - I1_LM * (kI1O_LM + kI1C_LM + kI1I2_LM)                   '
        'dI2_LMdt = C_LM * fC_INa * kC2I2_LM + I1_LM * kI1I2_LM + Is_LM * kIsb_LM - I2_LM * (kI2C_LM + kIsf_LM)    '
        'dIs_LMdt = I2_LM * kIsf_LM + C_LM * kIsf_LM - Is_LM * 2 * kIsb_LM '

        deltaprobability(d12_LM, C_LM, O_LM, fC_INa * kC2O_INa, kOC_INa, dt)
        deltaprobability(d23_LM, O_LM, I1_LM, kOI1_LM, kI1O_LM, dt)
        deltaprobability(d31_LM, I1_LM, C_LM, kI1C_LM, fC_INa * kC2I1_LM, dt)
        deltaprobability(d34_LM, I1_LM, I2_LM, kI1I2_LM, kI2I1_LM, dt)
        deltaprobability(d41_LM, I2_LM, C_LM, kI2C_LM, fC_INa * kC2I2_LM, dt)
        deltaprobability(d45_LM, I2_LM, Is_LM, kIsf_LM, kIsb_LM, dt)
        deltaprobability(d51_LM, Is_LM, C_LM, kIsb_LM, kIsf_LM, dt)

    End Sub


    'Public Sub calculateINarate()
    '    '************************************ pure Voltage-dependent gating rate ******************************************************* 
    '    VshiftNa = orgVshiftNamag * vshiftNasf

    '    Dim v As Double = Vm - VshiftNa
    '    kC2O_INa = 1 / (0.0025 * Math.Exp(-v / 8) + 0.15 * Math.Exp(-v / 100))   'according to the HuVEC model
    '    'kC2O_INa = 1  / (0.0025 * Exp(-v / 6.5) + 0.15 * Exp(-v / 100))           '
    '    kOC_INa = 1 / (30 * Exp(v / 12.0) + 0.53 * Exp(v / 50))                    'the same as the HuVEC model

    '    kOI2_TM = 1 / (0.0433 * Exp(-v / 27) + 0.34 * Exp(-v / 2000))           'the same as the HuVEC model
    '    kI2O_TM = 0.0001312                                                                                  '

    '    kC2I2_TM = 0.5 / (1.0 + kI2O_TM * kOC_INa / kOI2_TM / kC2O_INa)           'the same as the HuVEC model
    '    kI2C_TM = 0.5 - kC2I2_TM                                                                                      '

    '    kIsb_TM = 1 / (300000 * Exp(v / 10) + 50000 * Exp(v / 16))             ' the same as the HuVEC model    
    '    kIsf_TM = 1 / (0.016 * Exp(-v / 9.9) + 8 * Exp(-v / 45))                          '

    '    'fC_INa = 1 / (1 + Exp(-(v + 48 - 20) / 7))                               '
    '    'fC = 1 / (1 + Math.Exp((v + 48) / -8))
    '    '*******************  Calculate state transitions of TM ***************************************************************
    '    C_TM = 1.0 - Is_TM - O_TM - I2_TM                          '' 


    '    'dO_TMdt = I2_TM * kI2O_TM + fC_INa * C_TM * kC2O_INa - pO_TM * (kOC_INa + kOI2_TM)                                                             '
    '    'dI2_TMdt = fC_INa * C_TM * kC2I2_TM + pO_TM * kOI2_TM + Is_TM * kIsb_TM - I2_TM * (kI2C_TM + kI2O_TM + kIsf_TM)    '
    '    'dIs_TMdt = I2_TM * kIsf_TM + C_TM * kIsf_TM - Is_TM * 2 * kIsb_TM
    '    '
    '    deltaprobability(d12_TM, C_TM, O_TM, fC_INa * kC2O_INa, kOC_INa, dt)
    '    deltaprobability(d23_TM, O_TM, I2_TM, kOI2_TM, kI2O_TM, dt)
    '    deltaprobability(d31_TM, I2_TM, C_TM, kI2C_TM, fC_INa * kC2I2_TM, dt)
    '    deltaprobability(d34_TM, I2_TM, Is_TM, kIsf_TM, kIsb_TM, dt)
    '    deltaprobability(d41_TM, Is_TM, C_TM, kIsb_TM, kIsf_TM, dt)

    '    '****************  determine the gating of LSM *********** using common C1-C2-O gating with the transient mode ****************************
    '    kI1I2_LM = 0.00534 * sclI1I2 * fixzero_INaLslowgate                                   '
    '    kI2I1_LM = kI1I2_LM / 1000
    '    kOI1_LM = kOI2_TM                                                                                '
    '    kI1O_LM = 0.01                                                                                         '
    '    kI1C_LM = kI2C_TM                                                                               '
    '    kC2I1_LM = kC2I2_TM                                                                          '

    '    kC2I2_LM = kC2I2_TM * fixzero_INaLslowgate                 '
    '    kI2C_LM = kI2C_TM * fixzero_INaLslowgate                      '

    '    kIsb_LM = 1.0 / (300000 * Exp(v / 10) + 50000.0 * Exp(v / 16.0)) * fixzero_INaLslowgate   ' set #1    
    '    kIsf_LM = 1.0 / (0.016 * Exp(-v / 9.9) + 8 * Exp(-v / 45)) * fixzero_INaLslowgate

    '    '****************  Calculate state transitions of LSM ***************************************
    '    C_LM = 1.0 - Is_LM - O_LM - I1_LM - I2_LM

    '    'dO_LMdt = I1_LM * kI1O_LM + C_LM * fC_INa * kC2O_INa - pO_LM * (kOC_INa + kOI1_LM)                                        '
    '    'dI1_LMdt = pO_LM * kOI1_LM + C_LM * fC_INa * kC2I1_LM - I1_LM * (kI1O_LM + kI1C_LM + kI1I2_LM)                   '
    '    'dI2_LMdt = C_LM * fC_INa * kC2I2_LM + I1_LM * kI1I2_LM + Is_LM * kIsb_LM - I2_LM * (kI2C_LM + kIsf_LM)    '
    '    'dIs_LMdt = I2_LM * kIsf_LM + C_LM * kIsf_LM - Is_LM * 2 * kIsb_LM '

    '    deltaprobability(d12_LM, C_LM, O_LM, fC_INa * kC2O_INa, kOC_INa, dt)
    '    deltaprobability(d23_LM, O_LM, I1_LM, kOI1_LM, kI1O_LM, dt)
    '    deltaprobability(d31_LM, I1_LM, C_LM, kI1C_LM, fC_INa * kC2I1_LM, dt)
    '    deltaprobability(d34_LM, I1_LM, I2_LM, kI1I2_LM, kI2I1_LM, dt)
    '    deltaprobability(d41_LM, I2_LM, C_LM, kI2C_LM, fC_INa * kC2I2_LM, dt)
    '    deltaprobability(d45_LM, I2_LM, Is_LM, kIsf_LM, kIsb_LM, dt)
    '    deltaprobability(d51_LM, Is_LM, C_LM, kIsb_LM, kIsf_LM, dt)

    'End Sub


    Public INamag As Double = 1

    Private Sub calculateINamag(ByVal GHKNa As Double, ByVal GHKK As Double, ByRef INa_TM_Na_cyt As Double, ByRef INa_TM_K_cyt As Double, ByRef INaL_Na_cyt As Double, ByRef INaL_K_cyt As Double)
        ' ************************ Current amplitude of INa_total based on the previous gating condition *************************************
        INaT_Na_cyt = PNa * INamag * (1 - INaLratio) * GHKNa * O_TM * INasf                                       'current component carried by Na+
        INaT_K_cyt = PNa * relativePK_INa * INamag * (1 - INaLratio) * GHKK * O_TM * INasf            'current component carried by K+

        INaL_Na_cyt = PNa * INamag * INaLratio * GHKNa * O_LM * INasf                                            'current component carried by Na+
        INaL_K_cyt = PNa * relativePK_INa * INamag * INaLratio * GHKK * O_LM * INasf                'current component carried by K+
    End Sub

    '********** ICaL **********
    Public Ycc_iz As Double     'state of LCC  gateVm closed, gateCa closed
    Public Yco_iz As Double     'state of LCC  gateVm closed, gateCa open
    Public Yoc_iz As Double     'state of LCC  gateVm open, gateCa closed
    Public Yoo_iz As Double     'state of LCC  gateVm open, gateCa open

    Public dYco_izdt As Double     'state of LCC  gateVm closed, gateCa open
    Public dYoc_izdt As Double     'state of LCC  gateVm open, gateCa closed
    Public dYoo_izdt As Double     'state of LCC  gateVm open, gateCa open

    Public Ycc_blk As Double     'state of LCC  gateVm closed, gateCa closed
    Public Yco_blk As Double     'state of LCC  gateVm closed, gateCa open
    Public Yoc_blk As Double     'state of LCC  gateVm open, gateCa closed
    Public Yoo_blk As Double     'state of LCC  gateVm open, gateCa open

    Public dYco_blkdt As Double     'state of LCC  gateVm closed, gateCa open
    Public dYoc_blkdt As Double     'state of LCC  gateVm open, gateCa closed
    Public dYoo_blkdt As Double     'state of LCC  gateVm open, gateCa open

    Public kcocc_iz As Double              'Ca inactivation
    Public koooc_iz As Double              'Ca inactivation

    Public kcocc_blk As Double              'Ca inactivation
    Public koooc_blk As Double              'Ca inactivation

    Public kccco As Double              'removal of Ca inactivation, only Vm-dependent 
    Public kocoo As Double             'removal of Ca inactivation, only Vm-dependent 
    'Public kccco_blk As Double              'removal of Ca inactivation, only Vm-dependent 
    'Public kocoo_blk As Double             'removal of Ca inactivation, only Vm-dependent 

    Public kcooo_iz As Double              'Vm activation
    Public kooco_iz As Double              'removal of Vm activation
    Public kcooo_blk As Double              'Vm activation
    Public kooco_blk As Double              'removal of Vm activation

    Public kccoc_iz As Double               'Vm activation
    Public koccc_iz As Double               'removal of Vm activation
    Public kccoc_blk As Double               'Vm activation
    Public koccc_blk As Double               'removal of Vm activation

    Public Ca_loc_iz As Double             'local [Ca] at the Ca binding site (calmodulin tethered to LCC)   
    Public Ca_loc_blk As Double             'local [Ca] at the Ca binding site (calmodulin tethered to LCC)   

    Public product_CaL As Double

    Public fixCaconc As Boolean                              'fix [Ca] in all compartments to simulate Ca chelating exp.
    Public orgVshiftCaL As Double = 3         '13 mV
    Public VshiftCaL As Double = orgVshiftCaL
    Public VshiftCaLsf As Double = 1                  'regulation via PatternSearch()

    Public kslowf_CaL As Double        '/ms-1,  forward rate of the ultraslow gate of the ICaL
    Public kslowb_CaL As Double        '/ms-1,  backward rate of the ultraslow gate of the ICaL
    Public us_CaL As Double               '            gating variable of the ultraslow gate of the ICaL
    Public dus_CaLdt As Double               'dy/dt of gating variable us_CaL
    Public kusCaL As Double = 0.55         ' = 1, standard rate factor of ultra slow inactivation adjusted by AN in 210210 to fit the cell 07 AP plateau
    Public FusCaLmag As Double = 1
    Public FusCaLsf As Double = 1                'scaling factor

    Public Sub calculateICaLslowGate()
        kslowf_CaL = kusCaL * FusCaLmag * FusCaLsf / (5000000 * Math.Exp(Vm / 6) + 100 * Math.Exp(Vm / 65))     'ultra slow gate
        kslowb_CaL = kusCaL * FusCaLmag * FusCaLsf / (1000 * Math.Exp(Vm / -14) + 66 * Math.Exp(Vm / -65))        'ultra slow gate
        dus_CaLdt = kslowf_CaL * (1 - us_CaL) - kslowb_CaL * us_CaL                'dy/dt
    End Sub

    'Public Sub calculateICaLslowGate()
    '    Dim v As Double = Vm - VshiftCaL
    '    kslowf_CaL = 1 / (5000000 * Math.Exp(v / 6) + 100 * Math.Exp(v / 65))     'ultra slow gate
    '    kslowb_CaL = 1 / (1000 * Math.Exp(v / -14) + 66 * Math.Exp(v / -65))        'ultra slow gate
    '    dus_CaLdt = kslowf_CaL * (1 - us_CaL) - kslowb_CaL * us_CaL                'dy/dt
    'End Sub

    Public cfkocVm_CaL As Double = 0.5      'adjusting parameter

    Public Sub calculateICaLVmgate(ByRef kcoVm As Double, ByRef kocVm As Double)
        '********** fast Vm gate ***********************************************************
        VshiftCaL = orgVshiftCaL * VshiftCaLsf                        'control VshiftCaL (-3 mV)
        Dim v As Double = Vm - VshiftCaL
        kcoVm = 1.0 / (0.763 * Math.Exp(-v / 8.5) + 0.348 * Math.Exp(-v / 3500))    '
        kocVm = 1 * cfkocVm_CaL / (4.65 * Math.Exp(v / 15) + 1.363 * Math.Exp(v / 100))        'modified on 210217 by AN
        'kocVm = 0.1 / (4.65 * Math.Exp(Vm / 15) + 1.363 * Math.Exp(Vm / 100))                       '

        '********** calculateICaL4stateGate() ***************
        kcooo_blk = kcoVm                'LCC activation
        kooco_blk = kocVm                'deactivation
        kccoc_blk = kcoVm                'LCC activation
        koccc_blk = kocVm                'deactivation

        kcooo_iz = kcoVm                'LCC activation
        kooco_iz = kocVm                'deactivation
        kccoc_iz = kcoVm                'LCC activation
        koccc_iz = kocVm                'deactivation

        '********** CaRU() ***** 8 states transition ************************
        kYcooYooo = kcoVm           'LCC activation
        kYoooYcoo = kocVm           'deactivatin
        kYcocYooc = kcoVm           'LCC activation
        kYoocYcoc = kocVm           'deactivatin

        kYccoYoco = kcoVm           'LCC activation
        kYocoYcco = kocVm           'deactivatin
        kYcccYocc = kcoVm            'LCC activation
        kYoccYccc = kocVm            'deactivation

    End Sub

    'Public Sub calculateICaLVmgateSimple()
    '    dus_CaLdt = 0.0

    '    '********** fast Vm gate ***********************************************************
    '    VshiftCaL = orgVshiftCaL * VshiftCaLsf                       'orgVshiftCaL (-3 mV) * scaling factor
    '    fVmAct = 1.0 / (0.763 * Math.Exp(-(Vm - VshiftCaL) / 8.5) + 0.348 * Math.Exp(-(Vm - VshiftCaL) / 3500))    ' include the VshiftCaLsf
    '    kcoVm = fVmAct                                                                                          '
    '    kocVm = 1.0 / (4.65 * Exp(Vm / 15) + 1.363 * Exp(Vm / 100))                       '

    '    '********** calculateICaL4stateGate() ***************
    '    kcooo_blk = kcoVm                'LCC activation
    '    kooco_blk = kocVm
    '    kccoc_blk = kcoVm                'LCC activation
    '    koccc_blk = kocVm

    '    kcooo_iz = kcoVm                'LCC activation
    '    kooco_iz = kocVm
    '    kccoc_iz = kcoVm                'LCC activation
    '    koccc_iz = kocVm

    '    '********** CaRU() ***** 8 states transition ************************
    '    kYcooYooo = kcoVm           'LCC activation
    '    kYoooYcoo = kocVm
    '    kYcocYooc = kcoVm           'LCC activation
    '    kYoocYcoc = kocVm

    '    kYccoYoco = kcoVm           'LCC activation
    '    kYocoYcco = kocVm
    '    kYcccYocc = kcoVm            'LCC activation
    '    kYoccYccc = kocVm

    'End Sub


    'Public VtoA As Double = 0.25
    Public orgKL As Double = 4.4 / 1000 * 0.9                         '4.4 / 1000       'mM, dissociation constant 
    Public KL As Double
    Public KLmag As Double = 2.5 * 1.002                'revised by AN on 180918

    Public fVmAct As Double
    Dim mEtan1 As Double                                    'component of kco
    Dim mEtan2 As Double
    Dim TL As Double = 147.51                                               '147.51　　　

    Public Sub calculateICaLCagate(ByVal ca As Double, ByRef kocCa As Double)              '201114 revised
        '*****************  Ca-mediated inactivation of LCC dependent on both Ca and fVmAct **********************
        KL = orgKL * KLmag * KLsf
        kocCa = (ca / KL / TL) * 0.35 * kcoVm                                           'Ca-mediated inactivation rate  Eta+
    End Sub

    Public Sub izblkLCC_dydt()              'calculate for both the iz- and blk-spaces
        '**********  Finally, determine dy/dt for three states, CO, OO, OC by using the Vm-dependent rates **************************
        Ycc_iz = 1 - (Yco_iz + Yoo_iz + Yoc_iz)
        dYco_izdt = kccco * Ycc_iz + kooco_iz * Yoo_iz - (kcocc_iz + kcooo_iz) * Yco_iz
        dYoo_izdt = kcooo_iz * Yco_iz + kocoo * Yoc_iz - (kooco_iz + koooc_iz) * Yoo_iz
        dYoc_izdt = koooc_iz * Yoo_iz + kccoc_iz * Ycc_iz - (kocoo + koccc_iz) * Yoc_iz

        Ycc_blk = 1 - (Yco_blk + Yoo_blk + Yoc_blk)
        dYco_blkdt = kccco * Ycc_blk + kooco_blk * Yoo_blk - (kcocc_blk + kcooo_blk) * Yco_blk
        dYoo_blkdt = kcooo_blk * Yco_blk + kocoo * Yoc_blk - (kooco_blk + koooc_blk) * Yoo_blk
        dYoc_blkdt = koooc_blk * Yoo_blk + kccoc_blk * Ycc_blk - (kocoo + koccc_blk) * Yoc_blk
    End Sub

    Public Sub calculateICaLEta(ByRef kccco As Double, ByRef kocoo As Double)
        '*****************  removal of Ca-mediated inactivation (only Vm-dependent)  **********************
        mEtan1 = 1 / (8084 * Math.Exp(Vm / 10) + 158 * Math.Exp(Vm / 1000))        '
        mEtan2 = 1 / (134736 * Math.Exp(Vm / -5) + 337 * Math.Exp(Vm / -2000))   '
        kccco = mEtan1 + mEtan2                        '4 sate model
        kocoo = mEtan1 + mEtan2                        '4 sate model

        kYccoYcoo = kccco                                     'CaRU
        kYcccYcoc = kccco                                     'CaRU
        kYocoYooo = kocoo                                     'CaRU
        kYoccYooc = kocoo                                     'CaRU
    End Sub

    '*************************
    Public orgPCaL As Double = 12.67 * 0.4              '5.068      Ca Permeability factor for L-Ca channel
    Public PCaL As Double                                         'Ca Permeability factor for L-Ca channel
    Public ICaLmag As Double = 1

    Public perNa_ICaL As Double
    Public perK_ICaL As Double                      'the reversal potential of ICaL is shifted negative by this modification
    Public ATPCaL As Double = 6                    'a concentration of ATP for the CaL gating
    Public ATPfactor As Double                        'run-down due to ATP depletion

    Public ICaL As Double                              'L-type Ca current composed of Ca, Na, and K components
    Public ICaL_jnc As Double                       'L-type Ca current composed of Ca, Na, and K components
    Public ICaL_iz As Double                          'L-type Ca current composed of Ca, Na, and K components
    Public ICaL_blk As Double                       'L-type Ca current composed of Ca, Na, and K components

    Public ICaLCa_LR_jnc As Double              'ICaL due to Yooo in the CaRU
    Public ICaLCa_L0_jnc As Double               'ICaL due to Yooc in the CaRU

    Public ICaLCa_jnc As Double
    Public ICaLCa_iz As Double
    Public ICaLCa_blk As Double

    Public ICaLNa_jnc As Double
    Public ICaLNa_iz As Double
    Public ICaLNa_blk As Double

    Public ICaLK_jnc As Double
    Public ICaLK_iz As Double
    Public ICaLK_blk As Double

    Public Sub calculateICaL4stateAmp(ByVal fraction As Double, ByVal pOpen As Double, ByVal ghkCa As Double, ByVal ghkNa As Double, ByVal ghkK As Double, ByRef ICaLCa As Double, ByRef ICaLNa As Double, ByRef ICaLK As Double)
        '****** Calculate the amplitude of ICaL components *******************************************
        PCaL = orgPCaL * ICaLmag * ICaLsf                                      '* fPCaL          

        perNa_ICaL = 0.0000185 * PCaL                                                                   '
        perK_ICaL = 0.000367 * PCaL                                               '= 0.000367 * 0.05 * PCaL                                                                   '
        ATPfactor = 1.0 / (1.0 + ((1.4 / ATPCaL) ^ 3))     '

        product_CaL = pOpen * us_CaL * ATPfactor                               '

        ICaLCa = fraction * PCaL * ghkCa * product_CaL              ' fraction * PCaL blk* ghkCa * product         'pA / pF, amplitude of Ca component  
        ICaLNa = fraction * perNa_ICaL * ghkNa * product_CaL                                                                    'pA / pF, amplitude of  Na component
        ICaLK = fraction * perK_ICaL * ghkK * product_CaL                                                                         'pA / pF, amplitude of K component
    End Sub

    '********************************************************************************
    '****  8 States of the CaRU (voltage-gate LCC, Ca-gate LCC, RyR gate) ************
    Public Yooo As Double
    Public Yooc As Double
    'Public Yooi As Double

    Public Yoco As Double
    Public Yocc As Double
    'Public Yoci As Double

    Public Ycoo As Double
    Public Ycoc As Double
    'Public Ycoi As Double

    Public Ycco As Double
    Public Yccc As Double
    'Public Ycci As Double

    '****************** dy/dt for 8 states
    Public dYooodt As Double
    Public dYoocdt As Double
    'Public dYooidt As Double

    Public dYocodt As Double
    Public dYoccdt As Double
    'Public dYocidt As Double

    Public dYcoodt As Double
    Public dYcocdt As Double
    'Public dYcoidt As Double

    Public dYccodt As Double
    Public dYcccdt As Double
    'Public dYccidt As Double

    '***************  Transiton rate
    Public kYoooYooc As Double      'RyR gate 
    Public kYoocYooo As Double

    Public kYocoYocc As Double
    Public kYoccYoco As Double

    Public kYcooYcoc As Double
    Public kYcocYcoo As Double

    Public kYccoYccc As Double
    Public kYcccYcco As Double

    Public kYoooYcoo As Double      'ICaL gate (Voltage)
    Public kYcooYooo As Double
    Public kYoocYcoc As Double
    Public kYcocYooc As Double

    Public kYocoYcco As Double
    Public kYccoYoco As Double
    Public kYoccYccc As Double
    Public kYcccYocc As Double

    Public kYoooYoco As Double      'ICaL gate (Ca)
    Public kYocoYooo As Double
    Public kYoocYocc As Double
    Public kYoccYooc As Double

    Public kYcooYcco As Double
    Public kYccoYcoo As Double
    Public kYcocYccc As Double
    Public kYcccYcoc As Double

    Public Ca_clamp As Double = 0.00001              'mM, [Ca] under the voltage clamp condition using artificial Ca buffers in the pipette solution.
    Public Ca_ndLR As Double                  'mM, [Ca]nd when both R and L are open
    Public Ca_nd0R As Double                   'mM, [Ca]nd when only R is open
    Public Ca_ndL0 As Double                    'mM, [Ca]nd when only L is open
    Public Ca_nd00 As Double                    'mM, [Ca]nd when both R and L are closed
    Public minCa_ndLR As Double            'peak search for  minimum [Ca]nd when both R and L are open

    Public kRoc As Double        'rate for RyR 
    Public kRco As Double

    Public kcoVm As Double          'rate for activation for LCC
    Public kocVm As Double          'rate for deactivation for LCC

    Public JLCC As Double = 0.000913             'um3ms-1 conductance of a unit  LCC
    Public JRyR As Double = 0.02                       ' um3ms-1 conductance of a unit RyR  in a single CaRU

    Public gD_nd As Double = 0.065
    Public gD_free As Double = 0.065               ' um3ms-1 Diffusion const between channel mouth and  iz-space or blk_space
    Public po_LCC As Double                               'open probability of LCC
    Public po_RyR As Double                               'open probability of RYR
    Public pot_RyR As Double


    Public Sub CaRU_Rate()
        '(1) Rate for the opening and closing rate of couplon (RyR cluster)
        ' ******  Case of LCC is open ********
        RyRkineticsStern2013(Ca_ndL0, Ca_ndLR, kRoc, kRco)     'LCC is open
        kYoooYooc = kRoc                'trigger step closing rate
        kYoocYooo = kRco                'trigger step opening rate

        '' ******  Case of LCC is closed ********
        RyRkineticsStern2013(Ca_nd00, Ca_nd0R, kRoc, kRco)       'LCC is closed
        kYcooYcoc = kRoc               'trigger step closing rate     Ycoo
        kYcocYcoo = kRco               'trigger step opening rate

        kYccoYccc = kRoc               'trigger step closing rate   Ycco
        kYcccYcco = kRco               'trigger step opening rate

        kYocoYocc = kRoc               'trigger step closing rate   Yoco
        kYoccYoco = kRco

        '(2) opening and closing rate of LCC within the CaRU
        Dim kcaoc As Double

        calculateICaLCagate(Ca_ndLR, kcaoc)                            '[Ca]nd when both RyR and LCC are open 
        kYoooYoco = kcaoc           'inactivation step of Ca-gate of Yooo

        calculateICaLCagate(Ca_ndL0, kcaoc)                            '[Ca]nd when  LCC is open
        kYoocYocc = kcaoc            'inactivation step of Ca-gate of Yooc

        calculateICaLCagate(Ca_nd0R, kcaoc)                           '[Ca]nd when  RyR is open
        kYcooYcco = kcaoc             'inactivation step of Ca-gate of Ycoo

        calculateICaLCagate(Ca_nd00, kcaoc)                           '[Ca]nd when both RyR and LCC are closed
        kYcocYccc = kcaoc            'inactivation step of Ca-gate of Ycoc

    End Sub

    Public Sub CaRU_RateSimple()
        '******Calculate the Vm- and Ca-dependent state transition rates of the CaRU *************
        ''''****  prepare Ca_nd for the 4 different combinations of RyR and LCC   *******************
        CaNanoDomain()                       'determine [Ca]nd at various combinations

        '**  Ca gate of RyR (original Hinch model)  *****
        ' ******  Case of LCC is open ********
        RyRkineticsStern2013(Ca_ndL0, Ca_ndLR, kRoc, kRco)     'LCC is open
        kYoooYooc = kRoc                'trigger step closing rate
        kYoocYooo = kRco                'trigger step opening rate

        '' ******  Case of LCC is closed ********
        RyRkineticsStern2013(Ca_nd00, Ca_nd0R, kRoc, kRco)       'LCC is closed
        kYcooYcoc = kRoc               'trigger step closing rate     Ycoo
        kYcocYcoo = kRco               'trigger step opening rate

        kYccoYccc = kRoc               'trigger step closing rate   Ycco
        kYcccYcco = kRco               'trigger step opening rate

        kYocoYocc = kRoc               'trigger step closing rate   Yoco
        kYoccYoco = kRco

        ''*********** the Vm gate of LCC ***********
        'all rate of state transition have been determined by the StoreVarnowtoArray() and RecoverVarlastFromArray()

        '************   Ca gate of LCC , common for all LCC in both nanodomain and bulk spaces ******************

        'Dim kcaco As Double
        Dim kcaoc As Double

        calculateICaLCagate(Ca_ndLR, kcaoc)                            '[Ca]nd when both RyR and LCC are open 
        kYoooYoco = kcaoc           'inactivation step of Ca-gate of Yooo
        kYocoYooo = kocoo          'removal of inactivation 

        calculateICaLCagate(Ca_ndL0, kcaoc)                            '[Ca]nd when  LCC is open
        kYoocYocc = kcaoc            'inactivation step of Ca-gate of Yooc
        kYoccYooc = kocoo            'removal of inactivation

        calculateICaLCagate(Ca_nd0R, kcaoc)                           '[Ca]nd when  RyR is open
        kYcooYcco = kcaoc             'inactivation step of Ca-gate of Ycoo
        kYccoYcoo = kccco             'removal of inactivation

        calculateICaLCagate(Ca_nd00, kcaoc)                           '[Ca]nd when both RyR and LCC are closed
        kYcocYccc = kcaoc            'inactivation step of Ca-gate of Ycoc
        kYcccYcoc = kccco            'removal of inactivation

    End Sub

    Public Sub CaRU_IonFluxes(ByVal vm As Double, ByVal Fraction As Double, ByVal ghkNa As Double, ByVal ghkK As Double, ByRef ICaLCa_LR As Double, ByRef ICaLCa_L0 As Double, ByRef ICaLNa As Double, ByRef ICaLK As Double, ByRef Jrel_SR As Double)

        ''''************    pOpen,     ICaL_jnc and Irel_RyR ***************************************************
        po_LCC = Yooo + Yooc                                    'open probability of LCC
        po_RyR = Yooo + Ycoo + Ycco + Yoco              'open probability of RyR

        '****** conductances *****************************
        PCaL = orgPCaL * ICaLmag      '* fPCaL                                      '0.75  Current density
        perNa_ICaL = 0.0000185 * PCaL                                                                   '
        perK_ICaL = 0.000367 * PCaL                                                                       '
        ATPfactor = 1.0 / (1.0 + ((1.4 / ATPCaL) ^ 3))     '

        '*****  Current amplitude and Ca fluxes *************************
        ICaLCa_LR = Fraction * PCaL * GHKCa_LR * Yooo * us_CaL * ATPfactor             'Ca gradient between ext. and nd  
        ICaLCa_L0 = Fraction * PCaL * GHKCa_L0 * Yooc * us_CaL * ATPfactor             '  Ca gradient between ext. and nd
        ICaLNa = Fraction * perNa_ICaL * ghkNa * po_LCC * us_CaL * ATPfactor             '
        ICaLK = Fraction * perK_ICaL * ghkK * po_LCC * us_CaL * ATPfactor                      '

        pot_RyR = po_RyR + po_baseRyR             'basal open probability is added, in place of a Ca leak out of SR

        Jrel_SR = PRyR * pot_RyR * (Ca_SRrl - Cafree_jnc) * freezeSR             '       

    End Sub

    Public Vshift_base As Double = -5.0                'correction of Vshift
    Public orgExtraVshift As Double = -6.5
    Public extraVshiftsf As Double = 1

    Public IK1sf As Double = 1
    Public IKrsf As Double = 1
    Public IKssf As Double = 1
    Public IKursf As Double = 1
    Public IKtosf As Double = 1

    Public INasf As Double = 1
    Public ICaLsf As Double = 1
    Public KLsf As Double = 1
    Public ICaTsf As Double = 0

    Public Ihasf As Double = 1
    Public Istsf As Double = 1
    Public IbNSCsf As Double = 1

    '**************** InitialIxsf ****************************
    Public initialIK1sf As Double = 1
    Public initialIKrsf As Double = 1
    Public initialIKssf As Double = 1
    Public initialIKursf As Double = 1
    Public initialIKtosf As Double = 1

    Public initialINasf As Double = 1
    Public initialICaLsf As Double = 1
    Public initialKLsf As Double = 1
    Public initialICaTsf As Double = 0

    Public initialIhasf As Double = 1
    Public initialIstsf As Double = 1
    Public initialIbNSCsf As Double = 1

    Public initialVshiftCaLsf As Double = 1
    Public initialVshiftNasf As Double = 1


    Public Sub renewIxmag()         '実行画面の中で、txtIxmag.textを最終的なもので書き換える。
        '************ those which were regulated by pattern search through NP() ******************
        ExtraVshift = orgExtraVshift * extraVshiftsf    'Vshift of the experimental Vm record

        IKrmag = IKrmag * IKrsf                           'PC parameter
        IbNSCmag = IbNSCmag * IbNSCsf           'PC parameter
        IKurmag = IKurmag * IKursf                  'PC parameter
        ICaLmag = ICaLmag * ICaLsf                   'PC parameter
        KLmag = KLmag * KLsf                           'PC parameter
        VshiftCaL = orgVshiftCaL * VshiftCaLsf          'PC parameter
        Pstmag = Pstmag * Istsf

        '****************************************************
        INamag = INamag * INasf
        VshiftNa = orgVshiftNa * VshiftNasf
        IK1mag = IK1mag * IK1sf
        Ihamag = Ihamag * Ihasf

        '*********** not applied by the PS ************************************
        IKsmag = IKsmag * IKssf
        IKtomag = IKtomag * IKtosf

        ICaTmag = ICaTmag * ICaTsf

        'INaKmag = INaKmag * crf_NaK  'exceptional treatment
        'INCXmag = INCXmag * INCXsf   'exceptional treatment
        ' ****** reset the Ixsf to the unit ****************
        extraVshiftsf = 1

        IKrsf = 1
        IbNSCsf = 1
        IKursf = 1
        ICaLsf = 1
        KLsf = 1
        VshiftCaLsf = 1

        IK1sf = 1
        IKssf = 1
        IKtosf = 1
        INasf = 1
        VshiftNasf = 1
        Ihasf = 1
        Istsf = 1
        ICaTsf = 1

    End Sub

    'Dim DCao As Double
    Dim DVm As Double
    Dim expF As Double
    Public ratioBS As Double = 1
    Dim gD_bs As Double       'diffusion conductivity of Ca between the blink space and SRrl
    Dim fbs As Double
    Dim fR As Double
    Dim fL As Double

    Public BlinkS As Boolean = False              'indicate if blink space is defined or not
    Public Cabs_0R As Double
    Public Cabs_LR As Double

    Private Sub CaNanoDomain()
        If Vm > -0.00001 And Vm < 0.00001 Then Vm = 0.00001 '0 cannot be used.
        DVm = Vm / RTF2
        expF = Math.Exp(-DVm)

        gD_bs = gD_nd * ratioBS                    'a new gD_bs is calculated
        fR = JRyR / gD_nd
        fL = JLCC / gD_nd
        fbs = JRyR / gD_bs

        '********  local [Ca] at the vicinity of LCC in the iz- and blk-spacesd dependent on both [Ca]cytsol and ICaL ** note gD-free = gD_nd **********
        Ca_loc_iz = (Cafree_iz + fL * Cao * (DVm * expF) / (1 - expF)) / (1 + fL * DVm / (1 - expF))
        Ca_loc_blk = (Cafree_blk + fL * Cao * (DVm * expF) / (1 - expF)) / (1 + fL * DVm / (1 - expF))

        Ca_ndLR = (Cafree_jnc + Ca_SRrl * fR + expF * Cao * fL * DVm / (1 - expF)) / (1 + fR + fL * DVm / (1 - expF))
        Ca_nd0R = (Cafree_jnc + Ca_SRrl * fR) / (1 + fR)
        Ca_ndL0 = (Cafree_jnc + expF * Cao * fL * DVm / (1 - expF)) / (1 + fL * DVm / (1 - expF))               ' Ca_nd with L open 
        Ca_nd00 = Cafree_jnc                                 'Ca_nd without R and L open


    End Sub

    Public Sub CaRU_dydt()                   'State transition of CaRU
        Yccc = 1 - (Yooo + Yooc + Ycoo + Ycoc + Ycco + Yoco + Yocc)

        dYooodt = kYoocYooo * Yooc + kYcooYooo * Ycoo + kYocoYooo * Yoco - (kYoooYooc + kYoooYcoo + kYoooYoco) * Yooo
        dYoocdt = kYcocYooc * Ycoc + kYoooYooc * Yooo + kYoccYooc * Yocc - (kYoocYcoc + kYoocYooo + kYoocYocc) * Yooc

        dYcoodt = kYcocYcoo * Ycoc + kYoooYcoo * Yooo + kYccoYcoo * Ycco - (kYcooYcoc + kYcooYooo + kYcooYcco) * Ycoo
        dYcocdt = kYcooYcoc * Ycoo + kYoocYcoc * Yooc + kYcccYcoc * Yccc - (kYcocYcoo + kYcocYooc + kYcocYccc) * Ycoc

        dYccodt = kYcccYcco * Yccc + kYocoYcco * Yoco + kYcooYcco * Ycoo - (kYccoYccc + kYccoYoco + kYccoYcoo) * Ycco
        'dYcccdt = kYcciYccc * Ycci + kYccoYccc * Ycco + kYoccYccc * Yocc + kYcocYccc * Ycoc - (kYcccYcci + kYcccYcco + kYcccYocc + kYcccYcoc) * Yccc

        dYocodt = kYoccYoco * Yocc + kYccoYoco * Ycco + kYoooYoco * Yooo - (kYocoYocc + kYocoYcco + kYocoYooo) * Yoco
        dYoccdt = kYcccYocc * Yccc + kYocoYocc * Yoco + kYoocYocc * Yooc - (kYoccYccc + kYoccYoco + kYoccYooc) * Yocc
    End Sub


    Public factive As Double                                       'fraction of open RyR by [Ca]nd_0L
    'Public Km_RyR As Double
    Dim microMCaT As Double                                'uM, trigger [Ca]nd
    Dim microMCaA As Double                                'uM, autonomous [Ca]nd

    '****************** Stern et al. 2013 *********************

    Public ft As Double                                       'fraction of open RyR in the triggering step
    'Dim fn As Double = 3    'V-like cells   5  '7 , representing the nearest neighbor influence of Ca released from one RyR during the regenerative phase
    Dim fn As Double = 7    'N-like cells    '5  '7 , representing the nearest neighbor influence of Ca released from one RyR during the regenerative phase
    Public Km_RyR As Double

    Dim RyRkco As Double                         'kco of a single RyR
    Dim RyRkoc As Double                          'koc of a single RyR
    Public fixedkco As Double
    Dim Qten As Double = 3                         ' adjust for temperature assuming that recordings of the single RyR channel have been conducted at the room temp. 
    Dim sloc0 As Double = 0.1                      'original = 0.1 mM      biasing factor defined in Stern et al 2013
    Dim hry As Double = 2.7                         '2.65 (3 introduced on March 27) original Stern et al 2013
    Dim kom_RyR As Double = 0.5664         '3.0    closing rate     modified from Stern et al 2013
    Dim kon_RyR As Double = 0.4              '0.4  ms-1 mM-1
    Dim khalf_RyR As Double = 0.025        '0.025mM
    Dim pC As Double                                'steady-state probability of closed state used to convert RyR_koc to couplon_kroc 
    Dim Nc As Double = 10                        'number of RyRs in a couplon used to convert RyR_koc to couplon_kroc
    Dim Akrco As Double                           'temporaly couplon activation rate, 
    Dim Ckcop As Double
    Dim ftpC As Double

    Public Sub RyRkineticsStern2013(ByVal caT As Double, ByVal caA As Double, ByRef kroc As Double, ByRef krco As Double)
        '**** triggering  step with caT ********************
        RyRkco = Qten * kon_RyR / (1 + Math.Pow(khalf_RyR / caT, hry))         '/ms, opening rate of a single RyR channel 
        RyRkoc = Qten * kom_RyR                                                                        '/ms, closing rate of a single RyR channel                                                                             ' 
        ft = RyRkco / (RyRkco + RyRkoc)                                                       'assuming an instantaneous equilibrium 

        '**** RyR-RyR cooperative step with caA ***, positive feedback step ********************
        Akrco = Qten * kon_RyR * (sloc0 + Ca_SRrl) / (1 + Math.Pow(khalf_RyR / caA, hry))         ' opening rate of couplon  dependent on [Ca]SRrl  with caT
        krco = fn * ft * Akrco                              '7,  overall opening rate of a couplon with caA

        Ckcop = Qten * kon_RyR / (1 + Math.Pow(khalf_RyR / Ca_nd00, hry))         '/ms, opening rate of a single RyR channel 
        ftpC = Ckcop / (Ckcop + RyRkoc)
        pC = RyRkoc / (ftpC * Akrco + RyRkoc)                       'steady-state probability of closed state

        kroc = RyRkoc * pC ^ ((Nc - 1) * 0.74)                         'closing rate of couplon

        'fixedkco = fn * ft * Qten * kon_RyR * (0.7337) / (1 + Math.Pow(khalf_RyR / caA, hry))         ' opening rate of couplon  dependent on [Ca]SRrl  with caT

    End Sub


    '********** PMCA  **********
    Const KmPMCA As Double = 0.0005       'mM
    Public IPMCA As Double
    Public IPMCA_Ca_iz As Double
    Public IPMCA_Ca_blk As Double

    Public dATP_PMCA_iz As Double        'mM   consumption of ATP by PMCA
    Public dATP_PMCA_blk As Double

    Private Sub calculateIPMCA(ByVal frc, ByVal Ca, ByRef IPMCA, ByRef dNATP_PMCA)
        IPMCA = frc * maxIPMCA * Math.Pow(Ca, 1.6) / (Math.Pow(KmPMCA, 1.6) + Math.Pow(Ca, 1.6))
        dNATP_PMCA = IPMCA * Cm / 2.0 / Faraday                         ' amole / ms
    End Sub

    '********** contration of the NL model 2008  **********
    Public IsoMetric As Boolean = True
    Public Ca As Double                             'mM,  concentration of free Ca 

    Public halfSL As Double = 1.0                      'um, half sarcomere length
    Public crsBrLnX As Double = 0.994              'um, X in the NL model

    Public TSt As Double = 23 / 6                  'uM  total TS,  = number of Tn / nCaBS
    Public nCaBS As Double = 3                 'number of T in a troponin system

    Public TS As Double                              'uM, troponin-system Ca-free
    Public TSCa3 As Double                        'uM, TS bound with 3 Ca
    Public TSCa3W As Double                     'uM, 
    Public TSCa3S As Double                      'uM, 
    Public TSS As Double                            'uM, 
    Public TSW As Double                           'uM, 

    Public dydtTS As Double                        'uM/msec,  rate of change 
    Public dTSCa3dt As Double                    'uM/msec,
    Public dTSCa3Wdt As Double                 'uM/msec,
    Public dTSCa3Sdt As Double                  'uM/msec
    Public dTSSdt As Double                         'uM/msec,
    Public dTSWdt As Double                        'uM/msec,

    Public dydtCa As Double                          'uM/msec, rate of Ca change

    Public Yb As Double = 0.1816                    ' / uM3/ ms, Ca-binding rate
    Public Zb As Double = 0.1397                     '/ms,    Ca-unbinding rate

    Public ratef As Double = 0.0023                 '/ ms, rate for weak CB
    Public eqvhalfSL As Double = 1.15            'um,  a half sarcomere length, giving a maximum rate f
    Public convertF As Double = 15                 'L dependency of rate f

    Public Yp As Double = 0.1397                    '/ms, forming strong CB
    Public Zp As Double = 0.2095                    '/ms/mM, relaxing to weak CB

    Public Yr As Double = 0.1397                  'Ca unbinding rate from strong CB
    Public Zr As Double = 7.2626               'Ca binding rate to strong CB  
    Public Yq As Double = 0.2328               'relaxing to weak CB without Ca
    Public Zq As Double = 0.3724               'forming strong CB without Ca

    Public Za As Double = 0.0023
    Public Yv As Double = 1.5
    Public Yvd As Double = Yv                '1.5   dependency on h of gd, magnifying factor

    Public rateg As Double                       '/ms,  releasing CB weak
    Public rategd As Double                     '/ms, relaxation rate
    Public Yd As Double = 0.0333           'primary rate of gd
    Public Yc As Double = 1                     'dependency on half sarcomere length of Yd 
    Public Lc As Double = 1.2                  'reference length of sarcomere length

    Public Fb As Double                           ' mN mm-2  force of cross bridge
    Public Fbw As Double                        ' mN mm-2  force of cross bridge
    Public Fbp As Double                         ' mN mm-2  force of cross bridge
    Public hw As Double                           'um, elongation of weak cross bridge
    Public dhwdt As Double                      'um/ms, rate of hw change
    Public hwr As Double = 0.0001           'um, reference value of h
    Public hp As Double                             'um, elongation of strong cross bridge
    Public dhpdt As Double                        'um/ms, rate of hw change
    Public hpr As Double = 0.006               'um, reference value of h
    Public rateB As Double = 0.5               '1.2    '0.5    'amplitude factor for h change

    Public Ap As Double = 2700                 'mN mm-2 um-1 uM-1      force of myofilaments strong cross bridge
    Public Aw As Double = 540                   'mN mm-2 um-1 uM-1       weak cross bridge force of myofilaments

    Public Fp As Double                              'mN mm-2                           force of parallel elastic component
    Public Ke As Double = 105000               'mN mm-2 um-5
    Public L0 As Double = 0.97                    'um    resting half sarcomere length giving Fp = 0
    Public Le As Double = 10                        'mN mm-2 um-5                 'spring constant for a linear elastic component
    Public Fext As Double                             'mN mm-2                external force
    Public isometricL As Double = 1.0                    'um, 
    Public dATP_contraction As Double        'uM/ms            usage of ATP by contraction
    Const RatioATP_contraction As Double = 0.08             '0.08    0.025 a tentative scaling factor to adjust the ATP consumption   
    Public JCaTnC_3Ca As Double                         'Ca flux due to Ca binding to Troponin system 


    Dim d12 As Double
    Dim d23 As Double
    Dim d34 As Double
    Dim d45 As Double
    Dim d56 As Double
    Dim d61 As Double


    Public Sub ContractionNL2008(ByVal mMCa As Double)
        Dim uMCa As Double = mMCa * 1000                          'uM,  convert from mM to microM

        crsBrLnX = halfSL - hp                                                      'um, renew crsBrLnX
        TS = TSt - TSCa3 - TSCa3W - TSCa3S - TSS - TSW           'uM, renew TS

        Dim propFh As Double = 28000                                    'um-2,  converting factor
        rateg = Za + Yv * (1 - Exp(-propFh * (hw - hwr) ^ 2))                                              '/ms, rateg dependent on h 
        rategd = Yd + Yc * (halfSL - Lc) ^ 2 + Yvd * (1 - Exp(-propFh * (hw - hwr) ^ 2))    '/ms, rate gd dependent on both L and h

        '************ State transition according to a new [Ca] *******************************
        deltaprobability(d12, TS, TSCa3, (Yb * uMCa ^ nCaBS), Zb, dt)
        deltaprobability(d23, TSCa3, TSCa3W, ratef * Exp(-convertF * (halfSL - eqvhalfSL) ^ 2), rateg, dt)
        deltaprobability(d34, TSCa3W, TSCa3S, Yp, Zp, dt)
        deltaprobability(d45, TSCa3S, TSS, Yr, (Zr * uMCa ^ nCaBS), dt)
        deltaprobability(d56, TSS, TSW, Yq, Zq, dt)
        deltaprobability(d61, TSW, TS, rategd, rategd * 0.01, dt)

        'dTSCa3dt = Yb * TS * uMCa ^ nCaBS - Zb * TSCa3 + rateg * TSCa3W - ratef * Exp(-convertF * (halfSL - eqvhalfSL) ^ 2) * TSCa3  'uM/ms, 
        'dTSCa3Wdt = ratef * Exp(-convertF * (halfSL - eqvhalfSL) ^ 2) * TSCa3 - rateg * TSCa3W + Zp * TSCa3S - Yp * TSCa3W      'uM/ms
        'dTSCa3Sdt = Yp * TSCa3W - Zp * TSCa3S + Zr * TSS * uMCa ^ nCaBS - Yr * TSCa3S       'uM/ms
        'dTSSdt = Yr * TSCa3S - Zr * TSS * uMCa ^ nCaBS + Zq * TSW - Yq * TSS      'uM/ms
        'dTSWdt = Yq * TSS - Zq * TSW - rategd * TSW                                         'uM/ms

        '************ Time-dependent changes in hw and hs  *******************************
        dhwdt = -rateB * (hw - hwr)            'um/ms,
        dhpdt = -rateB * (hp - hpr)              'um/ms,

        dATP_contraction = RatioATP_contraction * (Yp * TSCa3W + Zq * TSW) * Vol_cyt            ' atto mole  (10-15)/ ms

        If IsoMetric = True Then                       'determine halfSL according to the isometric or isotonic contraction
            halfSL = isometricL                      'um,  
        Else

        End If

        '*************** developed tension ********************************************
        FbFp(halfSL, hp, hw, Fb, Fp)

    End Sub

    Public Sub FbFp(ByVal hsl As Double, ByVal thp As Double, ByVal thw As Double, ByRef Fb As Double, ByRef Fp As Double)
        '*************** developed tension ********************************************
        Fbw = nCaBS * Aw * (TSCa3W + TSW) * thw                   'thw = hw valid only in this sub class
        Fbp = nCaBS * Ap * (TSCa3S + TSS) * thp          'thp = hp valid only in this sub class
        Fb = Fbw + Fbp                       'force of cross bridge, The proportion factors Aw and Ap were scaled by 3 beforehand
        Fp = Le * (hsl - L0) + Ke * (hsl - -L0) ^ 5      'force of parallel elastic component
    End Sub

    Public dab As Double       'uM, net change in the clockwise direction
    Public Sub deltaprobability(ByRef dab As Double, ByVal statePre As Double, ByVal statePost As Double, ByVal Fward As Double, ByVal Bward As Double, ByVal dt As Double)
        Dim SumTwo As Double = statePre + statePost     'uM, total amount of pre and post 
        Dim Y0 As Double = statePost / SumTwo               'ratio 
        Dim tau As Double = 1 / (Fward + Bward)
        Dim Yinf As Double = Fward * tau
        dab = SumTwo * (Yinf - Y0) * (1 - Math.Exp(-dt / tau))
    End Sub


    Public optVm As Double = 0.06        'mV, an optimum or standard dVm  
    Public upperLimitDt As Double = 0.15     'ms, upper limit of Dt       max upperlimitDt = 0.14 
    Public lowerLimitDt As Double = 0.002           'ms, lower limit of Dt
    Public Sub adjustDT(ByRef dt As Double)
        Dim temporaryDT As Double                                  'ms, temporary dt
        temporaryDT = Abs(optVm / dVmdt)                       'ms, determine a temporary dt
        dt = Int(1000 * temporaryDT) / 1000                   'ms, calculate to three decimal places
        If dt >= upperLimitDt Then dt = upperLimitDt 'ms, constrain by the upper limit of dt
        If dt <= lowerLimitDt Then dt = lowerLimitDt 'ms, constrain by the lower limit of dt
    End Sub

    '******************************** cytosol ********************************
    Public HBuff_cyt As Double                 'H+ buffer power 
    Public pH_cyt As Double

    Public Pifree_cyt As Double                      'mM concentration
    Public AMP_cyt As Double                  'mM concentration
    Public ADPfr_cyt As Double                'mM concentration
    Public MgADP_cyt As Double             'mM concentration
    Public ATPfr_cyt As Double                 'mM concentration
    Public MgATP_cyt As Double              'mM concentration
    Public Cr_cyt As Double                      'mM concentration

    Public Mg_Mitmodel As Double = 4                                             'mM   only for the Mitochondria model

    Public effiADP As Double

    Public Sub substrateConcentrations()
        pH_cyt = 7.0
        HBuff_cyt = 108695.65217391304
        Pifree_cyt = 0.17149891579359286
        AMP_cyt = 0.00017449886446918583
        ADPfr_cyt = 0.0018220494350193984
        MgADP_cyt = 0.0210034517005118
        ATPfr_cyt = 0.03982306163021869
        MgATP_cyt = 6.6371769383697812
        Cr_cyt = 11.1879765283269

    End Sub

    Public Sub InitialValuesCell38()

        Vm = -20.05196424984273
        Nai = 6.100014384285112
        Ki = 143.89256988279371
        Catot_blk = 0.015115796212935604
        CaMCa = 0.0010916823069858032
        TnChCa = 0.0064557237481047136
        SRCa = 0.006279554554988233
        Catot_iz = 0.13157640708310089
        Catot_jnc = 0.24050343577388711
        Ca_SRup = 1.2261823499563009
        Catot_SRrl = 2.7058087083253852
        Yooo = 0.013714433652059644
        Yooc = 0.026348717464784104
        Ycoo = 0.0013883571188226494
        Ycoc = 0.038884230751316827
        Ycco = 0.0000087728613206462436
        Yoco = 0.000033157080830614874
        Yocc = 0.46045135744776444
        us_CaL = 0.60881166534754794
        Yco_iz = 0.0926774070692049
        Yoc_iz = 0.40823091515141863
        Yoo_iz = 0.092316750494020175
        Yco_blk = 0.093201077765900725
        Yoc_blk = 0.40770934565671946
        Yoo_blk = 0.092838319988719317
        O_TM = 0.0000068312585456323338
        I2_TM = 0.00025462651284768905
        Is_TM = 0.99973382418267731
        O_LM = 0.00016083383510359889
        I1_LM = 0.036493035036205074
        I2_LM = 0.0024349914935781246
        Is_LM = 0.960899872715378
        d_CaT = 0.67977515037882263
        f_CaT = 0.00028163386638480005
        d_st = 0.911976645120246
        f_st = 0.05937548570802282
        us_st = 0.2964772143934955
        C1_ha = 0.99999925499417219
        C2_ha = 0.00000074271692458692563
        O1_ha = 0.000000002282783338746808
        O2_ha = 0.0000000000061053830520744843
        Pbspm = 0.9594326804360791
        y1_IKr = 0.84795557552639833
        y2_IKr = 0.96103641503906923
        y3_IKr = 0.22269244770408811
        ua = 0.95651593899115417
        ui = 0.94714292949327661
        Ov_IKs = 0.29189552672097785
        C2IKs_iz = 0.0096312606596045472
        OcIKs_iz = 0.012772217641117434
        C2IKs_blk = 0.0096653436370830077
        OcIKs_blk = 0.011582040758377108
        y1_IKto = 0.0508124627899777
        y2_IKto = 0.056559024428325562
        O_ACh = 0.015463362624230061
        E1NCX_iz = 0.382461561903937
        I1NCX_iz = 0.15865792851880209
        I2NCX_iz = 0.44789932633727353
        E1NCX_blk = 0.35361659258783745
        I1NCX_blk = 0.16360600376017614
        I2NCX_blk = 0.473263345649443
        P1_6_NaK = 0.56727274613136225
        P7_NaK = 0.09754745525592072
        P8_13_NaK = 0.029804702539668
        TSCa3 = 0.13106641936491945
        TSCa3W = 0.12173784154243034
        TSCa3S = 0.078863247828962757
        TSS = 0.054464028020116968
        TSW = 0.033282472493233781
        hw = 0.0001
        hp = 0.006
        halfSL = 1.0

        totalCai = Catot_jnc * Vol_jnc + Catot_iz * Vol_iz + Catot_blk * Vol_blk + Ca_SRup * Vol_SRup + Catot_SRrl * Vol_SRrl


        '********* revised on 210409 +++********************
        Vm = -20.069562921151526
        Nai = 6.1000003722288456
        Ki = 143.89259005797783
        Catot_blk = 0.014451654482692117
        CaMCa = 0.0009595943888938321
        TnChCa = 0.00653624347995555
        SRCa = 0.0059415914514628311
        Catot_iz = 0.11904428786838589
        Catot_jnc = 0.2175906734919392
        Ca_SRup = 1.2633420840382916
        Catot_SRrl = 3.3262076213531389
        Yooo = 0.0083058498978546436
        Yooc = 0.0128393464734082
        Ycoo = 0.000892469971593947
        Ycoc = 0.020541056359994515
        Ycco = 0.000007589270406732176
        Yoco = 0.000026893677608841452
        Yocc = 0.47776809622812144
        us_CaL = 0.57775378446080339
        Yco_iz = 0.058760220947810957
        Yoc_iz = 0.44083700091681344
        Yoo_iz = 0.058103185360179672
        Yco_blk = 0.059064098766264124
        Yoc_blk = 0.4405365928089881
        Yoo_blk = 0.058403593468004923
        O_TM = 0.0000068181464276234408
        I2_TM = 0.00023441954274681465
        Is_TM = 0.99975431320620733
        O_LM = 0.00016006543497500592
        I1_LM = 0.036226731538150765
        I2_LM = 0.0023493555863980911
        Is_LM = 0.96125399796965871
        d_CaT = 0.68041842002196118
        f_CaT = 0.00027755752071340523
        d_st = 0.91211031905524731
        f_st = 0.047205743982573822
        us_st = 0.259737234453675
        C1_ha = 0.99999931438874556
        C2_ha = 0.00000068365881948147854
        O1_ha = 0.0000000019476590322478347
        O2_ha = 0.0000000000047656882978349467
        Pbspm = 0.95843059010816578
        y1_IKr = 0.86320846374675186
        y2_IKr = 0.9650093995926684
        y3_IKr = 0.22206059042927057
        ua = 0.95647705016792
        ui = 0.94624130619791957
        Ov_IKs = 0.31411087816404132
        C2IKs_iz = 0.009672830983387044
        OcIKs_iz = 0.014839177259332823
        C2IKs_blk = 0.009694876408396863
        OcIKs_blk = 0.013690952661489566
        y1_IKto = 0.05098189076705
        y2_IKto = 0.052348627000544781
        O_ACh = 0.015158920568915357
        E1NCX_iz = 0.3498073033048314
        I1NCX_iz = 0.15712326705700985
        I2NCX_iz = 0.4838032895613254
        E1NCX_blk = 0.32867361501905418
        I1NCX_blk = 0.16097340640347682
        I2NCX_blk = 0.50203535777981978
        P1_6_NaK = 0.566986546502447
        P7_NaK = 0.097914024668963456
        P8_13_NaK = 0.02981778104960927
        TSCa3 = 0.08617425938898253
        TSCa3W = 0.10276009214128276
        TSCa3S = 0.065215369248098912
        TSS = 0.066426744685386008
        TSW = 0.039683085612442062
        hw = 0.0001
        hp = 0.006
        halfSL = 1.0


        totalCai = Catot_jnc * Vol_jnc + Catot_iz * Vol_iz + Catot_blk * Vol_blk + Ca_SRup * Vol_SRup + Catot_SRrl * Vol_SRrl


    End Sub

    Public Sub InitialValuesCell10()
        Vm = -20.156798392077597
        Nai = 6.1000783270438061
        Ki = 143.89248753750056
        Catot_blk = 0.014887268920262865
        CaMCa = 0.0010943648214736034
        TnChCa = 0.006622870655270046
        SRCa = 0.0059496934592921354
        Catot_iz = 0.13008937464716064
        Catot_jnc = 0.23772846578743626
        Ca_SRup = 1.2514140449798554
        Catot_SRrl = 2.680759891165637
        Yooo = 0.010784400204065264
        Yooc = 0.020817019561970802
        Ycoo = 0.0010883846395494206
        Ycoc = 0.028509904420787654
        Ycco = 0.0000085950774582805342
        Yoco = 0.000033653323824999657
        Yocc = 0.487133828471146
        us_CaL = 0.76821768271858359
        Yco_iz = 0.066013099103476788
        Yoc_iz = 0.44817401889032665
        Yoo_iz = 0.0705948826706803
        Yco_blk = 0.066238315691346042
        Yoc_blk = 0.44793322228667454
        Yoo_blk = 0.070835679274332264
        O_TM = 0.0000065883263541192984
        I2_TM = 0.00021194810604058327
        Is_TM = 0.99977641496512692
        O_LM = 0.00010811843165812939
        I1_LM = 0.024271641901980005
        I2_LM = 0.0015745719073057142
        Is_LM = 0.97403545083208232
        d_CaT = 0.683373847448214
        f_CaT = 0.00026948701788104067
        d_st = 0.91277884521878416
        f_st = 0.08891557358852438
        us_st = 0.50234179643501942
        C1_ha = 0.99999938354833873
        C2_ha = 0.00000061485084715214525
        O1_ha = 0.0000000015972778117520817
        O2_ha = 0.0000000000035294177163896856
        Pbspm = 0.96656232457646329
        y1_IKr = 0.8763337545270049
        y2_IKr = 0.96608136332084638
        y3_IKr = 0.22037965708967225
        ua = 0.95665521881798543
        ui = 0.953557307559391
        Ov_IKs = 0.23269269416519123
        C2IKs_iz = 0.0096368806170875187
        OcIKs_iz = 0.018738291664166719
        C2IKs_blk = 0.00966230135702537
        OcIKs_blk = 0.017231125188159682
        y1_IKto = 0.051415020897868167
        y2_IKto = 0.053693531573048951
        O_ACh = 0.015243937860034442
        E1NCX_iz = 0.38063900370970133
        I1NCX_iz = 0.1539897565357832
        I2NCX_iz = 0.454531046205335
        E1NCX_blk = 0.35870238848525215
        I1NCX_blk = 0.1580332284327054
        I2NCX_blk = 0.47353573952691708
        P1_6_NaK = 0.566669521325671
        P7_NaK = 0.0983631577895669
        P8_13_NaK = 0.029769006176616736
        TSCa3 = 0.13814309963135682
        TSCa3W = 0.10196265624918477
        TSCa3S = 0.066744095494094
        TSS = 0.0455024570127217
        TSW = 0.028718930214877366
        hw = 0.0001
        hp = 0.006
        halfSL = 1
    End Sub

    Public Sub InitialValuesCell91()

        Vm = -20.064487801839494
        Nai = 6.0999876437224216
        Ki = 143.89259621008597
        Catot_blk = 0.014454562860791156
        CaMCa = 0.00095989556975639722
        TnChCa = 0.0065362365676971089
        SRCa = 0.0059423318798829429
        Catot_iz = 0.1190623177426339
        Catot_jnc = 0.21762782757758764
        Ca_SRup = 1.263343503652874
        Catot_SRrl = 3.3260087460959489
        Yooo = 0.0083625286591381843
        Yooc = 0.013083662116647396
        Ycoo = 0.00084298738655880386
        Ycoc = 0.020219205327111408
        Ycco = 0.0000074397759791158365
        Yoco = 0.000027912079275284691
        Yocc = 0.48538974188521439
        us_CaL = 0.57776107417519174
        Yco_iz = 0.057806389053814348
        Yoc_iz = 0.44786545963504848
        Yoo_iz = 0.058998385105226717
        Yco_blk = 0.0581055304550902
        Yoc_blk = 0.44756021704595722
        Yoo_blk = 0.05930362769431792
        O_TM = 0.0000068683346028343
        I2_TM = 0.00023358086703806103
        Is_TM = 0.9997551993044631
        O_LM = 0.00016047938522051954
        I1_LM = 0.036235853037921831
        I2_LM = 0.0023478673479486287
        Is_LM = 0.96124643124686993
        d_CaT = 0.68180710161128566
        f_CaT = 0.00027610886602758163
        d_st = 0.91244115094825284
        f_st = 0.047131798079102658
        us_st = 0.25974412812479414
        C1_ha = 0.99999931802136577
        C2_ha = 0.00000068004274156454173
        O1_ha = 0.0000000019311712842327791
        O2_ha = 0.0000000000047113267310678262
        Pbspm = 0.95841735397517591
        y1_IKr = 0.86350813431402218
        y2_IKr = 0.96505973610107354
        y3_IKr = 0.22158335182166719
        ua = 0.95665639694356586
        ui = 0.946234978219953
        Ov_IKs = 0.314101298872809
        C2IKs_iz = 0.0096727622375409415
        OcIKs_iz = 0.014838999644748017
        C2IKs_blk = 0.0096948094411263112
        OcIKs_blk = 0.013690805625535497
        y1_IKto = 0.051090135192698075
        y2_IKto = 0.052303498076855942
        O_ACh = 0.015154301034485194
        E1NCX_iz = 0.34986233808218592
        I1NCX_iz = 0.15712321246879937
        I2NCX_iz = 0.48373490857876489
        E1NCX_blk = 0.32873244906698973
        I1NCX_blk = 0.16097333416192819
        I2NCX_blk = 0.5019642177330722
        P1_6_NaK = 0.56697440978757774
        P7_NaK = 0.097932272907336027
        P8_13_NaK = 0.029796283464695927
        TSCa3 = 0.086259910150405045
        TSCa3W = 0.10279914477959942
        TSCa3S = 0.065245066476178892
        TSS = 0.066396472602149079
        TSW = 0.0396674143482549
        hw = 0.0001
        hp = 0.006
        halfSL = 1.0

        totalCai = Catot_jnc * Vol_jnc + Catot_iz * Vol_iz + Catot_blk * Vol_blk + Ca_SRup * Vol_SRup + Catot_SRrl * Vol_SRrl

    End Sub


    Public Sub InitialValuesCell86()
        Vm = -20.05196424984273
        Nai = 6.100014384285112
        Ki = 143.89256988279371
        Catot_blk = 0.015115796212935604
        CaMCa = 0.0010916823069858032
        TnChCa = 0.0064557237481047136
        SRCa = 0.006279554554988233
        Catot_iz = 0.13157640708310089
        Catot_jnc = 0.24050343577388711
        Ca_SRup = 1.2261823499563009
        Catot_SRrl = 2.7058087083253852
        Yooo = 0.013714433652059644
        Yooc = 0.026348717464784104
        Ycoo = 0.0013883571188226494
        Ycoc = 0.038884230751316827
        Ycco = 0.0000087728613206462436
        Yoco = 0.000033157080830614874
        Yocc = 0.46045135744776444
        us_CaL = 0.60881166534754794
        Yco_iz = 0.0926774070692049
        Yoc_iz = 0.40823091515141863
        Yoo_iz = 0.092316750494020175
        Yco_blk = 0.093201077765900725
        Yoc_blk = 0.40770934565671946
        Yoo_blk = 0.092838319988719317
        O_TM = 0.0000068312585456323338
        I2_TM = 0.00025462651284768905
        Is_TM = 0.99973382418267731
        O_LM = 0.00016083383510359889
        I1_LM = 0.036493035036205074
        I2_LM = 0.0024349914935781246
        Is_LM = 0.960899872715378
        d_CaT = 0.67977515037882263
        f_CaT = 0.00028163386638480005
        d_st = 0.911976645120246
        f_st = 0.05937548570802282
        us_st = 0.2964772143934955
        C1_ha = 0.99999925499417219
        C2_ha = 0.00000074271692458692563
        O1_ha = 0.000000002282783338746808
        O2_ha = 0.0000000000061053830520744843
        Pbspm = 0.9594326804360791
        y1_IKr = 0.84795557552639833
        y2_IKr = 0.96103641503906923
        y3_IKr = 0.22269244770408811
        ua = 0.95651593899115417
        ui = 0.94714292949327661
        Ov_IKs = 0.29189552672097785
        C2IKs_iz = 0.0096312606596045472
        OcIKs_iz = 0.012772217641117434
        C2IKs_blk = 0.0096653436370830077
        OcIKs_blk = 0.011582040758377108
        y1_IKto = 0.0508124627899777
        y2_IKto = 0.056559024428325562
        O_ACh = 0.015463362624230061
        E1NCX_iz = 0.382461561903937
        I1NCX_iz = 0.15865792851880209
        I2NCX_iz = 0.44789932633727353
        E1NCX_blk = 0.35361659258783745
        I1NCX_blk = 0.16360600376017614
        I2NCX_blk = 0.473263345649443
        P1_6_NaK = 0.56727274613136225
        P7_NaK = 0.09754745525592072
        P8_13_NaK = 0.029804702539668
        TSCa3 = 0.13106641936491945
        TSCa3W = 0.12173784154243034
        TSCa3S = 0.078863247828962757
        TSS = 0.054464028020116968
        TSW = 0.033282472493233781
        hw = 0.0001
        hp = 0.006
        halfSL = 1.0

        totalCai = Catot_jnc * Vol_jnc + Catot_iz * Vol_iz + Catot_blk * Vol_blk + Ca_SRup * Vol_SRup + Catot_SRrl * Vol_SRrl

    End Sub


    Public Sub defaultSTARTV()
        '***********InitialVariables091V()  *******************
        Vm = -20.030330894887673
        Nai = 6.402488558988102
        Ki = 141.96409377483937
        Catot_blk = 0.025109626322079606
        CaMCa = 0.0024237600092523
        TnChCa = 0.0068038705011758413
        SRCa = 0.00979912862197902
        Catot_iz = 0.18885329391255573
        Catot_jnc = 0.34458582501720553
        Ca_SRup = 1.8646759379050419
        Catot_SRrl = 4.010313027211942
        Yooo = 0.0053371808075338692
        Yooc = 0.010349687915133405
        Ycoo = 0.0012314488345703803
        Ycoc = 0.0337557969949814
        Ycco = 0.000062761954172918389
        Yoco = 0.000039822950258941887
        Yocc = 0.29499084955978244
        us_CaL = 0.42393063866500386
        Yco_iz = 0.12266708169028295
        Yoc_iz = 0.25562950871841933
        Yoo_iz = 0.055088032514289433
        Yco_blk = 0.12364662993090736
        Yoc_blk = 0.25518963365724978
        Yoo_blk = 0.055527907575459036
        O_TM = 0.000015897087741892657
        I2_TM = 0.0006983041465924455
        Is_TM = 0.9992729135343934
        O_LM = 0.0001935061388938792
        I1_LM = 0.039726747129604355
        I2_LM = 0.0035120053785012103
        Is_LM = 0.956542947167315
        d_CaT = 0.689398936891797
        f_CaT = 0.00026118126970431377
        d_st = 0.9141846278131025
        f_st = 0.04794303960434132
        us_st = 0.31343471408662626
        C1_ha = 0.999999415100073
        C2_ha = 0.0000005834359024655724
        O1_ha = 0.0000000014608990627701759
        O2_ha = 0.0000000000031197841671389192
        Pbspm = 0.950400149339782
        y1_IKr = 0.87762920548151557
        y2_IKr = 0.96609120999413678
        y3_IKr = 0.21796742686687348
        ua = 0.957326766249491
        ui = 0.94784429902839806
        Ov_IKs = 0.2930519163208013
        C2IKs_iz = 0.0093292334687923046
        OcIKs_iz = 0.032368707572828677
        C2IKs_blk = 0.0093581022323286269
        OcIKs_blk = 0.030688615589083295
        y1_IKto = 0.052006397553420576
        y2_IKto = 0.071789330455696951
        O_ACh = 0.015072747160145094
        E1NCX_iz = 0.53961187054900628
        I1NCX_iz = 0.14046881743788464
        I2NCX_iz = 0.29481288305505909
        E1NCX_blk = 0.52820817376727691
        I1NCX_blk = 0.14368580992508298
        I2NCX_blk = 0.3043127251958565
        P1_6_NaK = 0.55553336810767939
        P7_NaK = 0.11274656925482612
        P8_13_NaK = 0.03244231787451117
        TSCa3 = 0.96917503758614065
        TSCa3W = 0.49435538748682106
        TSCa3S = 0.32929719156603476
        TSS = 0.018190586171508309
        TSW = 0.012512085587589913
        hw = 0.0001
        hp = 0.006
        halfSL = 1.0

    End Sub

    Public Sub defaultSTARTA()

        Vm = -73.261962405254934
        Nai = 6.3544170921728274
        Ki = 141.85099855969611
        Catot_blk = 0.011217965556457833
        CaMCa = 0.00045210069073215771
        TnChCa = 0.0066458454471230357
        SRCa = 0.0032993814869566483
        Catot_iz = 0.0745020930812615
        Catot_jnc = 0.13666108625012649
        Ca_SRup = 1.485124250319698

        Catot_SRrl = 8.1430731604405526
        'Yooo = 0.00020253669428842405
        'Yooc = 0.00012586070783551491
        'Ycoo = 0.00017647414490221352
        'Ycoc = 0.99311947176338911
        'Ycco = 0.000000040324240633941138
        'Yoco = 0.000000007108662528364609
        'Yocc = 0.0000021186469054078025
        us_CaL = 0.99972007240039673
        Yco_iz = 0.99383496009503713

        Yoc_iz = 0.0000019371856741299545
        Yoo_iz = 0.00032858597201774521
        Yco_blk = 0.99384232454847854
        Yoc_blk = 0.0000019347508070066086
        Yoo_blk = 0.00032858840688486851
        O_TM = 0.0000005409268138235363
        I2_TM = 0.016654931481386005
        Is_TM = 0.83016496066441769
        O_LM = 0.0000023747125465770216
        I1_LM = 0.001370043404471766

        I2_LM = 0.016746782914056318
        Is_LM = 0.82915652611479584
        d_CaT = 0.00016261322228289861
        f_CaT = 0.44900248485290029
        d_st = 0.028583444490459756
        f_st = 0.96031474429165442
        us_st = 0.99862710093384377
        C1_ha = 0.97488226144001833
        C2_ha = 0.002820907457687366
        O1_ha = 0.0031484666642099626

        O2_ha = 0.0047793908174814971
        O_IK1 = 0.4651698
        Pbspm = 0.86962998614798059
        y1_IKr = 0.0045264981101403009
        y2_IKr = 0.09091112579532537
        y3_IKr = 0.9938018090199191
        ua = 0.0088451475850216781
        ui = 0.97091758813468176
        Ov_IKs = 0.0210874638535362
        C2IKs_iz = 0.009797980495198956

        OcIKs_iz = 0.019854801182127756
        C2IKs_blk = 0.0098010335871594571
        OcIKs_blk = 0.018859196820498303
        y1_IKto = 0.0025535181237643628
        y2_IKto = 0.65403199010083157
        O_ACh = 0.037606368462374695
        E1NCX_iz = 0.20432606394387906
        I1NCX_iz = 0.1584577127394789
        I2NCX_iz = 0.63636256964372928
        E1NCX_blk = 0.19907097176889385

        I1NCX_blk = 0.16198650607989243
        I2NCX_blk = 0.63811996778245328
        P1_6_NaK = 0.55296589058788381
        P7_NaK = 0.08119993261668286
        P8_13_NaK = 0.098679856454751386

        TSCa3 = 0.0391776006630368
        TSCa3W = 0.0016262362338393967
        TSCa3S = 0.00078102706265326734
        TSS = 0.0018319921802936519
        TSW = 0.00096100174804855112

        hw = 0.0001
        hp = 0.006
        halfSL = 1

    End Sub

    Public Sub defaultSTARTN()
        Vm = -73.261962405254934
        Nai = 6.3544170921728274
        Ki = 141.85099855969611
        Catot_blk = 0.011217965556457833
        CaMCa = 0.00045210069073215771
        TnChCa = 0.0066458454471230357
        SRCa = 0.0032993814869566483
        Catot_iz = 0.0745020930812615
        Catot_jnc = 0.13666108625012649
        Ca_SRup = 1.485124250319698

        Catot_SRrl = 8.1430731604405526
        'Yooo = 0.00020253669428842405
        'Yooc = 0.00012586070783551491
        'Ycoo = 0.00017647414490221352
        'Ycoc = 0.99311947176338911
        'Ycco = 0.000000040324240633941138
        'Yoco = 0.000000007108662528364609
        'Yocc = 0.0000021186469054078025
        us_CaL = 0.99972007240039673
        Yco_iz = 0.99383496009503713

        Yoc_iz = 0.0000019371856741299545
        Yoo_iz = 0.00032858597201774521
        Yco_blk = 0.99384232454847854
        Yoc_blk = 0.0000019347508070066086
        Yoo_blk = 0.00032858840688486851
        O_TM = 0.0000005409268138235363
        I2_TM = 0.016654931481386005
        Is_TM = 0.83016496066441769
        O_LM = 0.0000023747125465770216
        I1_LM = 0.001370043404471766

        I2_LM = 0.016746782914056318
        Is_LM = 0.82915652611479584
        d_CaT = 0.00016261322228289861
        f_CaT = 0.44900248485290029
        d_st = 0.028583444490459756
        f_st = 0.96031474429165442
        us_st = 0.99862710093384377
        C1_ha = 0.97488226144001833
        C2_ha = 0.002820907457687366
        O1_ha = 0.0031484666642099626

        O2_ha = 0.0047793908174814971
        O_IK1 = 0.4651698
        Pbspm = 0.86962998614798059
        y1_IKr = 0.0045264981101403009
        y2_IKr = 0.09091112579532537
        y3_IKr = 0.9938018090199191
        ua = 0.0088451475850216781
        ui = 0.97091758813468176
        Ov_IKs = 0.0210874638535362
        C2IKs_iz = 0.009797980495198956

        OcIKs_iz = 0.019854801182127756
        C2IKs_blk = 0.0098010335871594571
        OcIKs_blk = 0.018859196820498303
        y1_IKto = 0.0025535181237643628
        y2_IKto = 0.65403199010083157
        O_ACh = 0.037606368462374695
        E1NCX_iz = 0.20432606394387906
        I1NCX_iz = 0.1584577127394789
        I2NCX_iz = 0.63636256964372928
        E1NCX_blk = 0.19907097176889385

        I1NCX_blk = 0.16198650607989243
        I2NCX_blk = 0.63811996778245328
        P1_6_NaK = 0.55296589058788381
        P7_NaK = 0.08119993261668286
        P8_13_NaK = 0.098679856454751386
        TSCa3 = 0.0391776006630368
        TSCa3W = 0.0016262362338393967
        TSCa3S = 0.00078102706265326734
        TSS = 0.0018319921802936519
        TSW = 0.00096100174804855112

        hw = 0.0001
        hp = 0.006
        halfSL = 1
    End Sub


End Class
```
