## Supplementary material for "Gradient-based parameter optimization to determine membrane ionic current composition of human induced pluripotent stem cell-derived cardiomyocytes": optimizing program: Application.Designer.vb

```
'------------------------------------------------------------------------------
' <auto-generated>
'     This code was generated by a tool.
'     Runtime Version:4.0.30319.42000
'
'     Changes to this file may cause incorrect behavior and will be lost if
'     the code is regenerated.
' </auto-generated>
'------------------------------------------------------------------------------

Option Strict On
Option Explicit On


Namespace My

    'NOTE: This file is auto-generated; do not modify it directly.  To make changes,
    ' or if you encounter build errors in this file, go to the Project Designer
    ' (go to Project Properties or double-click the My Project node in
    ' Solution Explorer), and make changes on the Application tab.
    '
    Partial Friend Class MyApplication

        <Global.System.Diagnostics.DebuggerStepThroughAttribute()> _
        Public Sub New()
            MyBase.New(Global.Microsoft.VisualBasic.ApplicationServices.AuthenticationMode.Windows)
            Me.IsSingleInstance = false
            Me.EnableVisualStyles = true
            Me.SaveMySettingsOnExit = true
            Me.ShutDownStyle = Global.Microsoft.VisualBasic.ApplicationServices.ShutdownMode.AfterMainFormCloses
        End Sub

        <Global.System.Diagnostics.DebuggerStepThroughAttribute()> _
        Protected Overrides Sub OnCreateMainForm()
            Me.MainForm = Global.ORMPEM.Protocol1
        End Sub
    End Class
End Namespace
```
