## Supplementary material for "Gradient-based parameter optimization to determine membrane ionic current composition of human induced pluripotent stem cell-derived cardiomyocytes": optimizing program: AssemblyInfo.vb

```
Imports System
Imports System.Reflection
Imports System.Runtime.InteropServices

' アセンブリに関する一般情報は以下の属性セットをとおして制御されます。
' アセンブリに関連付けられている情報を変更するには、
' これらの属性値を変更してください。

' アセンブリ属性の値を確認します

<Assembly: AssemblyTitle("ORMPEM")>
<Assembly: AssemblyDescription("")>
<Assembly: AssemblyCompany("")>
<Assembly: AssemblyProduct("ORMPEM")>
<Assembly: AssemblyCopyright("Copyright ©  2021")>
<Assembly: AssemblyTrademark("")>

<Assembly: ComVisible(False)>

'このプロジェクトが COM に公開される場合、次の GUID が typelib の ID になります
<Assembly: Guid("48e41f29-10be-4264-bc25-992c62f89dec")>

' アセンブリのバージョン情報は次の 4 つの値で構成されています:
'
'      メジャー バージョン
'      マイナー バージョン
'      ビルド番号
'      Revision
'
' すべての値を指定するか、次を使用してビルド番号とリビジョン番号を既定に設定できます
' 既定値にすることができます:
' <Assembly: AssemblyVersion("1.0.*")>

<Assembly: AssemblyVersion("1.0.0.0")>
<Assembly: AssemblyFileVersion("1.0.0.0")>
```
