## Supplementary material for "Gradient-based parameter optimization to determine membrane ionic current composition of human induced pluripotent stem cell-derived cardiomyocytes": optimizing program: Resources.Designer.vb

```
'------------------------------------------------------------------------------
' <auto-generated>
'     This code was generated by a tool.
'     Runtime Version:4.0.30319.42000
'
'     Changes to this file may cause incorrect behavior and will be lost if
'     the code is regenerated.
' </auto-generated>
'------------------------------------------------------------------------------

Option Strict On
Option Explicit On


Namespace My.Resources

    'This class was auto-generated by the StronglyTypedResourceBuilder
    'class via a tool like ResGen or Visual Studio.
    'To add or remove a member, edit your .ResX file then rerun ResGen
    'with the /str option, or rebuild your VS project.
    '''<summary>
    '''  A strongly-typed resource class, for looking up localized strings, etc.
    '''</summary>
    <Global.System.CodeDom.Compiler.GeneratedCodeAttribute("System.Resources.Tools.StronglyTypedResourceBuilder", "4.0.0.0"), _
     Global.System.Diagnostics.DebuggerNonUserCodeAttribute(), _
     Global.System.Runtime.CompilerServices.CompilerGeneratedAttribute(), _
     Global.Microsoft.VisualBasic.HideModuleNameAttribute()> _
    Friend Module Resources

        Private resourceMan As Global.System.Resources.ResourceManager

        Private resourceCulture As Global.System.Globalization.CultureInfo

        '''<summary>
        '''  Returns the cached ResourceManager instance used by this class.
        '''</summary>
        <Global.System.ComponentModel.EditorBrowsableAttribute(Global.System.ComponentModel.EditorBrowsableState.Advanced)> _
        Friend ReadOnly Property ResourceManager() As Global.System.Resources.ResourceManager
            Get
                If Object.ReferenceEquals(resourceMan, Nothing) Then
                    Dim temp As Global.System.Resources.ResourceManager = New Global.System.Resources.ResourceManager("ORMPEM.Resources", GetType(Resources).Assembly)
                    resourceMan = temp
                End If
                Return resourceMan
            End Get
        End Property

        '''<summary>
        '''  Overrides the current thread's CurrentUICulture property for all
        '''  resource lookups using this strongly typed resource class.
        '''</summary>
        <Global.System.ComponentModel.EditorBrowsableAttribute(Global.System.ComponentModel.EditorBrowsableState.Advanced)> _
        Friend Property Culture() As Global.System.Globalization.CultureInfo
            Get
                Return resourceCulture
            End Get
            Set(ByVal value As Global.System.Globalization.CultureInfo)
                resourceCulture = value
            End Set
        End Property
    End Module
End Namespace
```
