## Supplementary material for "Gradient-based parameter optimization to determine membrane ionic current composition of human induced pluripotent stem cell-derived cardiomyocytes": optimizing program: Settings.Designer.vb

```
'------------------------------------------------------------------------------
' <auto-generated>
'     This code was generated by a tool.
'     Runtime Version:4.0.30319.42000
'
'     Changes to this file may cause incorrect behavior and will be lost if
'     the code is regenerated.
' </auto-generated>
'------------------------------------------------------------------------------

Option Strict On
Option Explicit On


Namespace My

    <Global.System.Runtime.CompilerServices.CompilerGeneratedAttribute(), _
     Global.System.CodeDom.Compiler.GeneratedCodeAttribute("Microsoft.VisualStudio.Editors.SettingsDesigner.SettingsSingleFileGenerator", "11.0.0.0"), _
     Global.System.ComponentModel.EditorBrowsableAttribute(Global.System.ComponentModel.EditorBrowsableState.Advanced)> _
    Partial Friend NotInheritable Class MySettings
        Inherits Global.System.Configuration.ApplicationSettingsBase

        Private Shared defaultInstance As MySettings = CType(Global.System.Configuration.ApplicationSettingsBase.Synchronized(New MySettings), MySettings)

#Region "My.Settings Auto-Save Functionality"
#If _MyType = "WindowsForms" Then
        Private Shared addedHandler As Boolean

        Private Shared addedHandlerLockObject As New Object

        <Global.System.Diagnostics.DebuggerNonUserCodeAttribute(), Global.System.ComponentModel.EditorBrowsableAttribute(Global.System.ComponentModel.EditorBrowsableState.Advanced)> _
        Private Shared Sub AutoSaveSettings(ByVal sender As Global.System.Object, ByVal e As Global.System.EventArgs)
            If My.Application.SaveMySettingsOnExit Then
                My.Settings.Save()
            End If
        End Sub
#End If
#End Region

        Public Shared ReadOnly Property [Default]() As MySettings
            Get

#If _MyType = "WindowsForms" Then
                   If Not addedHandler Then
                        SyncLock addedHandlerLockObject
                            If Not addedHandler Then
                                AddHandler My.Application.Shutdown, AddressOf AutoSaveSettings
                                addedHandler = True
                            End If
                        End SyncLock
                    End If
#End If
                Return defaultInstance
            End Get
        End Property
    End Class
End Namespace

Namespace My

    <Global.Microsoft.VisualBasic.HideModuleNameAttribute(), _
     Global.System.Diagnostics.DebuggerNonUserCodeAttribute(), _
     Global.System.Runtime.CompilerServices.CompilerGeneratedAttribute()> _
    Friend Module MySettingsProperty

        <Global.System.ComponentModel.Design.HelpKeywordAttribute("My.Settings")> _
        Friend ReadOnly Property Settings() As Global.ORMPEM.My.MySettings
            Get
                Return Global.ORMPEM.My.MySettings.Default
            End Get
        End Property
    End Module
End Namespace
```
