## Supplementary material for "Gradient-based parameter optimization to determine membrane ionic current composition of human induced pluripotent stem cell-derived cardiomyocytes": optimizing program: Protocol1.Designer.vb

```
<Global.Microsoft.VisualBasic.CompilerServices.DesignerGenerated()>
Partial Class Protocol1
    Inherits System.Windows.Forms.Form

    'フォームがコンポーネントの一覧をクリーンアップするために dispose をオーバーライドします。
    <System.Diagnostics.DebuggerNonUserCode()>
    Protected Overrides Sub Dispose(ByVal disposing As Boolean)
        Try
            If disposing AndAlso components IsNot Nothing Then
                components.Dispose()
            End If
        Finally
            MyBase.Dispose(disposing)
        End Try
    End Sub

    'Windows フォーム デザイナーで必要です。
    Private components As System.ComponentModel.IContainer

    'メモ: 以下のプロシージャは Windows フォーム デザイナーで必要です。
    'Windows フォーム デザイナーを使用して変更できます。  
    'コード エディターを使って変更しないでください。
    <System.Diagnostics.DebuggerStepThrough()>
    Private Sub InitializeComponent()
        Me.Label4 = New System.Windows.Forms.Label()
        Me.cbEnd = New System.Windows.Forms.Button()
        Me.pbVm1 = New System.Windows.Forms.PictureBox()
        Me.cbSelect = New System.Windows.Forms.Button()
        Me.cbPlot = New System.Windows.Forms.Button()
        Me.cbCsvFileList = New System.Windows.Forms.ComboBox()
        Me.cbselectCSVfile = New System.Windows.Forms.Button()
        Me.cbPS = New System.Windows.Forms.Button()
        Me.cbManualFit = New System.Windows.Forms.Button()
        Me.GroupBox3 = New System.Windows.Forms.GroupBox()
        Me.chkRand2 = New System.Windows.Forms.CheckBox()
        Me.Label36 = New System.Windows.Forms.Label()
        Me.txtadr4R = New System.Windows.Forms.TextBox()
        Me.Label33 = New System.Windows.Forms.Label()
        Me.txtWeight = New System.Windows.Forms.TextBox()
        Me.Label78 = New System.Windows.Forms.Label()
        Me.txtPrecision = New System.Windows.Forms.TextBox()
        Me.Label77 = New System.Windows.Forms.Label()
        Me.txtScf2 = New System.Windows.Forms.TextBox()
        Me.TextBox2 = New System.Windows.Forms.TextBox()
        Me.TextBox3 = New System.Windows.Forms.TextBox()
        Me.Label64 = New System.Windows.Forms.Label()
        Me.Label72 = New System.Windows.Forms.Label()
        Me.txtScf1 = New System.Windows.Forms.TextBox()
        Me.Label44 = New System.Windows.Forms.Label()
        Me.chkNaKRegulation = New System.Windows.Forms.CheckBox()
        Me.chkWriteFile = New System.Windows.Forms.CheckBox()
        Me.chkDefault = New System.Windows.Forms.CheckBox()
        Me.Label17 = New System.Windows.Forms.Label()
        Me.txtNP = New System.Windows.Forms.TextBox()
        Me.txtStimulus = New System.Windows.Forms.TextBox()
        Me.txtEndms = New System.Windows.Forms.TextBox()
        Me.txt4th = New System.Windows.Forms.TextBox()
        Me.txt3rd = New System.Windows.Forms.TextBox()
        Me.txtStartms = New System.Windows.Forms.TextBox()
        Me.Label52 = New System.Windows.Forms.Label()
        Me.txt2nd = New System.Windows.Forms.TextBox()
        Me.Label16 = New System.Windows.Forms.Label()
        Me.txtEval = New System.Windows.Forms.TextBox()
        Me.txt1st = New System.Windows.Forms.TextBox()
        Me.txtRgR = New System.Windows.Forms.TextBox()
        Me.txtRgL = New System.Windows.Forms.TextBox()
        Me.txtEvalMax = New System.Windows.Forms.TextBox()
        Me.Label50 = New System.Windows.Forms.Label()
        Me.Label19 = New System.Windows.Forms.Label()
        Me.txtMaxRepeat = New System.Windows.Forms.TextBox()
        Me.Label11 = New System.Windows.Forms.Label()
        Me.Label7 = New System.Windows.Forms.Label()
        Me.Label26 = New System.Windows.Forms.Label()
        Me.controlBox4 = New System.Windows.Forms.GroupBox()
        Me.Label80 = New System.Windows.Forms.Label()
        Me.Label79 = New System.Windows.Forms.Label()
        Me.txt2R = New System.Windows.Forms.TextBox()
        Me.txtR3 = New System.Windows.Forms.TextBox()
        Me.txtR1 = New System.Windows.Forms.TextBox()
        Me.txtF4 = New System.Windows.Forms.TextBox()
        Me.txtF3 = New System.Windows.Forms.TextBox()
        Me.txtF2 = New System.Windows.Forms.TextBox()
        Me.txtF1 = New System.Windows.Forms.TextBox()
        Me.Label2 = New System.Windows.Forms.Label()
        Me.Label71 = New System.Windows.Forms.Label()
        Me.Label70 = New System.Windows.Forms.Label()
        Me.Label56 = New System.Windows.Forms.Label()
        Me.txtNai = New System.Windows.Forms.TextBox()
        Me.chkIbNSC = New System.Windows.Forms.CheckBox()
        Me.txtINCXmag = New System.Windows.Forms.TextBox()
        Me.txtINaKmag = New System.Windows.Forms.TextBox()
        Me.chkINCX = New System.Windows.Forms.CheckBox()
        Me.chkINaK = New System.Windows.Forms.CheckBox()
        Me.chkIst = New System.Windows.Forms.CheckBox()
        Me.chkIKto = New System.Windows.Forms.CheckBox()
        Me.chkIKs = New System.Windows.Forms.CheckBox()
        Me.chkIha = New System.Windows.Forms.CheckBox()
        Me.chkIKur = New System.Windows.Forms.CheckBox()
        Me.chkINa = New System.Windows.Forms.CheckBox()
        Me.chkICaL = New System.Windows.Forms.CheckBox()
        Me.chkIK1 = New System.Windows.Forms.CheckBox()
        Me.chkIKr = New System.Windows.Forms.CheckBox()
        Me.NFusCaLmag = New System.Windows.Forms.TextBox()
        Me.rFusCaL = New System.Windows.Forms.TextBox()
        Me.Label60 = New System.Windows.Forms.Label()
        Me.txtstdtotCai = New System.Windows.Forms.TextBox()
        Me.txtstdNai = New System.Windows.Forms.TextBox()
        Me.Label59 = New System.Windows.Forms.Label()
        Me.txtFusCaLmag = New System.Windows.Forms.TextBox()
        Me.txtVshiftNa = New System.Windows.Forms.TextBox()
        Me.Label23 = New System.Windows.Forms.Label()
        Me.Label3 = New System.Windows.Forms.Label()
        Me.Label5 = New System.Windows.Forms.Label()
        Me.txtVshiftCaL = New System.Windows.Forms.TextBox()
        Me.Label43 = New System.Windows.Forms.Label()
        Me.Label35 = New System.Windows.Forms.Label()
        Me.txtKLmag = New System.Windows.Forms.TextBox()
        Me.Label25 = New System.Windows.Forms.Label()
        Me.Label30 = New System.Windows.Forms.Label()
        Me.IKr = New System.Windows.Forms.Label()
        Me.txtICaLmag = New System.Windows.Forms.TextBox()
        Me.txtIK1mag = New System.Windows.Forms.TextBox()
        Me.Label15 = New System.Windows.Forms.Label()
        Me.txtIKrmag = New System.Windows.Forms.TextBox()
        Me.Label53 = New System.Windows.Forms.Label()
        Me.txtNaLratio = New System.Windows.Forms.TextBox()
        Me.Label51 = New System.Windows.Forms.Label()
        Me.Label6 = New System.Windows.Forms.Label()
        Me.txtBaseVshift = New System.Windows.Forms.TextBox()
        Me.Label48 = New System.Windows.Forms.Label()
        Me.Label9 = New System.Windows.Forms.Label()
        Me.txtorgExtraVshift = New System.Windows.Forms.TextBox()
        Me.txtICaTmag = New System.Windows.Forms.TextBox()
        Me.Label34 = New System.Windows.Forms.Label()
        Me.txtIKurmag = New System.Windows.Forms.TextBox()
        Me.txtOS = New System.Windows.Forms.TextBox()
        Me.Label10 = New System.Windows.Forms.Label()
        Me.Label13 = New System.Windows.Forms.Label()
        Me.Label8 = New System.Windows.Forms.Label()
        Me.txtIKsmag = New System.Windows.Forms.TextBox()
        Me.txtMDP = New System.Windows.Forms.TextBox()
        Me.Label27 = New System.Windows.Forms.Label()
        Me.Label14 = New System.Windows.Forms.Label()
        Me.rNa = New System.Windows.Forms.TextBox()
        Me.Label1 = New System.Windows.Forms.Label()
        Me.Label18 = New System.Windows.Forms.Label()
        Me.txtCycleL = New System.Windows.Forms.TextBox()
        Me.Label28 = New System.Windows.Forms.Label()
        Me.txtIKtomag = New System.Windows.Forms.TextBox()
        Me.txtAPD = New System.Windows.Forms.TextBox()
        Me.rKr = New System.Windows.Forms.TextBox()
        Me.txtINamag = New System.Windows.Forms.TextBox()
        Me.txtTotalCai = New System.Windows.Forms.TextBox()
        Me.rK1 = New System.Windows.Forms.TextBox()
        Me.txtcrfNCX = New System.Windows.Forms.TextBox()
        Me.Label22 = New System.Windows.Forms.Label()
        Me.rVshiftCaL = New System.Windows.Forms.TextBox()
        Me.txtcrfNaK = New System.Windows.Forms.TextBox()
        Me.rCaL = New System.Windows.Forms.TextBox()
        Me.txtIbNSCmag = New System.Windows.Forms.TextBox()
        Me.Label24 = New System.Windows.Forms.Label()
        Me.txtKi = New System.Windows.Forms.TextBox()
        Me.rCaT = New System.Windows.Forms.TextBox()
        Me.Label29 = New System.Windows.Forms.Label()
        Me.rextraVshift = New System.Windows.Forms.TextBox()
        Me.rst = New System.Windows.Forms.TextBox()
        Me.txtIhamag = New System.Windows.Forms.TextBox()
        Me.rbNSC = New System.Windows.Forms.TextBox()
        Me.NVshiftCaL = New System.Windows.Forms.TextBox()
        Me.Label47 = New System.Windows.Forms.Label()
        Me.rKL = New System.Windows.Forms.TextBox()
        Me.NextraVshift = New System.Windows.Forms.TextBox()
        Me.rKto = New System.Windows.Forms.TextBox()
        Me.NVshiftNa = New System.Windows.Forms.TextBox()
        Me.rVshiftNa = New System.Windows.Forms.TextBox()
        Me.txtIstmag = New System.Windows.Forms.TextBox()
        Me.NIhamag = New System.Windows.Forms.TextBox()
        Me.rKs = New System.Windows.Forms.TextBox()
        Me.NICaTmag = New System.Windows.Forms.TextBox()
        Me.rha = New System.Windows.Forms.TextBox()
        Me.NIKurmag = New System.Windows.Forms.TextBox()
        Me.rKur = New System.Windows.Forms.TextBox()
        Me.Label20 = New System.Windows.Forms.Label()
        Me.NIKsmag = New System.Windows.Forms.TextBox()
        Me.NIKtomag = New System.Windows.Forms.TextBox()
        Me.NIstmag = New System.Windows.Forms.TextBox()
        Me.Label21 = New System.Windows.Forms.Label()
        Me.NKLmag = New System.Windows.Forms.TextBox()
        Me.NINamag = New System.Windows.Forms.TextBox()
        Me.NIbNSCmag = New System.Windows.Forms.TextBox()
        Me.NIKrmag = New System.Windows.Forms.TextBox()
        Me.NIK1mag = New System.Windows.Forms.TextBox()
        Me.NICaLmag = New System.Windows.Forms.TextBox()
        Me.Label55 = New System.Windows.Forms.Label()
        Me.chkVmadaptive = New System.Windows.Forms.CheckBox()
        Me.chkVmreference = New System.Windows.Forms.CheckBox()
        Me.pbInwardI = New System.Windows.Forms.PictureBox()
        Me.txtVmF = New System.Windows.Forms.TextBox()
        Me.Label40 = New System.Windows.Forms.Label()
        Me.Label38 = New System.Windows.Forms.Label()
        Me.pbpsVm = New System.Windows.Forms.PictureBox()
        Me.pbOutwardI = New System.Windows.Forms.PictureBox()
        Me.Label31 = New System.Windows.Forms.Label()
        Me.pbBP5 = New System.Windows.Forms.PictureBox()
        Me.Label68 = New System.Windows.Forms.Label()
        Me.txtMeanBP4 = New System.Windows.Forms.TextBox()
        Me.Label67 = New System.Windows.Forms.Label()
        Me.Label42 = New System.Windows.Forms.Label()
        Me.Label54 = New System.Windows.Forms.Label()
        Me.pbBP4 = New System.Windows.Forms.PictureBox()
        Me.TextBox1 = New System.Windows.Forms.TextBox()
        Me.txtMeanBP2 = New System.Windows.Forms.TextBox()
        Me.txtMeanBP1 = New System.Windows.Forms.TextBox()
        Me.Label41 = New System.Windows.Forms.Label()
        Me.Label49 = New System.Windows.Forms.Label()
        Me.Label12 = New System.Windows.Forms.Label()
        Me.Label61 = New System.Windows.Forms.Label()
        Me.pbBP1 = New System.Windows.Forms.PictureBox()
        Me.pbPB2 = New System.Windows.Forms.PictureBox()
        Me.Label66 = New System.Windows.Forms.Label()
        Me.txtMeanBP5 = New System.Windows.Forms.TextBox()
        Me.Label45 = New System.Windows.Forms.Label()
        Me.Label37 = New System.Windows.Forms.Label()
        Me.txtStp = New System.Windows.Forms.TextBox()
        Me.Label69 = New System.Windows.Forms.Label()
        Me.txtdetectINa = New System.Windows.Forms.TextBox()
        Me.Label65 = New System.Windows.Forms.Label()
        Me.txtPScycle = New System.Windows.Forms.TextBox()
        Me.Label32 = New System.Windows.Forms.Label()
        Me.txtNBPSQS = New System.Windows.Forms.TextBox()
        Me.txtLogMSE = New System.Windows.Forms.TextBox()
        Me.pbBP6 = New System.Windows.Forms.PictureBox()
        Me.Label58 = New System.Windows.Forms.Label()
        Me.txtMSE = New System.Windows.Forms.TextBox()
        Me.pbSQS = New System.Windows.Forms.PictureBox()
        Me.Label46 = New System.Windows.Forms.Label()
        Me.Label62 = New System.Windows.Forms.Label()
        Me.txtCellNo = New System.Windows.Forms.TextBox()
        Me.txtMeanBP6 = New System.Windows.Forms.TextBox()
        Me.Label74 = New System.Windows.Forms.Label()
        Me.pbBP3 = New System.Windows.Forms.PictureBox()
        Me.Label75 = New System.Windows.Forms.Label()
        Me.txtR2 = New System.Windows.Forms.TextBox()
        Me.Label76 = New System.Windows.Forms.Label()
        Me.txtMeanBP3 = New System.Windows.Forms.TextBox()
        Me.txtI1 = New System.Windows.Forms.TextBox()
        Me.txtI2 = New System.Windows.Forms.TextBox()
        Me.txtI3 = New System.Windows.Forms.TextBox()
        Me.txtI4 = New System.Windows.Forms.TextBox()
        Me.txtI5 = New System.Windows.Forms.TextBox()
        Me.txtI6 = New System.Windows.Forms.TextBox()
        Me.txtVmOnset = New System.Windows.Forms.TextBox()
        Me.txtError = New System.Windows.Forms.TextBox()
        Me.Label39 = New System.Windows.Forms.Label()
        Me.Label57 = New System.Windows.Forms.Label()
        Me.txtSbp1 = New System.Windows.Forms.TextBox()
        Me.txtSbp2 = New System.Windows.Forms.TextBox()
        Me.txtSbp3 = New System.Windows.Forms.TextBox()
        Me.txtSbp4 = New System.Windows.Forms.TextBox()
        Me.txtSbp5 = New System.Windows.Forms.TextBox()
        Me.txtSbp6 = New System.Windows.Forms.TextBox()
        Me.pbSensitivity = New System.Windows.Forms.PictureBox()
        Me.Label63 = New System.Windows.Forms.Label()
        Me.Label73 = New System.Windows.Forms.Label()
        Me.Label81 = New System.Windows.Forms.Label()
        Me.Label82 = New System.Windows.Forms.Label()
        Me.Label83 = New System.Windows.Forms.Label()
        Me.Label84 = New System.Windows.Forms.Label()
        Me.Label85 = New System.Windows.Forms.Label()
        Me.txtCLB = New System.Windows.Forms.TextBox()
        Me.txtPreviousMIN = New System.Windows.Forms.TextBox()
        Me.Label86 = New System.Windows.Forms.Label()
        Me.rbEMTest = New System.Windows.Forms.RadioButton()
        Me.rbMMtest = New System.Windows.Forms.RadioButton()
        CType(Me.pbVm1, System.ComponentModel.ISupportInitialize).BeginInit()
        Me.GroupBox3.SuspendLayout()
        Me.controlBox4.SuspendLayout()
        CType(Me.pbInwardI, System.ComponentModel.ISupportInitialize).BeginInit()
        CType(Me.pbpsVm, System.ComponentModel.ISupportInitialize).BeginInit()
        CType(Me.pbOutwardI, System.ComponentModel.ISupportInitialize).BeginInit()
        CType(Me.pbBP5, System.ComponentModel.ISupportInitialize).BeginInit()
        CType(Me.pbBP4, System.ComponentModel.ISupportInitialize).BeginInit()
        CType(Me.pbBP1, System.ComponentModel.ISupportInitialize).BeginInit()
        CType(Me.pbPB2, System.ComponentModel.ISupportInitialize).BeginInit()
        CType(Me.pbBP6, System.ComponentModel.ISupportInitialize).BeginInit()
        CType(Me.pbSQS, System.ComponentModel.ISupportInitialize).BeginInit()
        CType(Me.pbBP3, System.ComponentModel.ISupportInitialize).BeginInit()
        CType(Me.pbSensitivity, System.ComponentModel.ISupportInitialize).BeginInit()
        Me.SuspendLayout()
        '
        'Label4
        '
        Me.Label4.AutoSize = True
        Me.Label4.BackColor = System.Drawing.Color.White
        Me.Label4.Font = New System.Drawing.Font("Verdana", 14.25!, System.Drawing.FontStyle.Regular, System.Drawing.GraphicsUnit.Point, CType(0, Byte))
        Me.Label4.ForeColor = System.Drawing.Color.Red
        Me.Label4.Location = New System.Drawing.Point(113, 25)
        Me.Label4.Name = "Label4"
        Me.Label4.Size = New System.Drawing.Size(22, 23)
        Me.Label4.TabIndex = 932
        Me.Label4.Text = "2"
        '
        'cbEnd
        '
        Me.cbEnd.BackColor = System.Drawing.SystemColors.ButtonHighlight
        Me.cbEnd.Font = New System.Drawing.Font("Verdana", 15.75!, System.Drawing.FontStyle.Regular, System.Drawing.GraphicsUnit.Point, CType(0, Byte))
        Me.cbEnd.ForeColor = System.Drawing.Color.Blue
        Me.cbEnd.Location = New System.Drawing.Point(8, 79)
        Me.cbEnd.Name = "cbEnd"
        Me.cbEnd.Size = New System.Drawing.Size(114, 42)
        Me.cbEnd.TabIndex = 931
        Me.cbEnd.Text = "End"
        Me.cbEnd.UseVisualStyleBackColor = False
        '
        'pbVm1
        '
        Me.pbVm1.BackColor = System.Drawing.SystemColors.ButtonHighlight
        Me.pbVm1.Location = New System.Drawing.Point(837, 13)
        Me.pbVm1.Name = "pbVm1"
        Me.pbVm1.Size = New System.Drawing.Size(1008, 166)
        Me.pbVm1.TabIndex = 929
        Me.pbVm1.TabStop = False
        '
        'cbSelect
        '
        Me.cbSelect.BackColor = System.Drawing.SystemColors.ButtonHighlight
        Me.cbSelect.Font = New System.Drawing.Font("Verdana", 15.75!, System.Drawing.FontStyle.Regular, System.Drawing.GraphicsUnit.Point, CType(0, Byte))
        Me.cbSelect.ForeColor = System.Drawing.Color.Red
        Me.cbSelect.Location = New System.Drawing.Point(416, -2)
        Me.cbSelect.Name = "cbSelect"
        Me.cbSelect.Size = New System.Drawing.Size(129, 41)
        Me.cbSelect.TabIndex = 928
        Me.cbSelect.Text = "4 Select"
        Me.cbSelect.UseVisualStyleBackColor = False
        Me.cbSelect.Visible = False
        '
        'cbPlot
        '
        Me.cbPlot.BackColor = System.Drawing.SystemColors.ButtonHighlight
        Me.cbPlot.Font = New System.Drawing.Font("Verdana", 15.75!, System.Drawing.FontStyle.Regular, System.Drawing.GraphicsUnit.Point, CType(0, Byte))
        Me.cbPlot.ForeColor = System.Drawing.Color.Red
        Me.cbPlot.Location = New System.Drawing.Point(305, -2)
        Me.cbPlot.Name = "cbPlot"
        Me.cbPlot.Size = New System.Drawing.Size(114, 41)
        Me.cbPlot.TabIndex = 927
        Me.cbPlot.Text = "3 Plot"
        Me.cbPlot.UseVisualStyleBackColor = False
        Me.cbPlot.Visible = False
        '
        'cbCsvFileList
        '
        Me.cbCsvFileList.Font = New System.Drawing.Font("Verdana", 11.25!, System.Drawing.FontStyle.Regular, System.Drawing.GraphicsUnit.Point, CType(0, Byte))
        Me.cbCsvFileList.FormattingEnabled = True
        Me.cbCsvFileList.Location = New System.Drawing.Point(104, 3)
        Me.cbCsvFileList.Name = "cbCsvFileList"
        Me.cbCsvFileList.Size = New System.Drawing.Size(204, 26)
        Me.cbCsvFileList.TabIndex = 926
        '
        'cbselectCSVfile
        '
        Me.cbselectCSVfile.BackColor = System.Drawing.SystemColors.ButtonHighlight
        Me.cbselectCSVfile.Font = New System.Drawing.Font("Verdana", 12.0!, System.Drawing.FontStyle.Regular, System.Drawing.GraphicsUnit.Point, CType(0, Byte))
        Me.cbselectCSVfile.ForeColor = System.Drawing.Color.Red
        Me.cbselectCSVfile.Location = New System.Drawing.Point(6, 0)
        Me.cbselectCSVfile.Name = "cbselectCSVfile"
        Me.cbselectCSVfile.Size = New System.Drawing.Size(95, 48)
        Me.cbselectCSVfile.TabIndex = 925
        Me.cbselectCSVfile.Text = " 1 Select csvFile"
        Me.cbselectCSVfile.UseVisualStyleBackColor = False
        '
        'cbPS
        '
        Me.cbPS.BackColor = System.Drawing.Color.White
        Me.cbPS.Font = New System.Drawing.Font("Verdana", 15.75!, System.Drawing.FontStyle.Regular, System.Drawing.GraphicsUnit.Point, CType(0, Byte))
        Me.cbPS.ForeColor = System.Drawing.Color.Blue
        Me.cbPS.Location = New System.Drawing.Point(704, -2)
        Me.cbPS.Name = "cbPS"
        Me.cbPS.Size = New System.Drawing.Size(191, 45)
        Me.cbPS.TabIndex = 924
        Me.cbPS.Text = "6 PatternSearch"
        Me.cbPS.UseVisualStyleBackColor = False
        Me.cbPS.Visible = False
        '
        'cbManualFit
        '
        Me.cbManualFit.BackColor = System.Drawing.Color.White
        Me.cbManualFit.Font = New System.Drawing.Font("Verdana", 15.75!, System.Drawing.FontStyle.Regular, System.Drawing.GraphicsUnit.Point, CType(0, Byte))
        Me.cbManualFit.ForeColor = System.Drawing.Color.Red
        Me.cbManualFit.Location = New System.Drawing.Point(543, 0)
        Me.cbManualFit.Name = "cbManualFit"
        Me.cbManualFit.Size = New System.Drawing.Size(166, 35)
        Me.cbManualFit.TabIndex = 923
        Me.cbManualFit.Text = "5 manual  Fit"
        Me.cbManualFit.UseVisualStyleBackColor = False
        Me.cbManualFit.Visible = False
        '
        'GroupBox3
        '
        Me.GroupBox3.BackColor = System.Drawing.Color.FromArgb(CType(CType(220, Byte), Integer), CType(CType(255, Byte), Integer), CType(CType(255, Byte), Integer))
        Me.GroupBox3.Controls.Add(Me.chkRand2)
        Me.GroupBox3.Controls.Add(Me.Label36)
        Me.GroupBox3.Controls.Add(Me.txtadr4R)
        Me.GroupBox3.Controls.Add(Me.Label33)
        Me.GroupBox3.Controls.Add(Me.txtWeight)
        Me.GroupBox3.Controls.Add(Me.Label78)
        Me.GroupBox3.Controls.Add(Me.txtPrecision)
        Me.GroupBox3.Controls.Add(Me.Label77)
        Me.GroupBox3.Controls.Add(Me.txtScf2)
        Me.GroupBox3.Controls.Add(Me.TextBox2)
        Me.GroupBox3.Controls.Add(Me.TextBox3)
        Me.GroupBox3.Controls.Add(Me.Label64)
        Me.GroupBox3.Controls.Add(Me.Label72)
        Me.GroupBox3.Controls.Add(Me.txtScf1)
        Me.GroupBox3.Controls.Add(Me.Label44)
        Me.GroupBox3.Controls.Add(Me.chkNaKRegulation)
        Me.GroupBox3.Controls.Add(Me.chkWriteFile)
        Me.GroupBox3.Controls.Add(Me.chkDefault)
        Me.GroupBox3.Controls.Add(Me.Label17)
        Me.GroupBox3.Controls.Add(Me.txtNP)
        Me.GroupBox3.Controls.Add(Me.txtStimulus)
        Me.GroupBox3.Controls.Add(Me.txtEndms)
        Me.GroupBox3.Controls.Add(Me.txt4th)
        Me.GroupBox3.Controls.Add(Me.txt3rd)
        Me.GroupBox3.Controls.Add(Me.txtStartms)
        Me.GroupBox3.Controls.Add(Me.Label52)
        Me.GroupBox3.Controls.Add(Me.txt2nd)
        Me.GroupBox3.Controls.Add(Me.Label16)
        Me.GroupBox3.Controls.Add(Me.txtEval)
        Me.GroupBox3.Controls.Add(Me.txt1st)
        Me.GroupBox3.Controls.Add(Me.txtRgR)
        Me.GroupBox3.Controls.Add(Me.txtRgL)
        Me.GroupBox3.Controls.Add(Me.txtEvalMax)
        Me.GroupBox3.Controls.Add(Me.Label50)
        Me.GroupBox3.Controls.Add(Me.Label19)
        Me.GroupBox3.Controls.Add(Me.txtMaxRepeat)
        Me.GroupBox3.Controls.Add(Me.Label11)
        Me.GroupBox3.Controls.Add(Me.Label7)
        Me.GroupBox3.Controls.Add(Me.Label26)
        Me.GroupBox3.Font = New System.Drawing.Font("MS UI Gothic", 8.25!, System.Drawing.FontStyle.Regular, System.Drawing.GraphicsUnit.Point, CType(128, Byte))
        Me.GroupBox3.Location = New System.Drawing.Point(3, 127)
        Me.GroupBox3.Name = "GroupBox3"
        Me.GroupBox3.Size = New System.Drawing.Size(284, 257)
        Me.GroupBox3.TabIndex = 933
        Me.GroupBox3.TabStop = False
        Me.GroupBox3.Text = "00"
        '
        'chkRand2
        '
        Me.chkRand2.AutoSize = True
        Me.chkRand2.Checked = True
        Me.chkRand2.CheckState = System.Windows.Forms.CheckState.Checked
        Me.chkRand2.Font = New System.Drawing.Font("Verdana", 9.0!, System.Drawing.FontStyle.Regular, System.Drawing.GraphicsUnit.Point, CType(0, Byte))
        Me.chkRand2.Location = New System.Drawing.Point(195, 14)
        Me.chkRand2.Name = "chkRand2"
        Me.chkRand2.Size = New System.Drawing.Size(66, 18)
        Me.chkRand2.TabIndex = 1028
        Me.chkRand2.Text = "Rand2"
        Me.chkRand2.TextImageRelation = System.Windows.Forms.TextImageRelation.TextAboveImage
        Me.chkRand2.UseVisualStyleBackColor = True
        '
        'Label36
        '
        Me.Label36.AutoSize = True
        Me.Label36.Font = New System.Drawing.Font("Verdana", 9.75!, System.Drawing.FontStyle.Regular, System.Drawing.GraphicsUnit.Point, CType(0, Byte))
        Me.Label36.Location = New System.Drawing.Point(119, 111)
        Me.Label36.Name = "Label36"
        Me.Label36.Size = New System.Drawing.Size(100, 16)
        Me.Label36.TabIndex = 940
        Me.Label36.Text = "adr4R (730) ="
        '
        'txtadr4R
        '
        Me.txtadr4R.AccessibleDescription = "370"
        Me.txtadr4R.Font = New System.Drawing.Font("Verdana", 9.75!, System.Drawing.FontStyle.Regular, System.Drawing.GraphicsUnit.Point, CType(0, Byte))
        Me.txtadr4R.Location = New System.Drawing.Point(219, 107)
        Me.txtadr4R.Name = "txtadr4R"
        Me.txtadr4R.Size = New System.Drawing.Size(59, 23)
        Me.txtadr4R.TabIndex = 939
        Me.txtadr4R.Tag = ""
        Me.txtadr4R.Text = "730"
        '
        'Label33
        '
        Me.Label33.AutoSize = True
        Me.Label33.Font = New System.Drawing.Font("Verdana", 9.75!, System.Drawing.FontStyle.Regular, System.Drawing.GraphicsUnit.Point, CType(0, Byte))
        Me.Label33.Location = New System.Drawing.Point(1, 109)
        Me.Label33.Name = "Label33"
        Me.Label33.Size = New System.Drawing.Size(78, 16)
        Me.Label33.TabIndex = 938
        Me.Label33.Text = "weightIx ="
        '
        'txtWeight
        '
        Me.txtWeight.Font = New System.Drawing.Font("Verdana", 9.75!, System.Drawing.FontStyle.Regular, System.Drawing.GraphicsUnit.Point, CType(0, Byte))
        Me.txtWeight.Location = New System.Drawing.Point(79, 107)
        Me.txtWeight.Name = "txtWeight"
        Me.txtWeight.Size = New System.Drawing.Size(38, 23)
        Me.txtWeight.TabIndex = 937
        Me.txtWeight.Tag = ""
        '
        'Label78
        '
        Me.Label78.AutoSize = True
        Me.Label78.Font = New System.Drawing.Font("Verdana", 9.75!, System.Drawing.FontStyle.Regular, System.Drawing.GraphicsUnit.Point, CType(0, Byte))
        Me.Label78.Location = New System.Drawing.Point(6, 82)
        Me.Label78.Name = "Label78"
        Me.Label78.Size = New System.Drawing.Size(66, 16)
        Me.Label78.TabIndex = 936
        Me.Label78.Text = "precision"
        '
        'txtPrecision
        '
        Me.txtPrecision.Font = New System.Drawing.Font("Verdana", 9.75!, System.Drawing.FontStyle.Regular, System.Drawing.GraphicsUnit.Point, CType(0, Byte))
        Me.txtPrecision.Location = New System.Drawing.Point(79, 80)
        Me.txtPrecision.Name = "txtPrecision"
        Me.txtPrecision.Size = New System.Drawing.Size(95, 23)
        Me.txtPrecision.TabIndex = 935
        Me.txtPrecision.Tag = "0.3"
        Me.txtPrecision.Text = "0.00001"
        '
        'Label77
        '
        Me.Label77.AutoSize = True
        Me.Label77.Font = New System.Drawing.Font("Verdana", 9.75!, System.Drawing.FontStyle.Regular, System.Drawing.GraphicsUnit.Point, CType(0, Byte))
        Me.Label77.Location = New System.Drawing.Point(156, 58)
        Me.Label77.Name = "Label77"
        Me.Label77.Size = New System.Drawing.Size(69, 16)
        Me.Label77.TabIndex = 932
        Me.Label77.Text = "random 2"
        '
        'txtScf2
        '
        Me.txtScf2.Font = New System.Drawing.Font("Verdana", 9.75!, System.Drawing.FontStyle.Regular, System.Drawing.GraphicsUnit.Point, CType(0, Byte))
        Me.txtScf2.Location = New System.Drawing.Point(229, 56)
        Me.txtScf2.Name = "txtScf2"
        Me.txtScf2.Size = New System.Drawing.Size(49, 23)
        Me.txtScf2.TabIndex = 931
        Me.txtScf2.Tag = "0.3"
        Me.txtScf2.Text = "0.015"
        '
        'TextBox2
        '
        Me.TextBox2.Font = New System.Drawing.Font("Verdana", 9.0!, System.Drawing.FontStyle.Regular, System.Drawing.GraphicsUnit.Point, CType(0, Byte))
        Me.TextBox2.Location = New System.Drawing.Point(147, 234)
        Me.TextBox2.Name = "TextBox2"
        Me.TextBox2.Size = New System.Drawing.Size(28, 22)
        Me.TextBox2.TabIndex = 922
        Me.TextBox2.Text = "6"
        '
        'TextBox3
        '
        Me.TextBox3.Font = New System.Drawing.Font("Verdana", 9.0!, System.Drawing.FontStyle.Regular, System.Drawing.GraphicsUnit.Point, CType(0, Byte))
        Me.TextBox3.Location = New System.Drawing.Point(119, 234)
        Me.TextBox3.Name = "TextBox3"
        Me.TextBox3.Size = New System.Drawing.Size(28, 22)
        Me.TextBox3.TabIndex = 921
        Me.TextBox3.Text = "5"
        '
        'Label64
        '
        Me.Label64.AutoSize = True
        Me.Label64.Font = New System.Drawing.Font("Verdana", 9.75!, System.Drawing.FontStyle.Regular, System.Drawing.GraphicsUnit.Point, CType(0, Byte))
        Me.Label64.Location = New System.Drawing.Point(13, 218)
        Me.Label64.Name = "Label64"
        Me.Label64.Size = New System.Drawing.Size(155, 16)
        Me.Label64.TabIndex = 919
        Me.Label64.Text = "randomized sequence "
        '
        'Label72
        '
        Me.Label72.AutoSize = True
        Me.Label72.Font = New System.Drawing.Font("Verdana", 9.75!, System.Drawing.FontStyle.Regular, System.Drawing.GraphicsUnit.Point, CType(0, Byte))
        Me.Label72.Location = New System.Drawing.Point(8, 56)
        Me.Label72.Name = "Label72"
        Me.Label72.Size = New System.Drawing.Size(89, 16)
        Me.Label72.TabIndex = 920
        Me.Label72.Text = "random scf1"
        '
        'txtScf1
        '
        Me.txtScf1.Font = New System.Drawing.Font("Verdana", 9.75!, System.Drawing.FontStyle.Regular, System.Drawing.GraphicsUnit.Point, CType(0, Byte))
        Me.txtScf1.Location = New System.Drawing.Point(104, 54)
        Me.txtScf1.Name = "txtScf1"
        Me.txtScf1.Size = New System.Drawing.Size(42, 23)
        Me.txtScf1.TabIndex = 919
        Me.txtScf1.Tag = ""
        Me.txtScf1.Text = "0.2"
        '
        'Label44
        '
        Me.Label44.AutoSize = True
        Me.Label44.Font = New System.Drawing.Font("Verdana", 9.75!, System.Drawing.FontStyle.Regular, System.Drawing.GraphicsUnit.Point, CType(0, Byte))
        Me.Label44.Location = New System.Drawing.Point(178, 236)
        Me.Label44.Name = "Label44"
        Me.Label44.Size = New System.Drawing.Size(54, 16)
        Me.Label44.TabIndex = 918
        Me.Label44.Text = " now ="
        '
        'chkNaKRegulation
        '
        Me.chkNaKRegulation.AutoSize = True
        Me.chkNaKRegulation.Checked = True
        Me.chkNaKRegulation.CheckState = System.Windows.Forms.CheckState.Checked
        Me.chkNaKRegulation.Font = New System.Drawing.Font("Verdana", 9.0!, System.Drawing.FontStyle.Regular, System.Drawing.GraphicsUnit.Point, CType(0, Byte))
        Me.chkNaKRegulation.Location = New System.Drawing.Point(7, 32)
        Me.chkNaKRegulation.Name = "chkNaKRegulation"
        Me.chkNaKRegulation.Size = New System.Drawing.Size(119, 18)
        Me.chkNaKRegulation.TabIndex = 508
        Me.chkNaKRegulation.Text = "NaK regulation"
        Me.chkNaKRegulation.UseVisualStyleBackColor = True
        '
        'chkWriteFile
        '
        Me.chkWriteFile.AutoSize = True
        Me.chkWriteFile.Font = New System.Drawing.Font("Verdana", 9.0!, System.Drawing.FontStyle.Regular, System.Drawing.GraphicsUnit.Point, CType(0, Byte))
        Me.chkWriteFile.Location = New System.Drawing.Point(109, 13)
        Me.chkWriteFile.Name = "chkWriteFile"
        Me.chkWriteFile.Size = New System.Drawing.Size(79, 18)
        Me.chkWriteFile.TabIndex = 506
        Me.chkWriteFile.Text = "writeFile"
        Me.chkWriteFile.UseVisualStyleBackColor = True
        '
        'chkDefault
        '
        Me.chkDefault.AutoSize = True
        Me.chkDefault.Checked = True
        Me.chkDefault.CheckState = System.Windows.Forms.CheckState.Checked
        Me.chkDefault.Font = New System.Drawing.Font("Verdana", 9.0!, System.Drawing.FontStyle.Regular, System.Drawing.GraphicsUnit.Point, CType(0, Byte))
        Me.chkDefault.Location = New System.Drawing.Point(7, 13)
        Me.chkDefault.Name = "chkDefault"
        Me.chkDefault.Size = New System.Drawing.Size(102, 18)
        Me.chkDefault.TabIndex = 505
        Me.chkDefault.Text = "DefaultStart"
        Me.chkDefault.UseVisualStyleBackColor = True
        '
        'Label17
        '
        Me.Label17.AutoSize = True
        Me.Label17.Font = New System.Drawing.Font("Verdana", 9.75!, System.Drawing.FontStyle.Regular, System.Drawing.GraphicsUnit.Point, CType(0, Byte))
        Me.Label17.Location = New System.Drawing.Point(131, 32)
        Me.Label17.Name = "Label17"
        Me.Label17.Size = New System.Drawing.Size(102, 16)
        Me.Label17.TabIndex = 2
        Me.Label17.Text = "Stimulus (pA) "
        '
        'txtNP
        '
        Me.txtNP.Font = New System.Drawing.Font("Verdana", 9.75!, System.Drawing.FontStyle.Regular, System.Drawing.GraphicsUnit.Point, CType(0, Byte))
        Me.txtNP.ForeColor = System.Drawing.Color.Black
        Me.txtNP.Location = New System.Drawing.Point(234, 233)
        Me.txtNP.Name = "txtNP"
        Me.txtNP.Size = New System.Drawing.Size(30, 23)
        Me.txtNP.TabIndex = 740
        Me.txtNP.Text = "1"
        '
        'txtStimulus
        '
        Me.txtStimulus.Font = New System.Drawing.Font("Verdana", 8.25!, System.Drawing.FontStyle.Regular, System.Drawing.GraphicsUnit.Point, CType(0, Byte))
        Me.txtStimulus.Location = New System.Drawing.Point(236, 33)
        Me.txtStimulus.Name = "txtStimulus"
        Me.txtStimulus.Size = New System.Drawing.Size(37, 21)
        Me.txtStimulus.TabIndex = 1
        Me.txtStimulus.Text = "0"
        '
        'txtEndms
        '
        Me.txtEndms.AccessibleRole = System.Windows.Forms.AccessibleRole.None
        Me.txtEndms.Font = New System.Drawing.Font("Verdana", 9.0!, System.Drawing.FontStyle.Regular, System.Drawing.GraphicsUnit.Point, CType(0, Byte))
        Me.txtEndms.Location = New System.Drawing.Point(211, 163)
        Me.txtEndms.Name = "txtEndms"
        Me.txtEndms.Size = New System.Drawing.Size(57, 22)
        Me.txtEndms.TabIndex = 800
        Me.txtEndms.Text = "6000"
        '
        'txt4th
        '
        Me.txt4th.Font = New System.Drawing.Font("Verdana", 9.0!, System.Drawing.FontStyle.Regular, System.Drawing.GraphicsUnit.Point, CType(0, Byte))
        Me.txt4th.Location = New System.Drawing.Point(89, 234)
        Me.txt4th.Name = "txt4th"
        Me.txt4th.Size = New System.Drawing.Size(28, 22)
        Me.txt4th.TabIndex = 847
        Me.txt4th.Text = "4"
        '
        'txt3rd
        '
        Me.txt3rd.Font = New System.Drawing.Font("Verdana", 9.0!, System.Drawing.FontStyle.Regular, System.Drawing.GraphicsUnit.Point, CType(0, Byte))
        Me.txt3rd.Location = New System.Drawing.Point(60, 234)
        Me.txt3rd.Name = "txt3rd"
        Me.txt3rd.Size = New System.Drawing.Size(28, 22)
        Me.txt3rd.TabIndex = 737
        Me.txt3rd.Text = "3"
        '
        'txtStartms
        '
        Me.txtStartms.AccessibleRole = System.Windows.Forms.AccessibleRole.None
        Me.txtStartms.Font = New System.Drawing.Font("Verdana", 9.0!, System.Drawing.FontStyle.Regular, System.Drawing.GraphicsUnit.Point, CType(0, Byte))
        Me.txtStartms.Location = New System.Drawing.Point(75, 162)
        Me.txtStartms.Name = "txtStartms"
        Me.txtStartms.Size = New System.Drawing.Size(57, 22)
        Me.txtStartms.TabIndex = 797
        Me.txtStartms.Text = "500"
        '
        'Label52
        '
        Me.Label52.AccessibleRole = System.Windows.Forms.AccessibleRole.None
        Me.Label52.AutoSize = True
        Me.Label52.Font = New System.Drawing.Font("Verdana", 11.25!, System.Drawing.FontStyle.Regular, System.Drawing.GraphicsUnit.Point, CType(0, Byte))
        Me.Label52.Location = New System.Drawing.Point(5, 162)
        Me.Label52.Name = "Label52"
        Me.Label52.Size = New System.Drawing.Size(68, 18)
        Me.Label52.TabIndex = 798
        Me.Label52.Text = "Startms"
        '
        'txt2nd
        '
        Me.txt2nd.Font = New System.Drawing.Font("Verdana", 9.0!, System.Drawing.FontStyle.Regular, System.Drawing.GraphicsUnit.Point, CType(0, Byte))
        Me.txt2nd.Location = New System.Drawing.Point(31, 234)
        Me.txt2nd.Name = "txt2nd"
        Me.txt2nd.Size = New System.Drawing.Size(28, 22)
        Me.txt2nd.TabIndex = 736
        Me.txt2nd.Text = "2"
        '
        'Label16
        '
        Me.Label16.AccessibleRole = System.Windows.Forms.AccessibleRole.None
        Me.Label16.AutoSize = True
        Me.Label16.Font = New System.Drawing.Font("Verdana", 11.25!, System.Drawing.FontStyle.Regular, System.Drawing.GraphicsUnit.Point, CType(0, Byte))
        Me.Label16.Location = New System.Drawing.Point(142, 162)
        Me.Label16.Name = "Label16"
        Me.Label16.Size = New System.Drawing.Size(58, 18)
        Me.Label16.TabIndex = 801
        Me.Label16.Text = "Endms"
        '
        'txtEval
        '
        Me.txtEval.Font = New System.Drawing.Font("Verdana", 9.75!, System.Drawing.FontStyle.Regular, System.Drawing.GraphicsUnit.Point, CType(0, Byte))
        Me.txtEval.Location = New System.Drawing.Point(220, 210)
        Me.txtEval.Name = "txtEval"
        Me.txtEval.Size = New System.Drawing.Size(58, 23)
        Me.txtEval.TabIndex = 861
        '
        'txt1st
        '
        Me.txt1st.Font = New System.Drawing.Font("Verdana", 9.0!, System.Drawing.FontStyle.Regular, System.Drawing.GraphicsUnit.Point, CType(0, Byte))
        Me.txt1st.Location = New System.Drawing.Point(3, 234)
        Me.txt1st.Name = "txt1st"
        Me.txt1st.Size = New System.Drawing.Size(28, 22)
        Me.txt1st.TabIndex = 735
        Me.txt1st.Text = "1"
        '
        'txtRgR
        '
        Me.txtRgR.Font = New System.Drawing.Font("Verdana", 9.75!, System.Drawing.FontStyle.Regular, System.Drawing.GraphicsUnit.Point, CType(0, Byte))
        Me.txtRgR.Location = New System.Drawing.Point(192, 185)
        Me.txtRgR.Name = "txtRgR"
        Me.txtRgR.Size = New System.Drawing.Size(43, 23)
        Me.txtRgR.TabIndex = 930
        Me.txtRgR.Text = "1600"
        '
        'txtRgL
        '
        Me.txtRgL.Font = New System.Drawing.Font("Verdana", 9.75!, System.Drawing.FontStyle.Regular, System.Drawing.GraphicsUnit.Point, CType(0, Byte))
        Me.txtRgL.Location = New System.Drawing.Point(130, 185)
        Me.txtRgL.Name = "txtRgL"
        Me.txtRgL.Size = New System.Drawing.Size(43, 23)
        Me.txtRgL.TabIndex = 854
        Me.txtRgL.Text = "10"
        '
        'txtEvalMax
        '
        Me.txtEvalMax.Font = New System.Drawing.Font("Verdana", 9.75!, System.Drawing.FontStyle.Regular, System.Drawing.GraphicsUnit.Point, CType(0, Byte))
        Me.txtEvalMax.Location = New System.Drawing.Point(229, 136)
        Me.txtEvalMax.Name = "txtEvalMax"
        Me.txtEvalMax.Size = New System.Drawing.Size(44, 23)
        Me.txtEvalMax.TabIndex = 855
        Me.txtEvalMax.Text = "2000"
        '
        'Label50
        '
        Me.Label50.AutoSize = True
        Me.Label50.Font = New System.Drawing.Font("Verdana", 9.75!, System.Drawing.FontStyle.Regular, System.Drawing.GraphicsUnit.Point, CType(0, Byte))
        Me.Label50.Location = New System.Drawing.Point(141, 141)
        Me.Label50.Name = "Label50"
        Me.Label50.Size = New System.Drawing.Size(82, 16)
        Me.Label50.TabIndex = 856
        Me.Label50.Text = "Eval-Max ="
        '
        'Label19
        '
        Me.Label19.AutoSize = True
        Me.Label19.Font = New System.Drawing.Font("Verdana", 9.75!, System.Drawing.FontStyle.Regular, System.Drawing.GraphicsUnit.Point, CType(0, Byte))
        Me.Label19.Location = New System.Drawing.Point(173, 188)
        Me.Label19.Name = "Label19"
        Me.Label19.Size = New System.Drawing.Size(19, 16)
        Me.Label19.TabIndex = 857
        Me.Label19.Text = "~"
        '
        'txtMaxRepeat
        '
        Me.txtMaxRepeat.Font = New System.Drawing.Font("Verdana", 9.75!, System.Drawing.FontStyle.Regular, System.Drawing.GraphicsUnit.Point, CType(0, Byte))
        Me.txtMaxRepeat.Location = New System.Drawing.Point(103, 136)
        Me.txtMaxRepeat.Name = "txtMaxRepeat"
        Me.txtMaxRepeat.Size = New System.Drawing.Size(34, 23)
        Me.txtMaxRepeat.TabIndex = 859
        Me.txtMaxRepeat.Text = "30"
        '
        'Label11
        '
        Me.Label11.AutoSize = True
        Me.Label11.Font = New System.Drawing.Font("Verdana", 9.75!, System.Drawing.FontStyle.Regular, System.Drawing.GraphicsUnit.Point, CType(0, Byte))
        Me.Label11.Location = New System.Drawing.Point(2, 141)
        Me.Label11.Name = "Label11"
        Me.Label11.Size = New System.Drawing.Size(94, 16)
        Me.Label11.TabIndex = 860
        Me.Label11.Text = "maxRepeat ="
        '
        'Label7
        '
        Me.Label7.AutoSize = True
        Me.Label7.Font = New System.Drawing.Font("Verdana", 9.75!, System.Drawing.FontStyle.Regular, System.Drawing.GraphicsUnit.Point, CType(0, Byte))
        Me.Label7.Location = New System.Drawing.Point(180, 211)
        Me.Label7.Name = "Label7"
        Me.Label7.Size = New System.Drawing.Size(35, 16)
        Me.Label7.TabIndex = 675
        Me.Label7.Text = "Eval"
        '
        'Label26
        '
        Me.Label26.AutoSize = True
        Me.Label26.Font = New System.Drawing.Font("Verdana", 9.75!, System.Drawing.FontStyle.Regular, System.Drawing.GraphicsUnit.Point, CType(0, Byte))
        Me.Label26.Location = New System.Drawing.Point(3, 188)
        Me.Label26.Name = "Label26"
        Me.Label26.Size = New System.Drawing.Size(123, 16)
        Me.Label26.TabIndex = 853
        Me.Label26.Text = "SQS time window"
        '
        'controlBox4
        '
        Me.controlBox4.BackColor = System.Drawing.Color.FromArgb(CType(CType(255, Byte), Integer), CType(CType(255, Byte), Integer), CType(CType(192, Byte), Integer))
        Me.controlBox4.Controls.Add(Me.Label80)
        Me.controlBox4.Controls.Add(Me.Label79)
        Me.controlBox4.Controls.Add(Me.txt2R)
        Me.controlBox4.Controls.Add(Me.txtR3)
        Me.controlBox4.Controls.Add(Me.txtR1)
        Me.controlBox4.Controls.Add(Me.txtF4)
        Me.controlBox4.Controls.Add(Me.txtF3)
        Me.controlBox4.Controls.Add(Me.txtF2)
        Me.controlBox4.Controls.Add(Me.txtF1)
        Me.controlBox4.Controls.Add(Me.Label2)
        Me.controlBox4.Controls.Add(Me.Label71)
        Me.controlBox4.Controls.Add(Me.Label70)
        Me.controlBox4.Controls.Add(Me.Label56)
        Me.controlBox4.Controls.Add(Me.txtNai)
        Me.controlBox4.Controls.Add(Me.chkIbNSC)
        Me.controlBox4.Controls.Add(Me.txtINCXmag)
        Me.controlBox4.Controls.Add(Me.txtINaKmag)
        Me.controlBox4.Controls.Add(Me.chkINCX)
        Me.controlBox4.Controls.Add(Me.chkINaK)
        Me.controlBox4.Controls.Add(Me.chkIst)
        Me.controlBox4.Controls.Add(Me.chkIKto)
        Me.controlBox4.Controls.Add(Me.chkIKs)
        Me.controlBox4.Controls.Add(Me.chkIha)
        Me.controlBox4.Controls.Add(Me.chkIKur)
        Me.controlBox4.Controls.Add(Me.chkINa)
        Me.controlBox4.Controls.Add(Me.chkICaL)
        Me.controlBox4.Controls.Add(Me.chkIK1)
        Me.controlBox4.Controls.Add(Me.chkIKr)
        Me.controlBox4.Controls.Add(Me.NFusCaLmag)
        Me.controlBox4.Controls.Add(Me.rFusCaL)
        Me.controlBox4.Controls.Add(Me.Label60)
        Me.controlBox4.Controls.Add(Me.txtstdtotCai)
        Me.controlBox4.Controls.Add(Me.txtstdNai)
        Me.controlBox4.Controls.Add(Me.Label59)
        Me.controlBox4.Controls.Add(Me.txtFusCaLmag)
        Me.controlBox4.Controls.Add(Me.txtVshiftNa)
        Me.controlBox4.Controls.Add(Me.Label23)
        Me.controlBox4.Controls.Add(Me.Label3)
        Me.controlBox4.Controls.Add(Me.Label5)
        Me.controlBox4.Controls.Add(Me.txtVshiftCaL)
        Me.controlBox4.Controls.Add(Me.Label43)
        Me.controlBox4.Controls.Add(Me.Label35)
        Me.controlBox4.Controls.Add(Me.txtKLmag)
        Me.controlBox4.Controls.Add(Me.Label25)
        Me.controlBox4.Controls.Add(Me.Label30)
        Me.controlBox4.Controls.Add(Me.IKr)
        Me.controlBox4.Controls.Add(Me.txtICaLmag)
        Me.controlBox4.Controls.Add(Me.txtIK1mag)
        Me.controlBox4.Controls.Add(Me.Label15)
        Me.controlBox4.Controls.Add(Me.txtIKrmag)
        Me.controlBox4.Controls.Add(Me.Label53)
        Me.controlBox4.Controls.Add(Me.txtNaLratio)
        Me.controlBox4.Controls.Add(Me.Label51)
        Me.controlBox4.Controls.Add(Me.Label6)
        Me.controlBox4.Controls.Add(Me.txtBaseVshift)
        Me.controlBox4.Controls.Add(Me.Label48)
        Me.controlBox4.Controls.Add(Me.Label9)
        Me.controlBox4.Controls.Add(Me.txtorgExtraVshift)
        Me.controlBox4.Controls.Add(Me.txtICaTmag)
        Me.controlBox4.Controls.Add(Me.Label34)
        Me.controlBox4.Controls.Add(Me.txtIKurmag)
        Me.controlBox4.Controls.Add(Me.txtOS)
        Me.controlBox4.Controls.Add(Me.Label10)
        Me.controlBox4.Controls.Add(Me.Label13)
        Me.controlBox4.Controls.Add(Me.Label8)
        Me.controlBox4.Controls.Add(Me.txtIKsmag)
        Me.controlBox4.Controls.Add(Me.txtMDP)
        Me.controlBox4.Controls.Add(Me.Label27)
        Me.controlBox4.Controls.Add(Me.Label14)
        Me.controlBox4.Controls.Add(Me.rNa)
        Me.controlBox4.Controls.Add(Me.Label1)
        Me.controlBox4.Controls.Add(Me.Label18)
        Me.controlBox4.Controls.Add(Me.txtCycleL)
        Me.controlBox4.Controls.Add(Me.Label28)
        Me.controlBox4.Controls.Add(Me.txtIKtomag)
        Me.controlBox4.Controls.Add(Me.txtAPD)
        Me.controlBox4.Controls.Add(Me.rKr)
        Me.controlBox4.Controls.Add(Me.txtINamag)
        Me.controlBox4.Controls.Add(Me.txtTotalCai)
        Me.controlBox4.Controls.Add(Me.rK1)
        Me.controlBox4.Controls.Add(Me.txtcrfNCX)
        Me.controlBox4.Controls.Add(Me.Label22)
        Me.controlBox4.Controls.Add(Me.rVshiftCaL)
        Me.controlBox4.Controls.Add(Me.txtcrfNaK)
        Me.controlBox4.Controls.Add(Me.rCaL)
        Me.controlBox4.Controls.Add(Me.txtIbNSCmag)
        Me.controlBox4.Controls.Add(Me.Label24)
        Me.controlBox4.Controls.Add(Me.txtKi)
        Me.controlBox4.Controls.Add(Me.rCaT)
        Me.controlBox4.Controls.Add(Me.Label29)
        Me.controlBox4.Controls.Add(Me.rextraVshift)
        Me.controlBox4.Controls.Add(Me.rst)
        Me.controlBox4.Controls.Add(Me.txtIhamag)
        Me.controlBox4.Controls.Add(Me.rbNSC)
        Me.controlBox4.Controls.Add(Me.NVshiftCaL)
        Me.controlBox4.Controls.Add(Me.Label47)
        Me.controlBox4.Controls.Add(Me.rKL)
        Me.controlBox4.Controls.Add(Me.NextraVshift)
        Me.controlBox4.Controls.Add(Me.rKto)
        Me.controlBox4.Controls.Add(Me.NVshiftNa)
        Me.controlBox4.Controls.Add(Me.rVshiftNa)
        Me.controlBox4.Controls.Add(Me.txtIstmag)
        Me.controlBox4.Controls.Add(Me.NIhamag)
        Me.controlBox4.Controls.Add(Me.rKs)
        Me.controlBox4.Controls.Add(Me.NICaTmag)
        Me.controlBox4.Controls.Add(Me.rha)
        Me.controlBox4.Controls.Add(Me.NIKurmag)
        Me.controlBox4.Controls.Add(Me.rKur)
        Me.controlBox4.Controls.Add(Me.Label20)
        Me.controlBox4.Controls.Add(Me.NIKsmag)
        Me.controlBox4.Controls.Add(Me.NIKtomag)
        Me.controlBox4.Controls.Add(Me.NIstmag)
        Me.controlBox4.Controls.Add(Me.Label21)
        Me.controlBox4.Controls.Add(Me.NKLmag)
        Me.controlBox4.Controls.Add(Me.NINamag)
        Me.controlBox4.Controls.Add(Me.NIbNSCmag)
        Me.controlBox4.Controls.Add(Me.NIKrmag)
        Me.controlBox4.Controls.Add(Me.NIK1mag)
        Me.controlBox4.Controls.Add(Me.NICaLmag)
        Me.controlBox4.Location = New System.Drawing.Point(5, 394)
        Me.controlBox4.Name = "controlBox4"
        Me.controlBox4.Size = New System.Drawing.Size(285, 731)
        Me.controlBox4.TabIndex = 934
        Me.controlBox4.TabStop = False
        Me.controlBox4.Text = "Measurements"
        '
        'Label80
        '
        Me.Label80.AutoSize = True
        Me.Label80.Font = New System.Drawing.Font("Verdana", 9.75!, System.Drawing.FontStyle.Regular, System.Drawing.GraphicsUnit.Point, CType(0, Byte))
        Me.Label80.Location = New System.Drawing.Point(3, 705)
        Me.Label80.Name = "Label80"
        Me.Label80.Size = New System.Drawing.Size(16, 16)
        Me.Label80.TabIndex = 956
        Me.Label80.Text = "R"
        '
        'Label79
        '
        Me.Label79.AutoSize = True
        Me.Label79.Font = New System.Drawing.Font("Verdana", 9.75!, System.Drawing.FontStyle.Regular, System.Drawing.GraphicsUnit.Point, CType(0, Byte))
        Me.Label79.Location = New System.Drawing.Point(4, 686)
        Me.Label79.Name = "Label79"
        Me.Label79.Size = New System.Drawing.Size(16, 16)
        Me.Label79.TabIndex = 955
        Me.Label79.Text = "F"
        '
        'txt2R
        '
        Me.txt2R.Font = New System.Drawing.Font("Verdana", 9.75!, System.Drawing.FontStyle.Regular, System.Drawing.GraphicsUnit.Point, CType(0, Byte))
        Me.txt2R.Location = New System.Drawing.Point(87, 702)
        Me.txt2R.Name = "txt2R"
        Me.txt2R.Size = New System.Drawing.Size(56, 23)
        Me.txt2R.TabIndex = 954
        Me.txt2R.Text = "0"
        '
        'txtR3
        '
        Me.txtR3.Font = New System.Drawing.Font("Verdana", 9.75!, System.Drawing.FontStyle.Regular, System.Drawing.GraphicsUnit.Point, CType(0, Byte))
        Me.txtR3.Location = New System.Drawing.Point(149, 702)
        Me.txtR3.Name = "txtR3"
        Me.txtR3.Size = New System.Drawing.Size(56, 23)
        Me.txtR3.TabIndex = 953
        Me.txtR3.Text = "0"
        '
        'txtR1
        '
        Me.txtR1.Font = New System.Drawing.Font("Verdana", 9.75!, System.Drawing.FontStyle.Regular, System.Drawing.GraphicsUnit.Point, CType(0, Byte))
        Me.txtR1.Location = New System.Drawing.Point(25, 702)
        Me.txtR1.Name = "txtR1"
        Me.txtR1.Size = New System.Drawing.Size(56, 23)
        Me.txtR1.TabIndex = 952
        Me.txtR1.Text = "0"
        '
        'txtF4
        '
        Me.txtF4.Font = New System.Drawing.Font("Verdana", 9.75!, System.Drawing.FontStyle.Regular, System.Drawing.GraphicsUnit.Point, CType(0, Byte))
        Me.txtF4.Location = New System.Drawing.Point(209, 679)
        Me.txtF4.Name = "txtF4"
        Me.txtF4.Size = New System.Drawing.Size(56, 23)
        Me.txtF4.TabIndex = 951
        Me.txtF4.Text = "0"
        '
        'txtF3
        '
        Me.txtF3.Font = New System.Drawing.Font("Verdana", 9.75!, System.Drawing.FontStyle.Regular, System.Drawing.GraphicsUnit.Point, CType(0, Byte))
        Me.txtF3.Location = New System.Drawing.Point(150, 679)
        Me.txtF3.Name = "txtF3"
        Me.txtF3.Size = New System.Drawing.Size(56, 23)
        Me.txtF3.TabIndex = 950
        Me.txtF3.Text = "0"
        '
        'txtF2
        '
        Me.txtF2.Font = New System.Drawing.Font("Verdana", 9.75!, System.Drawing.FontStyle.Regular, System.Drawing.GraphicsUnit.Point, CType(0, Byte))
        Me.txtF2.Location = New System.Drawing.Point(88, 679)
        Me.txtF2.Name = "txtF2"
        Me.txtF2.Size = New System.Drawing.Size(56, 23)
        Me.txtF2.TabIndex = 949
        Me.txtF2.Text = "0"
        '
        'txtF1
        '
        Me.txtF1.Font = New System.Drawing.Font("Verdana", 9.75!, System.Drawing.FontStyle.Regular, System.Drawing.GraphicsUnit.Point, CType(0, Byte))
        Me.txtF1.Location = New System.Drawing.Point(26, 679)
        Me.txtF1.Name = "txtF1"
        Me.txtF1.Size = New System.Drawing.Size(56, 23)
        Me.txtF1.TabIndex = 948
        Me.txtF1.Text = "0"
        '
        'Label2
        '
        Me.Label2.AutoSize = True
        Me.Label2.Font = New System.Drawing.Font("Verdana", 9.75!, System.Drawing.FontStyle.Regular, System.Drawing.GraphicsUnit.Point, CType(0, Byte))
        Me.Label2.Location = New System.Drawing.Point(1, 89)
        Me.Label2.Name = "Label2"
        Me.Label2.Size = New System.Drawing.Size(50, 16)
        Me.Label2.TabIndex = 918
        Me.Label2.Text = "plot Ix"
        '
        'Label71
        '
        Me.Label71.AutoSize = True
        Me.Label71.Font = New System.Drawing.Font("Verdana", 9.75!, System.Drawing.FontStyle.Regular, System.Drawing.GraphicsUnit.Point, CType(0, Byte))
        Me.Label71.Location = New System.Drawing.Point(190, 88)
        Me.Label71.Name = "Label71"
        Me.Label71.Size = New System.Drawing.Size(91, 16)
        Me.Label71.TabIndex = 917
        Me.Label71.Text = " Ixmag x issf"
        '
        'Label70
        '
        Me.Label70.AutoSize = True
        Me.Label70.Font = New System.Drawing.Font("Verdana", 9.75!, System.Drawing.FontStyle.Regular, System.Drawing.GraphicsUnit.Point, CType(0, Byte))
        Me.Label70.Location = New System.Drawing.Point(140, 89)
        Me.Label70.Name = "Label70"
        Me.Label70.Size = New System.Drawing.Size(32, 16)
        Me.Label70.TabIndex = 916
        Me.Label70.Text = "Ixsf"
        '
        'Label56
        '
        Me.Label56.AutoSize = True
        Me.Label56.Font = New System.Drawing.Font("Verdana", 9.75!, System.Drawing.FontStyle.Regular, System.Drawing.GraphicsUnit.Point, CType(0, Byte))
        Me.Label56.Location = New System.Drawing.Point(68, 88)
        Me.Label56.Name = "Label56"
        Me.Label56.Size = New System.Drawing.Size(47, 16)
        Me.Label56.TabIndex = 915
        Me.Label56.Text = "Ixmag"
        '
        'txtNai
        '
        Me.txtNai.Font = New System.Drawing.Font("Verdana", 12.0!, System.Drawing.FontStyle.Bold, System.Drawing.GraphicsUnit.Point, CType(0, Byte))
        Me.txtNai.Location = New System.Drawing.Point(211, 549)
        Me.txtNai.Name = "txtNai"
        Me.txtNai.Size = New System.Drawing.Size(71, 27)
        Me.txtNai.TabIndex = 914
        Me.txtNai.Text = "1"
        '
        'chkIbNSC
        '
        Me.chkIbNSC.AutoSize = True
        Me.chkIbNSC.Checked = True
        Me.chkIbNSC.CheckState = System.Windows.Forms.CheckState.Checked
        Me.chkIbNSC.Location = New System.Drawing.Point(6, 260)
        Me.chkIbNSC.Name = "chkIbNSC"
        Me.chkIbNSC.Size = New System.Drawing.Size(15, 14)
        Me.chkIbNSC.TabIndex = 913
        Me.chkIbNSC.UseVisualStyleBackColor = True
        '
        'txtINCXmag
        '
        Me.txtINCXmag.Font = New System.Drawing.Font("Verdana", 9.0!, System.Drawing.FontStyle.Regular, System.Drawing.GraphicsUnit.Point, CType(0, Byte))
        Me.txtINCXmag.Location = New System.Drawing.Point(61, 500)
        Me.txtINCXmag.Name = "txtINCXmag"
        Me.txtINCXmag.Size = New System.Drawing.Size(67, 22)
        Me.txtINCXmag.TabIndex = 734
        Me.txtINCXmag.Text = "2"
        '
        'txtINaKmag
        '
        Me.txtINaKmag.Font = New System.Drawing.Font("Verdana", 9.0!, System.Drawing.FontStyle.Regular, System.Drawing.GraphicsUnit.Point, CType(0, Byte))
        Me.txtINaKmag.Location = New System.Drawing.Point(62, 476)
        Me.txtINaKmag.Name = "txtINaKmag"
        Me.txtINaKmag.Size = New System.Drawing.Size(67, 22)
        Me.txtINaKmag.TabIndex = 730
        Me.txtINaKmag.Tag = "28.5634"
        Me.txtINaKmag.Text = "2"
        '
        'chkINCX
        '
        Me.chkINCX.AutoSize = True
        Me.chkINCX.Checked = True
        Me.chkINCX.CheckState = System.Windows.Forms.CheckState.Checked
        Me.chkINCX.Location = New System.Drawing.Point(6, 505)
        Me.chkINCX.Name = "chkINCX"
        Me.chkINCX.Size = New System.Drawing.Size(15, 14)
        Me.chkINCX.TabIndex = 729
        Me.chkINCX.UseVisualStyleBackColor = True
        '
        'chkINaK
        '
        Me.chkINaK.AutoSize = True
        Me.chkINaK.Checked = True
        Me.chkINaK.CheckState = System.Windows.Forms.CheckState.Checked
        Me.chkINaK.Location = New System.Drawing.Point(7, 479)
        Me.chkINaK.Name = "chkINaK"
        Me.chkINaK.Size = New System.Drawing.Size(15, 14)
        Me.chkINaK.TabIndex = 728
        Me.chkINaK.UseVisualStyleBackColor = True
        '
        'chkIst
        '
        Me.chkIst.AutoSize = True
        Me.chkIst.Checked = True
        Me.chkIst.CheckState = System.Windows.Forms.CheckState.Checked
        Me.chkIst.Location = New System.Drawing.Point(6, 457)
        Me.chkIst.Name = "chkIst"
        Me.chkIst.Size = New System.Drawing.Size(15, 14)
        Me.chkIst.TabIndex = 727
        Me.chkIst.UseVisualStyleBackColor = True
        '
        'chkIKto
        '
        Me.chkIKto.AutoSize = True
        Me.chkIKto.Checked = True
        Me.chkIKto.CheckState = System.Windows.Forms.CheckState.Checked
        Me.chkIKto.Location = New System.Drawing.Point(7, 335)
        Me.chkIKto.Name = "chkIKto"
        Me.chkIKto.Size = New System.Drawing.Size(15, 14)
        Me.chkIKto.TabIndex = 726
        Me.chkIKto.UseVisualStyleBackColor = True
        '
        'chkIKs
        '
        Me.chkIKs.AutoSize = True
        Me.chkIKs.Checked = True
        Me.chkIKs.CheckState = System.Windows.Forms.CheckState.Checked
        Me.chkIKs.Location = New System.Drawing.Point(8, 308)
        Me.chkIKs.Name = "chkIKs"
        Me.chkIKs.Size = New System.Drawing.Size(15, 14)
        Me.chkIKs.TabIndex = 678
        Me.chkIKs.UseVisualStyleBackColor = True
        '
        'chkIha
        '
        Me.chkIha.AutoSize = True
        Me.chkIha.Checked = True
        Me.chkIha.CheckState = System.Windows.Forms.CheckState.Checked
        Me.chkIha.Location = New System.Drawing.Point(8, 284)
        Me.chkIha.Name = "chkIha"
        Me.chkIha.Size = New System.Drawing.Size(15, 14)
        Me.chkIha.TabIndex = 678
        Me.chkIha.UseVisualStyleBackColor = True
        '
        'chkIKur
        '
        Me.chkIKur.AutoSize = True
        Me.chkIKur.Checked = True
        Me.chkIKur.CheckState = System.Windows.Forms.CheckState.Checked
        Me.chkIKur.Location = New System.Drawing.Point(10, 359)
        Me.chkIKur.Name = "chkIKur"
        Me.chkIKur.Size = New System.Drawing.Size(15, 14)
        Me.chkIKur.TabIndex = 725
        Me.chkIKur.UseVisualStyleBackColor = True
        '
        'chkINa
        '
        Me.chkINa.AutoSize = True
        Me.chkINa.Checked = True
        Me.chkINa.CheckState = System.Windows.Forms.CheckState.Checked
        Me.chkINa.Location = New System.Drawing.Point(8, 383)
        Me.chkINa.Name = "chkINa"
        Me.chkINa.Size = New System.Drawing.Size(15, 14)
        Me.chkINa.TabIndex = 724
        Me.chkINa.UseVisualStyleBackColor = True
        '
        'chkICaL
        '
        Me.chkICaL.AutoSize = True
        Me.chkICaL.Checked = True
        Me.chkICaL.CheckState = System.Windows.Forms.CheckState.Checked
        Me.chkICaL.Location = New System.Drawing.Point(10, 159)
        Me.chkICaL.Name = "chkICaL"
        Me.chkICaL.Size = New System.Drawing.Size(15, 14)
        Me.chkICaL.TabIndex = 677
        Me.chkICaL.UseVisualStyleBackColor = True
        '
        'chkIK1
        '
        Me.chkIK1.AutoSize = True
        Me.chkIK1.Checked = True
        Me.chkIK1.CheckState = System.Windows.Forms.CheckState.Checked
        Me.chkIK1.Location = New System.Drawing.Point(11, 136)
        Me.chkIK1.Name = "chkIK1"
        Me.chkIK1.Size = New System.Drawing.Size(15, 14)
        Me.chkIK1.TabIndex = 676
        Me.chkIK1.UseVisualStyleBackColor = True
        '
        'chkIKr
        '
        Me.chkIKr.AutoSize = True
        Me.chkIKr.Checked = True
        Me.chkIKr.CheckState = System.Windows.Forms.CheckState.Checked
        Me.chkIKr.Location = New System.Drawing.Point(11, 114)
        Me.chkIKr.Name = "chkIKr"
        Me.chkIKr.Size = New System.Drawing.Size(15, 14)
        Me.chkIKr.TabIndex = 676
        Me.chkIKr.UseVisualStyleBackColor = True
        '
        'NFusCaLmag
        '
        Me.NFusCaLmag.Font = New System.Drawing.Font("Verdana", 9.75!, System.Drawing.FontStyle.Regular, System.Drawing.GraphicsUnit.Point, CType(0, Byte))
        Me.NFusCaLmag.ForeColor = System.Drawing.Color.Black
        Me.NFusCaLmag.Location = New System.Drawing.Point(206, 232)
        Me.NFusCaLmag.Name = "NFusCaLmag"
        Me.NFusCaLmag.Size = New System.Drawing.Size(63, 23)
        Me.NFusCaLmag.TabIndex = 723
        Me.NFusCaLmag.Text = "0"
        '
        'rFusCaL
        '
        Me.rFusCaL.Font = New System.Drawing.Font("Verdana", 9.75!, System.Drawing.FontStyle.Regular, System.Drawing.GraphicsUnit.Point, CType(0, Byte))
        Me.rFusCaL.ForeColor = System.Drawing.Color.Black
        Me.rFusCaL.Location = New System.Drawing.Point(138, 232)
        Me.rFusCaL.Name = "rFusCaL"
        Me.rFusCaL.Size = New System.Drawing.Size(63, 23)
        Me.rFusCaL.TabIndex = 722
        Me.rFusCaL.Text = "1"
        '
        'Label60
        '
        Me.Label60.AutoSize = True
        Me.Label60.Font = New System.Drawing.Font("Verdana", 9.75!, System.Drawing.FontStyle.Regular, System.Drawing.GraphicsUnit.Point, CType(0, Byte))
        Me.Label60.Location = New System.Drawing.Point(17, 605)
        Me.Label60.Name = "Label60"
        Me.Label60.Size = New System.Drawing.Size(54, 16)
        Me.Label60.TabIndex = 721
        Me.Label60.Text = "stdCai "
        '
        'txtstdtotCai
        '
        Me.txtstdtotCai.Font = New System.Drawing.Font("Verdana", 9.75!, System.Drawing.FontStyle.Regular, System.Drawing.GraphicsUnit.Point, CType(0, Byte))
        Me.txtstdtotCai.Location = New System.Drawing.Point(80, 602)
        Me.txtstdtotCai.Name = "txtstdtotCai"
        Me.txtstdtotCai.Size = New System.Drawing.Size(58, 23)
        Me.txtstdtotCai.TabIndex = 720
        Me.txtstdtotCai.Text = "79"
        '
        'txtstdNai
        '
        Me.txtstdNai.Font = New System.Drawing.Font("Verdana", 9.0!, System.Drawing.FontStyle.Regular, System.Drawing.GraphicsUnit.Point, CType(0, Byte))
        Me.txtstdNai.Location = New System.Drawing.Point(76, 552)
        Me.txtstdNai.Name = "txtstdNai"
        Me.txtstdNai.Size = New System.Drawing.Size(60, 22)
        Me.txtstdNai.TabIndex = 719
        Me.txtstdNai.Text = "6.1"
        '
        'Label59
        '
        Me.Label59.AutoSize = True
        Me.Label59.Font = New System.Drawing.Font("Verdana", 9.0!, System.Drawing.FontStyle.Regular, System.Drawing.GraphicsUnit.Point, CType(0, Byte))
        Me.Label59.Location = New System.Drawing.Point(19, 556)
        Me.Label59.Name = "Label59"
        Me.Label59.Size = New System.Drawing.Size(47, 14)
        Me.Label59.TabIndex = 718
        Me.Label59.Text = "stdNai"
        '
        'txtFusCaLmag
        '
        Me.txtFusCaLmag.Font = New System.Drawing.Font("Verdana", 9.75!, System.Drawing.FontStyle.Regular, System.Drawing.GraphicsUnit.Point, CType(0, Byte))
        Me.txtFusCaLmag.Location = New System.Drawing.Point(67, 231)
        Me.txtFusCaLmag.Name = "txtFusCaLmag"
        Me.txtFusCaLmag.Size = New System.Drawing.Size(66, 23)
        Me.txtFusCaLmag.TabIndex = 716
        Me.txtFusCaLmag.Text = "1"
        '
        'txtVshiftNa
        '
        Me.txtVshiftNa.Font = New System.Drawing.Font("Verdana", 9.75!, System.Drawing.FontStyle.Regular, System.Drawing.GraphicsUnit.Point, CType(0, Byte))
        Me.txtVshiftNa.Location = New System.Drawing.Point(63, 405)
        Me.txtVshiftNa.Name = "txtVshiftNa"
        Me.txtVshiftNa.Size = New System.Drawing.Size(67, 23)
        Me.txtVshiftNa.TabIndex = 714
        Me.txtVshiftNa.Text = "6.4328"
        '
        'Label23
        '
        Me.Label23.AutoSize = True
        Me.Label23.Font = New System.Drawing.Font("Verdana", 9.75!, System.Drawing.FontStyle.Regular, System.Drawing.GraphicsUnit.Point, CType(0, Byte))
        Me.Label23.ForeColor = System.Drawing.Color.Red
        Me.Label23.Location = New System.Drawing.Point(20, 502)
        Me.Label23.Name = "Label23"
        Me.Label23.Size = New System.Drawing.Size(40, 16)
        Me.Label23.TabIndex = 713
        Me.Label23.Text = "INCX"
        '
        'Label3
        '
        Me.Label3.AutoSize = True
        Me.Label3.Font = New System.Drawing.Font("Verdana", 9.75!, System.Drawing.FontStyle.Regular, System.Drawing.GraphicsUnit.Point, CType(0, Byte))
        Me.Label3.Location = New System.Drawing.Point(9, 234)
        Me.Label3.Name = "Label3"
        Me.Label3.Size = New System.Drawing.Size(60, 16)
        Me.Label3.TabIndex = 715
        Me.Label3.Text = "FusCaL "
        '
        'Label5
        '
        Me.Label5.AutoSize = True
        Me.Label5.Font = New System.Drawing.Font("Verdana", 9.75!, System.Drawing.FontStyle.Regular, System.Drawing.GraphicsUnit.Point, CType(0, Byte))
        Me.Label5.Location = New System.Drawing.Point(2, 209)
        Me.Label5.Name = "Label5"
        Me.Label5.Size = New System.Drawing.Size(63, 16)
        Me.Label5.TabIndex = 708
        Me.Label5.Text = "VshiftCa"
        '
        'txtVshiftCaL
        '
        Me.txtVshiftCaL.Font = New System.Drawing.Font("Verdana", 9.75!, System.Drawing.FontStyle.Regular, System.Drawing.GraphicsUnit.Point, CType(0, Byte))
        Me.txtVshiftCaL.Location = New System.Drawing.Point(66, 206)
        Me.txtVshiftCaL.Name = "txtVshiftCaL"
        Me.txtVshiftCaL.Size = New System.Drawing.Size(63, 23)
        Me.txtVshiftCaL.TabIndex = 707
        Me.txtVshiftCaL.Text = "0.2031"
        '
        'Label43
        '
        Me.Label43.AutoSize = True
        Me.Label43.Font = New System.Drawing.Font("Verdana", 9.75!, System.Drawing.FontStyle.Regular, System.Drawing.GraphicsUnit.Point, CType(0, Byte))
        Me.Label43.ForeColor = System.Drawing.Color.FromArgb(CType(CType(255, Byte), Integer), CType(CType(128, Byte), Integer), CType(CType(255, Byte), Integer))
        Me.Label43.Location = New System.Drawing.Point(22, 478)
        Me.Label43.Name = "Label43"
        Me.Label43.Size = New System.Drawing.Size(38, 16)
        Me.Label43.TabIndex = 711
        Me.Label43.Text = "INaK"
        '
        'Label35
        '
        Me.Label35.AutoSize = True
        Me.Label35.Font = New System.Drawing.Font("Verdana", 9.75!, System.Drawing.FontStyle.Regular, System.Drawing.GraphicsUnit.Point, CType(0, Byte))
        Me.Label35.Location = New System.Drawing.Point(27, 184)
        Me.Label35.Name = "Label35"
        Me.Label35.Size = New System.Drawing.Size(23, 16)
        Me.Label35.TabIndex = 696
        Me.Label35.Text = "KL"
        '
        'txtKLmag
        '
        Me.txtKLmag.Font = New System.Drawing.Font("Verdana", 9.75!, System.Drawing.FontStyle.Regular, System.Drawing.GraphicsUnit.Point, CType(0, Byte))
        Me.txtKLmag.Location = New System.Drawing.Point(66, 181)
        Me.txtKLmag.Name = "txtKLmag"
        Me.txtKLmag.Size = New System.Drawing.Size(66, 23)
        Me.txtKLmag.TabIndex = 694
        Me.txtKLmag.Tag = "1.8"
        Me.txtKLmag.Text = "0.5917"
        '
        'Label25
        '
        Me.Label25.AutoSize = True
        Me.Label25.Font = New System.Drawing.Font("Verdana", 9.75!, System.Drawing.FontStyle.Regular, System.Drawing.GraphicsUnit.Point, CType(0, Byte))
        Me.Label25.ForeColor = System.Drawing.Color.Red
        Me.Label25.Location = New System.Drawing.Point(27, 159)
        Me.Label25.Name = "Label25"
        Me.Label25.Size = New System.Drawing.Size(37, 16)
        Me.Label25.TabIndex = 689
        Me.Label25.Text = "ICaL"
        '
        'Label30
        '
        Me.Label30.AutoSize = True
        Me.Label30.Font = New System.Drawing.Font("Verdana", 9.75!, System.Drawing.FontStyle.Regular, System.Drawing.GraphicsUnit.Point, CType(0, Byte))
        Me.Label30.Location = New System.Drawing.Point(27, 134)
        Me.Label30.Name = "Label30"
        Me.Label30.Size = New System.Drawing.Size(29, 16)
        Me.Label30.TabIndex = 688
        Me.Label30.Text = "IK1"
        '
        'IKr
        '
        Me.IKr.AutoSize = True
        Me.IKr.Font = New System.Drawing.Font("Verdana", 9.75!, System.Drawing.FontStyle.Regular, System.Drawing.GraphicsUnit.Point, CType(0, Byte))
        Me.IKr.ForeColor = System.Drawing.Color.FromArgb(CType(CType(0, Byte), Integer), CType(CType(0, Byte), Integer), CType(CType(192, Byte), Integer))
        Me.IKr.Location = New System.Drawing.Point(27, 111)
        Me.IKr.Name = "IKr"
        Me.IKr.Size = New System.Drawing.Size(26, 16)
        Me.IKr.TabIndex = 687
        Me.IKr.Text = "IKr"
        '
        'txtICaLmag
        '
        Me.txtICaLmag.Font = New System.Drawing.Font("Verdana", 9.75!, System.Drawing.FontStyle.Regular, System.Drawing.GraphicsUnit.Point, CType(0, Byte))
        Me.txtICaLmag.Location = New System.Drawing.Point(66, 156)
        Me.txtICaLmag.Name = "txtICaLmag"
        Me.txtICaLmag.Size = New System.Drawing.Size(66, 23)
        Me.txtICaLmag.TabIndex = 686
        Me.txtICaLmag.Tag = "0.6809"
        Me.txtICaLmag.Text = "0.9991"
        '
        'txtIK1mag
        '
        Me.txtIK1mag.Font = New System.Drawing.Font("Verdana", 9.75!, System.Drawing.FontStyle.Regular, System.Drawing.GraphicsUnit.Point, CType(0, Byte))
        Me.txtIK1mag.Location = New System.Drawing.Point(66, 131)
        Me.txtIK1mag.Name = "txtIK1mag"
        Me.txtIK1mag.Size = New System.Drawing.Size(66, 23)
        Me.txtIK1mag.TabIndex = 685
        Me.txtIK1mag.Text = "0.15"
        '
        'Label15
        '
        Me.Label15.AutoSize = True
        Me.Label15.Font = New System.Drawing.Font("Verdana", 9.0!, System.Drawing.FontStyle.Regular, System.Drawing.GraphicsUnit.Point, CType(0, Byte))
        Me.Label15.Location = New System.Drawing.Point(4, 407)
        Me.Label15.Name = "Label15"
        Me.Label15.Size = New System.Drawing.Size(55, 14)
        Me.Label15.TabIndex = 709
        Me.Label15.Text = "VshitNa"
        '
        'txtIKrmag
        '
        Me.txtIKrmag.Font = New System.Drawing.Font("Verdana", 9.75!, System.Drawing.FontStyle.Regular, System.Drawing.GraphicsUnit.Point, CType(0, Byte))
        Me.txtIKrmag.Location = New System.Drawing.Point(66, 108)
        Me.txtIKrmag.Name = "txtIKrmag"
        Me.txtIKrmag.Size = New System.Drawing.Size(66, 23)
        Me.txtIKrmag.TabIndex = 684
        Me.txtIKrmag.Text = "0.8"
        '
        'Label53
        '
        Me.Label53.AutoSize = True
        Me.Label53.Font = New System.Drawing.Font("Verdana", 9.75!, System.Drawing.FontStyle.Regular, System.Drawing.GraphicsUnit.Point, CType(0, Byte))
        Me.Label53.Location = New System.Drawing.Point(8, 65)
        Me.Label53.Name = "Label53"
        Me.Label53.Size = New System.Drawing.Size(67, 16)
        Me.Label53.TabIndex = 681
        Me.Label53.Text = "NaL ratio"
        '
        'txtNaLratio
        '
        Me.txtNaLratio.Font = New System.Drawing.Font("Verdana", 9.75!, System.Drawing.FontStyle.Regular, System.Drawing.GraphicsUnit.Point, CType(0, Byte))
        Me.txtNaLratio.Location = New System.Drawing.Point(104, 61)
        Me.txtNaLratio.Name = "txtNaLratio"
        Me.txtNaLratio.Size = New System.Drawing.Size(49, 23)
        Me.txtNaLratio.TabIndex = 680
        Me.txtNaLratio.Text = "0.04"
        '
        'Label51
        '
        Me.Label51.AutoSize = True
        Me.Label51.Font = New System.Drawing.Font("Verdana", 9.75!, System.Drawing.FontStyle.Regular, System.Drawing.GraphicsUnit.Point, CType(0, Byte))
        Me.Label51.Location = New System.Drawing.Point(0, 39)
        Me.Label51.Name = "Label51"
        Me.Label51.Size = New System.Drawing.Size(101, 16)
        Me.Label51.TabIndex = 679
        Me.Label51.Text = "orgExtraVshift"
        '
        'Label6
        '
        Me.Label6.AutoSize = True
        Me.Label6.Font = New System.Drawing.Font("Verdana", 9.0!, System.Drawing.FontStyle.Regular, System.Drawing.GraphicsUnit.Point, CType(0, Byte))
        Me.Label6.Location = New System.Drawing.Point(21, 430)
        Me.Label6.Name = "Label6"
        Me.Label6.Size = New System.Drawing.Size(36, 14)
        Me.Label6.TabIndex = 706
        Me.Label6.Text = "ICaT"
        '
        'txtBaseVshift
        '
        Me.txtBaseVshift.Font = New System.Drawing.Font("Verdana", 9.0!, System.Drawing.FontStyle.Regular, System.Drawing.GraphicsUnit.Point, CType(0, Byte))
        Me.txtBaseVshift.Location = New System.Drawing.Point(105, 15)
        Me.txtBaseVshift.Name = "txtBaseVshift"
        Me.txtBaseVshift.Size = New System.Drawing.Size(49, 22)
        Me.txtBaseVshift.TabIndex = 678
        Me.txtBaseVshift.Text = "-3"
        '
        'Label48
        '
        Me.Label48.AutoSize = True
        Me.Label48.Font = New System.Drawing.Font("Verdana", 9.75!, System.Drawing.FontStyle.Regular, System.Drawing.GraphicsUnit.Point, CType(0, Byte))
        Me.Label48.Location = New System.Drawing.Point(27, 17)
        Me.Label48.Name = "Label48"
        Me.Label48.Size = New System.Drawing.Size(77, 16)
        Me.Label48.TabIndex = 677
        Me.Label48.Text = "baseVshift"
        '
        'Label9
        '
        Me.Label9.AutoSize = True
        Me.Label9.Font = New System.Drawing.Font("Verdana", 9.0!, System.Drawing.FontStyle.Regular, System.Drawing.GraphicsUnit.Point, CType(0, Byte))
        Me.Label9.ForeColor = System.Drawing.Color.FromArgb(CType(CType(255, Byte), Integer), CType(CType(128, Byte), Integer), CType(CType(0, Byte), Integer))
        Me.Label9.Location = New System.Drawing.Point(26, 358)
        Me.Label9.Name = "Label9"
        Me.Label9.Size = New System.Drawing.Size(33, 14)
        Me.Label9.TabIndex = 699
        Me.Label9.Text = "IKur"
        '
        'txtorgExtraVshift
        '
        Me.txtorgExtraVshift.Font = New System.Drawing.Font("Verdana", 9.0!, System.Drawing.FontStyle.Regular, System.Drawing.GraphicsUnit.Point, CType(0, Byte))
        Me.txtorgExtraVshift.Location = New System.Drawing.Point(105, 37)
        Me.txtorgExtraVshift.Name = "txtorgExtraVshift"
        Me.txtorgExtraVshift.Size = New System.Drawing.Size(49, 22)
        Me.txtorgExtraVshift.TabIndex = 676
        Me.txtorgExtraVshift.Tag = ""
        Me.txtorgExtraVshift.Text = "-6.5"
        '
        'txtICaTmag
        '
        Me.txtICaTmag.Font = New System.Drawing.Font("Verdana", 9.0!, System.Drawing.FontStyle.Regular, System.Drawing.GraphicsUnit.Point, CType(0, Byte))
        Me.txtICaTmag.Location = New System.Drawing.Point(63, 427)
        Me.txtICaTmag.Name = "txtICaTmag"
        Me.txtICaTmag.Size = New System.Drawing.Size(67, 22)
        Me.txtICaTmag.TabIndex = 705
        Me.txtICaTmag.Text = "0"
        '
        'Label34
        '
        Me.Label34.AutoSize = True
        Me.Label34.Font = New System.Drawing.Font("Verdana", 9.75!, System.Drawing.FontStyle.Regular, System.Drawing.GraphicsUnit.Point, CType(0, Byte))
        Me.Label34.Location = New System.Drawing.Point(8, 657)
        Me.Label34.Name = "Label34"
        Me.Label34.Size = New System.Drawing.Size(61, 16)
        Me.Label34.TabIndex = 672
        Me.Label34.Text = "MDP mV"
        '
        'txtIKurmag
        '
        Me.txtIKurmag.Font = New System.Drawing.Font("Verdana", 9.0!, System.Drawing.FontStyle.Regular, System.Drawing.GraphicsUnit.Point, CType(0, Byte))
        Me.txtIKurmag.Location = New System.Drawing.Point(63, 355)
        Me.txtIKurmag.Name = "txtIKurmag"
        Me.txtIKurmag.Size = New System.Drawing.Size(67, 22)
        Me.txtIKurmag.TabIndex = 698
        Me.txtIKurmag.Tag = "28.5634"
        Me.txtIKurmag.Text = "21.2547"
        '
        'txtOS
        '
        Me.txtOS.Font = New System.Drawing.Font("Verdana", 9.75!, System.Drawing.FontStyle.Regular, System.Drawing.GraphicsUnit.Point, CType(0, Byte))
        Me.txtOS.Location = New System.Drawing.Point(210, 627)
        Me.txtOS.Name = "txtOS"
        Me.txtOS.Size = New System.Drawing.Size(56, 23)
        Me.txtOS.TabIndex = 671
        Me.txtOS.Text = "0"
        '
        'Label10
        '
        Me.Label10.AutoSize = True
        Me.Label10.Font = New System.Drawing.Font("Verdana", 9.75!, System.Drawing.FontStyle.Regular, System.Drawing.GraphicsUnit.Point, CType(0, Byte))
        Me.Label10.Location = New System.Drawing.Point(147, 630)
        Me.Label10.Name = "Label10"
        Me.Label10.Size = New System.Drawing.Size(52, 16)
        Me.Label10.TabIndex = 670
        Me.Label10.Text = "OS mV"
        '
        'Label13
        '
        Me.Label13.AutoSize = True
        Me.Label13.Font = New System.Drawing.Font("Verdana", 9.75!, System.Drawing.FontStyle.Regular, System.Drawing.GraphicsUnit.Point, CType(0, Byte))
        Me.Label13.ForeColor = System.Drawing.Color.Fuchsia
        Me.Label13.Location = New System.Drawing.Point(29, 306)
        Me.Label13.Name = "Label13"
        Me.Label13.Size = New System.Drawing.Size(28, 16)
        Me.Label13.TabIndex = 700
        Me.Label13.Text = "IKs"
        '
        'Label8
        '
        Me.Label8.AutoSize = True
        Me.Label8.Font = New System.Drawing.Font("Verdana", 9.75!, System.Drawing.FontStyle.Regular, System.Drawing.GraphicsUnit.Point, CType(0, Byte))
        Me.Label8.Location = New System.Drawing.Point(149, 655)
        Me.Label8.Name = "Label8"
        Me.Label8.Size = New System.Drawing.Size(57, 16)
        Me.Label8.TabIndex = 669
        Me.Label8.Text = "APD ms"
        '
        'txtIKsmag
        '
        Me.txtIKsmag.Font = New System.Drawing.Font("Verdana", 9.75!, System.Drawing.FontStyle.Regular, System.Drawing.GraphicsUnit.Point, CType(0, Byte))
        Me.txtIKsmag.ForeColor = System.Drawing.Color.FromArgb(CType(CType(0, Byte), Integer), CType(CType(0, Byte), Integer), CType(CType(0, Byte), Integer), CType(CType(5, Byte), Integer))
        Me.txtIKsmag.Location = New System.Drawing.Point(65, 303)
        Me.txtIKsmag.Name = "txtIKsmag"
        Me.txtIKsmag.Size = New System.Drawing.Size(66, 23)
        Me.txtIKsmag.TabIndex = 697
        Me.txtIKsmag.Text = "1.3"
        '
        'txtMDP
        '
        Me.txtMDP.Font = New System.Drawing.Font("Verdana", 9.75!, System.Drawing.FontStyle.Regular, System.Drawing.GraphicsUnit.Point, CType(0, Byte))
        Me.txtMDP.Location = New System.Drawing.Point(87, 652)
        Me.txtMDP.Name = "txtMDP"
        Me.txtMDP.Size = New System.Drawing.Size(56, 23)
        Me.txtMDP.TabIndex = 673
        Me.txtMDP.Text = "0"
        '
        'Label27
        '
        Me.Label27.AutoSize = True
        Me.Label27.Font = New System.Drawing.Font("Verdana", 9.75!, System.Drawing.FontStyle.Regular, System.Drawing.GraphicsUnit.Point, CType(0, Byte))
        Me.Label27.Location = New System.Drawing.Point(10, 579)
        Me.Label27.Name = "Label27"
        Me.Label27.Size = New System.Drawing.Size(117, 16)
        Me.Label27.TabIndex = 664
        Me.Label27.Text = "correction f NCX"
        '
        'Label14
        '
        Me.Label14.AutoSize = True
        Me.Label14.Font = New System.Drawing.Font("Verdana", 9.0!, System.Drawing.FontStyle.Regular, System.Drawing.GraphicsUnit.Point, CType(0, Byte))
        Me.Label14.ForeColor = System.Drawing.Color.FromArgb(CType(CType(192, Byte), Integer), CType(CType(0, Byte), Integer), CType(CType(192, Byte), Integer))
        Me.Label14.Location = New System.Drawing.Point(27, 383)
        Me.Label14.Name = "Label14"
        Me.Label14.Size = New System.Drawing.Size(29, 14)
        Me.Label14.TabIndex = 704
        Me.Label14.Text = "INa"
        '
        'rNa
        '
        Me.rNa.Font = New System.Drawing.Font("Verdana", 9.75!, System.Drawing.FontStyle.Regular, System.Drawing.GraphicsUnit.Point, CType(0, Byte))
        Me.rNa.ForeColor = System.Drawing.Color.Red
        Me.rNa.Location = New System.Drawing.Point(140, 380)
        Me.rNa.Name = "rNa"
        Me.rNa.Size = New System.Drawing.Size(63, 23)
        Me.rNa.TabIndex = 3
        Me.rNa.Text = "1"
        '
        'Label1
        '
        Me.Label1.AutoSize = True
        Me.Label1.Font = New System.Drawing.Font("Verdana", 9.75!, System.Drawing.FontStyle.Regular, System.Drawing.GraphicsUnit.Point, CType(0, Byte))
        Me.Label1.Location = New System.Drawing.Point(6, 630)
        Me.Label1.Name = "Label1"
        Me.Label1.Size = New System.Drawing.Size(78, 16)
        Me.Label1.TabIndex = 557
        Me.Label1.Text = "cycle L ms"
        '
        'Label18
        '
        Me.Label18.AutoSize = True
        Me.Label18.Font = New System.Drawing.Font("Verdana", 9.75!, System.Drawing.FontStyle.Regular, System.Drawing.GraphicsUnit.Point, CType(0, Byte))
        Me.Label18.Location = New System.Drawing.Point(25, 335)
        Me.Label18.Name = "Label18"
        Me.Label18.Size = New System.Drawing.Size(35, 16)
        Me.Label18.TabIndex = 702
        Me.Label18.Text = "IKto"
        '
        'txtCycleL
        '
        Me.txtCycleL.Font = New System.Drawing.Font("Verdana", 9.75!, System.Drawing.FontStyle.Regular, System.Drawing.GraphicsUnit.Point, CType(0, Byte))
        Me.txtCycleL.Location = New System.Drawing.Point(87, 627)
        Me.txtCycleL.Name = "txtCycleL"
        Me.txtCycleL.Size = New System.Drawing.Size(56, 23)
        Me.txtCycleL.TabIndex = 668
        Me.txtCycleL.Text = "0"
        '
        'Label28
        '
        Me.Label28.AutoSize = True
        Me.Label28.Font = New System.Drawing.Font("Verdana", 9.75!, System.Drawing.FontStyle.Regular, System.Drawing.GraphicsUnit.Point, CType(0, Byte))
        Me.Label28.Location = New System.Drawing.Point(137, 605)
        Me.Label28.Name = "Label28"
        Me.Label28.Size = New System.Drawing.Size(69, 16)
        Me.Label28.TabIndex = 666
        Me.Label28.Text = "total Cai "
        '
        'txtIKtomag
        '
        Me.txtIKtomag.Font = New System.Drawing.Font("Verdana", 9.75!, System.Drawing.FontStyle.Regular, System.Drawing.GraphicsUnit.Point, CType(0, Byte))
        Me.txtIKtomag.Location = New System.Drawing.Point(62, 328)
        Me.txtIKtomag.Name = "txtIKtomag"
        Me.txtIKtomag.Size = New System.Drawing.Size(68, 23)
        Me.txtIKtomag.TabIndex = 701
        Me.txtIKtomag.Text = "6.2258"
        '
        'txtAPD
        '
        Me.txtAPD.Font = New System.Drawing.Font("Verdana", 9.75!, System.Drawing.FontStyle.Regular, System.Drawing.GraphicsUnit.Point, CType(0, Byte))
        Me.txtAPD.Location = New System.Drawing.Point(207, 652)
        Me.txtAPD.Name = "txtAPD"
        Me.txtAPD.Size = New System.Drawing.Size(56, 23)
        Me.txtAPD.TabIndex = 670
        Me.txtAPD.Text = "0"
        '
        'rKr
        '
        Me.rKr.Font = New System.Drawing.Font("Verdana", 9.75!, System.Drawing.FontStyle.Regular, System.Drawing.GraphicsUnit.Point, CType(0, Byte))
        Me.rKr.ForeColor = System.Drawing.Color.Red
        Me.rKr.Location = New System.Drawing.Point(139, 108)
        Me.rKr.Name = "rKr"
        Me.rKr.Size = New System.Drawing.Size(63, 23)
        Me.rKr.TabIndex = 14
        Me.rKr.Text = "1"
        '
        'txtINamag
        '
        Me.txtINamag.Font = New System.Drawing.Font("Verdana", 9.0!, System.Drawing.FontStyle.Regular, System.Drawing.GraphicsUnit.Point, CType(0, Byte))
        Me.txtINamag.Location = New System.Drawing.Point(63, 380)
        Me.txtINamag.Name = "txtINamag"
        Me.txtINamag.Size = New System.Drawing.Size(67, 22)
        Me.txtINamag.TabIndex = 703
        Me.txtINamag.Text = "0.33"
        '
        'txtTotalCai
        '
        Me.txtTotalCai.Font = New System.Drawing.Font("Verdana", 12.0!, System.Drawing.FontStyle.Bold, System.Drawing.GraphicsUnit.Point, CType(0, Byte))
        Me.txtTotalCai.Location = New System.Drawing.Point(211, 600)
        Me.txtTotalCai.Name = "txtTotalCai"
        Me.txtTotalCai.Size = New System.Drawing.Size(71, 27)
        Me.txtTotalCai.TabIndex = 665
        Me.txtTotalCai.Text = "1"
        '
        'rK1
        '
        Me.rK1.Font = New System.Drawing.Font("Verdana", 9.75!, System.Drawing.FontStyle.Regular, System.Drawing.GraphicsUnit.Point, CType(0, Byte))
        Me.rK1.ForeColor = System.Drawing.Color.Red
        Me.rK1.Location = New System.Drawing.Point(139, 129)
        Me.rK1.Name = "rK1"
        Me.rK1.Size = New System.Drawing.Size(63, 23)
        Me.rK1.TabIndex = 15
        Me.rK1.Text = "1"
        '
        'txtcrfNCX
        '
        Me.txtcrfNCX.Font = New System.Drawing.Font("Verdana", 9.75!, System.Drawing.FontStyle.Regular, System.Drawing.GraphicsUnit.Point, CType(0, Byte))
        Me.txtcrfNCX.Location = New System.Drawing.Point(141, 576)
        Me.txtcrfNCX.Name = "txtcrfNCX"
        Me.txtcrfNCX.Size = New System.Drawing.Size(59, 23)
        Me.txtcrfNCX.TabIndex = 663
        Me.txtcrfNCX.Text = "1"
        '
        'Label22
        '
        Me.Label22.AutoSize = True
        Me.Label22.Font = New System.Drawing.Font("Verdana", 9.75!, System.Drawing.FontStyle.Regular, System.Drawing.GraphicsUnit.Point, CType(0, Byte))
        Me.Label22.Location = New System.Drawing.Point(18, 258)
        Me.Label22.Name = "Label22"
        Me.Label22.Size = New System.Drawing.Size(48, 16)
        Me.Label22.TabIndex = 695
        Me.Label22.Text = "IbNSC"
        '
        'rVshiftCaL
        '
        Me.rVshiftCaL.Font = New System.Drawing.Font("Verdana", 9.75!, System.Drawing.FontStyle.Regular, System.Drawing.GraphicsUnit.Point, CType(0, Byte))
        Me.rVshiftCaL.ForeColor = System.Drawing.Color.Black
        Me.rVshiftCaL.Location = New System.Drawing.Point(139, 206)
        Me.rVshiftCaL.Name = "rVshiftCaL"
        Me.rVshiftCaL.Size = New System.Drawing.Size(63, 23)
        Me.rVshiftCaL.TabIndex = 549
        Me.rVshiftCaL.Text = "1"
        '
        'txtcrfNaK
        '
        Me.txtcrfNaK.Font = New System.Drawing.Font("Verdana", 9.75!, System.Drawing.FontStyle.Regular, System.Drawing.GraphicsUnit.Point, CType(0, Byte))
        Me.txtcrfNaK.Location = New System.Drawing.Point(141, 525)
        Me.txtcrfNaK.Name = "txtcrfNaK"
        Me.txtcrfNaK.Size = New System.Drawing.Size(62, 23)
        Me.txtcrfNaK.TabIndex = 542
        Me.txtcrfNaK.Text = "1.1279"
        '
        'rCaL
        '
        Me.rCaL.Font = New System.Drawing.Font("Verdana", 9.75!, System.Drawing.FontStyle.Regular, System.Drawing.GraphicsUnit.Point, CType(0, Byte))
        Me.rCaL.ForeColor = System.Drawing.Color.Red
        Me.rCaL.Location = New System.Drawing.Point(139, 154)
        Me.rCaL.Name = "rCaL"
        Me.rCaL.Size = New System.Drawing.Size(63, 23)
        Me.rCaL.TabIndex = 16
        Me.rCaL.Text = "1"
        '
        'txtIbNSCmag
        '
        Me.txtIbNSCmag.Font = New System.Drawing.Font("Verdana", 9.75!, System.Drawing.FontStyle.Regular, System.Drawing.GraphicsUnit.Point, CType(0, Byte))
        Me.txtIbNSCmag.ForeColor = System.Drawing.Color.Black
        Me.txtIbNSCmag.Location = New System.Drawing.Point(66, 255)
        Me.txtIbNSCmag.Name = "txtIbNSCmag"
        Me.txtIbNSCmag.Size = New System.Drawing.Size(66, 23)
        Me.txtIbNSCmag.TabIndex = 693
        Me.txtIbNSCmag.Text = "3.1"
        '
        'Label24
        '
        Me.Label24.AutoSize = True
        Me.Label24.Font = New System.Drawing.Font("Verdana", 9.0!, System.Drawing.FontStyle.Regular, System.Drawing.GraphicsUnit.Point, CType(0, Byte))
        Me.Label24.ForeColor = System.Drawing.Color.Maroon
        Me.Label24.Location = New System.Drawing.Point(26, 455)
        Me.Label24.Name = "Label24"
        Me.Label24.Size = New System.Drawing.Size(24, 14)
        Me.Label24.TabIndex = 692
        Me.Label24.Text = "Ist"
        '
        'txtKi
        '
        Me.txtKi.Font = New System.Drawing.Font("Verdana", 9.0!, System.Drawing.FontStyle.Regular, System.Drawing.GraphicsUnit.Point, CType(0, Byte))
        Me.txtKi.Location = New System.Drawing.Point(202, 492)
        Me.txtKi.Name = "txtKi"
        Me.txtKi.Size = New System.Drawing.Size(60, 22)
        Me.txtKi.TabIndex = 521
        Me.txtKi.Text = "0"
        '
        'rCaT
        '
        Me.rCaT.Font = New System.Drawing.Font("Verdana", 9.0!, System.Drawing.FontStyle.Regular, System.Drawing.GraphicsUnit.Point, CType(0, Byte))
        Me.rCaT.Location = New System.Drawing.Point(138, 428)
        Me.rCaT.Name = "rCaT"
        Me.rCaT.Size = New System.Drawing.Size(64, 22)
        Me.rCaT.TabIndex = 519
        Me.rCaT.Text = "1"
        '
        'Label29
        '
        Me.Label29.AutoSize = True
        Me.Label29.Font = New System.Drawing.Font("Verdana", 9.75!, System.Drawing.FontStyle.Regular, System.Drawing.GraphicsUnit.Point, CType(0, Byte))
        Me.Label29.ForeColor = System.Drawing.Color.FromArgb(CType(CType(0, Byte), Integer), CType(CType(192, Byte), Integer), CType(CType(0, Byte), Integer))
        Me.Label29.Location = New System.Drawing.Point(30, 282)
        Me.Label29.Name = "Label29"
        Me.Label29.Size = New System.Drawing.Size(29, 16)
        Me.Label29.TabIndex = 691
        Me.Label29.Text = "Iha"
        '
        'rextraVshift
        '
        Me.rextraVshift.Font = New System.Drawing.Font("Verdana", 9.0!, System.Drawing.FontStyle.Regular, System.Drawing.GraphicsUnit.Point, CType(0, Byte))
        Me.rextraVshift.ForeColor = System.Drawing.Color.Black
        Me.rextraVshift.Location = New System.Drawing.Point(156, 37)
        Me.rextraVshift.Name = "rextraVshift"
        Me.rextraVshift.Size = New System.Drawing.Size(62, 22)
        Me.rextraVshift.TabIndex = 548
        Me.rextraVshift.Text = "1"
        '
        'rst
        '
        Me.rst.Font = New System.Drawing.Font("Verdana", 9.0!, System.Drawing.FontStyle.Regular, System.Drawing.GraphicsUnit.Point, CType(0, Byte))
        Me.rst.ForeColor = System.Drawing.Color.Black
        Me.rst.Location = New System.Drawing.Point(138, 452)
        Me.rst.Name = "rst"
        Me.rst.Size = New System.Drawing.Size(64, 22)
        Me.rst.TabIndex = 4
        Me.rst.Text = "1"
        '
        'txtIhamag
        '
        Me.txtIhamag.Font = New System.Drawing.Font("Verdana", 9.75!, System.Drawing.FontStyle.Regular, System.Drawing.GraphicsUnit.Point, CType(0, Byte))
        Me.txtIhamag.Location = New System.Drawing.Point(65, 278)
        Me.txtIhamag.Name = "txtIhamag"
        Me.txtIhamag.Size = New System.Drawing.Size(66, 23)
        Me.txtIhamag.TabIndex = 690
        Me.txtIhamag.Text = "3.6"
        '
        'rbNSC
        '
        Me.rbNSC.Font = New System.Drawing.Font("Verdana", 9.75!, System.Drawing.FontStyle.Regular, System.Drawing.GraphicsUnit.Point, CType(0, Byte))
        Me.rbNSC.ForeColor = System.Drawing.Color.Black
        Me.rbNSC.Location = New System.Drawing.Point(139, 254)
        Me.rbNSC.Name = "rbNSC"
        Me.rbNSC.Size = New System.Drawing.Size(63, 23)
        Me.rbNSC.TabIndex = 490
        Me.rbNSC.Text = "1"
        '
        'NVshiftCaL
        '
        Me.NVshiftCaL.Font = New System.Drawing.Font("Verdana", 9.75!, System.Drawing.FontStyle.Regular, System.Drawing.GraphicsUnit.Point, CType(0, Byte))
        Me.NVshiftCaL.ForeColor = System.Drawing.Color.Black
        Me.NVshiftCaL.Location = New System.Drawing.Point(206, 206)
        Me.NVshiftCaL.Name = "NVshiftCaL"
        Me.NVshiftCaL.Size = New System.Drawing.Size(63, 23)
        Me.NVshiftCaL.TabIndex = 661
        Me.NVshiftCaL.Text = "0"
        '
        'Label47
        '
        Me.Label47.AutoSize = True
        Me.Label47.Font = New System.Drawing.Font("Verdana", 9.75!, System.Drawing.FontStyle.Regular, System.Drawing.GraphicsUnit.Point, CType(0, Byte))
        Me.Label47.Location = New System.Drawing.Point(12, 528)
        Me.Label47.Name = "Label47"
        Me.Label47.Size = New System.Drawing.Size(115, 16)
        Me.Label47.TabIndex = 543
        Me.Label47.Text = "correction f NaK"
        '
        'rKL
        '
        Me.rKL.Font = New System.Drawing.Font("Verdana", 9.75!, System.Drawing.FontStyle.Regular, System.Drawing.GraphicsUnit.Point, CType(0, Byte))
        Me.rKL.ForeColor = System.Drawing.Color.Black
        Me.rKL.Location = New System.Drawing.Point(139, 179)
        Me.rKL.Name = "rKL"
        Me.rKL.Size = New System.Drawing.Size(63, 23)
        Me.rKL.TabIndex = 496
        Me.rKL.Text = "1"
        '
        'NextraVshift
        '
        Me.NextraVshift.Font = New System.Drawing.Font("Verdana", 9.0!, System.Drawing.FontStyle.Regular, System.Drawing.GraphicsUnit.Point, CType(0, Byte))
        Me.NextraVshift.ForeColor = System.Drawing.Color.Black
        Me.NextraVshift.Location = New System.Drawing.Point(223, 36)
        Me.NextraVshift.Name = "NextraVshift"
        Me.NextraVshift.Size = New System.Drawing.Size(62, 22)
        Me.NextraVshift.TabIndex = 660
        Me.NextraVshift.Text = "1"
        '
        'rKto
        '
        Me.rKto.Font = New System.Drawing.Font("Verdana", 9.75!, System.Drawing.FontStyle.Regular, System.Drawing.GraphicsUnit.Point, CType(0, Byte))
        Me.rKto.ForeColor = System.Drawing.Color.Black
        Me.rKto.Location = New System.Drawing.Point(140, 329)
        Me.rKto.Name = "rKto"
        Me.rKto.Size = New System.Drawing.Size(63, 23)
        Me.rKto.TabIndex = 508
        Me.rKto.Text = "1"
        '
        'NVshiftNa
        '
        Me.NVshiftNa.Font = New System.Drawing.Font("Verdana", 9.75!, System.Drawing.FontStyle.Regular, System.Drawing.GraphicsUnit.Point, CType(0, Byte))
        Me.NVshiftNa.ForeColor = System.Drawing.Color.Black
        Me.NVshiftNa.Location = New System.Drawing.Point(206, 403)
        Me.NVshiftNa.Name = "NVshiftNa"
        Me.NVshiftNa.Size = New System.Drawing.Size(63, 23)
        Me.NVshiftNa.TabIndex = 662
        Me.NVshiftNa.Text = "0"
        '
        'rVshiftNa
        '
        Me.rVshiftNa.Font = New System.Drawing.Font("Verdana", 9.75!, System.Drawing.FontStyle.Regular, System.Drawing.GraphicsUnit.Point, CType(0, Byte))
        Me.rVshiftNa.ForeColor = System.Drawing.Color.Black
        Me.rVshiftNa.Location = New System.Drawing.Point(139, 403)
        Me.rVshiftNa.Name = "rVshiftNa"
        Me.rVshiftNa.Size = New System.Drawing.Size(63, 23)
        Me.rVshiftNa.TabIndex = 596
        Me.rVshiftNa.Text = "1"
        '
        'txtIstmag
        '
        Me.txtIstmag.Font = New System.Drawing.Font("Verdana", 9.0!, System.Drawing.FontStyle.Regular, System.Drawing.GraphicsUnit.Point, CType(0, Byte))
        Me.txtIstmag.Location = New System.Drawing.Point(63, 452)
        Me.txtIstmag.Name = "txtIstmag"
        Me.txtIstmag.Size = New System.Drawing.Size(67, 22)
        Me.txtIstmag.TabIndex = 683
        Me.txtIstmag.Text = "0.99507"
        '
        'NIhamag
        '
        Me.NIhamag.Font = New System.Drawing.Font("Verdana", 9.75!, System.Drawing.FontStyle.Regular, System.Drawing.GraphicsUnit.Point, CType(0, Byte))
        Me.NIhamag.ForeColor = System.Drawing.Color.Black
        Me.NIhamag.Location = New System.Drawing.Point(205, 279)
        Me.NIhamag.Name = "NIhamag"
        Me.NIhamag.Size = New System.Drawing.Size(63, 23)
        Me.NIhamag.TabIndex = 659
        Me.NIhamag.Text = "0"
        '
        'rKs
        '
        Me.rKs.Font = New System.Drawing.Font("Verdana", 9.75!, System.Drawing.FontStyle.Regular, System.Drawing.GraphicsUnit.Point, CType(0, Byte))
        Me.rKs.ForeColor = System.Drawing.Color.Black
        Me.rKs.Location = New System.Drawing.Point(139, 304)
        Me.rKs.Name = "rKs"
        Me.rKs.Size = New System.Drawing.Size(63, 23)
        Me.rKs.TabIndex = 506
        Me.rKs.Text = "1"
        '
        'NICaTmag
        '
        Me.NICaTmag.Font = New System.Drawing.Font("Verdana", 9.0!, System.Drawing.FontStyle.Regular, System.Drawing.GraphicsUnit.Point, CType(0, Byte))
        Me.NICaTmag.Location = New System.Drawing.Point(206, 427)
        Me.NICaTmag.Name = "NICaTmag"
        Me.NICaTmag.Size = New System.Drawing.Size(64, 22)
        Me.NICaTmag.TabIndex = 658
        Me.NICaTmag.Text = "0"
        '
        'rha
        '
        Me.rha.Font = New System.Drawing.Font("Verdana", 9.75!, System.Drawing.FontStyle.Regular, System.Drawing.GraphicsUnit.Point, CType(0, Byte))
        Me.rha.ForeColor = System.Drawing.Color.Black
        Me.rha.Location = New System.Drawing.Point(138, 279)
        Me.rha.Name = "rha"
        Me.rha.Size = New System.Drawing.Size(63, 23)
        Me.rha.TabIndex = 545
        Me.rha.Text = "1"
        '
        'NIKurmag
        '
        Me.NIKurmag.Font = New System.Drawing.Font("Verdana", 9.0!, System.Drawing.FontStyle.Regular, System.Drawing.GraphicsUnit.Point, CType(0, Byte))
        Me.NIKurmag.ForeColor = System.Drawing.Color.Black
        Me.NIKurmag.Location = New System.Drawing.Point(206, 355)
        Me.NIKurmag.Name = "NIKurmag"
        Me.NIKurmag.Size = New System.Drawing.Size(64, 22)
        Me.NIKurmag.TabIndex = 655
        Me.NIKurmag.Text = "0"
        '
        'rKur
        '
        Me.rKur.Font = New System.Drawing.Font("Verdana", 9.0!, System.Drawing.FontStyle.Regular, System.Drawing.GraphicsUnit.Point, CType(0, Byte))
        Me.rKur.ForeColor = System.Drawing.Color.Black
        Me.rKur.Location = New System.Drawing.Point(139, 355)
        Me.rKur.Name = "rKur"
        Me.rKur.Size = New System.Drawing.Size(64, 22)
        Me.rKur.TabIndex = 506
        Me.rKur.Text = "1"
        '
        'Label20
        '
        Me.Label20.AutoSize = True
        Me.Label20.Font = New System.Drawing.Font("Verdana", 9.0!, System.Drawing.FontStyle.Regular, System.Drawing.GraphicsUnit.Point, CType(0, Byte))
        Me.Label20.Location = New System.Drawing.Point(143, 495)
        Me.Label20.Name = "Label20"
        Me.Label20.Size = New System.Drawing.Size(53, 14)
        Me.Label20.TabIndex = 522
        Me.Label20.Text = "Ki (mM)"
        '
        'NIKsmag
        '
        Me.NIKsmag.Font = New System.Drawing.Font("Verdana", 9.75!, System.Drawing.FontStyle.Regular, System.Drawing.GraphicsUnit.Point, CType(0, Byte))
        Me.NIKsmag.ForeColor = System.Drawing.Color.Black
        Me.NIKsmag.Location = New System.Drawing.Point(206, 304)
        Me.NIKsmag.Name = "NIKsmag"
        Me.NIKsmag.Size = New System.Drawing.Size(63, 23)
        Me.NIKsmag.TabIndex = 656
        Me.NIKsmag.Text = "0"
        '
        'NIKtomag
        '
        Me.NIKtomag.Font = New System.Drawing.Font("Verdana", 9.75!, System.Drawing.FontStyle.Regular, System.Drawing.GraphicsUnit.Point, CType(0, Byte))
        Me.NIKtomag.ForeColor = System.Drawing.Color.Black
        Me.NIKtomag.Location = New System.Drawing.Point(207, 329)
        Me.NIKtomag.Name = "NIKtomag"
        Me.NIKtomag.Size = New System.Drawing.Size(63, 23)
        Me.NIKtomag.TabIndex = 657
        Me.NIKtomag.Text = "0"
        '
        'NIstmag
        '
        Me.NIstmag.Font = New System.Drawing.Font("Verdana", 9.0!, System.Drawing.FontStyle.Regular, System.Drawing.GraphicsUnit.Point, CType(0, Byte))
        Me.NIstmag.ForeColor = System.Drawing.Color.Black
        Me.NIstmag.Location = New System.Drawing.Point(206, 452)
        Me.NIstmag.Name = "NIstmag"
        Me.NIstmag.Size = New System.Drawing.Size(64, 22)
        Me.NIstmag.TabIndex = 649
        Me.NIstmag.Text = "0"
        '
        'Label21
        '
        Me.Label21.AutoSize = True
        Me.Label21.Font = New System.Drawing.Font("Verdana", 9.0!, System.Drawing.FontStyle.Regular, System.Drawing.GraphicsUnit.Point, CType(0, Byte))
        Me.Label21.Location = New System.Drawing.Point(142, 552)
        Me.Label21.Name = "Label21"
        Me.Label21.Size = New System.Drawing.Size(62, 14)
        Me.Label21.TabIndex = 521
        Me.Label21.Text = "Nai (mM)"
        '
        'NKLmag
        '
        Me.NKLmag.Font = New System.Drawing.Font("Verdana", 9.75!, System.Drawing.FontStyle.Regular, System.Drawing.GraphicsUnit.Point, CType(0, Byte))
        Me.NKLmag.ForeColor = System.Drawing.Color.Black
        Me.NKLmag.Location = New System.Drawing.Point(206, 179)
        Me.NKLmag.Name = "NKLmag"
        Me.NKLmag.Size = New System.Drawing.Size(63, 23)
        Me.NKLmag.TabIndex = 654
        Me.NKLmag.Text = "0"
        '
        'NINamag
        '
        Me.NINamag.Font = New System.Drawing.Font("Verdana", 9.75!, System.Drawing.FontStyle.Regular, System.Drawing.GraphicsUnit.Point, CType(0, Byte))
        Me.NINamag.ForeColor = System.Drawing.Color.Red
        Me.NINamag.Location = New System.Drawing.Point(207, 380)
        Me.NINamag.Name = "NINamag"
        Me.NINamag.Size = New System.Drawing.Size(63, 23)
        Me.NINamag.TabIndex = 648
        Me.NINamag.Text = "0"
        '
        'NIbNSCmag
        '
        Me.NIbNSCmag.Font = New System.Drawing.Font("Verdana", 9.75!, System.Drawing.FontStyle.Regular, System.Drawing.GraphicsUnit.Point, CType(0, Byte))
        Me.NIbNSCmag.ForeColor = System.Drawing.Color.Black
        Me.NIbNSCmag.Location = New System.Drawing.Point(206, 254)
        Me.NIbNSCmag.Name = "NIbNSCmag"
        Me.NIbNSCmag.Size = New System.Drawing.Size(63, 23)
        Me.NIbNSCmag.TabIndex = 653
        Me.NIbNSCmag.Text = "0"
        '
        'NIKrmag
        '
        Me.NIKrmag.Font = New System.Drawing.Font("Verdana", 9.75!, System.Drawing.FontStyle.Regular, System.Drawing.GraphicsUnit.Point, CType(0, Byte))
        Me.NIKrmag.ForeColor = System.Drawing.Color.Red
        Me.NIKrmag.Location = New System.Drawing.Point(206, 107)
        Me.NIKrmag.Name = "NIKrmag"
        Me.NIKrmag.Size = New System.Drawing.Size(63, 23)
        Me.NIKrmag.TabIndex = 650
        Me.NIKrmag.Text = "0"
        '
        'NIK1mag
        '
        Me.NIK1mag.Font = New System.Drawing.Font("Verdana", 9.75!, System.Drawing.FontStyle.Regular, System.Drawing.GraphicsUnit.Point, CType(0, Byte))
        Me.NIK1mag.ForeColor = System.Drawing.Color.Red
        Me.NIK1mag.Location = New System.Drawing.Point(206, 128)
        Me.NIK1mag.Name = "NIK1mag"
        Me.NIK1mag.Size = New System.Drawing.Size(63, 23)
        Me.NIK1mag.TabIndex = 651
        Me.NIK1mag.Text = "0"
        '
        'NICaLmag
        '
        Me.NICaLmag.Font = New System.Drawing.Font("Verdana", 9.75!, System.Drawing.FontStyle.Regular, System.Drawing.GraphicsUnit.Point, CType(0, Byte))
        Me.NICaLmag.ForeColor = System.Drawing.Color.Red
        Me.NICaLmag.Location = New System.Drawing.Point(206, 153)
        Me.NICaLmag.Name = "NICaLmag"
        Me.NICaLmag.Size = New System.Drawing.Size(63, 23)
        Me.NICaLmag.TabIndex = 652
        Me.NICaLmag.Text = "0"
        '
        'Label55
        '
        Me.Label55.AutoSize = True
        Me.Label55.BackColor = System.Drawing.Color.White
        Me.Label55.Font = New System.Drawing.Font("Verdana", 24.0!, System.Drawing.FontStyle.Regular, System.Drawing.GraphicsUnit.Point, CType(0, Byte))
        Me.Label55.Location = New System.Drawing.Point(298, 592)
        Me.Label55.Name = "Label55"
        Me.Label55.Size = New System.Drawing.Size(39, 38)
        Me.Label55.TabIndex = 943
        Me.Label55.Text = "B"
        '
        'chkVmadaptive
        '
        Me.chkVmadaptive.AutoSize = True
        Me.chkVmadaptive.BackColor = System.Drawing.Color.White
        Me.chkVmadaptive.Checked = True
        Me.chkVmadaptive.CheckState = System.Windows.Forms.CheckState.Checked
        Me.chkVmadaptive.Font = New System.Drawing.Font("Verdana", 9.0!, System.Drawing.FontStyle.Regular, System.Drawing.GraphicsUnit.Point, CType(0, Byte))
        Me.chkVmadaptive.ForeColor = System.Drawing.Color.Black
        Me.chkVmadaptive.Location = New System.Drawing.Point(789, 42)
        Me.chkVmadaptive.Name = "chkVmadaptive"
        Me.chkVmadaptive.Size = New System.Drawing.Size(104, 18)
        Me.chkVmadaptive.TabIndex = 936
        Me.chkVmadaptive.Text = "Vm adaptive"
        Me.chkVmadaptive.UseVisualStyleBackColor = False
        '
        'chkVmreference
        '
        Me.chkVmreference.AutoSize = True
        Me.chkVmreference.BackColor = System.Drawing.Color.White
        Me.chkVmreference.Checked = True
        Me.chkVmreference.CheckState = System.Windows.Forms.CheckState.Checked
        Me.chkVmreference.Font = New System.Drawing.Font("Verdana", 9.0!, System.Drawing.FontStyle.Regular, System.Drawing.GraphicsUnit.Point, CType(0, Byte))
        Me.chkVmreference.ForeColor = System.Drawing.Color.Red
        Me.chkVmreference.Location = New System.Drawing.Point(674, 44)
        Me.chkVmreference.Name = "chkVmreference"
        Me.chkVmreference.Size = New System.Drawing.Size(109, 18)
        Me.chkVmreference.TabIndex = 935
        Me.chkVmreference.Text = "Vm reference"
        Me.chkVmreference.UseVisualStyleBackColor = False
        '
        'pbInwardI
        '
        Me.pbInwardI.BackColor = System.Drawing.SystemColors.ButtonHighlight
        Me.pbInwardI.Location = New System.Drawing.Point(296, 835)
        Me.pbInwardI.Name = "pbInwardI"
        Me.pbInwardI.Size = New System.Drawing.Size(600, 300)
        Me.pbInwardI.TabIndex = 942
        Me.pbInwardI.TabStop = False
        '
        'txtVmF
        '
        Me.txtVmF.Font = New System.Drawing.Font("Verdana", 12.0!, System.Drawing.FontStyle.Regular, System.Drawing.GraphicsUnit.Point, CType(0, Byte))
        Me.txtVmF.Location = New System.Drawing.Point(336, 36)
        Me.txtVmF.Name = "txtVmF"
        Me.txtVmF.Size = New System.Drawing.Size(100, 27)
        Me.txtVmF.TabIndex = 940
        Me.txtVmF.Text = "-"
        '
        'Label40
        '
        Me.Label40.AutoSize = True
        Me.Label40.BackColor = System.Drawing.Color.White
        Me.Label40.Font = New System.Drawing.Font("Verdana", 24.0!, System.Drawing.FontStyle.Regular, System.Drawing.GraphicsUnit.Point, CType(0, Byte))
        Me.Label40.Location = New System.Drawing.Point(298, 44)
        Me.Label40.Name = "Label40"
        Me.Label40.Size = New System.Drawing.Size(39, 38)
        Me.Label40.TabIndex = 939
        Me.Label40.Text = "A"
        '
        'Label38
        '
        Me.Label38.AutoSize = True
        Me.Label38.BackColor = System.Drawing.Color.White
        Me.Label38.Font = New System.Drawing.Font("Verdana", 12.0!, System.Drawing.FontStyle.Regular, System.Drawing.GraphicsUnit.Point, CType(0, Byte))
        Me.Label38.Location = New System.Drawing.Point(654, 576)
        Me.Label38.Name = "Label38"
        Me.Label38.Size = New System.Drawing.Size(50, 18)
        Me.Label38.TabIndex = 938
        Me.Label38.Text = "msec"
        '
        'pbpsVm
        '
        Me.pbpsVm.BackColor = System.Drawing.SystemColors.ButtonHighlight
        Me.pbpsVm.Location = New System.Drawing.Point(296, 30)
        Me.pbpsVm.Name = "pbpsVm"
        Me.pbpsVm.Size = New System.Drawing.Size(600, 544)
        Me.pbpsVm.TabIndex = 937
        Me.pbpsVm.TabStop = False
        '
        'pbOutwardI
        '
        Me.pbOutwardI.BackColor = System.Drawing.SystemColors.ButtonHighlight
        Me.pbOutwardI.Location = New System.Drawing.Point(296, 559)
        Me.pbOutwardI.Name = "pbOutwardI"
        Me.pbOutwardI.Size = New System.Drawing.Size(600, 300)
        Me.pbOutwardI.TabIndex = 941
        Me.pbOutwardI.TabStop = False
        '
        'Label31
        '
        Me.Label31.AutoSize = True
        Me.Label31.BackColor = System.Drawing.Color.White
        Me.Label31.Font = New System.Drawing.Font("Verdana", 11.25!, System.Drawing.FontStyle.Bold, System.Drawing.GraphicsUnit.Point, CType(0, Byte))
        Me.Label31.Location = New System.Drawing.Point(1225, 1105)
        Me.Label31.Name = "Label31"
        Me.Label31.Size = New System.Drawing.Size(41, 18)
        Me.Label31.TabIndex = 953
        Me.Label31.Text = "BP5"
        '
        'pbBP5
        '
        Me.pbBP5.BackColor = System.Drawing.SystemColors.ButtonHighlight
        Me.pbBP5.Location = New System.Drawing.Point(1126, 685)
        Me.pbBP5.Name = "pbBP5"
        Me.pbBP5.Size = New System.Drawing.Size(260, 450)
        Me.pbBP5.TabIndex = 952
        Me.pbBP5.TabStop = False
        '
        'Label68
        '
        Me.Label68.AutoSize = True
        Me.Label68.BackColor = System.Drawing.Color.White
        Me.Label68.Font = New System.Drawing.Font("Verdana", 11.25!, System.Drawing.FontStyle.Bold, System.Drawing.GraphicsUnit.Point, CType(0, Byte))
        Me.Label68.Location = New System.Drawing.Point(1021, 1104)
        Me.Label68.Name = "Label68"
        Me.Label68.Size = New System.Drawing.Size(41, 18)
        Me.Label68.TabIndex = 951
        Me.Label68.Text = "BP4"
        '
        'txtMeanBP4
        '
        Me.txtMeanBP4.Font = New System.Drawing.Font("Verdana", 12.0!, System.Drawing.FontStyle.Regular, System.Drawing.GraphicsUnit.Point, CType(0, Byte))
        Me.txtMeanBP4.Location = New System.Drawing.Point(1054, 649)
        Me.txtMeanBP4.Name = "txtMeanBP4"
        Me.txtMeanBP4.Size = New System.Drawing.Size(95, 27)
        Me.txtMeanBP4.TabIndex = 950
        '
        'Label67
        '
        Me.Label67.AutoSize = True
        Me.Label67.BackColor = System.Drawing.Color.White
        Me.Label67.Font = New System.Drawing.Font("Verdana", 11.25!, System.Drawing.FontStyle.Bold, System.Drawing.GraphicsUnit.Point, CType(0, Byte))
        Me.Label67.Location = New System.Drawing.Point(886, 153)
        Me.Label67.Name = "Label67"
        Me.Label67.Size = New System.Drawing.Size(94, 18)
        Me.Label67.TabIndex = 949
        Me.Label67.Text = "Log( MSE)"
        '
        'Label42
        '
        Me.Label42.AutoSize = True
        Me.Label42.BackColor = System.Drawing.Color.White
        Me.Label42.Font = New System.Drawing.Font("Verdana", 24.0!, System.Drawing.FontStyle.Regular, System.Drawing.GraphicsUnit.Point, CType(0, Byte))
        Me.Label42.Location = New System.Drawing.Point(874, 657)
        Me.Label42.Name = "Label42"
        Me.Label42.Size = New System.Drawing.Size(42, 38)
        Me.Label42.TabIndex = 945
        Me.Label42.Text = "G"
        '
        'Label54
        '
        Me.Label54.AutoSize = True
        Me.Label54.BackColor = System.Drawing.Color.White
        Me.Label54.Font = New System.Drawing.Font("Verdana", 24.0!, System.Drawing.FontStyle.Regular, System.Drawing.GraphicsUnit.Point, CType(0, Byte))
        Me.Label54.Location = New System.Drawing.Point(1145, 653)
        Me.Label54.Name = "Label54"
        Me.Label54.Size = New System.Drawing.Size(41, 38)
        Me.Label54.TabIndex = 944
        Me.Label54.Text = "H"
        '
        'pbBP4
        '
        Me.pbBP4.BackColor = System.Drawing.SystemColors.ButtonHighlight
        Me.pbBP4.Location = New System.Drawing.Point(889, 684)
        Me.pbBP4.Name = "pbBP4"
        Me.pbBP4.Size = New System.Drawing.Size(260, 450)
        Me.pbBP4.TabIndex = 947
        Me.pbBP4.TabStop = False
        '
        'TextBox1
        '
        Me.TextBox1.Font = New System.Drawing.Font("Verdana", 11.25!, System.Drawing.FontStyle.Regular, System.Drawing.GraphicsUnit.Point, CType(0, Byte))
        Me.TextBox1.Location = New System.Drawing.Point(15, 1126)
        Me.TextBox1.Name = "TextBox1"
        Me.TextBox1.Size = New System.Drawing.Size(2035, 26)
        Me.TextBox1.TabIndex = 954
        '
        'txtMeanBP2
        '
        Me.txtMeanBP2.Font = New System.Drawing.Font("Verdana", 12.0!, System.Drawing.FontStyle.Regular, System.Drawing.GraphicsUnit.Point, CType(0, Byte))
        Me.txtMeanBP2.Location = New System.Drawing.Point(1245, 164)
        Me.txtMeanBP2.Name = "txtMeanBP2"
        Me.txtMeanBP2.Size = New System.Drawing.Size(95, 27)
        Me.txtMeanBP2.TabIndex = 962
        '
        'txtMeanBP1
        '
        Me.txtMeanBP1.Font = New System.Drawing.Font("Verdana", 12.0!, System.Drawing.FontStyle.Regular, System.Drawing.GraphicsUnit.Point, CType(0, Byte))
        Me.txtMeanBP1.Location = New System.Drawing.Point(1054, 164)
        Me.txtMeanBP1.Name = "txtMeanBP1"
        Me.txtMeanBP1.Size = New System.Drawing.Size(95, 27)
        Me.txtMeanBP1.TabIndex = 961
        '
        'Label41
        '
        Me.Label41.AutoSize = True
        Me.Label41.BackColor = System.Drawing.Color.White
        Me.Label41.Font = New System.Drawing.Font("Verdana", 24.0!, System.Drawing.FontStyle.Regular, System.Drawing.GraphicsUnit.Point, CType(0, Byte))
        Me.Label41.Location = New System.Drawing.Point(1146, 172)
        Me.Label41.Name = "Label41"
        Me.Label41.Size = New System.Drawing.Size(37, 38)
        Me.Label41.TabIndex = 960
        Me.Label41.Text = "E"
        '
        'Label49
        '
        Me.Label49.AutoSize = True
        Me.Label49.BackColor = System.Drawing.Color.White
        Me.Label49.Font = New System.Drawing.Font("Verdana", 24.0!, System.Drawing.FontStyle.Regular, System.Drawing.GraphicsUnit.Point, CType(0, Byte))
        Me.Label49.Location = New System.Drawing.Point(883, 171)
        Me.Label49.Name = "Label49"
        Me.Label49.Size = New System.Drawing.Size(42, 38)
        Me.Label49.TabIndex = 959
        Me.Label49.Text = "D"
        '
        'Label12
        '
        Me.Label12.AutoSize = True
        Me.Label12.BackColor = System.Drawing.Color.White
        Me.Label12.Font = New System.Drawing.Font("Verdana", 11.25!, System.Drawing.FontStyle.Bold, System.Drawing.GraphicsUnit.Point, CType(0, Byte))
        Me.Label12.Location = New System.Drawing.Point(1245, 633)
        Me.Label12.Name = "Label12"
        Me.Label12.Size = New System.Drawing.Size(41, 18)
        Me.Label12.TabIndex = 958
        Me.Label12.Text = "BP2"
        '
        'Label61
        '
        Me.Label61.AutoSize = True
        Me.Label61.BackColor = System.Drawing.Color.White
        Me.Label61.Font = New System.Drawing.Font("Verdana", 11.25!, System.Drawing.FontStyle.Bold, System.Drawing.GraphicsUnit.Point, CType(0, Byte))
        Me.Label61.Location = New System.Drawing.Point(1019, 633)
        Me.Label61.Name = "Label61"
        Me.Label61.Size = New System.Drawing.Size(41, 18)
        Me.Label61.TabIndex = 957
        Me.Label61.Text = "BP1"
        '
        'pbBP1
        '
        Me.pbBP1.BackColor = System.Drawing.SystemColors.ButtonHighlight
        Me.pbBP1.Location = New System.Drawing.Point(889, 206)
        Me.pbBP1.Name = "pbBP1"
        Me.pbBP1.Size = New System.Drawing.Size(260, 450)
        Me.pbBP1.TabIndex = 955
        Me.pbBP1.TabStop = False
        '
        'pbPB2
        '
        Me.pbPB2.BackColor = System.Drawing.SystemColors.ButtonHighlight
        Me.pbPB2.Location = New System.Drawing.Point(1126, 206)
        Me.pbPB2.Name = "pbPB2"
        Me.pbPB2.Size = New System.Drawing.Size(260, 450)
        Me.pbPB2.TabIndex = 956
        Me.pbPB2.TabStop = False
        '
        'Label66
        '
        Me.Label66.AutoSize = True
        Me.Label66.BackColor = System.Drawing.Color.White
        Me.Label66.Font = New System.Drawing.Font("Verdana", 24.0!, System.Drawing.FontStyle.Regular, System.Drawing.GraphicsUnit.Point, CType(0, Byte))
        Me.Label66.Location = New System.Drawing.Point(1391, 171)
        Me.Label66.Name = "Label66"
        Me.Label66.Size = New System.Drawing.Size(35, 38)
        Me.Label66.TabIndex = 981
        Me.Label66.Text = "F"
        '
        'txtMeanBP5
        '
        Me.txtMeanBP5.Font = New System.Drawing.Font("Verdana", 12.0!, System.Drawing.FontStyle.Regular, System.Drawing.GraphicsUnit.Point, CType(0, Byte))
        Me.txtMeanBP5.Location = New System.Drawing.Point(1250, 649)
        Me.txtMeanBP5.Name = "txtMeanBP5"
        Me.txtMeanBP5.Size = New System.Drawing.Size(95, 27)
        Me.txtMeanBP5.TabIndex = 980
        '
        'Label45
        '
        Me.Label45.AutoSize = True
        Me.Label45.BackColor = System.Drawing.Color.White
        Me.Label45.Font = New System.Drawing.Font("Verdana", 24.0!, System.Drawing.FontStyle.Regular, System.Drawing.GraphicsUnit.Point, CType(0, Byte))
        Me.Label45.Location = New System.Drawing.Point(1391, 654)
        Me.Label45.Name = "Label45"
        Me.Label45.Size = New System.Drawing.Size(30, 38)
        Me.Label45.TabIndex = 978
        Me.Label45.Text = "I"
        '
        'Label37
        '
        Me.Label37.AutoSize = True
        Me.Label37.BackColor = System.Drawing.Color.White
        Me.Label37.Font = New System.Drawing.Font("Verdana", 9.75!, System.Drawing.FontStyle.Regular, System.Drawing.GraphicsUnit.Point, CType(0, Byte))
        Me.Label37.Location = New System.Drawing.Point(1772, 232)
        Me.Label37.Name = "Label37"
        Me.Label37.Size = New System.Drawing.Size(45, 16)
        Me.Label37.TabIndex = 977
        Me.Label37.Text = "Stp ="
        '
        'txtStp
        '
        Me.txtStp.Font = New System.Drawing.Font("Verdana", 9.75!, System.Drawing.FontStyle.Regular, System.Drawing.GraphicsUnit.Point, CType(0, Byte))
        Me.txtStp.Location = New System.Drawing.Point(1823, 229)
        Me.txtStp.Name = "txtStp"
        Me.txtStp.Size = New System.Drawing.Size(68, 23)
        Me.txtStp.TabIndex = 976
        '
        'Label69
        '
        Me.Label69.AutoSize = True
        Me.Label69.BackColor = System.Drawing.Color.White
        Me.Label69.Font = New System.Drawing.Font("Verdana", 9.75!, System.Drawing.FontStyle.Regular, System.Drawing.GraphicsUnit.Point, CType(0, Byte))
        Me.Label69.Location = New System.Drawing.Point(1747, 144)
        Me.Label69.Name = "Label69"
        Me.Label69.Size = New System.Drawing.Size(92, 16)
        Me.Label69.TabIndex = 975
        Me.Label69.Text = "removed N ="
        '
        'txtdetectINa
        '
        Me.txtdetectINa.Font = New System.Drawing.Font("Verdana", 9.75!, System.Drawing.FontStyle.Regular, System.Drawing.GraphicsUnit.Point, CType(0, Byte))
        Me.txtdetectINa.Location = New System.Drawing.Point(1845, 141)
        Me.txtdetectINa.Name = "txtdetectINa"
        Me.txtdetectINa.Size = New System.Drawing.Size(42, 23)
        Me.txtdetectINa.TabIndex = 974
        '
        'Label65
        '
        Me.Label65.AutoSize = True
        Me.Label65.Font = New System.Drawing.Font("Verdana", 9.75!, System.Drawing.FontStyle.Regular, System.Drawing.GraphicsUnit.Point, CType(0, Byte))
        Me.Label65.Location = New System.Drawing.Point(1768, 254)
        Me.Label65.Name = "Label65"
        Me.Label65.Size = New System.Drawing.Size(77, 16)
        Me.Label65.TabIndex = 972
        Me.Label65.Text = "N_PScycle"
        '
        'txtPScycle
        '
        Me.txtPScycle.Font = New System.Drawing.Font("Verdana", 9.75!, System.Drawing.FontStyle.Regular, System.Drawing.GraphicsUnit.Point, CType(0, Byte))
        Me.txtPScycle.Location = New System.Drawing.Point(1842, 250)
        Me.txtPScycle.Name = "txtPScycle"
        Me.txtPScycle.Size = New System.Drawing.Size(49, 23)
        Me.txtPScycle.TabIndex = 971
        '
        'Label32
        '
        Me.Label32.AutoSize = True
        Me.Label32.Font = New System.Drawing.Font("Verdana", 9.75!, System.Drawing.FontStyle.Regular, System.Drawing.GraphicsUnit.Point, CType(0, Byte))
        Me.Label32.Location = New System.Drawing.Point(1752, 208)
        Me.Label32.Name = "Label32"
        Me.Label32.Size = New System.Drawing.Size(69, 16)
        Me.Label32.TabIndex = 970
        Me.Label32.Text = "N_BPSQS"
        '
        'txtNBPSQS
        '
        Me.txtNBPSQS.Font = New System.Drawing.Font("Verdana", 9.75!, System.Drawing.FontStyle.Regular, System.Drawing.GraphicsUnit.Point, CType(0, Byte))
        Me.txtNBPSQS.Location = New System.Drawing.Point(1823, 205)
        Me.txtNBPSQS.Name = "txtNBPSQS"
        Me.txtNBPSQS.Size = New System.Drawing.Size(49, 23)
        Me.txtNBPSQS.TabIndex = 969
        '
        'txtLogMSE
        '
        Me.txtLogMSE.Font = New System.Drawing.Font("Verdana", 9.75!, System.Drawing.FontStyle.Regular, System.Drawing.GraphicsUnit.Point, CType(0, Byte))
        Me.txtLogMSE.Location = New System.Drawing.Point(1804, 164)
        Me.txtLogMSE.Name = "txtLogMSE"
        Me.txtLogMSE.Size = New System.Drawing.Size(83, 23)
        Me.txtLogMSE.TabIndex = 968
        Me.txtLogMSE.Text = "1"
        '
        'pbBP6
        '
        Me.pbBP6.BackColor = System.Drawing.SystemColors.ButtonHighlight
        Me.pbBP6.Location = New System.Drawing.Point(1365, 685)
        Me.pbBP6.Name = "pbBP6"
        Me.pbBP6.Size = New System.Drawing.Size(260, 450)
        Me.pbBP6.TabIndex = 967
        Me.pbBP6.TabStop = False
        '
        'Label58
        '
        Me.Label58.AutoSize = True
        Me.Label58.Font = New System.Drawing.Font("Verdana", 9.75!, System.Drawing.FontStyle.Regular, System.Drawing.GraphicsUnit.Point, CType(0, Byte))
        Me.Label58.Location = New System.Drawing.Point(1643, 171)
        Me.Label58.Name = "Label58"
        Me.Label58.Size = New System.Drawing.Size(50, 16)
        Me.Label58.TabIndex = 966
        Me.Label58.Text = "MSE ="
        '
        'txtMSE
        '
        Me.txtMSE.Font = New System.Drawing.Font("Verdana", 9.75!, System.Drawing.FontStyle.Regular, System.Drawing.GraphicsUnit.Point, CType(0, Byte))
        Me.txtMSE.Location = New System.Drawing.Point(1719, 164)
        Me.txtMSE.Name = "txtMSE"
        Me.txtMSE.Size = New System.Drawing.Size(83, 23)
        Me.txtMSE.TabIndex = 965
        Me.txtMSE.Text = "1"
        '
        'pbSQS
        '
        Me.pbSQS.BackColor = System.Drawing.SystemColors.ButtonHighlight
        Me.pbSQS.Location = New System.Drawing.Point(1625, 205)
        Me.pbSQS.Name = "pbSQS"
        Me.pbSQS.Size = New System.Drawing.Size(267, 617)
        Me.pbSQS.TabIndex = 973
        Me.pbSQS.TabStop = False
        '
        'Label46
        '
        Me.Label46.AutoSize = True
        Me.Label46.BackColor = System.Drawing.Color.White
        Me.Label46.Font = New System.Drawing.Font("Verdana", 14.25!, System.Drawing.FontStyle.Regular, System.Drawing.GraphicsUnit.Point, CType(0, Byte))
        Me.Label46.Location = New System.Drawing.Point(1520, 626)
        Me.Label46.Name = "Label46"
        Me.Label46.Size = New System.Drawing.Size(105, 23)
        Me.Label46.TabIndex = 982
        Me.Label46.Text = "time (ms)"
        '
        'Label62
        '
        Me.Label62.AutoSize = True
        Me.Label62.BackColor = System.Drawing.Color.White
        Me.Label62.Font = New System.Drawing.Font("Verdana", 11.25!, System.Drawing.FontStyle.Bold, System.Drawing.GraphicsUnit.Point, CType(0, Byte))
        Me.Label62.Location = New System.Drawing.Point(1507, 1105)
        Me.Label62.Name = "Label62"
        Me.Label62.Size = New System.Drawing.Size(41, 18)
        Me.Label62.TabIndex = 983
        Me.Label62.Text = "BP6"
        '
        'txtCellNo
        '
        Me.txtCellNo.Font = New System.Drawing.Font("Verdana", 15.75!, System.Drawing.FontStyle.Regular, System.Drawing.GraphicsUnit.Point, CType(0, Byte))
        Me.txtCellNo.Location = New System.Drawing.Point(125, 79)
        Me.txtCellNo.Name = "txtCellNo"
        Me.txtCellNo.Size = New System.Drawing.Size(167, 33)
        Me.txtCellNo.TabIndex = 984
        '
        'txtMeanBP6
        '
        Me.txtMeanBP6.Font = New System.Drawing.Font("Verdana", 12.0!, System.Drawing.FontStyle.Regular, System.Drawing.GraphicsUnit.Point, CType(0, Byte))
        Me.txtMeanBP6.Location = New System.Drawing.Point(1474, 649)
        Me.txtMeanBP6.Name = "txtMeanBP6"
        Me.txtMeanBP6.Size = New System.Drawing.Size(95, 27)
        Me.txtMeanBP6.TabIndex = 987
        '
        'Label74
        '
        Me.Label74.AutoSize = True
        Me.Label74.BackColor = System.Drawing.Color.White
        Me.Label74.Font = New System.Drawing.Font("Verdana", 24.0!, System.Drawing.FontStyle.Regular, System.Drawing.GraphicsUnit.Point, CType(0, Byte))
        Me.Label74.Location = New System.Drawing.Point(1598, 200)
        Me.Label74.Name = "Label74"
        Me.Label74.Size = New System.Drawing.Size(37, 38)
        Me.Label74.TabIndex = 986
        Me.Label74.Text = "E"
        '
        'pbBP3
        '
        Me.pbBP3.BackColor = System.Drawing.SystemColors.ButtonHighlight
        Me.pbBP3.Location = New System.Drawing.Point(1365, 206)
        Me.pbBP3.Name = "pbBP3"
        Me.pbBP3.Size = New System.Drawing.Size(260, 450)
        Me.pbBP3.TabIndex = 985
        Me.pbBP3.TabStop = False
        '
        'Label75
        '
        Me.Label75.AutoSize = True
        Me.Label75.BackColor = System.Drawing.Color.White
        Me.Label75.Font = New System.Drawing.Font("Verdana", 9.75!, System.Drawing.FontStyle.Regular, System.Drawing.GraphicsUnit.Point, CType(0, Byte))
        Me.Label75.Location = New System.Drawing.Point(1797, 274)
        Me.Label75.Name = "Label75"
        Me.Label75.Size = New System.Drawing.Size(52, 16)
        Me.Label75.TabIndex = 990
        Me.Label75.Text = " CR2 ="
        '
        'txtR2
        '
        Me.txtR2.Font = New System.Drawing.Font("Verdana", 9.75!, System.Drawing.FontStyle.Regular, System.Drawing.GraphicsUnit.Point, CType(0, Byte))
        Me.txtR2.Location = New System.Drawing.Point(1849, 272)
        Me.txtR2.Name = "txtR2"
        Me.txtR2.Size = New System.Drawing.Size(42, 23)
        Me.txtR2.TabIndex = 989
        '
        'Label76
        '
        Me.Label76.AutoSize = True
        Me.Label76.BackColor = System.Drawing.Color.White
        Me.Label76.Font = New System.Drawing.Font("Verdana", 11.25!, System.Drawing.FontStyle.Bold, System.Drawing.GraphicsUnit.Point, CType(0, Byte))
        Me.Label76.Location = New System.Drawing.Point(1479, 632)
        Me.Label76.Name = "Label76"
        Me.Label76.Size = New System.Drawing.Size(41, 18)
        Me.Label76.TabIndex = 991
        Me.Label76.Text = "BP3"
        '
        'txtMeanBP3
        '
        Me.txtMeanBP3.Font = New System.Drawing.Font("Verdana", 12.0!, System.Drawing.FontStyle.Regular, System.Drawing.GraphicsUnit.Point, CType(0, Byte))
        Me.txtMeanBP3.Location = New System.Drawing.Point(1493, 166)
        Me.txtMeanBP3.Name = "txtMeanBP3"
        Me.txtMeanBP3.Size = New System.Drawing.Size(95, 27)
        Me.txtMeanBP3.TabIndex = 998
        '
        'txtI1
        '
        Me.txtI1.Font = New System.Drawing.Font("Verdana", 12.0!, System.Drawing.FontStyle.Regular, System.Drawing.GraphicsUnit.Point, CType(0, Byte))
        Me.txtI1.Location = New System.Drawing.Point(997, 174)
        Me.txtI1.Name = "txtI1"
        Me.txtI1.Size = New System.Drawing.Size(55, 27)
        Me.txtI1.TabIndex = 999
        '
        'txtI2
        '
        Me.txtI2.Font = New System.Drawing.Font("Verdana", 12.0!, System.Drawing.FontStyle.Regular, System.Drawing.GraphicsUnit.Point, CType(0, Byte))
        Me.txtI2.Location = New System.Drawing.Point(1189, 175)
        Me.txtI2.Name = "txtI2"
        Me.txtI2.Size = New System.Drawing.Size(55, 27)
        Me.txtI2.TabIndex = 1000
        '
        'txtI3
        '
        Me.txtI3.Font = New System.Drawing.Font("Verdana", 12.0!, System.Drawing.FontStyle.Regular, System.Drawing.GraphicsUnit.Point, CType(0, Byte))
        Me.txtI3.Location = New System.Drawing.Point(1432, 178)
        Me.txtI3.Name = "txtI3"
        Me.txtI3.Size = New System.Drawing.Size(55, 27)
        Me.txtI3.TabIndex = 1001
        '
        'txtI4
        '
        Me.txtI4.Font = New System.Drawing.Font("Verdana", 12.0!, System.Drawing.FontStyle.Regular, System.Drawing.GraphicsUnit.Point, CType(0, Byte))
        Me.txtI4.Location = New System.Drawing.Point(995, 657)
        Me.txtI4.Name = "txtI4"
        Me.txtI4.Size = New System.Drawing.Size(55, 27)
        Me.txtI4.TabIndex = 1002
        '
        'txtI5
        '
        Me.txtI5.Font = New System.Drawing.Font("Verdana", 12.0!, System.Drawing.FontStyle.Regular, System.Drawing.GraphicsUnit.Point, CType(0, Byte))
        Me.txtI5.Location = New System.Drawing.Point(1193, 657)
        Me.txtI5.Name = "txtI5"
        Me.txtI5.Size = New System.Drawing.Size(55, 27)
        Me.txtI5.TabIndex = 1003
        '
        'txtI6
        '
        Me.txtI6.Font = New System.Drawing.Font("Verdana", 12.0!, System.Drawing.FontStyle.Regular, System.Drawing.GraphicsUnit.Point, CType(0, Byte))
        Me.txtI6.Location = New System.Drawing.Point(1418, 657)
        Me.txtI6.Name = "txtI6"
        Me.txtI6.Size = New System.Drawing.Size(55, 27)
        Me.txtI6.TabIndex = 1004
        '
        'txtVmOnset
        '
        Me.txtVmOnset.Font = New System.Drawing.Font("Verdana", 12.0!, System.Drawing.FontStyle.Regular, System.Drawing.GraphicsUnit.Point, CType(0, Byte))
        Me.txtVmOnset.Location = New System.Drawing.Point(446, 37)
        Me.txtVmOnset.Name = "txtVmOnset"
        Me.txtVmOnset.Size = New System.Drawing.Size(102, 27)
        Me.txtVmOnset.TabIndex = 1005
        Me.txtVmOnset.Text = "-"
        '
        'txtError
        '
        Me.txtError.Font = New System.Drawing.Font("Verdana", 12.0!, System.Drawing.FontStyle.Regular, System.Drawing.GraphicsUnit.Point, CType(0, Byte))
        Me.txtError.Location = New System.Drawing.Point(563, 36)
        Me.txtError.Name = "txtError"
        Me.txtError.Size = New System.Drawing.Size(106, 27)
        Me.txtError.TabIndex = 1006
        Me.txtError.Text = "-"
        '
        'Label39
        '
        Me.Label39.AutoSize = True
        Me.Label39.BackColor = System.Drawing.Color.White
        Me.Label39.Font = New System.Drawing.Font("Verdana", 9.75!, System.Drawing.FontStyle.Regular, System.Drawing.GraphicsUnit.Point, CType(0, Byte))
        Me.Label39.Location = New System.Drawing.Point(548, 41)
        Me.Label39.Name = "Label39"
        Me.Label39.Size = New System.Drawing.Size(17, 16)
        Me.Label39.TabIndex = 1007
        Me.Label39.Text = "="
        '
        'Label57
        '
        Me.Label57.AutoSize = True
        Me.Label57.BackColor = System.Drawing.Color.White
        Me.Label57.Font = New System.Drawing.Font("Verdana", 9.75!, System.Drawing.FontStyle.Regular, System.Drawing.GraphicsUnit.Point, CType(0, Byte))
        Me.Label57.Location = New System.Drawing.Point(434, 42)
        Me.Label57.Name = "Label57"
        Me.Label57.Size = New System.Drawing.Size(15, 16)
        Me.Label57.TabIndex = 1008
        Me.Label57.Text = "-"
        '
        'txtSbp1
        '
        Me.txtSbp1.Font = New System.Drawing.Font("Verdana", 8.0!)
        Me.txtSbp1.Location = New System.Drawing.Point(1054, 192)
        Me.txtSbp1.Name = "txtSbp1"
        Me.txtSbp1.Size = New System.Drawing.Size(95, 20)
        Me.txtSbp1.TabIndex = 1009
        '
        'txtSbp2
        '
        Me.txtSbp2.Font = New System.Drawing.Font("Verdana", 8.0!)
        Me.txtSbp2.Location = New System.Drawing.Point(1245, 189)
        Me.txtSbp2.Name = "txtSbp2"
        Me.txtSbp2.Size = New System.Drawing.Size(95, 20)
        Me.txtSbp2.TabIndex = 1010
        '
        'txtSbp3
        '
        Me.txtSbp3.Font = New System.Drawing.Font("Verdana", 8.0!)
        Me.txtSbp3.Location = New System.Drawing.Point(1493, 194)
        Me.txtSbp3.Name = "txtSbp3"
        Me.txtSbp3.Size = New System.Drawing.Size(95, 20)
        Me.txtSbp3.TabIndex = 1011
        '
        'txtSbp4
        '
        Me.txtSbp4.Font = New System.Drawing.Font("Verdana", 8.0!)
        Me.txtSbp4.Location = New System.Drawing.Point(1054, 676)
        Me.txtSbp4.Name = "txtSbp4"
        Me.txtSbp4.Size = New System.Drawing.Size(95, 20)
        Me.txtSbp4.TabIndex = 1012
        '
        'txtSbp5
        '
        Me.txtSbp5.Font = New System.Drawing.Font("Verdana", 8.0!)
        Me.txtSbp5.Location = New System.Drawing.Point(1250, 676)
        Me.txtSbp5.Name = "txtSbp5"
        Me.txtSbp5.Size = New System.Drawing.Size(95, 20)
        Me.txtSbp5.TabIndex = 1013
        '
        'txtSbp6
        '
        Me.txtSbp6.Font = New System.Drawing.Font("Verdana", 8.0!)
        Me.txtSbp6.Location = New System.Drawing.Point(1474, 676)
        Me.txtSbp6.Name = "txtSbp6"
        Me.txtSbp6.Size = New System.Drawing.Size(95, 20)
        Me.txtSbp6.TabIndex = 1014
        '
        'pbSensitivity
        '
        Me.pbSensitivity.BackColor = System.Drawing.SystemColors.ButtonHighlight
        Me.pbSensitivity.Location = New System.Drawing.Point(1625, 824)
        Me.pbSensitivity.Name = "pbSensitivity"
        Me.pbSensitivity.Size = New System.Drawing.Size(267, 301)
        Me.pbSensitivity.TabIndex = 1015
        Me.pbSensitivity.TabStop = False
        '
        'Label63
        '
        Me.Label63.AutoSize = True
        Me.Label63.BackColor = System.Drawing.Color.White
        Me.Label63.Font = New System.Drawing.Font("Verdana", 11.25!, System.Drawing.FontStyle.Bold, System.Drawing.GraphicsUnit.Point, CType(0, Byte))
        Me.Label63.ForeColor = System.Drawing.Color.Gray
        Me.Label63.Location = New System.Drawing.Point(1677, 804)
        Me.Label63.Name = "Label63"
        Me.Label63.Size = New System.Drawing.Size(41, 18)
        Me.Label63.TabIndex = 1016
        Me.Label63.Text = "BP2"
        '
        'Label73
        '
        Me.Label73.AutoSize = True
        Me.Label73.BackColor = System.Drawing.Color.White
        Me.Label73.Font = New System.Drawing.Font("Verdana", 11.25!, System.Drawing.FontStyle.Bold, System.Drawing.GraphicsUnit.Point, CType(0, Byte))
        Me.Label73.Location = New System.Drawing.Point(1635, 804)
        Me.Label73.Name = "Label73"
        Me.Label73.Size = New System.Drawing.Size(41, 18)
        Me.Label73.TabIndex = 1017
        Me.Label73.Text = "BP1"
        '
        'Label81
        '
        Me.Label81.AutoSize = True
        Me.Label81.BackColor = System.Drawing.Color.White
        Me.Label81.Font = New System.Drawing.Font("Verdana", 11.25!, System.Drawing.FontStyle.Bold, System.Drawing.GraphicsUnit.Point, CType(0, Byte))
        Me.Label81.ForeColor = System.Drawing.Color.Fuchsia
        Me.Label81.Location = New System.Drawing.Point(1718, 804)
        Me.Label81.Name = "Label81"
        Me.Label81.Size = New System.Drawing.Size(41, 18)
        Me.Label81.TabIndex = 1018
        Me.Label81.Text = "BP3"
        '
        'Label82
        '
        Me.Label82.AutoSize = True
        Me.Label82.BackColor = System.Drawing.Color.White
        Me.Label82.Font = New System.Drawing.Font("Verdana", 11.25!, System.Drawing.FontStyle.Bold, System.Drawing.GraphicsUnit.Point, CType(0, Byte))
        Me.Label82.ForeColor = System.Drawing.Color.Lime
        Me.Label82.Location = New System.Drawing.Point(1759, 805)
        Me.Label82.Name = "Label82"
        Me.Label82.Size = New System.Drawing.Size(41, 18)
        Me.Label82.TabIndex = 1019
        Me.Label82.Text = "BP4"
        '
        'Label83
        '
        Me.Label83.AutoSize = True
        Me.Label83.BackColor = System.Drawing.Color.White
        Me.Label83.Font = New System.Drawing.Font("Verdana", 11.25!, System.Drawing.FontStyle.Bold, System.Drawing.GraphicsUnit.Point, CType(0, Byte))
        Me.Label83.ForeColor = System.Drawing.Color.Red
        Me.Label83.Location = New System.Drawing.Point(1803, 804)
        Me.Label83.Name = "Label83"
        Me.Label83.Size = New System.Drawing.Size(41, 18)
        Me.Label83.TabIndex = 1020
        Me.Label83.Text = "BP5"
        '
        'Label84
        '
        Me.Label84.AutoSize = True
        Me.Label84.BackColor = System.Drawing.Color.White
        Me.Label84.Font = New System.Drawing.Font("Verdana", 11.25!, System.Drawing.FontStyle.Bold, System.Drawing.GraphicsUnit.Point, CType(0, Byte))
        Me.Label84.ForeColor = System.Drawing.Color.Blue
        Me.Label84.Location = New System.Drawing.Point(1848, 804)
        Me.Label84.Name = "Label84"
        Me.Label84.Size = New System.Drawing.Size(41, 18)
        Me.Label84.TabIndex = 1021
        Me.Label84.Text = "BP6"
        '
        'Label85
        '
        Me.Label85.AutoSize = True
        Me.Label85.BackColor = System.Drawing.Color.White
        Me.Label85.Font = New System.Drawing.Font("Verdana", 9.75!, System.Drawing.FontStyle.Regular, System.Drawing.GraphicsUnit.Point, CType(0, Byte))
        Me.Label85.Location = New System.Drawing.Point(1807, 296)
        Me.Label85.Name = "Label85"
        Me.Label85.Size = New System.Drawing.Size(38, 16)
        Me.Label85.TabIndex = 1023
        Me.Label85.Text = " C35"
        '
        'txtCLB
        '
        Me.txtCLB.Font = New System.Drawing.Font("Verdana", 9.75!, System.Drawing.FontStyle.Regular, System.Drawing.GraphicsUnit.Point, CType(0, Byte))
        Me.txtCLB.Location = New System.Drawing.Point(1849, 293)
        Me.txtCLB.Name = "txtCLB"
        Me.txtCLB.Size = New System.Drawing.Size(42, 23)
        Me.txtCLB.TabIndex = 1022
        '
        'txtPreviousMIN
        '
        Me.txtPreviousMIN.Font = New System.Drawing.Font("Verdana", 9.75!, System.Drawing.FontStyle.Regular, System.Drawing.GraphicsUnit.Point, CType(0, Byte))
        Me.txtPreviousMIN.Location = New System.Drawing.Point(1719, 184)
        Me.txtPreviousMIN.Name = "txtPreviousMIN"
        Me.txtPreviousMIN.Size = New System.Drawing.Size(83, 23)
        Me.txtPreviousMIN.TabIndex = 1024
        Me.txtPreviousMIN.Text = "1"
        '
        'Label86
        '
        Me.Label86.AutoSize = True
        Me.Label86.Font = New System.Drawing.Font("Verdana", 9.75!, System.Drawing.FontStyle.Regular, System.Drawing.GraphicsUnit.Point, CType(0, Byte))
        Me.Label86.Location = New System.Drawing.Point(1643, 189)
        Me.Label86.Name = "Label86"
        Me.Label86.Size = New System.Drawing.Size(77, 16)
        Me.Label86.TabIndex = 1025
        Me.Label86.Text = "Previous ="
        '
        'rbEMTest
        '
        Me.rbEMTest.AutoSize = True
        Me.rbEMTest.Font = New System.Drawing.Font("Verdana", 12.0!, System.Drawing.FontStyle.Regular, System.Drawing.GraphicsUnit.Point, CType(0, Byte))
        Me.rbEMTest.Location = New System.Drawing.Point(40, 54)
        Me.rbEMTest.Name = "rbEMTest"
        Me.rbEMTest.Size = New System.Drawing.Size(98, 22)
        Me.rbEMTest.TabIndex = 1026
        Me.rbEMTest.TabStop = True
        Me.rbEMTest.Text = "rbEMtest"
        Me.rbEMTest.UseVisualStyleBackColor = True
        '
        'rbMMtest
        '
        Me.rbMMtest.AutoSize = True
        Me.rbMMtest.Checked = True
        Me.rbMMtest.Font = New System.Drawing.Font("Verdana", 12.0!, System.Drawing.FontStyle.Regular, System.Drawing.GraphicsUnit.Point, CType(0, Byte))
        Me.rbMMtest.ForeColor = System.Drawing.Color.Red
        Me.rbMMtest.Location = New System.Drawing.Point(157, 54)
        Me.rbMMtest.Name = "rbMMtest"
        Me.rbMMtest.Size = New System.Drawing.Size(101, 22)
        Me.rbMMtest.TabIndex = 1027
        Me.rbMMtest.TabStop = True
        Me.rbMMtest.Text = "rbMMtest"
        Me.rbMMtest.UseVisualStyleBackColor = True
        '
        'Protocol1
        '
        Me.AutoScaleDimensions = New System.Drawing.SizeF(6.0!, 12.0!)
        Me.AutoScaleMode = System.Windows.Forms.AutoScaleMode.Font
        Me.ClientSize = New System.Drawing.Size(1924, 1181)
        Me.Controls.Add(Me.rbMMtest)
        Me.Controls.Add(Me.rbEMTest)
        Me.Controls.Add(Me.Label86)
        Me.Controls.Add(Me.txtPreviousMIN)
        Me.Controls.Add(Me.Label85)
        Me.Controls.Add(Me.txtCLB)
        Me.Controls.Add(Me.Label84)
        Me.Controls.Add(Me.Label83)
        Me.Controls.Add(Me.Label82)
        Me.Controls.Add(Me.Label81)
        Me.Controls.Add(Me.Label73)
        Me.Controls.Add(Me.Label63)
        Me.Controls.Add(Me.pbSensitivity)
        Me.Controls.Add(Me.txtSbp6)
        Me.Controls.Add(Me.txtSbp5)
        Me.Controls.Add(Me.txtSbp4)
        Me.Controls.Add(Me.txtSbp3)
        Me.Controls.Add(Me.txtSbp2)
        Me.Controls.Add(Me.txtSbp1)
        Me.Controls.Add(Me.Label57)
        Me.Controls.Add(Me.Label39)
        Me.Controls.Add(Me.txtMeanBP4)
        Me.Controls.Add(Me.txtError)
        Me.Controls.Add(Me.txtVmOnset)
        Me.Controls.Add(Me.txtI6)
        Me.Controls.Add(Me.txtI5)
        Me.Controls.Add(Me.txtI4)
        Me.Controls.Add(Me.txtI3)
        Me.Controls.Add(Me.txtI2)
        Me.Controls.Add(Me.txtI1)
        Me.Controls.Add(Me.txtMeanBP3)
        Me.Controls.Add(Me.TextBox1)
        Me.Controls.Add(Me.Label76)
        Me.Controls.Add(Me.Label46)
        Me.Controls.Add(Me.Label75)
        Me.Controls.Add(Me.txtR2)
        Me.Controls.Add(Me.txtMeanBP6)
        Me.Controls.Add(Me.Label74)
        Me.Controls.Add(Me.txtCellNo)
        Me.Controls.Add(Me.Label62)
        Me.Controls.Add(Me.Label66)
        Me.Controls.Add(Me.txtMeanBP5)
        Me.Controls.Add(Me.Label45)
        Me.Controls.Add(Me.Label37)
        Me.Controls.Add(Me.txtStp)
        Me.Controls.Add(Me.Label69)
        Me.Controls.Add(Me.txtdetectINa)
        Me.Controls.Add(Me.Label65)
        Me.Controls.Add(Me.txtPScycle)
        Me.Controls.Add(Me.Label32)
        Me.Controls.Add(Me.txtNBPSQS)
        Me.Controls.Add(Me.txtLogMSE)
        Me.Controls.Add(Me.Label58)
        Me.Controls.Add(Me.txtMSE)
        Me.Controls.Add(Me.pbSQS)
        Me.Controls.Add(Me.txtMeanBP2)
        Me.Controls.Add(Me.txtMeanBP1)
        Me.Controls.Add(Me.Label41)
        Me.Controls.Add(Me.Label49)
        Me.Controls.Add(Me.Label12)
        Me.Controls.Add(Me.Label61)
        Me.Controls.Add(Me.pbBP1)
        Me.Controls.Add(Me.pbPB2)
        Me.Controls.Add(Me.Label31)
        Me.Controls.Add(Me.Label68)
        Me.Controls.Add(Me.Label67)
        Me.Controls.Add(Me.Label42)
        Me.Controls.Add(Me.Label54)
        Me.Controls.Add(Me.pbBP4)
        Me.Controls.Add(Me.Label55)
        Me.Controls.Add(Me.chkVmadaptive)
        Me.Controls.Add(Me.chkVmreference)
        Me.Controls.Add(Me.pbInwardI)
        Me.Controls.Add(Me.txtVmF)
        Me.Controls.Add(Me.Label40)
        Me.Controls.Add(Me.Label38)
        Me.Controls.Add(Me.GroupBox3)
        Me.Controls.Add(Me.controlBox4)
        Me.Controls.Add(Me.Label4)
        Me.Controls.Add(Me.cbEnd)
        Me.Controls.Add(Me.cbSelect)
        Me.Controls.Add(Me.cbPlot)
        Me.Controls.Add(Me.cbCsvFileList)
        Me.Controls.Add(Me.cbselectCSVfile)
        Me.Controls.Add(Me.cbPS)
        Me.Controls.Add(Me.cbManualFit)
        Me.Controls.Add(Me.pbpsVm)
        Me.Controls.Add(Me.pbOutwardI)
        Me.Controls.Add(Me.pbVm1)
        Me.Controls.Add(Me.pbBP3)
        Me.Controls.Add(Me.pbBP5)
        Me.Controls.Add(Me.pbBP6)
        Me.Name = "Protocol1"
        Me.Text = "-5"
        CType(Me.pbVm1, System.ComponentModel.ISupportInitialize).EndInit()
        Me.GroupBox3.ResumeLayout(False)
        Me.GroupBox3.PerformLayout()
        Me.controlBox4.ResumeLayout(False)
        Me.controlBox4.PerformLayout()
        CType(Me.pbInwardI, System.ComponentModel.ISupportInitialize).EndInit()
        CType(Me.pbpsVm, System.ComponentModel.ISupportInitialize).EndInit()
        CType(Me.pbOutwardI, System.ComponentModel.ISupportInitialize).EndInit()
        CType(Me.pbBP5, System.ComponentModel.ISupportInitialize).EndInit()
        CType(Me.pbBP4, System.ComponentModel.ISupportInitialize).EndInit()
        CType(Me.pbBP1, System.ComponentModel.ISupportInitialize).EndInit()
        CType(Me.pbPB2, System.ComponentModel.ISupportInitialize).EndInit()
        CType(Me.pbBP6, System.ComponentModel.ISupportInitialize).EndInit()
        CType(Me.pbSQS, System.ComponentModel.ISupportInitialize).EndInit()
        CType(Me.pbBP3, System.ComponentModel.ISupportInitialize).EndInit()
        CType(Me.pbSensitivity, System.ComponentModel.ISupportInitialize).EndInit()
        Me.ResumeLayout(False)
        Me.PerformLayout()

    End Sub

    Friend WithEvents Label4 As Label
    Friend WithEvents cbEnd As Button
    Friend WithEvents pbVm1 As PictureBox
    Friend WithEvents cbSelect As Button
    Friend WithEvents cbPlot As Button
    Friend WithEvents cbCsvFileList As ComboBox
    Friend WithEvents cbselectCSVfile As Button
    Friend WithEvents cbPS As Button
    Friend WithEvents cbManualFit As Button
    Friend WithEvents GroupBox3 As GroupBox
    Friend WithEvents TextBox2 As TextBox
    Friend WithEvents TextBox3 As TextBox
    Friend WithEvents Label64 As Label
    Friend WithEvents Label72 As Label
    Friend WithEvents txtScf1 As TextBox
    Friend WithEvents Label44 As Label
    Friend WithEvents chkNaKRegulation As CheckBox
    Friend WithEvents chkWriteFile As CheckBox
    Friend WithEvents chkDefault As CheckBox
    Friend WithEvents Label17 As Label
    Friend WithEvents txtNP As TextBox
    Friend WithEvents txtStimulus As TextBox
    Friend WithEvents txtEndms As TextBox
    Friend WithEvents txt4th As TextBox
    Friend WithEvents txt3rd As TextBox
    Friend WithEvents txtStartms As TextBox
    Friend WithEvents Label52 As Label
    Friend WithEvents txt2nd As TextBox
    Friend WithEvents Label16 As Label
    Friend WithEvents txtEval As TextBox
    Friend WithEvents txt1st As TextBox
    Friend WithEvents txtRgR As TextBox
    Friend WithEvents txtRgL As TextBox
    Friend WithEvents txtEvalMax As TextBox
    Friend WithEvents Label50 As Label
    Friend WithEvents Label19 As Label
    Friend WithEvents txtMaxRepeat As TextBox
    Friend WithEvents Label11 As Label
    Friend WithEvents Label7 As Label
    Friend WithEvents Label26 As Label
    Friend WithEvents controlBox4 As GroupBox
    Friend WithEvents Label71 As Label
    Friend WithEvents Label70 As Label
    Friend WithEvents Label56 As Label
    Friend WithEvents txtNai As TextBox
    Friend WithEvents chkIbNSC As CheckBox
    Friend WithEvents txtINCXmag As TextBox
    Friend WithEvents txtINaKmag As TextBox
    Friend WithEvents chkINCX As CheckBox
    Friend WithEvents chkINaK As CheckBox
    Friend WithEvents chkIst As CheckBox
    Friend WithEvents chkIKto As CheckBox
    Friend WithEvents chkIKs As CheckBox
    Friend WithEvents chkIha As CheckBox
    Friend WithEvents chkIKur As CheckBox
    Friend WithEvents chkINa As CheckBox
    Friend WithEvents chkICaL As CheckBox
    Friend WithEvents chkIK1 As CheckBox
    Friend WithEvents chkIKr As CheckBox
    Friend WithEvents NFusCaLmag As TextBox
    Friend WithEvents rFusCaL As TextBox
    Friend WithEvents Label60 As Label
    Friend WithEvents txtstdtotCai As TextBox
    Friend WithEvents txtstdNai As TextBox
    Friend WithEvents Label59 As Label
    Friend WithEvents txtFusCaLmag As TextBox
    Friend WithEvents txtVshiftNa As TextBox
    Friend WithEvents Label23 As Label
    Friend WithEvents Label3 As Label
    Friend WithEvents Label5 As Label
    Friend WithEvents txtVshiftCaL As TextBox
    Friend WithEvents Label43 As Label
    Friend WithEvents Label35 As Label
    Friend WithEvents txtKLmag As TextBox
    Friend WithEvents Label25 As Label
    Friend WithEvents Label30 As Label
    Friend WithEvents IKr As Label
    Friend WithEvents txtICaLmag As TextBox
    Friend WithEvents txtIK1mag As TextBox
    Friend WithEvents Label15 As Label
    Friend WithEvents txtIKrmag As TextBox
    Friend WithEvents Label53 As Label
    Friend WithEvents txtNaLratio As TextBox
    Friend WithEvents Label51 As Label
    Friend WithEvents Label6 As Label
    Friend WithEvents txtBaseVshift As TextBox
    Friend WithEvents Label48 As Label
    Friend WithEvents Label9 As Label
    Friend WithEvents txtICaTmag As TextBox
    Friend WithEvents Label34 As Label
    Friend WithEvents txtIKurmag As TextBox
    Friend WithEvents txtOS As TextBox
    Friend WithEvents Label10 As Label
    Friend WithEvents Label13 As Label
    Friend WithEvents Label8 As Label
    Friend WithEvents txtIKsmag As TextBox
    Friend WithEvents txtMDP As TextBox
    Friend WithEvents Label27 As Label
    Friend WithEvents Label14 As Label
    Friend WithEvents rNa As TextBox
    Friend WithEvents Label1 As Label
    Friend WithEvents Label18 As Label
    Friend WithEvents txtCycleL As TextBox
    Friend WithEvents Label28 As Label
    Friend WithEvents txtIKtomag As TextBox
    Friend WithEvents txtAPD As TextBox
    Friend WithEvents rKr As TextBox
    Friend WithEvents txtINamag As TextBox
    Friend WithEvents txtTotalCai As TextBox
    Friend WithEvents rK1 As TextBox
    Friend WithEvents txtcrfNCX As TextBox
    Friend WithEvents Label22 As Label
    Friend WithEvents rVshiftCaL As TextBox
    Friend WithEvents txtcrfNaK As TextBox
    Friend WithEvents rCaL As TextBox
    Friend WithEvents txtIbNSCmag As TextBox
    Friend WithEvents Label24 As Label
    Friend WithEvents txtKi As TextBox
    Friend WithEvents rCaT As TextBox
    Friend WithEvents Label29 As Label
    Friend WithEvents rextraVshift As TextBox
    Friend WithEvents rst As TextBox
    Friend WithEvents txtIhamag As TextBox
    Friend WithEvents rbNSC As TextBox
    Friend WithEvents NVshiftCaL As TextBox
    Friend WithEvents Label47 As Label
    Friend WithEvents rKL As TextBox
    Friend WithEvents NextraVshift As TextBox
    Friend WithEvents rKto As TextBox
    Friend WithEvents NVshiftNa As TextBox
    Friend WithEvents rVshiftNa As TextBox
    Friend WithEvents txtIstmag As TextBox
    Friend WithEvents NIhamag As TextBox
    Friend WithEvents rKs As TextBox
    Friend WithEvents NICaTmag As TextBox
    Friend WithEvents rha As TextBox
    Friend WithEvents NIKurmag As TextBox
    Friend WithEvents rKur As TextBox
    Friend WithEvents Label20 As Label
    Friend WithEvents NIKsmag As TextBox
    Friend WithEvents NIKtomag As TextBox
    Friend WithEvents NIstmag As TextBox
    Friend WithEvents Label21 As Label
    Friend WithEvents NKLmag As TextBox
    Friend WithEvents NINamag As TextBox
    Friend WithEvents NIbNSCmag As TextBox
    Friend WithEvents NIKrmag As TextBox
    Friend WithEvents NIK1mag As TextBox
    Friend WithEvents NICaLmag As TextBox
    Friend WithEvents Label55 As Label
    Friend WithEvents chkVmadaptive As CheckBox
    Friend WithEvents chkVmreference As CheckBox
    Friend WithEvents pbInwardI As PictureBox
    Friend WithEvents txtVmF As TextBox
    Friend WithEvents Label40 As Label
    Friend WithEvents Label38 As Label
    Friend WithEvents pbpsVm As PictureBox
    Friend WithEvents pbOutwardI As PictureBox
    Friend WithEvents Label31 As Label
    Friend WithEvents pbBP5 As PictureBox
    Friend WithEvents Label68 As Label
    Friend WithEvents txtMeanBP4 As TextBox
    Friend WithEvents Label67 As Label
    Friend WithEvents Label42 As Label
    Friend WithEvents Label54 As Label
    Friend WithEvents pbBP4 As PictureBox
    Friend WithEvents TextBox1 As TextBox
    Friend WithEvents txtMeanBP2 As TextBox
    Friend WithEvents txtMeanBP1 As TextBox
    Friend WithEvents Label41 As Label
    Friend WithEvents Label49 As Label
    Friend WithEvents Label12 As Label
    Friend WithEvents Label61 As Label
    Friend WithEvents pbBP1 As PictureBox
    Friend WithEvents pbPB2 As PictureBox
    Friend WithEvents Label66 As Label
    Friend WithEvents txtMeanBP5 As TextBox
    Friend WithEvents Label45 As Label
    Friend WithEvents Label37 As Label
    Friend WithEvents txtStp As TextBox
    Friend WithEvents Label69 As Label
    Friend WithEvents txtdetectINa As TextBox
    Friend WithEvents Label65 As Label
    Friend WithEvents txtPScycle As TextBox
    Friend WithEvents Label32 As Label
    Friend WithEvents txtNBPSQS As TextBox
    Friend WithEvents txtLogMSE As TextBox
    Friend WithEvents pbBP6 As PictureBox
    Friend WithEvents Label58 As Label
    Friend WithEvents txtMSE As TextBox
    Friend WithEvents pbSQS As PictureBox
    Friend WithEvents Label46 As Label
    Friend WithEvents Label62 As Label
    Friend WithEvents txtCellNo As TextBox
    Friend WithEvents txtMeanBP6 As TextBox
    Friend WithEvents Label74 As Label
    Friend WithEvents pbBP3 As PictureBox
    Friend WithEvents Label75 As Label
    Friend WithEvents txtR2 As TextBox
    Friend WithEvents Label76 As Label
    Friend WithEvents Label77 As Label
    Friend WithEvents txtScf2 As TextBox
    Friend WithEvents Label2 As Label
    Friend WithEvents txtorgExtraVshift As TextBox
    Friend WithEvents Label80 As Label
    Friend WithEvents Label79 As Label
    Friend WithEvents txt2R As TextBox
    Friend WithEvents txtR3 As TextBox
    Friend WithEvents txtR1 As TextBox
    Friend WithEvents txtF4 As TextBox
    Friend WithEvents txtF3 As TextBox
    Friend WithEvents txtF2 As TextBox
    Friend WithEvents txtF1 As TextBox
    Friend WithEvents txtMeanBP3 As TextBox
    Friend WithEvents txtI1 As TextBox
    Friend WithEvents txtI2 As TextBox
    Friend WithEvents txtI3 As TextBox
    Friend WithEvents txtI4 As TextBox
    Friend WithEvents txtI5 As TextBox
    Friend WithEvents txtI6 As TextBox
    Friend WithEvents txtVmOnset As TextBox
    Friend WithEvents txtError As TextBox
    Friend WithEvents Label78 As Label
    Friend WithEvents txtPrecision As TextBox
    Friend WithEvents Label39 As Label
    Friend WithEvents Label57 As Label
    Friend WithEvents txtSbp1 As TextBox
    Friend WithEvents txtSbp2 As TextBox
    Friend WithEvents txtSbp3 As TextBox
    Friend WithEvents txtSbp4 As TextBox
    Friend WithEvents txtSbp5 As TextBox
    Friend WithEvents txtSbp6 As TextBox
    Friend WithEvents pbSensitivity As PictureBox
    Friend WithEvents Label63 As Label
    Friend WithEvents Label73 As Label
    Friend WithEvents Label81 As Label
    Friend WithEvents Label82 As Label
    Friend WithEvents Label83 As Label
    Friend WithEvents Label84 As Label
    Friend WithEvents Label85 As Label
    Friend WithEvents txtCLB As TextBox
    Friend WithEvents txtPreviousMIN As TextBox
    Friend WithEvents Label86 As Label
    Friend WithEvents Label36 As Label
    Friend WithEvents txtadr4R As TextBox
    Friend WithEvents Label33 As Label
    Friend WithEvents txtWeight As TextBox
    Friend WithEvents rbEMTest As RadioButton
    Friend WithEvents rbMMtest As RadioButton
    Friend WithEvents chkRand2 As CheckBox
End Class
```
