## Supplementary material for "Gradient-based parameter optimization to determine membrane ionic current composition of human induced pluripotent stem cell-derived cardiomyocytes": optimizing program: Protocol1.vb

```
Public Class Protocol1

#Region "Graph definition"
    '********** Graph option **********
    Dim graphVm1 As Graph2018
    Dim graphpsVm As Graph2018

    Dim graphOutwardI As Graph2018
    Dim graphInwardI As Graph2018

    Dim graphTimeCourse As Graph2018        ' time course of the scf_Ixmag

    Dim graphBP1MSE As Graph2018
    Dim graphBP2MSE As Graph2018
    Dim graphBP3MSE As Graph2018
    Dim graphBP4MSE As Graph2018
    Dim graphBP5MSE As Graph2018
    Dim graphBP6MSE As Graph2018

    Dim graphMSE As Graph2018
    Dim graphSensitivity As Graph2018
#End Region

#Region "Graph index, Max, Min, Grids"
    ''******************************   Graph parameters **********************************
    Dim MinX1 As Double = 0                               'msec, Abscissa of the graph
    Dim MaxX1 As Double = 10000                         '1000,   600
    Dim IntervalX1 As Double = 1000

    Dim MinY1 As Double = -90                          'mV, ordinate, graph of membrane potential
    Dim MaxY1 As Double = 40
    Dim IntervalY1 As Double = 10

    Dim MinX2 As Double = 0                               'msec, Abscissa of the graph
    Dim MaxX2 As Double = 5000         ' 20 sec 1000,   600
    Dim IntervalX2 As Double = 500     ' 

    Dim MinX3 As Double = 0                               'msec, Abscissa of the time course of SQS 
    Dim MaxX3 As Double = 2000                   '300           '1000
    Dim IntervalX3 As Double = 200  '20

    Dim MinY2 As Double = -0.5                      'time course of SQS
    Dim MaxY2 As Double = 3.01
    Dim IntervalY2 As Double = 0.5

    Dim MinX8 As Double = 0                               'msec, Abscissa of the PS Vm graph
    Dim MaxX8 As Double = 2500.01         'ms
    Dim IntervalX8 As Double = 200     ' 

    Dim MinY8 As Double = -1.2                      'pA/pF,  outward Current , graph of Current
    Dim MaxY8 As Double = 0
    Dim IntervalY8 As Double = 0.2

    Dim MinY9 As Double = 0                          'pA/pF,  inward Current , graph of Current
    Dim MaxY9 As Double = 1.2
    Dim IntervalY9 As Double = 0.2

    Dim MinX15 As Double = 0.8                          'BP     Ixsf
    Dim MaxX15 As Double = 1.201
    Dim IntervalX15 As Double = 0.05

    Dim MinX15W As Double = 0.5                          'BP     Ixsf  for a wider scale
    Dim MaxX15W As Double = 1.501
    Dim IntervalX15W As Double = 0.1

    Dim MinY15 As Double = -0.1                      'mV2,   BP_ SQS relationship
    Dim MaxY15 As Double = 1.01
    Dim IntervalY15 As Double = 0.1

    Dim MinX20 As Double = 0.8                          'BP     Ixsf
    Dim MaxX20 As Double = 1.201
    Dim IntervalX20 As Double = 0.1

    Dim MinX20W As Double = 0.5                          'BP     Ixsf  for a wider scale
    Dim MaxX20W As Double = 1.501
    Dim IntervalX20W As Double = 0.1


    Dim MinX21 As Double = 0                    'mV/s,   Sensitivity,  dMSE/dstep- Eval relationship
    Dim MaxX21 As Double = 300
    Dim IntervalX21 As Double = 50

    Dim MinY21 As Double = -4.0                     'uV,     Sensitivity,  dMSE/dstep- Eval relationship
    Dim MaxY21 As Double = 3.01
    Dim IntervalY21 As Double = 1


#End Region

#Region "CSVReadandWrite"

    Public iPS As New IPSmodel1022
    'Public iPS As New IPSmodel0217


    Dim CsvFileNameF As String = "csviPS.csv"
    Dim namesF() As String = New String() {"Vm", "Nai", "Ki", "Catot_blk", "CaMCa", "TnChCa", "SRCa", "Catot_iz", "HCa_iz", "Catot_jnc",
"Ca_SRup", "Catot_SRrl", "Yooo", "Yooc", "Ycoo", "Ycoc", "Ycco", "Yoco", "Yocc", "us_CaL", "Yco_iz",
"Yoc_iz", "Yoo_iz", "Yco_blk", "Yoc_blk", "Yoo_blk", "O_TM", "I2_TM", "Is_TM", "O_LM", "I1_LM",
"I2_LM", "Is_LM", "d_CaT", "f_CaT", "d_st", "f_st", "us_st", "C1_ha", "C2_ha", "O1_ha",
"O2_ha", "O_IK1", "Pbspm", "y1_IKr", "y2_IKr", "y3_IKr", "ua", "ui", "Ov_IKs", "C2IKs_iz",
"OcIKs_iz", "C2IKs_blk", "OcIKs_blk", "y1_IKto", "y2_IKto", "O_ACh", "E1NCX_iz", "I1NCX_iz", "I2NCX_iz", "E1NCX_blk",
"I1NCX_blk", "I2NCX_blk", "P1_6_NaK", "P7_NaK", "P8_13_NaK", "TSCa3", "TSCa3W", "TSCa3S", "TSS", "TSW", "hw", "hp", "halfSL",
"SPM", "ExtraVshift", "IKrmag", "IK1mag", "ICaLmag", "KLmag", "orgVshiftCaL", "IbNSCmag", "Ihamag", "IKsmag",
"IKtomag", "IKurmag", "INamag", "orgVshiftNa", "ICaTmag", "Pstmag", "INaKmag", "INCXmag", "crf_NaK", "crf_NCX"}


    Dim CsvFileNameIxpara As String = "csvIxpara.csv"       'Vm, [ion], Ca buffers,  Ix_parameters  膜イオン電流パラメータEuler 変数
    Dim namesIxpara() As String = New String() {"Vm", "Nai", "Ki", "Catot_blk", "CaMCa", "TnChCa", "SRCa", "Catot_iz", "Catot_jnc",
"Ca_SRup", "Catot_SRrl", "Yooo", "Yooc", "Ycoo", "Ycoc", "Ycco", "Yoco", "Yocc", "us_CaL", "Yco_iz",
"Yoc_iz", "Yoo_iz", "Yco_blk", "Yoc_blk", "Yoo_blk", "O_TM", "I2_TM", "Is_TM", "O_LM", "I1_LM",
"I2_LM", "Is_LM", "d_CaT", "f_CaT", "d_st", "f_st", "us_st", "C1_ha", "C2_ha", "O1_ha",
"O2_ha", "pO_IK1", "Pbspm", "y1_IKr", "y2_IKr", "y3_IKr", "ua", "ui", "Ov_IKs", "C2IKs_iz",
"OcIKs_iz", "C2IKs_blk", "OcIKs_blk", "y1_IKto", "y2_IKto", "O_ACh", "E1NCX_iz", "I1NCX_iz", "I2NCX_iz", "E1NCX_blk",
"I1NCX_blk", "I2NCX_blk", "P1_6_NaK", "P7_NaK", "P8_13_NaK", "TSCa3", "TSCa3W", "TSCa3S", "TSS", "TSW", "hw", "hp", "halfSL"}

    Dim CsvFileNameIxmag As String = "csvIxmag.csv"   '膜イオン電流振幅調節parameter　Ixmag
    Dim namesIxmag() As String = New String() {"IKrmag", "IKurmag", "IKtomag", "IK1mag", "IKsmag",
           "ICaLmag", "KLmag", "IbNSCmag", "Ihamag", "INamag", "INaLratio", "Pstmag", "ICaTmag",
           "INaKmag", "INCXmag", "orgExtraVshift", "baseVshift", "orgVshiftCaL", "orgVshiftNa", "SPM"}

    Dim CsvFileNameIxsf As String = "csvIxsf.csv"     'Ixsf,  パラメータ最適化関連scaling factor
    Dim namesIxsf() As String = New String() {"IKrsf", "IKursf", "IKtosf", "IK1sf", "IKssf", "ICaLsf", "KLsf", "IbNSCsf", "Ihasf",
             "INasf", "Istsf", "ICaTsf", "VshiftCaLsf", "VshiftNasf", "stdNai", "stdtotCai", "crf_NaK", "crf_NCX"}

    Dim CsvFileNameBPSQS As String = "csvBPSQS.csv"     'Random test parameter
    Dim namesBPSQS() As String = New String() {"PScycle", "LogMIN", "BP1", "BP2", "BP3", "BP4", "BP5", "BP6", "crf_NaK",
        "Nai", "crf_NCX", "wholeCai", "Eval", "CR2"}

#End Region

#Region "Physiology"    '********** Time step **********
    Public dt As Double = 0.1   '0.05   '0.05        '  msec, time step  already averaged for every 5 data points recorded with a dt = 0.05

    Public RepeatN As Double                       'Number of repeat the sweeps

    Public sweepTime As Double                 'msec   time for each sweep
    Public numSweep As Double                 'counter of sweeps
    Dim TtoPlot As Double                            'msec timing for plot the point
    Dim maxTtoPlot As Double = 0  'dt * 10          'plot all points

    Dim MaxX As Double = 200000        '************<<<<<<<
    Dim FirstDefinition As Boolean = True      'avoid repetitive definition of graphs and others within a series of simulation runs

    Dim ReconstT(MaxX) As Double               'simulated ms
    Dim ReconstVm(MaxX) As Double               'simulated Vm
    Dim K As Long = 0                                       'counter
    Dim ReconstK As Long = 0                          'number of data points

    Public recDuration As Double               'ms,  duration of the selected segment
    'Public referenceVm As Double = -20    '-57     'mV,
    Public refAdrF(100) As Long                 'adr of crossing the reference level during the falling phase
    Public refAdrR(100) As Long                 'adr of crossing the reference level during the rising phase

    Dim NcrossEventF As Long = 0
    Dim NcrossEventR As Long = 0

    Public targetPoints As Long                              'size of the targetVm array

    Public Lcycle_exp As Long                                 'length of a single cycle    Dim ItestOn As Double = 0
    Public Lcycle_sim As Long                                 'length of a single cycle    Dim ItestOn As Double = 0

    '********* Physiological variables ******************
    'Public Nc As Integer = 60000          '3000 ms in duration, number of cells　を宣言する
    'Public CM(Nc) As CMmodel

    Public pOArray(MaxX)                       'pOpen

    'Public Vm As Double                  'mV,
    Public Vshift As Double
    Public Vshift_temp As Double
    Public peakVm As Double               'mV
    Public VmatpeakpO(2, 1000) As Double                'array for Vm at individual peaks  2; Vshift  1 ;  peak of pO_IKr    0; Vm at individaual peaks
    'Public pOIKr As Double                  'open probability of IKr

    Public StartBit As Long                    'start address of selected segment  
    Public adr As Integer

    Public SweepPoint As Double                'counter within the single sweep
    Public endAdr As Long
    Public repeat As Double
    Public NdataPoints As Long         'number of data point in the Vmshift - Vm at peak pO relationship
    Public Points_PS As Long         'number of data point in the Vmshift - Vm at peak pO relationship
#End Region

#Region "Analysis"
    Public MAmax As Long = 400000                     'size of the memory arrayのサイズ
    Public Vmdataexp(MAmax) As Double              'memory array データを保存しておく二次元配列
    Public VmdataSelect(MAmax) As Double          'selected Dataを保存しておく二次元配列
    Public expVmSmooth(MAmax) As Double
    Public tmpVmRecord(100000) As Double            'temporary Vm array
    Public targetVm(100000) As Double                   'target Vm for reconstruction
    'Public tmpTargetVm(100000) As Double            'target Vm adjusted for Vshift
    Public tgPoints As Long                              'size of the targetVm array

    Public simVm(100000) As Double
    Public simIKr(100000) As Double
    Public simICaL(100000) As Double
    Public simINa(100000) As Double
    Public simIK1(100000) As Double
    Public simIbNSC(100000) As Double
    Public simIha(100000) As Double
    Public simIst(100000) As Double
    Public simIKur(100000) As Double
    Public simIKto(100000) As Double
    Public simIKs(100000) As Double

    Public simINaK(100000) As Double
    Public simINCX(100000) As Double

    Public weightEFvm(100000) As Double

    'Public Vshift_base As Double = -5.0                'correction of Vshift
    Public baseVshift As Double = -5.0
    Public orgExtraVshift As Double = -5.5       'this is defined by the textBox, 
    Public extraVshift As Double
    Public extraVshiftsf As Double = 1

    Public AdrF(100) As Integer                'for the tagetVm()
    Public AdrR(100) As Integer                'for the tagetVm()
    Public AdrF_sim(100) As Integer                'for the simVm()
    Public AdrR_sim(100) As Integer                'for the simVm()

    Public crossEventF As Integer           'counter of the events crossing the reference level during repolariztzion
    Public crossEventR As Integer          'counter of the events crossing the reference level during depolariztzion


    Dim SmoothVm(500000) As Double          'selected Dataを保存しておく二次元配列

    Public Ndpoints As Long = 0                             'total number of data points 

    Dim VmArray(MAmax) As Double            'Vm data

    Dim Startms As Double
    Dim Endms As Double

    Public I As Long

    Public InitialSetting As Boolean = True      'indicate the initial setting

#End Region

    Dim ABC As Double       'dummy
    Private Sub cbPlot_Click(sender As Object, e As EventArgs) Handles cbPlot.Click
        defineGraphs()
        Clear()
        DrawAxesEM()

        '***************** GetOriginalData() *********************************
        '*********** '一行を読み込んで配列に結果を受け取る 'ファイルの終わりまで一行ずつ処理   CSV readに匹敵する****************
        ' ************************************************   TextFieldParser  構文解析プログラムをインスタンス化します。
        reader = New FileIO.TextFieldParser(input_expdata, SHIFT_JIS)
        reader.TextFieldType = FileIO.FieldType.Delimited
        reader.SetDelimiters(",")

        count = 0
        While Not reader.EndOfData                                       '配列の最後まで、data pointを一つづつ取得する
            str = reader.ReadFields()                                             '文字データを配列str()に取得する？

            'Vmdataexp(count) = Double.Parse(str(0))                     'ms, exp. time　今回は必要なし　　実数dbl に変換
            Vmdataexp(count) = Double.Parse(str(1))                      'mV, Vm equivalent as CDbl(str(1))
            count = count + 1                                                           'number of data points  take  every 2 points
            'If count * dt > (MaxX1 - 10) Then Exit While 'limit of the number of data points for the analysis　　dt = 0.05 msec for the experimental recording
            If count * (dt / 2) > (MaxX1 - 10) Then Exit While 'the dt used in the analysis is 0.1 ms,  the number of data points for the analysis MaxX1
        End While

        Ndpoints = count / 1                            'delete the initial small segment, back by 1 ms = 20 * 0.05 ms

        I = 0
        L = 0
        Do
            My.Application.DoEvents()
            VmdataSelect(L) = Vmdataexp(I)
            graphVm1.PlotPoint(L * dt, VmdataSelect(L), Color.Black, 2)      'Display the data in the ms - Vm coordinate
            L = L + 1
            I = I + 2
        Loop While I < Ndpoints


        'txtCount.Text = CStr(Ndpoints)       'print the total number of points which is used for the Data Analysis
        '************************************************************************************************
        cbPlot.Visible = False
        cbSelect.Visible = True


    End Sub

    Dim halfWidth As Long = 20                 'half size of the window used for smoothing
    Dim fullWidth As Long = 2 * halfWidth                'half size of the window used for smoothing
    Dim sum As Double                               'sum of the Vm within the window
    Dim mean As Double                             'average over the width of the original Vm record

    Dim L As Integer
    Dim Quit As Boolean = False

    Public maxRepeat As Double
    Public NSweep As Long = 0

    Public segmentSelect(MAmax) As Double          'selected Dataを保存しておく二次元配列
    Private Sub cbSelect_Click(sender As Object, e As EventArgs) Handles cbSelect.Click
        cbSelect.Visible = False      'hide the command bottun

        txtCellNo.Text = "Cell 86"       'print the Cell No.

        '(process 1) **************************************
        CSVReadAndWriteE15.ReadCSV(CsvFileNameIxpara, iPS, namesIxpara)    'モデル初期値　　initial set of model variables and parameter
        '                                            iPS.InitialValuesCell10()      '初期値が原因で、計算が破綻する場合に使用する。通常は使わない
        CSVReadAndWriteE15.ReadCSV(CsvFileNameIxmag, iPS, namesIxmag)      'initial set of iPS.Ixmag
        printIxmag()                  'illustrate Ixmag in the textBox
        CSVReadAndWriteE15.ReadCSV(CsvFileNameIxsf, iPS, namesIxsf)              'initial set of iPS.Ixsf, totCa
        PrintIxscf()                    'illustrate Ixsf in the textBox 
        '************************************************

        Startms = CDbl(txtStartms.Text)                  'ms,   textBoxから計算対象範囲のStartmsを得る
        Endms = CDbl(txtEndms.Text)                     'ms,   textBoxから計算対象範囲のEndmsを得る

        graphVm1.PlotLine(Startms, MinY1, Startms, MaxY1, Color.Red, 2)
        graphVm1.PlotLine(Endms, MinY1, Endms, MaxY1, Color.Blue, 2)

        StartBit = Startms / dt                                  'points, 最初の点の位置
        countSelect = (Endms - Startms) / dt

        '           PrepareVmdataSelect()  '********* Vmdataexp()の中からVmdataSelect(）をコピーする。
        I = 0               'number of points within the selected dt  
        Do                         'also, the Vshift_temp is executed here !
            segmentSelect(I) = VmdataSelect(StartBit + I)             'mV,  ' Vmdataexp()の中からVmdataSelect(）をコピーする。
            I = I + 1                                        '
        Loop While I <= countSelect         'countSelect     segment for analysis 

        '(process 2)********** Prepare the targetVm **********************************************

        baseVshift = CDbl(txtBaseVshift.Text)   '-textBoxからbaseVshift取得, まず最初にbaseVshiftだけ実験記録をシフトする
        iPS.baseVshift = baseVshift                     'just to keep in the csvFile

        PrepareExpVmSmooth()                  'average (smoothing) and shift by the baseVshift, but no plotting  *******************

        orgExtraVshift = CDbl(txtorgExtraVshift.Text)     '直接textBoxからExtraVshiftをorgExtraVshiftとして取得
        extraVshift = orgExtraVshift * extraVshiftsf                            'BaseVshiftに更に調節可能なExtraVshiftを用いて、Vshiftの調節範囲とする。
        iPS.ExtraVshift = extraVshift                                  'just for recording in the csvFilewrite        PreparetmpTargetVm(expVmSmooth, extraVshift)      'prepare the targetVm consisting of 4 Lcycles of Vm change
        PreparetmpTargetVm(expVmSmooth, extraVshift)      'prepare the targetVm consisting of 4 Lcycles of Vm change, also plot on the screen memory

        My.Application.DoEvents()                                           'display on the graphic display

        APmetrics()                                  'measure the AP metrics, such as OS, AMP, APD, cycle Length, MDP   , display on the graphVm1

        L = 0                  'integer
        Do                                   'Display the smoothed Vm record shifted by the Vshift
            graphVm1.PlotPoint((Startms + L * dt), expVmSmooth(L) + extraVshift, Color.Pink, 1)
            L += 1
        Loop While (Startms + L * dt) < Endms

        DisplaytmpTargetVm()         'Vshift = baseVshift + orgExtraVshift, a temporary target which is further modified by adjusting ExtraVshift
        My.Application.DoEvents()

        defaultStart = False             'use the CSVfile as the initial set of variables of the numerical integration
        cbManualFit.Visible = True  'display the Manual fitting button
        cbSelect.Visible = False

    End Sub


    Public Sub PrepareVmdataSelect()
        '(process 1)   ********* Vmdataexp()の中からVmdataSelect(）を作り、プロットする。******************
        I = 0               'number of points within the selected interval   
        Do                         'also, the Vshift_temp is executed here !
            My.Application.DoEvents()
            VmdataSelect(I) = VmdataSelect(StartBit + I)             'mV, 

            I = I + 1                                        '
        Loop While I < (Endms - Startms) / dt           'countSelect      解析対象segment全長のポイント数
        countSelect = I                                            'number of selected data points
    End Sub

    Public Sub PrepareExpVmSmooth()   '********************* average (smoothing) and baseVshift *******************
        '********************* average (smoothing) and Vshift ****************************************
        sum = 0            'summation over the average window
        I = 0          '********** At the very beginning, execute Smoothing on the experimental record to remove the white noise 
        Do             'Just a single mean at the start      expVmSmooth(0)
            sum = sum + segmentSelect(I)     'summate CM(I) over the width of average window only at the first window 
            I = I + 1
        Loop While I < fullWidth               'length of the average window (41 points = 2 * halfWidth + 1)
        mean = sum / fullWidth               'average
        expVmSmooth(0) = mean + baseVshift       'expVmRecord() is the mean value after addition of  the default Vshift of -5 mV

        L = 1                       'repeat until L = totNum_CM - halfWidth
        Do
            'My.Application.DoEvents()
            Dim M As Integer = L - 1
            Dim N As Integer = L + (fullWidth - 1)
            sum = sum - segmentSelect(M) + segmentSelect(N)    'replace CM(I-1) by CM(I + fullWidth) in the sum
            mean = sum / fullWidth

            expVmSmooth(L) = mean + baseVshift                             '+ extraVshift
            L = L + 1
        Loop While L < MaxX2 / dt                 'stop when the graphVm2 is full 
    End Sub


    Public Sub DisplaytmpTargetVm()
        graphpsVm.Clear()                                                         'prepare graphVm
        graphpsVm.DrawAxis(IntervalX8, IntervalY1, "", 11)
        graphpsVm.PlotLine(Lcycle_exp * dt, MinY1, Lcycle_exp * dt, MaxY1, Color.Red, 1)
        graphpsVm.PlotLine(2 * Lcycle_exp * dt, MinY1, 2 * Lcycle_exp * dt, MaxY1, Color.Red, 1)
        graphpsVm.PlotLine(3 * Lcycle_exp * dt, MinY1, 3 * Lcycle_exp * dt, MaxY1, Color.Red, 1)

        I = 0
        Do                      'display the full length of the targetVm_Org
            My.Application.DoEvents()
            graphpsVm.PlotPoint(I * dt, targetVm(I), Color.Chocolate, 1)  'display the time course of Vm
            I = I + 1
        Loop While I < targetPoints

    End Sub


    Dim Ntot As Integer = 0
    Dim pre As Double            'technical variable
    Dim post As Double           'technical variable
    Public VmOnset As Double           'the first Vm point in the reference Trace

    Public OS As Double = -90         'mV, Overshoot potential of AP
    Public MDP As Double = 50    'mV, maximum diastolic potential

    Public Sub PreparetmpTargetVm(ByVal expVmSmooth() As Double, ByVal extraVshift As Double)      'when an extraVshift is applied
        'Step(1)  'apply the extraVshift to the smoothed Vm recording obtained in experiments  **********************

        L = 0
        Do                 'prepare new targetVm by modifying the expVmRecord() with the extraVshift
            tmpVmRecord(L) = expVmSmooth(L) + extraVshift     'the expVmSmooth() was prepared in cbSelect() 
            L = L + 1
        Loop While L < (MaxX2 / dt + 4000)              '4000  ms is added for safety to cover at least four cycles of spontaneous AP 
        Ntot = L                                                         'number of total points of selected segment of Vm record

        'Step(2A)  '******  determine adress of crossEventF, crossEventR of the targetVm and count four repolarization events **********************
        '********** the targetVm should start at the reference Vm (-20 mV) *****************
        Quit = False                  'reset the boolean 'Quit'

        'Step(2B) '************ detect Events of crossing the reference level (-20 mV)  ***************************
        referenceVm = -20        'mV   only at the beginning    after the modification using the (baseVshift + extraVshift)

        crossEventF = 0            'counter of F events Falling phase
        crossEventR = 0            'counter of R events Rising phase
        adr = 0
        Do           'determine total length of the data array 
            pre = (tmpVmRecord(adr) - referenceVm)
            post = (tmpVmRecord(adr + 1) - referenceVm)
            If pre > 0 And post < 0 Then   'detect crossing -20 mV during a falling phase
                crossEventF = crossEventF + 1        'proceed with the counter'
                AdrF(crossEventF) = adr                  'points, store the location of  crossing reference level (falling phase)
                graphVm1.PlotLine((Startms + adr * dt), MinY1, (Startms + adr * dt), MaxY1, Color.Green, 2)               'indicate the AdrF
                If crossEventF >= 4 Then Quit = True 'exit Do when crossEventF = 4
            ElseIf pre < 0 And post > 0 Then  'detect a rising phase
                crossEventR = crossEventR + 1       'proceed with the counter'
                AdrR(crossEventR) = adr                'store the location of  crossing reference level (rising phase)
                graphVm1.PlotLine((Startms + adr * dt), MinY1, (Startms + adr * dt), MaxY1, Color.Violet, 2)               'indicate the AdrR
            End If

            txtF1.Text = Format(Startms + AdrF(1) * dt, "###0.##")     'print the results of crossEvents
            txtF2.Text = Format(Startms + AdrF(2) * dt, "###0.##")
            txtF3.Text = Format(Startms + AdrF(3) * dt, "###0.##")
            txtF4.Text = Format(Startms + AdrF(4) * dt, "###0.##")

            txtR1.Text = Format(Startms + AdrR(1) * dt, "###0.##")
            txt2R.Text = Format(Startms + AdrR(2) * dt, "###0.##")
            txtR3.Text = Format(Startms + AdrR(3) * dt, "###0.##")

            If Quit = True Then Exit Do 'Exit the Do Loop
            adr = adr + 1
        Loop While adr < Ntot           'length of the smoothed Vm record
        My.Application.DoEvents()           'detect the data plot event

        APmetrics()                     'measure the AP metrics, such as OS, APD, MDP, Lcycle_exp, and plot one cycle in the graphVm1

        '(1) *****  obtain representative 4 cycles of Vm changes,  critical time point of falling and rising phases*************
        tgPoints = 0            ' counter of length of a  target Vm record 
        I = 0
        Do                'prepare the targetVm() consiting of 4 identical segments of Lcycle, Note, AdrF(2) is used in this case
            My.Application.DoEvents()           'detect the data plot event
            targetVm(I) = tmpVmRecord(I + AdrF(2))                                                'the 1st segment                  
            targetVm(I + Lcycle_exp) = tmpVmRecord(I + AdrF(2))                                  'the 2nd segment                    
            targetVm(I + Lcycle_exp + Lcycle_exp) = tmpVmRecord(I + AdrF(2))                    'the 3rd segment    
            targetVm(I + Lcycle_exp + Lcycle_exp + Lcycle_exp) = tmpVmRecord(I + AdrF(2))      'the 4th segment 

            tgPoints = tgPoints + 4                     'count the points of target segment
            I = I + 1
        Loop While I < Lcycle_exp
        targetPoints = tgPoints - 1             'number of data points over the 4 x 2 cycles

        VmOnset = targetVm(0)                 'mV,    determine the new VmOnset , which is very critical to calculate MSE    
        txtVmOnset.Text = Format(VmOnset, "#0.######")

        'I = 0
        'Do                                'plot the targetVm in the graphpsVm
        '    graphpsVm.PlotPoint(I * dt, targetVm(I), Color.Red, 1, 1)
        '    I += 1
        'Loop While I <= tgPoints
        'My.Application.DoEvents()           'detect the data plot event

    End Sub

    Private Sub APmetrics()   '************* measure Action potential parameters ******************* confirmed by AN  220106
        MDP = 50             'temporary MDP
        OS = -90               'temporary OS
        For I As Long = AdrF(2) To AdrF(3)            'measure the MDP and OS potential     One cycle of spontaneous activity
            If tmpVmRecord(I) <= MDP Then MDP = tmpVmRecord(I) 'maximum diastolic  potential
            If tmpVmRecord(I) > OS Then OS = tmpVmRecord(I) 'overshoot potential
            graphVm1.PlotPoint(Startms + I * dt, tmpVmRecord(I), Color.Red, 1, 1)      'Display the data in the ms - Vm coordinate
        Next

        txtMDP.Text = Format(MDP, "###0.#")
        txtOS.Text = Format(OS, "###0.#")
        txtAPD.Text = Format((AdrF(3) - AdrR(2)) * dt, "##0.# ms")

        Lcycle_exp = AdrF(3) - AdrF(2)  'points, length of a single cycle is determined according to an appropriate target segment within the tmpVmrecord(),   

        'With this numbering of the starting AdrF2 to AdrF3 , the second cycle is used, but this is an option
        txtCycleL.Text = Format(Lcycle_exp * dt, "###0.# ms")
        My.Application.DoEvents()
    End Sub


    Public defaultStart As Boolean = True

    Private Sub prepareGraphVmIm()
        graphpsVm.Clear()
        graphpsVm.DrawAxis(IntervalX8, IntervalY1, "", 9)

        If rbMMtest.Checked = True Then
            graphpsVm.PlotLine(limitL, MinY1, limitL, MaxY1, Color.Green, 1)   'ms,
            graphpsVm.PlotLine(limitR, MinY1, limitR, MaxY1, Color.Blue, 1)   'ms,

            adr4R = 0
            graphpsVm.PlotLine(adr0L, MinY1, adr0L, MaxY1, Color.Violet, 1)   'ms,
            graphpsVm.PlotLine(adr1R, MinY1, adr1R, MaxY1, Color.Violet, 1)   'ms,
            graphpsVm.PlotLine(adr2R, MinY1, adr2R, MaxY1, Color.Violet, 1)   'ms,
            graphpsVm.PlotLine(adr3R, MinY1, adr3R, MaxY1, Color.Violet, 1)   'ms,
            graphpsVm.PlotLine(adr4R, MinY1, adr4R, MaxY1, Color.Violet, 1)   'ms,
        End If

        graphOutwardI.Clear()
        graphOutwardI.DrawAxis(IntervalX8, IntervalY8, "", 9)
        graphOutwardI.PlotLine(adr0L, MinY9, adr0L, MaxY9, Color.Blue, 1)   'ms,
        graphOutwardI.PlotLine(adr1R, MinY9, adr1R, MaxY9, Color.Violet, 1)   'ms,
        graphOutwardI.PlotLine(adr2R, MinY9, adr2R, MaxY9, Color.Violet, 1)   'ms,
        graphOutwardI.PlotLine(adr3R, MinY9, adr3R, MaxY9, Color.Violet, 1)   'ms,
        'graphOutwardI.PlotLine(adr4R, MinY9, adr4R, MaxY9, Color.Violet, 1)   'ms,


        graphInwardI.Clear()
        graphInwardI.DrawAxis(IntervalX8, IntervalY8, "", 9)
        graphInwardI.PlotLine(adr0L, MinY8, adr0L, MaxY8, Color.Blue, 1)   'ms,
        graphInwardI.PlotLine(adr1R, MinY8, adr1R, MaxY8, Color.Violet, 1)   'ms,
        graphInwardI.PlotLine(adr2R, MinY8, adr2R, MaxY8, Color.Violet, 1)   'ms,
        graphInwardI.PlotLine(adr3R, MinY8, adr3R, MaxY8, Color.Violet, 1)   'ms,
        'graphInwardI.PlotLine(adr4R, MinY8, adr4R, MaxY8, Color.Violet, 1)   'ms,


    End Sub

    Private Sub Drawlines()
        If rbMMtest.Checked = True Then
            graphpsVm.PlotLine(adr0L, MinY1, adr0L, MaxY1, Color.Violet, 1)   'ms,
            graphpsVm.PlotLine(adr1R, MinY1, adr1R, MaxY1, Color.Violet, 1)   'ms,
            graphpsVm.PlotLine(adr2R, MinY1, adr2R, MaxY1, Color.Violet, 1)   'ms,
            'graphpsVm.PlotLine(adr3R, MinY1, adr3R, MaxY1, Color.Violet, 1)   'ms,
            'graphpsVm.PlotLine(adr4R, MinY1, adr4R, MaxY1, Color.Violet, 1)   'ms,
        End If

        graphOutwardI.PlotLine(adr0L, MinY9, adr0L, MaxY9, Color.Blue, 1)   'ms,
        graphOutwardI.PlotLine(adr1R, MinY9, adr1R, MaxY9, Color.Violet, 1)   'ms,
        graphOutwardI.PlotLine(adr2R, MinY9, adr2R, MaxY9, Color.Violet, 1)   'ms,
        'graphOutwardI.PlotLine(adr3R, MinY9, adr3R, MaxY9, Color.Violet, 1)   'ms,

        graphInwardI.PlotLine(adr0L, MinY8, adr0L, MaxY8, Color.Blue, 1)   'ms,
        graphInwardI.PlotLine(adr1R, MinY8, adr1R, MaxY8, Color.Violet, 1)   'ms,
        graphInwardI.PlotLine(adr2R, MinY8, adr2R, MaxY8, Color.Violet, 1)   'ms,
        'graphInwardI.PlotLine(adr3R, MinY8, adr3R, MaxY8, Color.Violet, 1)   'ms,

    End Sub


    '************* preliminary parameter optimization in a manual manner *************
    Dim constraints As Boolean = False        'use the usual full model equations of Ionic currents

    '********** memory array of expData and Pattern Search  ***********************************
    Public ArraySize As Long = 100000   ' = 5000 / 0.05            'assuming dt = 0.05

    Private Sub defineGraphs()
        graphVm1 = New Graph2018(pbVm1, MinX1, MaxX1, MinY1, MaxY1)
        graphpsVm = New Graph2018(pbpsVm, MinX8, MaxX8, MinY1, MaxY1)

        graphOutwardI = New Graph2018(pbOutwardI, MinX8, MaxX8, MinY9, MaxY9)
        graphInwardI = New Graph2018(pbInwardI, MinX8, MaxX8, MinY8, MaxY8)

        graphBP1MSE = New Graph2018(pbBP1, MinX15, MaxX15, MinY15, MaxY15)       'relationship between SQS and Ixsf
        graphBP2MSE = New Graph2018(pbPB2, MinX15, MaxX15, MinY15, MaxY15)   'relationship between SQS and Ixsf
        graphBP3MSE = New Graph2018(pbBP3, MinX15, MaxX15, MinY15, MaxY15)   'relationship between SQS and Ixsf
        graphBP4MSE = New Graph2018(pbBP4, MinX15, MaxX15, MinY15, MaxY15)        'relationship between SQS and Ixsff
        graphBP5MSE = New Graph2018(pbBP5, MinX15, MaxX15, MinY15, MaxY15)        'relationship between SQS and Ixsff
        graphBP6MSE = New Graph2018(pbBP6, MinX15, MaxX15, MinY15, MaxY15)        'relationship between SQS and Ixsff

        graphMSE = New Graph2018(pbSQS, MinX3, MaxX3, MinY2, MaxY2)
        graphSensitivity = New Graph2018(pbSensitivity, MinX21, MaxX21, MinY21, MaxY21)

    End Sub

    Private Sub defineGraphsMM()
        'graphVm1 = New Graph2018(pbVm1, MinX1, MaxX1, MinY1, MaxY1)
        'graphpsVm = New Graph2018(pbpsVm, MinX8, MaxX8, MinY1, MaxY1)

        'graphOutwardI = New Graph2018(pbOutwardI, MinX8, MaxX8, MinY9, MaxY9)
        'graphInwardI = New Graph2018(pbInwardI, MinX8, MaxX8, MinY8, MaxY8)

        graphBP1MSE = New Graph2018(pbBP1, MinX20, MaxX20, MinY21, MaxY21)       'relationship between SQS and Ixsf
        graphBP2MSE = New Graph2018(pbPB2, MinX20, MaxX20, MinY21, MaxY21)   'relationship between SQS and Ixsf
        graphBP3MSE = New Graph2018(pbBP3, MinX20, MaxX20, MinY21, MaxY21)   'relationship between SQS and Ixsf
        graphBP4MSE = New Graph2018(pbBP4, MinX20, MaxX20, MinY21, MaxY21)        'relationship between SQS and Ixsff
        graphBP5MSE = New Graph2018(pbBP5, MinX20, MaxX20, MinY21, MaxY21)        'relationship between SQS and Ixsff
        graphBP6MSE = New Graph2018(pbBP6, MinX20, MaxX20, MinY21, MaxY21)        'relationship between SQS and Ixsff

        'graphIK1sfSQS = New Graph2018(pbIK1sfSQS, MinX15, MaxX15, MinY15, MaxY15)        'relationship between SQS and Ixsff
        'graphIKtosfSQS = New Graph2018(pbIKtosfSQS, MinX15, MaxX15, MinY15, MaxY15)        'relationship between SQS and Ixsff
        'graphIKssfSQS = New Graph2018(pbBP6, MinX15, MaxX15, MinY15, MaxY15)        'relationship between SQS and Ixsff

        graphMSE = New Graph2018(pbSQS, MinX3, MaxX3, MinY21, MaxY21)

        'graphVshiftCaLsfSQS = New Graph2018(pbVshiftCaLsf, MinX15, MaxX15, MinY15, MaxY15)        'relationship between SQS and Ixs

    End Sub


    Private Sub DrawAxesEM()
        graphVm1.DrawAxis(IntervalX1, IntervalY1, "", 9)
        graphpsVm.DrawAxis(IntervalX8, IntervalY1, "", 9)

        graphOutwardI.DrawAxis(IntervalX8, IntervalY8, "", 9)
        graphInwardI.DrawAxis(IntervalX8, IntervalY8, "", 9)

        graphBP1MSE.DrawAxis(IntervalX15, IntervalY15, "", 10)                 'X- and Y-index
        graphBP2MSE.DrawAxis(IntervalX15, IntervalY15, "", 10)
        graphBP3MSE.DrawAxis(IntervalX15, IntervalY15, "", 10)
        graphBP4MSE.DrawAxis(IntervalX15, IntervalY15, "", 10)
        graphBP5MSE.DrawAxis(IntervalX15, IntervalY15, "", 10)
        graphBP6MSE.DrawAxis(IntervalX15, IntervalY15, "", 10)

        graphMSE.DrawAxis(IntervalX3, IntervalY2, "", 8)
        graphMSE.PlotLine(MinX3, crtMSEdepthA, MaxX3, crtMSEdepthA, Color.Chocolate, 1)             'zero hilizontal level of SQS = 1 mV^2

        graphBP1MSE.PlotLine(MinX15, 0, MaxX15, 0, Color.Black, 1)             'zero hilizontal level of SQS = 1 mV^2
        graphBP2MSE.PlotLine(MinX15, 0, MaxX15, 0, Color.Black, 1)
        graphBP3MSE.PlotLine(MinX15, 0, MaxX15, 0, Color.Black, 1)
        graphBP4MSE.PlotLine(MinX15, 0, MaxX15, 0, Color.Black, 1)
        graphBP5MSE.PlotLine(MinX15, 0, MaxX15, 0, Color.Black, 1)
        graphBP6MSE.PlotLine(MinX15, 0, MaxX15, 0, Color.Black, 1)

        graphBP1MSE.PlotLine(1, MinY15, 1, MaxY15, Color.Black, 1)            'zero vertidcal line  = 1 vertical line
        graphBP2MSE.PlotLine(1, MinY15, 1, MaxY15, Color.Black, 1)
        graphBP3MSE.PlotLine(1, MinY15, 1, MaxY15, Color.Black, 1)
        graphBP4MSE.PlotLine(1, MinY15, 1, MaxY15, Color.Black, 1)
        graphBP5MSE.PlotLine(1, MinY15, 1, MaxY15, Color.Black, 1)
        graphBP6MSE.PlotLine(1, MinY15, 1, MaxY15, Color.Black, 1)

        graphSensitivity.DrawAxis(IntervalX21, IntervalY21, "", 8)

    End Sub


    Private Sub DrawAxesMM()
        'graphVm1.DrawAxis(IntervalX1, IntervalY1, "", 9)
        'graphpsVm.DrawAxis(IntervalX8, IntervalY1, "", 9)

        'graphOutwardI.DrawAxis(IntervalX8, IntervalY8, "", 9)
        'graphInwardI.DrawAxis(IntervalX8, IntervalY8, "", 9)

        graphBP1MSE.DrawAxis(IntervalX20, IntervalY21, "", 10)                 'X- and Y-index
        graphBP2MSE.DrawAxis(IntervalX20, IntervalY21, "", 10)
        graphBP3MSE.DrawAxis(IntervalX20, IntervalY21, "", 10)
        graphBP4MSE.DrawAxis(IntervalX20, IntervalY21, "", 10)
        graphBP5MSE.DrawAxis(IntervalX20, IntervalY21, "", 10)
        graphBP6MSE.DrawAxis(IntervalX20, IntervalY21, "", 10)

        graphMSE.DrawAxis(IntervalX3, IntervalY21, "", 8)
        graphMSE.PlotLine(MinX3, crtMSEdepthA, MaxX3, crtMSEdepthA, Color.Chocolate, 1)             'zero hilizontal level of SQS = 1 mV^2
        graphMSE.PlotLine(MinX3, crtMSEdepthB, MaxX3, crtMSEdepthB, Color.Magenta, 1)             'zero hilizontal level of SQS = 1 mV^2

        graphBP1MSE.PlotLine(MinX20, 0, MaxX20, 0, Color.Black, 1)             'zero hilizontal level of SQS = 1 mV^2
        graphBP2MSE.PlotLine(MinX20, 0, MaxX20, 0, Color.Black, 1)
        graphBP3MSE.PlotLine(MinX20, 0, MaxX20, 0, Color.Black, 1)
        graphBP4MSE.PlotLine(MinX20, 0, MaxX20, 0, Color.Black, 1)
        graphBP5MSE.PlotLine(MinX20, 0, MaxX20, 0, Color.Black, 1)
        graphBP6MSE.PlotLine(MinX20, 0, MaxX20, 0, Color.Black, 1)

        graphBP1MSE.PlotLine(1, MinY21, 1, MaxY21, Color.Black, 1)            'zero vertidcal line  = 1 vertical line
        graphBP2MSE.PlotLine(1, MinY21, 1, MaxY21, Color.Black, 1)
        graphBP3MSE.PlotLine(1, MinY21, 1, MaxY21, Color.Black, 1)
        graphBP4MSE.PlotLine(1, MinY21, 1, MaxY21, Color.Black, 1)
        graphBP5MSE.PlotLine(1, MinY21, 1, MaxY21, Color.Black, 1)
        graphBP6MSE.PlotLine(1, MinY21, 1, MaxY21, Color.Black, 1)

    End Sub
    Private Sub Clear()
        'graphVm1.Clear()
        graphpsVm.Clear()

        graphInwardI.Clear()
        graphOutwardI.Clear()


    End Sub

    Private Sub ClearMM()
        graphBP1MSE.Clear()
        graphBP2MSE.Clear()
        graphBP3MSE.Clear()
        graphBP4MSE.Clear()
        graphBP5MSE.Clear()
        graphBP6MSE.Clear()

        graphMSE.Clear()

    End Sub

    '************   parameters use for the pattern search ********************* 
    Public BP(20) As Double          'base point for the best fit routine
    Public oldBP(20) As Double          'last base point for the best fit routine
    Public NP(20) As Double          'new point for the best fit routine
    Public oldNP(20) As Double          'previous point for the best fit routine
    Public Stp(20) As Double          'step size  for movement searching for smaller deviation
    Public RedFct(20) As Double    'reduction factor to decrease the step size
    Public CrtStp(20) As Double     'critical size of the step to quit the search 
    Public Advance(20) As Double        '        best fit routine

    Public N_BPSQS As Double = 0
    Public PScycle As Double = 1

    Dim Eval As Double = 0                        'number of trials (calculating the given equation)
    Dim falseEval As Double = 0

    Dim oldEval As Double = 1                'number of trials (calculating the given equation)
    Dim crtEval As Double = 100

    Dim crtLogMINdepth As Double
    Dim EvalMax As Double = 1000                 'number of trials (calculating the given equation)

    ' ************* variable and parameters for the PatternSearch()
    Dim PAT As Double
    Dim Difference As Double           'deviation from the theoretical value,  y - y(theory)
    Dim MSE As Double                     'mean square error
    Dim MSEp As Double                     'mean square error at NP() positive step
    Dim MSEn As Double                     'mean square error at NP() negative step
    Dim oldMSE As Double = 10                   'previous mean square error
    Dim Delta_sqs As Double                    'delta SQS

    Dim MIN As Double                    'minimum vaue of SQS
    Public MinSto As Double           'Overall minimum of SQS
    Dim EPS As Double = 1.0E+37   'latest Stp
    Dim Fails As Double
    Dim HOME As Double            'a new origin
    Dim Value As Double

    Dim Nplot As Double = 0
    Public scfNai As Double = 1            '0.3, 0.35, 0.37, 0.4  amplifying factor
    Public scfNaK As Double = 1

    Dim valueX As Double
    Dim Ndefinition As Integer = 0

    Private Sub cbManualFit_Click(sender As Object, e As EventArgs) Handles cbManualFit.Click
        '********  Just to indicate the name of this program name on the Desiner screen     This part is usually fixed *******************
        Dim appPath As String = System.Reflection.Assembly.GetExecutingAssembly().Location  '使用しているプログラムのDirectory
        TextBox1.Text = appPath                   'Print the whole directory to indicate which program is used
        '************** Analysis starts here ******************************

        cbManualFit.Visible = False        'hide the command 'cbFit'
        'chkWriteFile.Checked = True

        '(1)  get intial set of Model variables for the numerical integration **************************
        If InitialSetting = True Then                 'only at the beginning of the SQS calculation　　　最初に１回だけ初期値を定める
            maxRepeat = CLng(txtMaxRepeat.Text)      'max repeat in GetsimVm()   Sweep number 

            iPS.stdNai = CDbl(txtstdNai.Text)               'target stdNai for the 'sub regulateINaK()'
            iPS.stdtotCai = CDbl(txtstdtotCai.Text)      'target stdCa for the 'sub regulate INCX()'

            limitL = CDbl(txtRgL.Text)    'left border of the time window to calculate SQS
            limitR = CDbl(txtRgR.Text)    'left border of the time window to calculate SQS

            txtCellNo.Text = "Cell 86 "       'print the Cell No.

            CSVReadAndWriteE15.ReadCSV(CsvFileNameIxpara, iPS, namesIxpara)    'モデル初期値　　initial set of model variables and parameter
            '                                            iPS.InitialValuesCell10()      '初期値が原因で、計算が破綻する場合に使用する。通常は使わない
            CSVReadAndWriteE15.ReadCSV(CsvFileNameIxmag, iPS, namesIxmag)      'initial set of iPS.Ixmag
            printIxmag()                  'illustrate Ixmag in the textBox
            CSVReadAndWriteE15.ReadCSV(CsvFileNameIxsf, iPS, namesIxsf)              'initial set of iPS.Ixsf, totCa
            PrintIxscf()                    'illustrate Ixsf in the textBox 
            '*********************************************************************************************
            InitialSetting = False    'This initial setting is not used after this sub method
        End If


        '*******************  two options for the ormpEM and ormpMM tests ********************************

        If rbEMTest.Checked = True Then      'option ormpEM test
            If Ndefinition = 0 Then         'avoid repetitive definition of graphs
                defineGraphs()        'prepare Graphs for the EM test
                Clear()
                DrawAxesEM()

                Ndefinition += 1
            End If

            I = 0          'demonstrate the experimental Vm recording 
            L = 0
            Do
                graphVm1.PlotPoint(L * dt, VmdataSelect(L), Color.Black, 2)      'Display the data in the ms - Vm coordinate
                L += 1
            Loop While L < Ndpoints / 2
            My.Application.DoEvents()

            For I As Integer = 1 To 10             'determine the CrtStp
                CrtStp(I) = 8 / 20000
            Next I
            'scf2 = 0.01
            txtPrecision.Text = "0.000001"     'print the Precision
            crtMSEdepthA = 1.0    'critical level to activate the Random2
            graphMSE.PlotLine(MinX3, crtMSEdepthA, MaxX3, crtMSEdepthA, Color.Red, 1)
            rstIxmag = True          'Boolean,  enable resetting Ixmag when new MSE is better than before

            '****  In order to get a new Ixmag  or maxP of the current system Ix by directly changing the text boxes
            GetNewIxsf()        'take Ixsf only when it is different from the oldIxsf
            GetNewIxmag()    'renew  the set of preliminary Ixmags and a final orgextraVshift  for modifying the standard 'iPSC model'    
            PrintNewIxmag()   'calculate the (NIxmag * Ixsf) and print in the text box NIxmag

            extraVshift = orgExtraVshift * extraVshiftsf           'correct the targetVm() for the Vshift induced during the experimental recording        
            PreparetmpTargetVm(expVmSmooth, extraVshift)      'prepare new targetVm() and the AdrF()　and plot in the graph

            '(5) *********    Numeric integration of the model  to compare between expVm and simVm ****************
            RNaKNCX = True                                        'boolean, to indicate use, or not use the regulation of capacity of the NaK and NCX
            GetSimVm(RNaKNCX, maxRepeat)            'Numeric Integration of the Model using  simVm() for adjusting Ixmag through the textBox

        ElseIf rbMMtest.Checked = True Then            'if rbMMtest.checked = True,         option ormpMM test  

            If Ndefinition = 0 Then          'avoid repetitive definition of graphs
                defineGraphsMM()         'prepare Graphs for the MM test              
                Clear()
                ClearMM()
                DrawAxesMM()
                Ndefinition += 1
            End If

            For I As Integer = 1 To 10
                CrtStp(I) = 2 / 20000
            Next I

            'scf2 = 0.01
            txtPrecision.Text = "0.000001"
            crtMSEdepthA = -2.5        'critical Log(MSE) to activate the Random2
            'graphMSE.PlotLine(MinX3, crtMSEdepthA, MaxX3, crtMSEdepthA, Color.Red, 1)

            limitL = CDbl(txtRgL.Text)    'left border of the time window to calculate SQS
            limitR = CDbl(txtRgR.Text)    'left border of the time window to calculate SQS

            rstIxmag = False        'Boolean,  avoid resetting Ixmag irrespetively If a new MSE is better or worse than before

            RNaKNCX = True                                        'boolean, to indicate use, or not use the regulation of capacity of the NaK and NCX
            GetSimVm(RNaKNCX, maxRepeat)            'Numeric Integration of the Model using  simVm() for adjusting Ixmag through the textBox

            L = 0
            Do                                    'determine the targetVm()
                targetVm(L) = simVm(L)
                L += 1
            Loop While L < MaxX8 / dt
            My.Application.DoEvents()
        End If


        '***************  calculation of SQS ************************
        '********** provided that the targetVm() has been prepared from the experimental recording or by using the GetsimVm()

        CalculateMSE_MF()

        If chkWriteFile.Checked = True Then         'This part is necessary in the sub manualfitting(),  Note; the initial values for the numerical integration should not be change in the Pattern Search
            CSVReadAndWriteE15.CreateNewCSV(CsvFileNameIxpara, iPS, namesIxpara)   'Replace the old CSV file with a new one
            CSVReadAndWriteE15.WriteCSV(CsvFileNameIxpara, I, iPS, 1, namesIxpara)     '膜イオン電流パラメータEuler 変数
            CSVReadAndWriteE15.CreateNewCSV(CsvFileNameIxmag, iPS, namesIxmag)   'Replace the old CSV file with a new one
            CSVReadAndWriteE15.WriteCSV(CsvFileNameIxmag, I, iPS, 1, namesIxmag)     '膜イオン電流パラメータEuler 変数
            CSVReadAndWriteE15.CreateNewCSV(CsvFileNameIxsf, iPS, namesIxsf)   'Replace the old CSV file with a new one
            CSVReadAndWriteE15.WriteCSV(CsvFileNameIxsf, I, iPS, 1, namesIxsf)     '膜イオン電流パラメータEuler 変数
        End If

        cbManualFit.Visible = True                  'display the cbFit
        Nfitting = Nfitting + 1
        If Nfitting = 3 Then cbPS.Visible = True '3 numerical integration (minimal) might be ready for the pattern search. but this manual fitting can be continued for satisfaction.

    End Sub


    Private Sub GetNewIxmag()               ' cover all possible Ixmags for simplicity
        orgExtraVshift = CDbl(txtorgExtraVshift.Text)
        iPS.orgExtraVshift = orgExtraVshift                 'to save in the csvFileIxmag

        iPS.baseVshift = CDbl(txtBaseVshift.Text)

        '*********** data points of Ix are saved in each memory array, to make the program more visible ***************

        iPS.IKrmag = CDbl(txtIKrmag.Text)
        iPS.IKurmag = CDbl(txtIKurmag.Text)

        iPS.ICaLmag = CDbl(txtICaLmag.Text)
        iPS.orgVshiftCaL = CDbl(txtVshiftCaL.Text)         'original voltage shift can be modified by NP() in patternSearch()
        iPS.KLmag = CDbl(txtKLmag.Text)                              'Ca-dependency of ICaL inactivation

        iPS.IbNSCmag = CDbl(txtIbNSCmag.Text)                 'represents a net influence of several backgroun currents
        iPS.IK1mag = CDbl(txtIK1mag.Text)      'only when the MDP < -75 mV

        '*********** parameters without the pattern Search in the latest version ***************
        iPS.IKsmag = CDbl(txtIKsmag.Text)         'under the control condition IKs is relatively small and do not influence SDD in HuVEC model 
        iPS.Ihamag = CDbl(txtIhamag.Text)                'shuld be a target of PatternSearch() when MDP < -70 mV
        iPS.IKtomag = CDbl(txtIKtomag.Text)            'negligible  influence on the slow diastolic depolarization
        iPS.INamag = CDbl(txtINamag.Text)              'adjusted according to the maximum rate of rise of AP
        iPS.orgVshiftNa = CDbl(txtVshiftNa.Text)
        iPS.INaLratio = CDbl(txtNaLratio.Text)          'to adjust the EAD events    
        iPS.Pstmag = CDbl(txtIstmag.Text)                  'Ist permeability  relatively small influence on SDD

        '***************  Ion transporters ************************************
        iPS.INaKmag = CDbl(txtINaKmag.Text)     'adjust INa/Kmag via textBox. see sub iPS.RegulatemaxINaK(), which automatically adjusted outside of the pattern Search to keep [Na]i = 6.4 mM 
        iPS.INCXmag = CDbl(txtINCXmag.Text)    'fixed, but automatically changes significantly through changes in [Na]i 

    End Sub


    Dim oldextraVshiftsf As Double

    Dim oldIKrsf As Double
    Dim oldIKursf As Double
    Dim oldICaLsf As Double
    Dim oldVshiftCaLsf As Double
    Dim oldKLsf As Double
    Dim oldIbNSCsf As Double
    Dim oldIK1sf As Double
    Dim oldIKssf As Double
    Dim oldIhasf As Double
    Dim oldIKtosf As Double
    Dim oldINasf As Double
    Dim oldVshiftNasf As Double
    Dim oldIstsf As Double
    Dim oldICaTsf As Double

    Private Sub GetNewIxsf()
        extraVshiftsf = CDbl(rextraVshift.Text)                'iPS.extraVshiftsf　basically, vshiftはiPSC-CMモデルに含まれない

        iPS.IKrsf = CDbl(rKr.Text)                                           'iPS.IKrsf
        iPS.IKursf = CDbl(rKur.Text)
        iPS.ICaLsf = CDbl(rCaL.Text)
        iPS.VshiftCaLsf = CDbl(rVshiftCaL.Text)
        iPS.KLsf = CDbl(rKL.Text)
        iPS.IbNSCsf = CDbl(rbNSC.Text)
        iPS.IK1sf = CDbl(rK1.Text)
        iPS.IKssf = CDbl(rKs.Text)
        iPS.Ihasf = CDbl(rha.Text)
        iPS.IKtosf = CDbl(rKto.Text)
        iPS.INasf = CDbl(rNa.Text)
        iPS.VshiftNasf = CDbl(rVshiftNa.Text)
        iPS.Istsf = CDbl(rst.Text)

    End Sub

    Public Sub defineOldIxsf()
        oldextraVshiftsf = iPS.extraVshiftsf       '  210909

        oldIKrsf = iPS.IKrsf
        oldIKursf = iPS.IKursf
        oldICaLsf = iPS.ICaLsf
        oldVshiftCaLsf = iPS.VshiftCaLsf
        oldKLsf = iPS.KLsf
        oldIbNSCsf = iPS.IbNSCsf
        oldIK1sf = iPS.IK1sf
        oldIKssf = iPS.IKssf
        oldIhasf = iPS.Ihasf
        oldIKtosf = iPS.IKtosf
        oldINasf = iPS.INasf
        oldVshiftNasf = iPS.VshiftNasf
        oldIstsf = iPS.Istsf
        'oldICaTsf = iPS.ICaTsf

    End Sub

    Public Sub PrintNewIxmag()              'Ixmag is given by the product of (Ixmag x Ixsf)
        NextraVshift.Text = Format(orgExtraVshift * extraVshiftsf, "#0.#####")

        '************ print new Ixmag **********************
        NIKrmag.Text = Format(iPS.IKrmag * iPS.IKrsf, "#0.#####")   ' only to monitor the change in 'Ixmag'
        NIK1mag.Text = Format(iPS.IK1mag * iPS.IK1sf, "#0.#####")
        NICaLmag.Text = Format(iPS.ICaLmag * iPS.ICaLsf, "#0.#####")
        NKLmag.Text = Format(iPS.KLmag * iPS.KLsf, "#0.#####")
        NVshiftCaL.Text = Format(iPS.orgVshiftCaL * iPS.VshiftCaLsf, "#0.#####")
        NIbNSCmag.Text = Format(iPS.IbNSCmag * iPS.IbNSCsf, "#0.#####")
        NIhamag.Text = Format(iPS.Ihamag * iPS.Ihasf, "#0.#####")
        NIKsmag.Text = Format(iPS.IKsmag * iPS.IKssf, "#0.#####")
        NIKtomag.Text = Format(iPS.IKtomag * iPS.IKtosf, "#0.#####")
        NIKurmag.Text = Format(iPS.IKurmag * iPS.IKursf, "#0.#####")
        NINamag.Text = Format(iPS.INamag * iPS.INasf, "#0.#####")
        NVshiftNa.Text = Format(iPS.orgVshiftNa * iPS.VshiftNasf, "#0.#####")
        NIstmag.Text = Format(iPS.Pstmag * iPS.Istsf, "#0.#####")

        '********** print stp size *****************************
        txtStp.Text = Format(Stp(1), "#0.#####")   'step size of NP points

    End Sub


    Public VmF As Double                        'final Vm at the end of GetSimVm()
    Public VmA As Double
    Public VmB As Double
    Dim J As Long = 0                           'local counter
    Dim referenceVm As Double = -20.0         'mV,  threshold to decide Quit 
    Dim widthQ As Double = 0.002
    Dim referenceQ As Double = referenceVm - widthQ     '-20.003

    Private Sub GetSimVm(ByVal RNaKNCX As Boolean, ByVal maxRepeat As Long)                  'prepare SimVm(), which consists of 4 cycles of spontaneous AP, finish at the crossing event across the reference level.
        RepeatN = 0                     'reset the number of simulation cycles 
        Do
            '***************** Start of a Single Sweep  ********************************************************
            Quit = False
            crossEventF = 0            'reset counter crossEventF    This is used for synchronizing the start of a Vm Sweep
            crossEventR = 0            'reset counter crossEventF    This is used for synchronizing the start of a Vm Sweep   

            I = 0
            Do                'establish steady rhythm, a single sweep to adapt to the new set of Ixmags. 

                iPS.ModelCell(I * dt)
                iPS.EulerIntegrationStd(dt)
                simVm(I) = iPS.Vm                  'copy the new Vm in the simVm() array
                If RepeatN = (maxRepeat - 1) And I <= MaxX8 / dt Then     'to save time of graphic display
                    StoreIm(I)         'Data are kept in each memory array of individual current system  
                End If

                If I > 1 Then DetectEvents()            'Quit or not      adjust the Vm at the end of each sweep

                If Quit = True Then Exit Do 'exit this Do Loop of numerical integration   This step is essential

                I = I + 1                  'proceed with I
            Loop While I < (MaxX8 + 6000) / dt              'the extra 3000 msec is necessary to safely obtain the 3rd crossEventF
            'txtVmF.Text = Format(iPS.Vm, "##0.0######")     'print Vm at the end of sweep

            If RNaKNCX = True Then        'include the adjusting procedure of NaK and NCX activities to keep [Na]i and Catot in a limited range
                iPS.RegulatemaxINaK()    'iPS.crf_NaK = iPS.crf_NaK - (iPS.stdNai - iPS.Nai) * 0.4     INaKmag to fix [Na]i close to stdNai (6.4 mM)
                txtNai.Text = Format(iPS.Nai, "##0.### ")
                iPS.RegulateCaitotal()
                iPS.wholeCai = iPS.totalCai
                txtTotalCai.Text = Format(iPS.totalCai, "##0.### ")

                txtcrfNaK.Text = Format(iPS.crf_NaK, "##.0###")     'correction factor of Na/K pump to keep [Na]i = stdNai
                txtcrfNCX.Text = Format(iPS.crf_NCX, "##.0###")     'correction factor of Na/Ca exchanger to keep [Ca]i = stdCai
            End If

            RepeatN = RepeatN + 1                               'reset the number of simulation cycles 
        Loop While RepeatN < maxRepeat                'ideally repeat until a steady cycle of spontaneous activity is established
        '************************************* detailed adjusstment of FVm ***********************************************
        VmA = iPS.Vm                      'copy the last Vm

        referenceVm = VmOnset     ' finer adjestment
        referenceQ = VmOnset - widthQ    ' finer adjestment

        K = 0
        Quit = False
        Do                'decide how to quit the repetition of sweeps 
            iPS.ModelCell(I * dt)
            iPS.EulerIntegrationStd(dt)
            VmB = iPS.Vm                  'copy the new Vm in the simVm() array

            '*************  DetectEventsEach() *************************
            pre = VmA - referenceVm
            post = VmB - referenceVm
            If (pre * post) < 0 Then            'Vm has crossed the reference level (-20 mV)
                If pre > 0 Then                    'falling phase
                    Dim valueZ As Double = (VmB - referenceQ)
                    If valueZ >= 0 Then Quit = True                    'Note, the numerical integration  is quitted at the point of crossEventF  
                End If
            End If
            If Quit = True Then Exit Do     'Quit Do Loop >>>>>>>>>>>>>>>>>>>>>>>>

            VmA = VmB               'replace VmA by VmB
            '********************************************************
            I = I + 1
            K = K + 1                  'proceed with K
        Loop While K < 200000 / dt              'to get onsetVm (simVm(0)) closer to -20 mV
        'Loop While K < 60000 / dt

        VmF = iPS.Vm
        txtVmF.Text = Format(VmF, "##0.0###0")                                      'print the final Vm
        txtError.Text = Format((VmF - VmOnset), "##0.0###0")                  'print the deviation from the VmOnset

        '***********************************************************************************
        If Quit = True Then
            simVm(0) = VmF                  'copy the new Vm in the simVm() array
            I = 1
            Do                'obtain final simVm(), which is used as the adaptive Vm segment
                iPS.ModelCell(I * dt)
                iPS.EulerIntegrationStd(dt)
                simVm(I) = iPS.Vm                  'copy the new Vm in the simVm() array
                StoreIm(I)         'Data are kept in each memory array of individual current system  
                I += 1
            Loop While I < MaxX8 / dt              'the extra 3000 msec is necessary to safely obtain the 3rd crossEventF

            prepareGraphVmIm()              'Clear and drawAxes of Vm, outward  and inward current graphs before the new plot

        End If

        NSweep = NSweep + 1

        'My.Application.DoEvents()         'show the traces in the graph at once to avoid time for plotting individual points at every numerical integration

    End Sub


    Private Sub DetectEvents()  '********* quit numerical integration at the time point of crossing the reference leve
        pre = simVm(I - 1) - referenceVm
        post = simVm(I) - referenceVm

        If (pre * post) < 0 Then            'Vm is crossing the reference level
            If pre < 0 Then                    'Rising phase
                crossEventR = crossEventR + 1         'counting
                AdrR_sim(crossEventR) = I                'store the adress (location) of  crossing reference level
            Else                                    'Falling phase
                crossEventF = crossEventF + 1               'count the crossing eventF'
                AdrF_sim(crossEventF) = I                      'write the address in the memory array
                If crossEventF >= 3 Then Quit = True 'Note, the numerical integration is quitted at the point of crossEventF  
            End If
        End If

    End Sub

    Dim NINa As Double
    Public thresholdINa As Double = -15      'pA/pF,      used to remove points during the maximum rate of rise
    Dim weightEVm As Double

    Dim MSElevelA As Double = -0.5     'log(MSE)
    Dim MSELevelB As Double = -1.0    'log(MSE)
    Dim MSELevelC As Double = -1.5      'log(MSE)
    Dim MSELevelD As Double = -10    'log(MSE)
    Dim IndexY As Double                     'copy of the valueY

    Dim adr0L As Double = 8                        'ms, left limit of the interval for measuring MSE
    Dim adr1R As Double = 80                 'ms,   right limit of the interval for measuring MSE
    Dim adr2R As Double = 1025                   'ms,   right limit of the interval for measuring MSE
    Dim adr3R As Double = 1590                'ms     right limit of the interval for measuring MSE
    Dim adr4R As Double = 1080                 'ms       adjustable              'ms       adjustable

    Dim weightmagA As Double = 4         '********** the weight of calculatintg MSE For points around the MDP ****************************
    Dim weightmagB As Double = 3            '3   ********** the weight of calculatintg MSE for points around the MDP ****************************
    Dim weightmagC As Double = 2            '2   ********** the weight of calculatintg MSE for points around the MDP ****************************
    Dim weightmagD As Double = 1            '1   ********** the weight of calculatintg MSE for points around the MDP ****************************
    Dim weightmagE As Double = 0.7            '1   ********** the weight of calculatintg MSE for points around the MDP ****************************
    Dim weightmagF As Double = 0.3

    Dim sumMSE As Double = 0

    Private Sub CalculateMSE_PS()
        '********** Calculate MSE for each sweep of targetVm(I), and simVm(I) ************************
        weightsum = 0                            'counter of calculating SQS values
        NINa = 0                             'counter of the removed points
        sumMSE = 0                       'reset the summator

        '************ Determine weightEFvm(I) ********************************************************
        I = MinX8 / dt
        Do

            If I >= limitL / dt And I < limitR / dt Then                                 'For each point within a AP record 
                sweepTime = I * dt
                '***************** determine the weightmag at this calculating time ***********************
                If rbMMtest.Checked = True Then
                    If sweepTime <= adr1R Then                   'up to the foot of AP  　　　　　 しっかり調整して、IKrsfを調節する。
                        weightEFvm(I) = weightmagA            'B
                    ElseIf sweepTime <= adr2R Then               'SDDの調整によって、その他の電流振幅を決定する。　　　　
                        weightEFvm(I) = weightmagC            'C
                    ElseIf sweepTime <= limitR Then                                                   'AP intervalを調整して、INasfを決定する。
                        weightEFvm(I) = weightmagF            'F
                    End If
                Else
                    weightEFvm(I) = weightmagD         'D = 1
                End If

                If simINa(I) < thresholdINa Then                        'case; .INa is larger than -1 pA/pF , activated during the AP take off  
                    NINa += 1
                Else      '    calculate MSE          simINa(I) > thresholdINa  (= -15 pA)                                                               'case; ips.INa is at the resting level 
                    '************** determine weightEVm according to the address for each Vm point **********************************
                    Difference = targetVm(I) - simVm(I)               'calculate difference beforehand determining the weightEVm

                    If Difference <> 0 Then
                        sumMSE += weightEFvm(I) * Difference * Difference              'Sigma sqare Sum, to calculate the variance 
                        weightsum += weightEFvm(I)
                    End If
                End If
            End If
            PlotResultsVmIm(I)        'plot points of Vm and Im in the graphic memory, yet not display on the screen

            I += 1
        Loop While I < MaxX8 / dt    ''********* at a given sweep of MaxX8 msec is finished ******  plot and print results of PS, and simultaneously MSE is calculated *****************

        '********** determine the final LogMSE ****************************************** 
        If weightsum > 0 Then MSE = sumMSE / weightsum      'average of MSE at a single sweep
        txtMSE.Text = Format(MSE, "##0.#######0")

        If rbMMtest.Checked = True Then
            If MIN > 0 Then
                valueY = Math.Log10(MIN)                      'Math.Log10(MSE)                                         'valueY at a single sweep
                txtMSE.Text = Format(MIN, "##0.#######0")         'Format(MSE, "##0.#######0")
            End If
        Else
            If MSE > 0 Then
                valueY = Math.Log10(MSE)                      'Math.Log10(MSE)                                         'valueY at a single sweep
                txtMSE.Text = Format(MSE, "##0.#######0")         'Format(MSE, "##0.#######0")
            End If
        End If

        txtLogMSE.Text = Format(valueY, "##0.#######0")

        graphMSE.PlotPoint(Eval, valueY, Color.Black, 1, 1) 'plot the average SQS in the graphMSE
        If Eval = 1 Then
            graphBP1MSE.PlotPoint(BP(1), valueY, Color.Red, 2, 2)
            graphBP2MSE.PlotPoint(BP(2), valueY, Color.Red, 2, 2)
            graphBP3MSE.PlotPoint(BP(3), valueY, Color.Red, 2, 2)
            graphBP4MSE.PlotPoint(BP(4), valueY, Color.Red, 2, 2)
            'graphBP5MSE.PlotPoint(BP(5), valueY, Color.Red, 2, 2)
            'graphBP6MSE.PlotPoint(BP(6), valueY, Color.Red, 2, 2)
            oldvalueY = valueY
            Plot_MSEvsBPx()
        End If

        txtdetectINa.Text = Format(NINa, "###0")
        '********************************************************************************

        My.Application.DoEvents()

    End Sub

    Private Sub CalculateMSE_MF()                        'weight = 1
        '********** Calculate MSE for each sweep of targetVm(I), and simVm(I) ************************
        weightsum = 0                            'counter of calculating SQS values
        NINa = 0                             'counter of the removed points
        sumMSE = 0                       'reset the summator

        '************ Determine weightEFvm(I) ********************************************************
        I = MinX8 / dt
        Do

            If I >= limitL / dt And I < limitR / dt Then                                 'For each point within a AP record 
                sweepTime = I * dt
                '***************** determine the weightmag at this calculating time ***********************


                If simINa(I) < thresholdINa Then                        'case; .INa is larger than -1 pA/pF , activated during the AP take off  
                    NINa += 1
                Else      '    calculate MSE          simINa(I) > thresholdINa  (= -15 pA)                                                               'case; ips.INa is at the resting level 
                    '************** determine weightEVm according to the address for each Vm point **********************************
                    Difference = targetVm(I) - simVm(I)               'calculate difference beforehand determining the weightEVm

                    If Difference <> 0 Then
                        sumMSE += Difference * Difference              'Sigma sqare Sum, to calculate the variance 
                        weightsum += 1
                    End If
                End If
            End If
            PlotResultsVmIm(I)        'plot points of Vm and Im in the graphic memory, yet not display on the screen

            I += 1
        Loop While I < MaxX8 / dt    ''********* at a given sweep of MaxX8 msec is finished ******  plot and print results of PS, and simultaneously MSE is calculated *****************

        '********** determine the final LogMSE ****************************************** 
        If weightsum > 0 Then MSE = sumMSE / weightsum      'average of MSE at a single sweep
        txtMSE.Text = Format(MSE, "##0.#######0")

        If MSE > 0 Then
            valueY = Math.Log10(MSE)                                    '  Math.Log10(MIN)                             'valueY at a single sweep
            txtMSE.Text = Format(MSE, "##0.#######0")         'Format(MIN, "##0.#######0")
            txtLogMSE.Text = Format(valueY, "##0.#######0")
        End If
        graphMSE.PlotPoint(Eval, valueY, Color.Black, 1, 1) 'plot the average SQS in the graphMSE
        If Eval = 1 Then
            graphBP1MSE.PlotPoint(BP(1), valueY, Color.Red, 2, 2)
            graphBP2MSE.PlotPoint(BP(2), valueY, Color.Red, 2, 2)
            graphBP3MSE.PlotPoint(BP(3), valueY, Color.Red, 2, 2)
            graphBP4MSE.PlotPoint(BP(4), valueY, Color.Red, 2, 2)
            'graphBP5MSE.PlotPoint(BP(5), valueY, Color.Red, 2, 2)
            'graphBP6MSE.PlotPoint(BP(6), valueY, Color.Red, 2, 2)
            oldvalueY = valueY
            Plot_MSEvsBPx()
        End If

        txtdetectINa.Text = Format(NINa, "###0")
        '********************************************************************************

        My.Application.DoEvents()

    End Sub


    Private Sub StoreIm(ByVal i As Long)
        'keep data points of current systems during the numerical integration
        simIKr(i) = iPS.IKr
        simICaL(i) = iPS.ICaL
        simINa(i) = iPS.INa
        simIK1(i) = iPS.IK1

        simIKs(i) = iPS.IKs
        simIKur(i) = iPS.IKur
        simIKto(i) = iPS.IKto

        simIha(i) = iPS.Iha
        simIK1(i) = iPS.IK1
        simIbNSC(i) = iPS.IbNSC
        simIst(i) = iPS.Ist

        simINaK(i) = iPS.INaK
        simINCX(i) = iPS.INCX

    End Sub

    Private Sub PlotResultsVmIm(ByVal I As Long)
        sweepTime = I * dt
        If chkVmadaptive.Checked Then graphpsVm.PlotPoint(sweepTime, simVm(I), Color.Black, 1, 1)
        If chkVmreference.Checked Then graphpsVm.PlotPoint(sweepTime, targetVm(I), Color.Red, 1, 1)

        If rbMMtest.Checked = True Then graphpsVm.PlotPoint(sweepTime, 10 * weightEFvm(I), Color.Green, 1, 1)

        If chkIKr.Checked Then graphOutwardI.PlotPoint(sweepTime, simIKr(I), Color.Blue, 1, 1)
        If chkIK1.Checked Then graphOutwardI.PlotPoint(sweepTime, simIK1(I), Color.SteelBlue, 1, 1)
        If chkIKur.Checked Then graphOutwardI.PlotPoint(sweepTime, simIKur(I), Color.Chocolate, 1, 1)
        If chkIKto.Checked Then graphOutwardI.PlotPoint(sweepTime, simIKto(I), Color.Black, 1, 1)
        If chkIKs.Checked Then graphOutwardI.PlotPoint(sweepTime, simIKs(I), Color.Violet, 1, 1)
        If chkINaK.Checked Then graphOutwardI.PlotPoint(sweepTime, simINaK(I), Color.Pink, 1, 1)

        If chkICaL.Checked Then graphInwardI.PlotPoint(sweepTime, simICaL(I), Color.Red, 1, 1)
        If chkINa.Checked Then graphInwardI.PlotPoint(sweepTime, simINa(I), Color.Magenta, 1, 1)
        If chkIbNSC.Checked Then graphInwardI.PlotPoint(sweepTime, simIbNSC(I), Color.Black, 1, 1)
        If chkIha.Checked Then graphInwardI.PlotPoint(sweepTime, simIha(I), Color.Green, 1, 1)
        If chkIst.Checked Then graphInwardI.PlotPoint(sweepTime, simIst(I), Color.Brown, 1, 1)
        If chkINCX.Checked Then graphInwardI.PlotPoint(sweepTime, simINCX(I), Color.Red, 1, 1)
    End Sub


    Private Sub printIxmag()               ' cover all possible Ixmags for simplicity
        baseVshift = iPS.baseVshift
        extraVshift = iPS.orgExtraVshift
        txtBaseVshift.Text = Format(iPS.baseVshift, "#0.######")         'print the baseVshift
        txtorgExtraVshift.Text = Format(iPS.orgExtraVshift, "#0.######")       'print the extraVshift

        '*********** data points of Ix are saved in each memory array, to make the program more visible ***************

        txtIKrmag.Text = Format(iPS.IKrmag, "#0.######")
        txtIKurmag.Text = Format(iPS.IKurmag, "#0.######")
        txtICaLmag.Text = Format(iPS.ICaLmag, "#0.######")
        txtVshiftCaL.Text = Format(iPS.orgVshiftCaL, "#0.######")       'original voltage shift can be modified by NP() in patternSearch()
        txtKLmag.Text = Format(iPS.KLmag, "#0.######")                             'Ca-dependency of ICaL inactivation
        txtIbNSCmag.Text = Format(iPS.IbNSCmag, "#0.######")                 'represents a net influence of several backgroun currents
        txtIK1mag.Text = Format(iPS.IK1mag, "#0.######")     'only when the MDP < -75 mV

        '*********** parameters without the pattern Search in the latest version ***************
        txtIKsmag.Text = Format(iPS.IKsmag, "#0.######")       'under the control condition IKs is relatively small and do not influence SDD in HuVEC model 
        txtIhamag.Text = Format(iPS.Ihamag, "#0.######")              'shuld be a target of PatternSearch() when MDP < -70 mV
        txtIKtomag.Text = Format(iPS.IKtomag, "#0.######")           'negligible  influence on the slow diastolic depolarization
        txtINamag.Text = Format(iPS.INamag, "#0.######")             'adjusted according to the maximum rate of rise of AP
        txtVshiftNa.Text = Format(iPS.orgVshiftNa, "#0.######")
        txtNaLratio.Text = Format(iPS.INaLratio, "#0.######")         'to adjust the EAD events    
        txtIstmag.Text = Format(iPS.Pstmag, "#0.######")                  'Ist permeability  relatively small influence on SDD

        '***************  Ion transporters ************************************
        txtINaKmag.Text = Format(iPS.INaKmag, "#0.######")    'adjust INa/Kmag via textBox. see sub iPS.RegulatemaxINaK(), which automatically adjusted outside of the pattern Search to keep [Na]i = 6.4 mM 
        txtINCXmag.Text = Format(iPS.INCXmag, "#0.######")      'fixed, but automatically changes significantly through changes in [Na]i 
        My.Application.DoEvents()
    End Sub

    Private Sub PrintIxscf()
        rKr.Text = Format(iPS.IKrsf, "#0.######")
        rKur.Text = Format(iPS.IKursf, "#0.######")

        rCaL.Text = Format(iPS.ICaLsf, "#0.######")
        rVshiftCaL.Text = Format(iPS.VshiftCaLsf, "#0.######")
        rKL.Text = Format(iPS.KLsf, "#0.######")

        rbNSC.Text = Format(iPS.IbNSCsf, "#0.######")
        rK1.Text = Format(iPS.IK1sf, "#0.######")
        rKs.Text = Format(iPS.IKssf, "#0.######")        'under the control condition IKs is relatively small and do not influence SDD in HuVEC model 
        rha.Text = Format(iPS.Ihasf, "#0.######")                'shuld be a target of PatternSearch() when MDP < -70 mV
        rKto.Text = Format(iPS.IKtosf, "#0.######")           'negligible  influence on the slow diastolic depolarization

        rNa.Text = Format(iPS.INasf, "#0.######")            'adjusted according to the maximum rate of rise of AP
        rVshiftNa.Text = Format(iPS.VshiftNasf, "#0.######")             'adjusted according to the maximum rate of rise of AP
        rst.Text = Format(iPS.Istsf, "#0.######")                 'Ist permeability  relatively small influence on SDD
        'rCaT.Text = Format(iPS.ICaTsf, "#0.######")                'shuld be a target of PatternSearch() when MDP < -70 mV

        txtstdNai.Text = Format(iPS.stdNai, "#0.######")
        txtstdtotCai.Text = Format(iPS.stdtotCai, "#0.######")
        txtcrfNaK.Text = Format(iPS.crf_NaK, "#0.######")
        txtcrfNCX.Text = Format(iPS.crf_NCX, "#0.######")

    End Sub


    Dim valueAAA As Double
    Public GminimumMIN As Double = 0
    Public rstIxmag As Boolean = False
    Public dMSE_step(10) As Double                    'dMSE/ Stp   for each parameter

    Private Sub cbPS_Click(sender As Object, e As EventArgs) Handles cbPS.Click
        cbPS.Visible = False        'hide the command button, 'PatternSerch'
        cbManualFit.Visible = False        'hide the command button, 'Manual Fit'

        chkWriteFile.Checked = False                    'this must be False !!
        '*************************************************************************************************

        '**** prepare Ixmag and Ixsf using the final values in ManualFit() ***************
        calculateNewIxmag()             'calculate iPS.Ixmag = (Ixmag * Ixsf),            and 'print the new Ixmag  in the text box 
        ResetIxsftoUnit()                    'reset Ixsf = 1 and print

        '*************** csvFile write ,  as Ixmag, and Ixsf = 1,   These csvFile is used at the beggining of a new run of PS
        If chkWriteFile.Checked = True Then         'This part is necessary in the sub manualfitting(),  Note; the initial values for the numerical integration should not be change in the Pattern Search
            CSVReadAndWriteE15.CreateNewCSV(CsvFileNameIxpara, iPS, namesIxpara)   'Replace the old CSV file with a new one
            CSVReadAndWriteE15.WriteCSV(CsvFileNameIxpara, I, iPS, 1, namesIxpara)     '膜イオン電流パラメータEuler 変数

            CSVReadAndWriteE15.CreateNewCSV(CsvFileNameIxmag, iPS, namesIxmag)     'Replace the old CSV file with a new one
            CSVReadAndWriteE15.WriteCSV(CsvFileNameIxmag, I, iPS, 1, namesIxmag)      '膜イオン電流振幅調節parameter　Ixmag

            CSVReadAndWriteE15.CreateNewCSV(CsvFileNameIxsf, iPS, namesIxsf)              'initial set of Ixsf, totCa
            CSVReadAndWriteE15.WriteCSV(CsvFileNameIxsf, I, iPS, 1, namesIxsf)              'initial set of Ixsf, totCa
        End If

        '********************  Prepare summator, counter etc. **********************************************
        EvalMax = MaxX3
        txtEvalMax.Text = Format(EvalMax, "###0.#")          'limit of the cycle of Evaluation (givenEquation)   usually 500 is enough     

        Precision = CDbl(txtPrecision.Text)          'adjust the difference (SQS - MIN)     

        'adr2R = Double.Parse(txtWeight.Text)
        'adr3R = Double.Parse(txtRR3R.Text)

        'txtScf2.Text = Format(scf2, "#0.#####")           '  This is used only for the Random2 

        '******************************* Calculation ***********************************
        Sum_BP1 = 0          'clear the summator used in calculating average of repetitive PC cycles
        Sum_BP2 = 0
        Sum_BP3 = 0
        Sum_BP4 = 0
        Sum_BP5 = 0
        'Sum_BP6 = 0

        'oldMSE = 100   '' a maximum magnitude


        '************** Counter ***************************************

        NSweep = 0         'count the number of sweeps
        CountS = 0          'reset the counter for the Sensitivity
        PScycle = 1          'reset the PS cycle counter
        Do                        ' Do Loop for individual PS cycle

            '******** initialize counter, Ixmag, Ixsf etc. for each run of Pattern Search cycle ***************
            'ClearSumSQS()       'initial values of counter, oldSQS, oldvalueY, etc
            PrepareGraphPSVm()


            '******* recover the parameter sets, which were determined in the Manual fitting results ********************
            '*******        Note;  Ixmag is modified when the deeperInsight is used using the chkDeeperInsight *****************
            '*********** Initial set of BP() is given here ********************************************

            CSVReadAndWriteE15.ReadCSV(CsvFileNameIxpara, iPS, namesIxpara)    'initial set of model variables and parameter
            CSVReadAndWriteE15.ReadCSV(CsvFileNameIxmag, iPS, namesIxmag)      'initial set of Ixmag
            CSVReadAndWriteE15.ReadCSV(CsvFileNameIxsf, iPS, namesIxsf)              'initial set of Ixsf, totCa

            calculateNewIxmag()                     'print Ixmag * Ixsf in the text box NIxmag
            printIxmag()                                    'print new Ixmag
            ResetIxsftoUnit()                            'reset Ixsf = 1 and print

            '*************** randomize the model parameters to obtain a set of parameter independent of the initial values ********
            RandomizeInitialIxsf()            'Ixsf (= BP(x)) of selected Ix is  modified for the EMrandomtest using the scf　Ixsfをランダム化する

            For I As Integer = 1 To 10    'scaling factors
                NP(I) = BP(I)
                iPS.NP(I) = NP(I)                     'copy NP(I) as the model parameter
                Stp(I) = 1 / 100                    '75  '20 100   step size = (initial Value) / 100 ,   adjustable according to the project
                '******** CrtStp　はMannualFit()で決定される。*************************
                'CrtStp(I) = 8 / 20000          ' only when the DeeperInsighrt is used. critical step size to quit the optimization = (initial Value) / 1000   ,adjustable
                'CrtStp(I) = 2 / 20000          ' regular CrtStp        critical step size to quit the optimization = (initial Value) / 1000   ,adjustable
                RedFct(I) = 1 / 4                 '5, 4       'reduce the step size   , adjustable           
            Next I

            '********************* setting initial conditions for the PatternSearch() *****************************************
            MSE = 1000
            LogSQS = 3       'temporarily the value is unrealistic large value　is used for determining a minimum SQS
            iPS.LogMIN = LogSQS      'temporarily the value is unrealistic large value　is used for determining a minimum SQS

            Eval = 0                                    'counter of calculating 'givenEquation' are set at zero
            oldEval = 0
            oldvalueY = 3      'initial Log(SQS) is reset to 3, safely beyond the range of graph

            txtStp.Text = Format(Stp(1), "#0.#####")   'step size of NP points

            CountR2 = 0
            For I As Integer = 0 To 5       'initial setting of the oldMIN()
                oldMIN(I) = 1000
            Next


            countPlot = 0       'count the number of using Random2

            countcrossingLevelB = 0      'counter of the MSE, lower than the crtDepthMSEB
            txtCLB.Text = Format(countcrossingLevelB, "##0.")

            For I As Integer = MinX3 To MaxX3   ' weight for all through a single sweep
                weightEFvm(I) = 0
            Next I

            PatternSearch()       '>>>>>>>>>>>>>>> the PatternSearch cycle >>>>>>>>>>>>>>>>>>>>>>>>>>>>>        

            oldMIN(CountR2) = MIN        'keep the last MIN     Because of executing Randomize() for several times, there are several MIN
            MIN = oldMIN(0)                    'now take the minimum MIN among the oldMIN()
            For I As Integer = 1 To CountR2                                               'determine the final MIN
                If oldMIN(I) < oldMIN(I - 1) Then MIN = oldMIN(I)
            Next

            '******************** procedure after the Pattern Search ****************************
            iPS.Eval = Eval             'just to save in the csvFile using the Graph2018　　csvFileに書くためモデルの変数にコピー
            iPS.CR2 = CountR2      'to save in the csvFile

            PlotExit_MSEvsBPx()                     'Plot SQSvsBPx graph using the final MIN

            '******** keep the last BPSQS relation in the csvFile *************
            CSVReadAndWriteE15.WriteCSV(CsvFileNameBPSQS, N_BPSQS, iPS, 1, namesBPSQS)    'Disk write>>>>>>>>
            My.Application.DoEvents()


            N_BPSQS += 1         'proceed with the sequential number of BP points
            txtNBPSQS.Text = Format(N_BPSQS, "##0")   'indicate in the text box

            txtPScycle.Text = Format(PScycle, "##0")
            iPS.PScycle = PScycle        'for the csvFileWrite ***** PSのサイクルNo。

            PScycle += 1                  'counter of the PatternSearch()
        Loop While PScycle < 200               'max is 200 , can be modified    If the number of paralell computation = 10,  total pscycle = 2000

        cbPlot.Visible = True
        cbPS.Visible = True
        cbManualFit.Visible = True

    End Sub

    Private Sub PrepareGraphPSVm()
        txtEval.ForeColor = Color.Black
        graphMSE.Clear()                                                               'prepare the graphSQS at the beginning of each PS series
        graphMSE.DrawAxis(IntervalX3, IntervalY2, "", 8)       'prepare the graphSQS
        graphMSE.PlotLine(MinX3, crtMSEdepthA, MaxX3, crtMSEdepthA, Color.Chocolate, 1)             'MSEdepthA 
        graphMSE.PlotLine(MinX3, crtMSEdepthB, MaxX3, crtMSEdepthB, Color.Magenta, 1)             'MSEdepthB

        graphMSE.PlotLine(MinX3, MSElevelA, MaxX3, MSElevelA, Color.Magenta, 1)             'zero hilizontal level of SQS = 1 mV^2
        graphMSE.PlotLine(MinX3, MSELevelB, MaxX3, MSELevelB, Color.Magenta, 1)             'zero hilizontal level of SQS = 1 mV^2

        If rbMMtest.Checked = True Then
            graphpsVm.PlotLine(limitL, MinY1, limitL, MaxY1, Color.Green, 1)
            graphpsVm.PlotLine(limitR, MinY1, limitR, MaxY1, Color.Green, 1)

            'graphpsVm.PlotLine(adr0L, MinY1, adr0L, MaxY1, Color.Blue, 1)
            'graphpsVm.PlotLine(adr1R, MinY1, adr1R, MaxY1, Color.Violet, 1)
            'graphpsVm.PlotLine(adr2R, MinY1, adr2R, MaxY1, Color.Violet, 1)
            'graphpsVm.PlotLine(adr3R, MinY1, adr3R, MaxY1, Color.Violet, 1)
        End If


        graphSensitivity.Clear()
        graphSensitivity.DrawAxis(IntervalX21, IntervalY21, "", 8)

    End Sub


    Dim gRND As Random = New Random()      'ランダム関数を使用する宣言
    Dim valueRND As Double
    Dim valueZ As Double

    Public scf1 As Double     '= 0.1                      'the % randomization = scf * 50  %
    Public scf2 As Double      '= 0.01

    Private Sub RandomizeInitialIxsf()

        'txtScf1.Text = Format(scf1, "#0.###")                  ' obtain from txtScf        '0.1,  5% ,  0.2  10%,  0.3  15%,  0.4  20%,  0.5  25%
        'txtScf2.Text = Format(scf2, "#0.###")                   ' obtain from txtScf        '0.1,  5% ,  0.2  10%,  0.3  15%,  0.4  20%,  0.5  25%

        scf1 = Double.Parse(txtScf1.Text)
        scf2 = Double.Parse(txtScf2.Text)


        valueRND = gRND.NextDouble()          'randomize       IKr
        iPS.initialIKrsf = 1 + (valueRND - 0.5) * scf1
        iPS.IKrsf = iPS.initialIKrsf
        rKr.Text = Format(iPS.IKrsf, "#0.#####")                          'テキストボックスに書き込む

        valueRND = gRND.NextDouble()          'randomize         ICaL
        iPS.initialICaLsf = 1 + (valueRND - 0.5) * scf1
        iPS.ICaLsf = iPS.initialICaLsf
        rCaL.Text = Format(iPS.ICaLsf, "#0.#####")                       'テキストボックスに書き込む

        'valueRND = gRND.NextDouble()          'randomize               IbNSC
        'iPS.initialIbNSCsf = 1 + (valueRND - 0.5) * scf1
        'iPS.IbNSCsf = iPS.initialIbNSCsf
        'rbNSC.Text = Format(iPS.IbNSCsf, "#0.#####")                       'テキストボックスに書き込む 

        valueRND = gRND.NextDouble()          'randomize               INa
        iPS.initialINasf = 1 + (valueRND - 0.5) * scf1
        iPS.INasf = iPS.initialINasf
        rNa.Text = Format(iPS.INasf, "#0.###")


        valueRND = gRND.NextDouble()          'randomize            IK1
        iPS.initialIK1sf = 1 + (valueRND - 0.5) * scf1
        iPS.IK1sf = iPS.initialIK1sf
        rK1.Text = Format(iPS.IK1sf, "#0.###")                   'テキストボックスに書き込む 

        'valueRND = gRND.NextDouble()          'randomize               VshiftNa
        'iPS.initialVshiftNasf = 1 + (valueRND - 0.5) * scf1
        'iPS.VshiftNasf = iPS.initialVshiftNasf
        'rVshiftNa.Text = Format(iPS.VshiftNasf, "#0.###")

        'valueRND = gRND.NextDouble()          'randomize               IKur
        'iPS.initialIKursf = 1 + (valueRND - 0.5) * scf1
        'iPS.IKursf = iPS.initialIKursf
        'rKur.Text = Format(iPS.IKursf, "#0.###")

        'valueRND = gRND.NextDouble()          'randomize               IKs
        'iPS.initialIKssf = 1 + (valueRND - 0.5) * scf1
        'iPS.IKssf = iPS.initialIKssf
        'rKs.Text = Format(iPS.IKssf, "#0.###")

        'valueRND = gRND.NextDouble()          'randomize               KL
        'iPS.initialKLsf = 1 + (valueRND - 0.5) * scf1
        'iPS.KLsf = iPS.initialKLsf
        'rKL.Text = Format(iPS.KLsf, "#0.###")


        'valueRND = gRND.NextDouble()          'randomize            Iha
        'iPS.initialIhasf = 0.97699               '1 + (valueRND - 0.5) * scf1
        'iPS.Ihasf = iPS.initialIhasf
        'rst.Text = Format(iPS.Ihasf, "#0.###")                   'テキストボックスに書き込む


        'valueRND = gRND.NextDouble()          'randomize               IKto
        'iPS.initialIKtosf = 1 + (valueRND - 0.5) * scf1
        'iPS.IKtosf = iPS.initialIKtosf
        'rKto.Text = Format(iPS.IKtosf, "#0.###")

        BP(1) = iPS.IKrsf                  'prepare for the line plot in Panel D　それぞれのモデル変数を割り当てる
        BP(2) = iPS.ICaLsf
        BP(3) = iPS.INasf
        BP(4) = iPS.IK1sf
        'BP(5) = iPS.Ihasf
        'BP(6) = iPS.IbNSCsf


        oldBP(1) = BP(1)                 'prepare for the line plot in Panel D, if required　折れ線グラフを書くため、前回の値を記録する
        oldBP(2) = BP(2)     '最初の値
        oldBP(3) = BP(3)
        oldBP(4) = BP(4)
        'oldBP(5) = BP(5)
        'oldBP(6) = BP(6)
        'oldBP(7) = BP(7)


        iPS.BP1 = BP(1)              'for the csvFileWrite ***** the first line of csvFile indicates the initial value
        iPS.BP2 = BP(2)              'csvFileに書き出すために、テクニカルなコード
        iPS.BP3 = BP(3)
        iPS.BP4 = BP(4)
        'iPS.BP5 = BP(5)
        'iPS.BP6 = BP(6)
        'iPS.BP7 = BP(7)

        valueY = MaxY21 + 0.1        '************* plot the initial randomized points at the top of the graph ****************
        graphBP1MSE.PlotPoint(iPS.initialIKrsf, valueY, Color.Blue, 4, 4)            'グラフ上にランダム化されたBPをプロットする。
        graphBP2MSE.PlotPoint(iPS.initialICaLsf, valueY, Color.Red, 4, 4)
        graphBP3MSE.PlotPoint(iPS.initialINasf, valueY, Color.Violet, 4, 4)
        graphBP4MSE.PlotPoint(iPS.initialIK1sf, valueY, Color.SteelBlue, 4, 4)
        'graphBP5MSE.PlotPoint(iPS.initialIhasf, valueY, Color.Lime, 4, 4)
        'graphBP6MSE.PlotPoint(iPS.initialIbNSCsf, valueY, Color.Black, 4, 4)

    End Sub

    Public CountR2 As Double = 0         'counter for the number of randomization R

    Public oldMIN(5) As Double       'MIN just before the randomize2 


    Public Sub Randomize2()  'this is used to randomize the parameters to escape from the stucked state
        'If MIN < minimumMIN Then minimumMIN = MIN     'select the least MSE 
        'txtPreviousMIN.Text = Format(Math.Log10(minimumMIN), "#0.#####")

        I = 0
        Do                                          'ランドム関数を繰り返す　　　無駄かも知れない
            valueRND = gRND.NextDouble()
            I += 1
        Loop While I < 23

        '********** その時々のIxsfの値を変更する。＊＊＊＊＊＊＊＊＊＊＊
        'scf2 = Double.Parse(txtScf2.Text)                  ' obtain from txtScf        '0.1,  5% ,  0.2  10%,  0.3  15%,  0.4  20%,  0.5  25%

        valueRND = gRND.NextDouble()          'randomize       IKr
        iPS.IKrsf = iPS.IKrsf * (1 + (valueRND - 0.5) * scf2)
        rKr.Text = Format(iPS.IKrsf, "#0.#####")

        valueRND = gRND.NextDouble()          'randomize         ICaL
        iPS.ICaLsf = iPS.ICaLsf * (1 + (valueRND - 0.5) * scf2)
        rCaL.Text = Format(iPS.ICaLsf, "#0.#####")

        'valueRND = gRND.NextDouble()          'randomize               IbNSC
        'iPS.IbNSCsf = iPS.IbNSCsf * (1 + (valueRND - 0.5) * scf2)
        'rbNSC.Text = Format(iPS.IbNSCsf, "#0.#####")

        'valueRND = gRND.NextDouble()          'randomize               IKur
        'iPS.IKursf = iPS.IKursf * (1 + (valueRND - 0.5) * scf2)
        'rKur.Text = Format(iPS.IKursf, "#0.###")

        valueRND = gRND.NextDouble()          'randomize               INa  
        iPS.INasf = iPS.INasf * (1 + (valueRND - 0.5) * scf2 / 2)
        rNa.Text = Format(iPS.INasf, "#0.#####")

        valueRND = gRND.NextDouble()          'randomize            IK1
        iPS.IK1sf = iPS.IK1sf * (1 + (valueRND - 0.5) * scf2)
        rK1.Text = Format(iPS.IK1sf, "#0.#####")                   'テキストボックスに書き込む

        'valueRND = gRND.NextDouble()          'randomize               IK1
        'iPS.IK1sf = iPS.IK1sf * (1 + (valueRND - 0.5) * scf2)
        'rK1.Text = Format(iPS.IK1sf, "#0.#####")

        'valueRND = gRND.NextDouble()          'randomize               KL
        'iPS.KLsf = iPS.KLsf * (1 + (valueRND - 0.5) * scf2)
        'rKL.Text = Format(iPS.KLsf, "#0.###")


        'valueRND = gRND.NextDouble()          'randomize               IKs
        'iPS.IKssf = iPS.IKssf * (1 + (valueRND - 0.5) * scf2)
        'rKs.Text = Format(iPS.IKssf, "#0.###")

        'valueRND = gRND.NextDouble()          'randomize            Iha
        'iPS.Ihasf = iPS.Ihasf * (1 + (valueRND - 0.5) * scf2)
        'rha.Text = Format(iPS.Ihasf, "#0.###")                   'テキストボックスに書き込む


        'valueRND = gRND.NextDouble()          'randomize            Ist
        'iPS.Istsf = iPS.Istsf * (1 + (valueRND - 0.5) * scf2)
        'rst.Text = Format(iPS.Istsf, "#0.###")                   'テキストボックスに書き込む

        'valueRND = gRND.NextDouble()          'randomize               IKto
        'iPS.initialIKtosf = 1 + (valueRND - 0.5) * scf2
        'iPS.IKtosf = iPS.initialIKtosf
        'rKto.Text = Format(iPS.IKtosf, "#0.###")


        BP(1) = iPS.IKrsf                  'prepare for the line plot in Panel D　それぞれのモデル変数を割り当てる
        BP(2) = iPS.ICaLsf
        BP(3) = iPS.INasf
        BP(4) = iPS.IK1sf
        'BP(5) = iPS.Ihasf
        'BP(6) = iPS.IbNSCsf

        oldBP(1) = BP(1)                 'prepare for the line plot in Panel D, if required
        oldBP(2) = BP(2)
        oldBP(3) = BP(3)
        oldBP(4) = BP(4)
        'oldBP(5) = BP(5)
        'oldBP(6) = BP(6)
        'oldBP(6) = BP(7)

        NP(1) = BP(1)              'for the csvFileWrite ***** the first line of csvFile indicates the initial value
        NP(2) = BP(2)
        NP(3) = BP(3)
        NP(4) = BP(4)
        'NP(5) = BP(5)
        'NP(6) = BP(6)
        'iPS.BP6 = BP(7)

        graphMSE.PlotLine(Eval, MinY21, Eval, MaxY21, Color.Magenta, 1)   'random2使用のタイミングをグラフに表示する。

        givenEquation()　　　　　　'新たなMSEをNP(n)を使って計算する。

        MIN = MSE
        '********** plot results *******************************
        Plot_MSEvsBPxRandom2()           'plot the MSE, use the conventional method

    End Sub


    Public Sub Randomize3()  'this is used to randomize the parameters to escape from the stucked state
        'If MIN < minimumMIN Then minimumMIN = MIN     'select the least MSE 
        'txtPreviousMIN.Text = Format(Math.Log10(minimumMIN), "#0.#####")

        'I = 0
        'Do                                          'ランドム関数を繰り返す　　　無駄かも知れない
        '    valueRND = gRND.NextDouble()
        '    I += 1
        'Loop While I < 23

        '********** その時々のIxsfの値を変更する。＊＊＊＊＊＊＊＊＊＊＊
        'scf2 = Double.Parse(txtScf2.Text)                  ' obtain from txtScf        '0.1,  5% ,  0.2  10%,  0.3  15%,  0.4  20%,  0.5  25%

        valueRND = gRND.NextDouble()          'randomize       IKr
        iPS.IKrsf = iPS.IKrsf * (1 + (valueRND - 0.5) * scf2)
        rKr.Text = Format(iPS.IKrsf, "#0.#####")

        valueRND = gRND.NextDouble()          'randomize         ICaL
        iPS.ICaLsf = iPS.ICaLsf * (1 + (valueRND - 0.5) * scf2)
        rCaL.Text = Format(iPS.ICaLsf, "#0.#####")

        valueRND = gRND.NextDouble()          'randomize               IbNSC
        iPS.IbNSCsf = iPS.IbNSCsf * (1 + (valueRND - 0.5) * scf2)
        rbNSC.Text = Format(iPS.IbNSCsf, "#0.#####")

        valueRND = gRND.NextDouble()          'randomize               INa
        iPS.INasf = iPS.INasf * (1 + (valueRND - 0.5) * scf2)
        rNa.Text = Format(iPS.INasf, "#0.#####")

        valueRND = gRND.NextDouble()          'randomize            IK1
        iPS.IK1sf = iPS.IK1sf * (1 + (valueRND - 0.5) * scf2)
        rK1.Text = Format(iPS.IK1sf, "#0.#####")                   'テキストボックスに書き込む

        'valueRND = gRND.NextDouble()          'randomize               IK1
        'iPS.IK1sf = iPS.IK1sf * (1 + (valueRND - 0.5) * scf2)
        'rK1.Text = Format(iPS.IK1sf, "#0.#####")

        'valueRND = gRND.NextDouble()          'randomize               KL
        'iPS.KLsf = iPS.KLsf * (1 + (valueRND - 0.5) * scf2)
        'rKL.Text = Format(iPS.KLsf, "#0.###")

        'valueRND = gRND.NextDouble()          'randomize               IKur
        'iPS.IKursf = iPS.IKursf * (1 + (valueRND - 0.5) * scf2)
        'rKur.Text = Format(iPS.IKursf, "#0.###")

        'valueRND = gRND.NextDouble()          'randomize               IKs
        'iPS.IKssf = iPS.IKssf * (1 + (valueRND - 0.5) * scf2)
        'rKs.Text = Format(iPS.IKssf, "#0.###")

        'valueRND = gRND.NextDouble()          'randomize            Iha
        'iPS.Ihasf = iPS.Ihasf * (1 + (valueRND - 0.5) * scf2)
        'rha.Text = Format(iPS.Ihasf, "#0.###")                   'テキストボックスに書き込む


        'valueRND = gRND.NextDouble()          'randomize            Ist
        'iPS.Istsf = iPS.Istsf * (1 + (valueRND - 0.5) * scf2)
        'rst.Text = Format(iPS.Istsf, "#0.###")                   'テキストボックスに書き込む

        'valueRND = gRND.NextDouble()          'randomize               IKto
        'iPS.initialIKtosf = 1 + (valueRND - 0.5) * scf2
        'iPS.IKtosf = iPS.initialIKtosf
        'rKto.Text = Format(iPS.IKtosf, "#0.###")


        BP(1) = iPS.IKrsf                  'prepare initial BP(x) for PS
        BP(2) = iPS.ICaLsf
        BP(3) = iPS.IbNSCsf
        BP(4) = iPS.IKursf
        BP(5) = iPS.INasf
        'BP(6) = iPS.IKursf

        oldBP(1) = BP(1)                 'prepare for the line plot in Panel D, if required
        oldBP(2) = BP(2)
        oldBP(3) = BP(3)
        oldBP(4) = BP(4)
        oldBP(5) = BP(5)
        'oldBP(6) = BP(6)
        'oldBP(6) = BP(7)

        NP(1) = BP(1)              'for the csvFileWrite ***** the first line of csvFile indicates the initial value
        NP(2) = BP(2)
        NP(3) = BP(3)
        NP(4) = BP(4)
        NP(5) = BP(5)
        'NP(6) = BP(6)
        'iPS.BP6 = BP(7)

        'graphMSE.PlotLine(Eval, MinY21, Eval, MaxY21, Color.Magenta, 1)   'random2使用のタイミングをグラフに表示する。

        givenEquation()　　　　　　'新たなMSEをNP(n)を使って計算する。

        MIN = MSE
        valueY = Math.Log10(MIN)
        graphMSE.PlotPoint(Eval, valueY, Color.Black, 1, 1)   'random2使用のタイミングをグラフに表示する。


        '********** plot results *******************************
        'Plot_MSEvsBPxRandom2()           'plot the MSE, use the conventional method

    End Sub


    'Public rowsimVm(ArraySize) As Double         'Vm record in the pattern Search
    Dim crtMSEdepthA As Double              'threshold for the randomize2
    Dim countR2max As Double = 3
    Dim crtMSEdepthB As Double = -4.0             'threshold for the termination


    Public weightsum As Double = 0                              'number of data points used in calculating SQS
    Dim Nfitting As Integer = 0                                           'number of fitting
    Dim NumPara As Integer = 4   '7          'number of parameters to be determined

    Dim LogMIN As Double
    Dim crtLogMSE As Double = 10
    Dim R3Eval As Double = 0
    Dim countcrossingLevelB As Double = 0
    Dim CountStoMIN As Double = 0
    Dim countRepetition As Double = 0       'Counter of the repetitive global MIN

    Dim repeatRandom As Double


    Private Sub PatternSearch()               '*************** Start of the Pattern Search *********************
        My.Application.DoEvents()                           'quit when any event happens during simulation

        Dim I As Integer
        'PAT = 1
        givenEquation()                                 'calculate SQS
        MIN = MSE                       'minimum of SQS

D590:

        EPS = 1.0E+37                  ' ExPanded Step ? is prepared at the maximum double  This is useful to get a deeper level of SQS
        For I = 1 To NumPara
            NP(I) = BP(I)                                  'At the begining of the Pattern Search,  NP() is copied from the base point
            'If CrtStp(I) < EPS Then EPS = CrtStp(I) 'renew the critical step by the current one
        Next I

D680:
        EXPLORE()                  ' Determine a new NP points
        'graphFails.PlotLine(Eval, 0, Eval, Fails, Color.Black, 3)    'is not used in the present Analysis 

        If Fails <> NumPara Then GoTo D740                          'examine if examinations are finished for all parameters 

        '(option 1 MSE switch to Dexit) ********************************************************************
        valueY = Math.Log10(MIN)                                'define valueY as soon as possible after the given equation
        If valueY < crtMSEdepthB Then                        'LevelB = -4    
            countcrossingLevelB += 1                               'Level B defines the satisfactory minimum level of MSE
            txtCLB.Text = Format(countcrossingLevelB, "##0.")

            If countcrossingLevelB >= 2 Then GoTo Dexit           '  'Small enough to Exit
            My.Application.DoEvents()
        End If

        '(option 2 Stp switch to the next round  ************* Reset the next round after decreasing the Stp(1)   ********************************
        If Math.Abs(Stp(1)) > CrtStp(1) Then      ' if any Stp() > CrtStp() then
            For I = 1 To NumPara                            'usual procedure
                Stp(I) = Stp(I) * RedFct(I)                       'decrease the step size by a factor RedFct 
            Next I
            GoTo D680
        End If                                    'Case of Math.Abs(Stp(M)) =< CrtStp(M),  decide if it is appropriate to quit the parameter optimization


        '(option 3 the second MSE switch to Dexit) ************************************************************************
        If valueY < crtMSEdepthA Then GoTo Dexit            'LevelA = -2     limit of excuting randomize2()

        '(option 4 the R2 swich to Dexit) ************************************************************************
        If Eval > (MaxX3 - 200) Then GoTo Dexit                'No time to execute random2 because the time shortage for the random 2 completion

        ' ***** execute the random 2 *************************************************
        'If CountR2 < countR2max Then
        '    oldMIN(CountR2) = MIN                      'keep the last MIN determined before executing the randomize2() in the memory array
        '    txtR2.Text = Format(CountR2 + 1, "##0")   'print the number of repetition
        '    CountR2 += 1

        '    Randomize2()

        '    For I = 1 To NumPara            'option
        '        Stp(I) = 0.000625                 'retry using this larger Stp()
        '    Next I
        '    'Plot_MSEvsBPx()                     'Plot and Save the new BP
        '    GoTo D680                                'continue the Pattern Search
        'Else
        GoTo Dexit     '(option 5 to Dexit)  of unknown reason *****************************************
        'End If


        '*************  The start of Block 2 ***************************************************************************
D740:
        For I = 1 To NumPara
            Advance(I) = NP(I) - BP(I)              'Advance of each NP(I) = movement of NP(I) from BP(I) 
        Next I

D750:   'renew  Base Points
        For I = 1 To NumPara
            BP(I) = NP(I)                                  'renew the base point BP() by a new point NP(), irrespective of improvement or failure
        Next I

        Plot_MSEvsBPx()                     '*************  Plot and Save the new BP  *******************************************

        For I = 1 To NumPara
            NP(I) = BP(I) + Advance(I)
            If Advance(I) * Stp(I) <= 0 Then
                Stp(I) = -Stp(I)                             'invert the direction for search
            End If
        Next I

        '********************* limit of the evaluation cycle********************
        iPS.Eval = Eval

        If Eval > EvalMax Then GoTo Dexit                          'Eval switch to Dexit 
        'If falseEval > MaxX3 / 5 Then GoTo Dexit
        '********************  **************************************************

        MinSto = MIN                                      'keep the minimum SQS 

        givenEquation()               'calculate present SQS using the latest set of parameters,    for sure?
        MIN = MSE                      '

        EXPLORE()                                                                'attached at #D750

        If MIN <= MinSto Then
            For I = 1 To NumPara
                Advance(I) = NP(I) - BP(I)
            Next I

            For I = 1 To NumPara
                If Math.Abs(Advance(I)) > EPS Then GoTo D750
            Next I
        End If

        'txtMSE.Text = Format(Math.Log10(MinSto), "##.0######")      'print in the text Box

        MIN = MinSto

        '*************** New lines to calculate the global minimum of MSE,  220214 *************
        'If MIN < (GminimumMIN - Precision) Then     'small enough to decide
        '    GminimumMIN = MIN
        '    countRepetition = 0        'reset the counter of the continuous minimum level
        '    txtPreviousMIN.Text = Format(GminimumMIN, "#0.########")
        'Else
        '    countRepetition += 1       'proceed with the counter
        '    If countRepetition > 200 Then GoTo Dexit           'quit a given run of the Pattern Search
        'End If
        '*****************************************************************************

        'If chkRand2.Checked = True Then
        '    chkRand2.Checked = False
        '    If CountR2 < countR2max Then
        '        oldMIN(CountR2) = MIN                      'keep the last MIN determined before executing the randomize2() in the memory array
        '        txtR2.Text = Format(CountR2 + 1, "##0")   'print the number of repetition
        '        CountR2 += 1
        '    End If
        '    Randomize2()
        '    GoTo D680
        'End If


        '*****************************************************************************
        GoTo D590

Dexit:


    End Sub

    Dim Precision As Double = 0
    Dim M As Integer
    Dim cMSE As Double
    Dim dMIN(10) As Double
    Dim OlddMin(10) As Double
    Dim CountS As Long = 0
    Dim caseMSE(10) As Boolean
    Dim MINtemp(10) As Double

    Private Sub EXPLORE()                          'あるパラメータを＋－ステップ分移動してみて改善が見られるか否か調べる。
        'counter of converting the step sign, 1, or 2
        Fails = 0                                             'reset the counter of failure to obtain a decrease in SQS  
        'Dim J As Integer

        '************************  New option,   Case using RandomizeSequence()
        RandomizeSequence(previousNP)        'determine rdNP(1) ~ (4) is determined according to the preious No. of parameter
        rdNP(5) = 5                                           'outside of Randamization
        rdNP(6) = 6                                           'outside of Randamization
        rdNP(7) = 7                                           'outside of Randamization


        For I As Integer = 1 To NumPara        'check if a step in each parameter make an improvement
            M = rdNP(I)                              'No. of a parameter    Do not use L, which is used in the getsimVmMFPS()
            'L = rdNP(I)

            txtNP.Text = Format(M, "#0")          'print L to indicate the present NP number
            My.Application.DoEvents()             'indicate immediately the randamized new sequence

            ' ************************ Case not using RandomizeSequence
            'For M As Integer = 1 To NumPara     'No. of radomizeSequence         Be careful!  Next L　で終わること
            '********************************************************

            HOME = NP(M)                          ' A temporaly HOME is defined for a given parameter NP()  
D980:
            NP(M) = HOME + Stp(M)           'new point is determined as HOME + stp size
            givenEquation()               'GetsimVm()実行して、定常状態を得る。calculate the variance for the new point to decide it is improved or not
            MSEp = MSE　　　　　　'このNP点でのMSEをMSEｐとして取得

            NP(M) = HOME - Stp(M)           'new point is determined as HOME + stp size
            givenEquation()                'GetsimVm()実行して、定常状態を得るcalculate the variance for the new point to decide it is improved or not
            MSEn = MSE　　　　　　'このNP点でのMSEをMSEｎとして取得
            '*************************************************************
            If MSEp < MSEn Then                   'the positive step is better significantly
                caseMSE(M) = True

                If (MSEp - MIN) < Precision Then　　'precisionを与えて、MINと比較して、MSEを有意に減少する事ができた場合。
                    'CalculateSensitivity()
                    MIN = MSEp
                    NP(M) = HOME + Stp(M)　 'new NP()
                Else                                          'no improvement of MSE
                    NP(M) = HOME
                    Fails = Fails + 1
                End If

            Else              'the negative step is better siginificantly
                caseMSE(M) = False
                '
                If (MSEn - MIN) < Precision Then　　'precisionを与えて、MINと比較して、MSEを減少する事ができた場合。
                    'CalculateSensitivity()
                    MIN = MSEn
                    NP(M) = HOME - Stp(M)　 'new NP()
                Else                                           'no improvement of MSE
                    NP(M) = HOME
                    Fails = Fails + 1
                End If
            End If

            CalculateSensitivity()

        Next I
        '             Next M                                           'move to the next parameter 

    End Sub

    Dim preEval As Double


    Private Sub CalculateSensitivity()
        dMSE_step(M) = (MIN - MINtemp(M)) / Math.Abs(Stp(M))       'Slopeを計算する   based on the method of 'sensitivity analysis'
        If dMSE_step(M) > 0 Then
            dMIN(M) = dMSE_step(M) / 100   'for visibility, divide by 1000

            Dim Yvalue As Double = Math.Log10(dMIN(M))
            If M = 1 Then
                graphSensitivity.PlotPoint(Eval, Yvalue, Color.Black, 3, 3)
                txtSbp1.Text = Format(dMIN(1), "#0.######0")
            ElseIf M = 2 Then
                graphSensitivity.PlotPoint(Eval, Yvalue, Color.Gray, 3, 3)
                txtSbp2.Text = Format(dMIN(2), "#0.######0")
            ElseIf M = 3 Then
                graphSensitivity.PlotPoint(Eval, Yvalue, Color.Fuchsia, 3, 3)
                txtSbp3.Text = Format(dMIN(3), "#0.######0")
            ElseIf M = 4 Then
                graphSensitivity.PlotPoint(Eval, Yvalue, Color.Lime, 3, 3)
                txtSbp4.Text = Format(dMIN(4), "#0.######0")
            ElseIf M = 5 Then
                graphSensitivity.PlotPoint(Eval, Yvalue, Color.Red, 3, 3)
                txtSbp5.Text = Format(dMIN(5), "#0.######0")
                'ElseIf M = 6 Then
                '    graphSensitivity.PlotPoint(Eval, Yvalue, Color.Blue, 3, 3)
                '    txtSbp6.Text = Format(dMIN(6), "#0.######0")
            End If

        ElseIf dMSE_step(M) <= 0 Then
            If M = 1 Then txtSbp1.Text = "-----"     'Format("-")
            If M = 2 Then txtSbp2.Text = "-----"     'Format("-")
            If M = 3 Then txtSbp3.Text = "-----"     'Format("-")
            If M = 4 Then txtSbp4.Text = "-----"     'Format("-")
            If M = 5 Then txtSbp5.Text = "-----"     'Format("-")
            If M = 6 Then txtSbp6.Text = "-----"     'Format("-")
        End If


        CountS += 1     'proceed with the counter of the plot

    End Sub


    'Private Sub CalculateSensitivity()

    '    dMSE_step(M) = Math.Abs((MIN - MSE) / Stp(M))       'Slopeを計算する   based on the method of 'sensitivity analysis'

    '    If dMSE_step(M) > 0 Then

    '        dMIN(M) = dMSE_step(M) / 1000     'for visibility, divide by 1000
    '        Dim Yvalue As Double = Math.Log10(dMIN(M))
    '        If M = 1 Then graphSensitivity.PlotPoint(Eval, Yvalue, Color.Black, 3, 3)
    '        If M = 2 Then graphSensitivity.PlotPoint(Eval, Yvalue, Color.Gray, 3, 3)
    '        If M = 3 Then graphSensitivity.PlotPoint(Eval, Yvalue, Color.Fuchsia, 3, 3)
    '        If M = 4 Then graphSensitivity.PlotPoint(Eval, Yvalue, Color.Lime, 3, 3)
    '        If M = 5 Then graphSensitivity.PlotPoint(Eval, Yvalue, Color.Red, 3, 3)
    '        If M = 6 Then graphSensitivity.PlotPoint(Eval, Yvalue, Color.Blue, 3, 3)

    '        txtSbp1.Text = Format(dMIN(1), "#0.######0")
    '        txtSbp2.Text = Format(dMIN(2), "#0.######0")
    '        txtSbp3.Text = Format(dMIN(3), "#0.######0")
    '        txtSbp4.Text = Format(dMIN(4), "#0.######0")
    '        txtSbp5.Text = Format(dMIN(5), "#0.######0")
    '        txtSbp6.Text = Format(dMIN(6), "#0.######0")

    '    ElseIf dMSE_step(M) <= 0 Then
    '        txtSbp1.Text = Format(dMIN(1), "-")
    '        txtSbp2.Text = Format(dMIN(2), "-")
    '        txtSbp3.Text = Format(dMIN(3), "-")
    '        txtSbp4.Text = Format(dMIN(4), "-")
    '        txtSbp5.Text = Format(dMIN(5), "-")
    '        txtSbp6.Text = Format(dMIN(6), "-")
    '    End If

    '    CountS += 1     'proceed with the counter of the plot

    'End Sub


    Public limitL As Double
    Public limitR As Double
    Public RNaKNCX As Boolean

    Private Sub givenEquation()       'calculate sum of variance 

        '*********** Determine which NP(x), which is Ixsf,  is modified through the pattern Search *************   
        iPS.IKrsf = NP(1)               'Blue
        iPS.ICaLsf = NP(2)             'Red
        iPS.INasf = NP(3)               'pink
        iPS.IK1sf = NP(4)             'SteelBlue 
        'iPS.Ihasf = NP(5)               'Green
        'iPS.IbNSCsf = NP(6)             'black 

        'iPS.Ihasf = NP(4)            'green

        'iPS.IKursf = NP(4)         'Pink
        'iPS.Istsf = NP(6)           'chocolate

        'iPS.IKtosf = NP(6)            'black

        'iPS.Istsf = NP(5)                'chocolate

        If NSweep = 0 Then
            txtI1.Text = "IKr"
            txtI2.Text = "ICaL"
            txtI3.Text = "INa"
            txtI4.Text = "IK1"
            'txtI5.Text = "Iha"
            'txtI6.Text = "IbNSC"
            'txtI6.Text = 
            'txtI6.Text = "IKur"

            'txtI4.Text = "IKto"
        End If

        '************* Calculate Vm,a *******************************************
        RNaKNCX = True                         'regulate NaK, and NCX activity,     crfNaK, crfNCX
        GetSimVm(RNaKNCX, maxRepeat)                         'get simVm() by calculating the model after  a parameter modification 

        '*********** mean square difference (mV2) between targetVm() and simVm() *********************
        CalculateMSE_PS()

        '************ plot Results ********************************************************************
        PrintScf()                    'print the last scfs just to monitor Ixsf and Ixsf * Ixmag

        'If MSE < 10 Then
        Eval += 1
            txtEval.Text = Format(Eval, "###0")    'print the eval
        'End If

        '********************************************************************************
        My.Application.DoEvents()

    End Sub

    Public Sub calculateNewIxmag()              'Ixmag is given by the product of (Ixmag x Ixsf)
        '******* Calcurate new Ixmag *******************
        iPS.IKrmag = iPS.IKrmag * iPS.IKrsf
        iPS.IK1mag = iPS.IK1mag * iPS.IK1sf
        iPS.ICaLmag = iPS.ICaLmag * iPS.ICaLsf
        iPS.KLmag = iPS.KLmag * iPS.KLsf
        iPS.orgVshiftCaL = iPS.orgVshiftCaL * iPS.VshiftCaLsf
        iPS.IbNSCmag = iPS.IbNSCmag * iPS.IbNSCsf
        iPS.Ihamag = iPS.Ihamag * iPS.Ihasf
        iPS.IKsmag = iPS.IKsmag * iPS.IKssf
        iPS.IKtomag = iPS.IKtomag * iPS.IKtosf
        iPS.IKurmag = iPS.IKurmag * iPS.IKursf
        iPS.INamag = iPS.INamag * iPS.INasf
        iPS.orgVshiftNa = iPS.orgVshiftNa * iPS.VshiftNasf
        iPS.Pstmag = iPS.Pstmag * iPS.Istsf

        '************ print new Ixmag **********************
        NIKrmag.Text = Format(iPS.IKrmag, "#0.######")   ' only to monitor the change in 'Ixmag'
        NIK1mag.Text = Format(iPS.IK1mag, "#0.######")
        NICaLmag.Text = Format(iPS.ICaLmag, "#0.######")
        NKLmag.Text = Format(iPS.KLmag, "#0.######")
        NVshiftCaL.Text = Format(iPS.orgVshiftCaL, "#0.######")
        NIbNSCmag.Text = Format(iPS.IbNSCmag, "#0.######")
        NIhamag.Text = Format(iPS.Ihamag, "#0.######")
        NIKsmag.Text = Format(iPS.IKsmag, "#0.######")
        NIKtomag.Text = Format(iPS.IKtomag, "#0.######")
        NIKurmag.Text = Format(iPS.IKurmag, "#0.######")
        NINamag.Text = Format(iPS.INamag, "#0.######")
        NVshiftNa.Text = Format(iPS.orgVshiftNa, "#0.######")
        NIstmag.Text = Format(iPS.Pstmag, "#0.######")
    End Sub


    Private Sub ResetIxsftoUnit()
        '******* replace Ixsf *******************
        iPS.IKrsf = 1
        iPS.IK1sf = 1
        iPS.ICaLsf = 1
        iPS.KLsf = 1
        iPS.VshiftCaLsf = 1
        iPS.IbNSCsf = 1
        iPS.Ihasf = 1
        iPS.IKssf = 1
        iPS.IKtosf = 1
        iPS.IKursf = 1
        iPS.INasf = 1
        iPS.VshiftNasf = 1
        iPS.Istsf = 1

        '************ print new Ixsf **********************
        rKr.Text = Format(iPS.IKrsf, "#0.######")
        rK1.Text = Format(iPS.IK1sf, "#0.######")
        rCaL.Text = Format(iPS.ICaLsf, "#0.#####")
        rKL.Text = Format(iPS.KLsf, "#0.######")
        rVshiftCaL.Text = Format(iPS.VshiftCaLsf, "#0.######")
        rbNSC.Text = Format(iPS.IbNSCsf, "#0.######")
        rha.Text = Format(iPS.Ihasf, "#0.######")
        rKs.Text = Format(iPS.IKssf, "#0.######")
        rKto.Text = Format(iPS.IKtosf, "#0.######")
        rKur.Text = Format(iPS.IKursf, "#0.######")
        rNa.Text = Format(iPS.INasf, "#0.######")
        rVshiftNa.Text = Format(iPS.VshiftNasf, "#0.######")
        rst.Text = Format(iPS.Istsf, "#0.######")

    End Sub

    Public Sub PrintScf()                                'parameters to be adjusted
        txtStp.Text = Format(Stp(1), "#0.#####")   'step size of NP points

        rKr.Text = Format(iPS.IKrsf, "#0.#####")
        rCaL.Text = Format(iPS.ICaLsf, "#0.#####")
        rNa.Text = Format(iPS.INasf, "#0.#####")
        rK1.Text = Format(iPS.IK1sf, "#0.#####")
        'rha.Text = Format(iPS.Ihasf, "#0.#####")
        'rbNSC.Text = Format(iPS.IbNSCsf, "#0.#####")

        NIKrmag.Text = Format(iPS.IKrmag * iPS.IKrsf, "#0.#####")
        NICaLmag.Text = Format(iPS.ICaLmag * iPS.ICaLsf, "#0.#####")
        NINamag.Text = Format(iPS.INamag * iPS.INasf, "#0.#####")
        NIK1mag.Text = Format(iPS.IK1mag * iPS.IK1sf, "#0.#####")
        'NIhamag.Text = Format(iPS.Ihamag * iPS.Ihasf, "#0.#####")
        'NIbNSCmag.Text = Format(iPS.IbNSCmag * iPS.IbNSCsf, "#0.#####")


        'rha.Text = Format(iPS.Ihasf, "##.0###")

        'rNa.Text = Format(iPS.INasf, "##.0###")
        'rst.Text = Format(iPS.Istsf, "##.0###")

        'rKto.Text = Format(iPS.IKtosf, "##.0###")

        'rKs.Text = Format(iPS.IKssf, "##.0###")
        'rK1.Text = Format(iPS.IK1sf, "##.0###")
        'rKto.Text = Format(iPS.IKtosf, "##.0###")
        'rVshiftNa.Text = Format(iPS.VshiftNasf, "##.0###")

        'NIKrmag.Text = Format(iPS.IKrmag * iPS.IKrsf, "##0.####")
        'NICaLmag.Text = Format(iPS.ICaLmag * iPS.ICaLsf, "##0.####")
        'NIbNSCmag.Text = Format(iPS.IbNSCmag * iPS.IbNSCsf, "##0.####")
        'NINamag.Text = Format(iPS.INamag * iPS.INasf, "##0.####")
        ''NIKurmag.Text = Format(iPS.IKurmag * iPS.IKursf, "##0.####")
        'NKLmag.Text = Format(iPS.KLmag * iPS.KLsf, "##0.####")

        'NIhamag.Text = Format(iPS.Ihamag * iPS.Ihasf, "##0.####")

        'NINamag.Text = Format(iPS.INamag * iPS.INasf, "##0.####")
        'NIstmag.Text = Format(iPS.Pstmag * iPS.Istsf, "##0.####")

        oldEval = Eval                                       'replace oldEval
        For I As Integer = 1 To NumPara         'replace NP()
            oldBP(I) = BP(I)
        Next I

    End Sub

    Public countPlot As Long = 0
    Public LogSQS As Double
    Public valueY As Double
    Public oldvalueY As Double = 2

    Private Sub Plot_MSEvsBPx()
        '************ csvWrite of new BP points  ****************************
        If MIN > 0 Then
            valueY = Math.Log10(MIN)
            iPS.LogMIN = valueY
        End If
        IndexY = valueY                       'This is used in the calculateMSE()

        '(1)  ********* write  new BP(x) points in the csvFile *************************************
        'If valueY < 1 Then
        'iPS.LogMIN = valueY     'prepare saving  BP(x) values via csvRead&write() method

        iPS.BP1 = BP(1)
        iPS.BP2 = BP(2)
        iPS.BP3 = BP(3)
        iPS.BP4 = BP(4)
        'iPS.BP5 = BP(5)
        'iPS.BP6 = BP(6)
        'iPS.BP7 = BP(7)

        '********************* csvFile.write;   relationship between BP(x) and Log10(SQS) ***********************
        CSVReadAndWriteE15.WriteCSV(CsvFileNameBPSQS, N_BPSQS, iPS, 1, namesBPSQS)    'Disk write>>>>>>>>

        N_BPSQS += 1         'proceed with the sequential number of BP points
        txtNBPSQS.Text = Format(N_BPSQS, "##0")   'indicate in the text box
        'End If

        '' (2)  ********* plot BP(x) in each graph **************************************************
        'If N_BPSQS >= 0 And valueY <= MaxY15 Then                   'Plot BP(x) when valueY < the maxY of the graph
        '    graphBP1MSE.PlotPoint(BP(1), valueY, Color.Black, 2, 2)
        '    graphBP2MSE.PlotPoint(BP(2), valueY, Color.Black, 2, 2)
        '    graphBP3MSE.PlotPoint(BP(3), valueY, Color.Black, 2, 2)
        '    graphBP4MSE.PlotPoint(BP(4), valueY, Color.Black, 2, 2)
        '    graphBP5MSE.PlotPoint(BP(5), valueY, Color.Black, 2, 2)
        '    graphBP6MSE.PlotPoint(BP(6), valueY, Color.Black, 2, 2)

        'End If

        countPlot = countPlot + 1

        'If countPlot >= 1 Then                 'connect new BP to the old BP 
        graphBP1MSE.PlotLine(oldBP(1), oldvalueY, BP(1), valueY, Color.Black, 1)
        graphBP2MSE.PlotLine(oldBP(2), oldvalueY, BP(2), valueY, Color.Black, 1)
        graphBP3MSE.PlotLine(oldBP(3), oldvalueY, BP(3), valueY, Color.Black, 1)
        graphBP4MSE.PlotLine(oldBP(4), oldvalueY, BP(4), valueY, Color.Black, 1)
        'graphBP5MSE.PlotLine(oldBP(5), oldvalueY, BP(5), valueY, Color.Black, 1)
        'graphBP6MSE.PlotLine(oldBP(6), oldvalueY, BP(6), valueY, Color.Black, 1)

        graphBP1MSE.PlotPoint(BP(1), valueY, Color.Black, 3, 3)
        graphBP2MSE.PlotPoint(BP(2), valueY, Color.Black, 3, 3)
        graphBP3MSE.PlotPoint(BP(3), valueY, Color.Black, 3, 3)
        graphBP4MSE.PlotPoint(BP(4), valueY, Color.Black, 3, 3)
        'graphBP5MSE.PlotPoint(BP(5), valueY, Color.Black, 3, 3)
        'graphBP6MSE.PlotPoint(BP(6), valueY, Color.Black, 3, 3)

        '********** parameters for the line plot ******************
        oldBP(1) = BP(1)        'replace the old BP
        oldBP(2) = BP(2)        'replace the old BP
        oldBP(3) = BP(3)        'replace the old BP
        oldBP(4) = BP(4)        'replace the old BP
        'oldBP(5) = BP(5)        'replace the old BP
        'oldBP(6) = BP(6)        'replace the old BP
        'oldBP(7) = BP(7)        'replace the old BP
        oldvalueY = valueY        'replace the old valueY
        'End If
        'End If

        'End If

    End Sub

    Private Sub Plot_MSEvsBPxRandom2()
        '************ csvWrite of new BP points  ****************************
        If MIN > 0 Then
            valueY = Math.Log10(MIN)
            'iPS.LogMIN = valueY

            ''(1)  ********* write  new BP(x) points in the csvFile **************************************
            iPS.LogMIN = valueY     'prepare saving  BP(x) values via csvRead&write() method

            iPS.BP1 = BP(1)
            iPS.BP2 = BP(2)
            iPS.BP3 = BP(3)
            iPS.BP4 = BP(4)
            'iPS.BP5 = BP(5)
            'iPS.BP6 = BP(6)
            'iPS.BP7 = BP(7)

            '********************* csvFile.write;   relationship between BP(x) and Log10(SQS) ***********************
            'CSVReadAndWriteE15.WriteCSV(CsvFileNameBPSQS, N_BPSQS, iPS, 1, namesBPSQS)    'Disk write>>>>>>>>

            'N_BPSQS = N_BPSQS + 1         'proceed with the sequential number of BP points
            'txtNBPSQS.Text = Format(N_BPSQS, "##0")   'indicate in the text box

            ' (2)  ********* plot BP(x) in each graph **************************************************
            If N_BPSQS > 1 And valueY <= MaxY15 Then                   'Plot BP(x) when valueY < the maxY of the graph
                graphBP1MSE.PlotPoint(BP(1), valueY, Color.Black, 2, 2)
                graphBP2MSE.PlotPoint(BP(2), valueY, Color.Black, 2, 2)
                graphBP3MSE.PlotPoint(BP(3), valueY, Color.Black, 2, 2)
                graphBP4MSE.PlotPoint(BP(4), valueY, Color.Black, 2, 2)
                'graphBP5MSE.PlotPoint(BP(5), valueY, Color.Black, 2, 2)
                'graphBP6MSE.PlotPoint(BP(6), valueY, Color.Black, 2, 2)
                'Else
                '    graphBP1MSE.PlotPoint(BP(1), valueY, Color.Black, 2, 2)
                '    graphBP2MSE.PlotPoint(BP(2), valueY, Color.Black, 2, 2)
                '    graphBP3MSE.PlotPoint(BP(3), valueY, Color.Black, 2, 2)
                '    graphBP4MSE.PlotPoint(BP(4), valueY, Color.Black, 2, 2)
                'graphBP5MSE.PlotPoint(BP(5), valueY, Color.Black, 2, 2)
                '    'graphBP6MSE.PlotPoint(BP(6), valueY, Color.Black, 2, 2)
            End If

            countPlot += 1

            If countPlot >= 1 Then                 'connect new BP to the old BP 
                graphBP1MSE.PlotLine(oldBP(1), oldvalueY, BP(1), valueY, Color.Black, 1)
                graphBP2MSE.PlotLine(oldBP(2), oldvalueY, BP(2), valueY, Color.Black, 1)
                graphBP3MSE.PlotLine(oldBP(3), oldvalueY, BP(3), valueY, Color.Black, 1)
                graphBP4MSE.PlotLine(oldBP(4), oldvalueY, BP(4), valueY, Color.Black, 1)
                'graphBP5MSE.PlotLine(oldBP(5), oldvalueY, BP(5), valueY, Color.Black, 1)
                'graphBP6MSE.PlotLine(oldBP(6), oldvalueY, BP(6), valueY, Color.Black, 1)

                '********** parameters for the line plot ******************
                'oldBP(1) = BP(1)        'replace the old BP
                'oldBP(2) = BP(2)        'replace the old BP
                'oldBP(3) = BP(3)        'replace the old BP
                'oldBP(4) = BP(4)        'replace the old BP
                'oldBP(5) = BP(5)        'replace the old BP
                'oldBP(6) = BP(6)        'replace the old BP
                'oldBP(7) = BP(7)        'replace the old BP
                oldvalueY = valueY        'replace the old valueY
            End If
        End If

        'End If

    End Sub


    Dim Sum_BP1 As Double = 0
    Dim Sum_BP2 As Double = 0
    Dim Sum_BP3 As Double = 0
    Dim Sum_BP4 As Double = 0
    Dim Sum_BP5 As Double = 0
    Dim Sum_BP6 As Double = 0
    Dim sum_BP7 As Double = 0

    Dim Mean_BP1 As Double
    Dim Mean_BP2 As Double
    Dim Mean_BP3 As Double
    Dim Mean_BP4 As Double
    Dim Mean_BP5 As Double
    Dim Mean_BP6 As Double
    Dim Mean_BP7 As Double


    Private Sub PlotExit_MSEvsBPx()

        'iPS.LogMIN = Math.Log10(GminimumMIN)
        iPS.LogMIN = Math.Log10(MIN)
        valueY = iPS.LogMIN

        If valueY <> 0 Then
            '(1)  indicate the last BP points depicted in red when it stopped by Stp = CrtStp
            If Stp(1) <= CrtStp(1) Then
                graphBP1MSE.PlotPoint(BP(1), valueY, Color.Red, 3, 3)
                graphBP2MSE.PlotPoint(BP(2), valueY, Color.Red, 3, 3)
                graphBP3MSE.PlotPoint(BP(3), valueY, Color.Red, 3, 3)
                graphBP4MSE.PlotPoint(BP(4), valueY, Color.Red, 3, 3)
                'graphBP5MSE.PlotPoint(BP(5), valueY, Color.Red, 3, 3)
                'graphBP6MSE.PlotPoint(BP(6), valueY, Color.Red, 3, 3)
            Else                           'optimization stopped by Eval > EvalMax
                graphBP1MSE.PlotPoint(BP(1), valueY, Color.Chocolate, 3, 3)
                graphBP2MSE.PlotPoint(BP(2), valueY, Color.Chocolate, 3, 3)
                graphBP3MSE.PlotPoint(BP(3), valueY, Color.Chocolate, 3, 3)
                graphBP4MSE.PlotPoint(BP(4), valueY, Color.Chocolate, 3, 3)
                'graphBP5MSE.PlotPoint(BP(5), valueY, Color.Chocolate, 3, 3)
                'graphBP6MSE.PlotPoint(BP(6), valueY, Color.Chocolate, 3, 3)
            End If

            Sum_BP1 = Sum_BP1 + BP(1)     'Note, the average is made according to BP(x)
            Sum_BP2 = Sum_BP2 + BP(2)
            Sum_BP3 = Sum_BP3 + BP(3)
            Sum_BP4 = Sum_BP4 + BP(4)
            'Sum_BP5 = Sum_BP5 + BP(5)
            'Sum_BP6 = Sum_BP6 + BP(6)
            'Sum_BP7 = Sum_BP7 + BP(7)

            Mean_BP1 = Sum_BP1 / (PScycle)       'mean values of individual Ixsf (BP) is calculated
            Mean_BP2 = Sum_BP2 / (PScycle)
            Mean_BP3 = Sum_BP3 / (PScycle)
            Mean_BP4 = Sum_BP4 / (PScycle)
            'Mean_BP5 = Sum_BP5 / (PScycle)
            'Mean_BP6 = Sum_BP6 / (PScycle)
            'Mean_BP7 = sum_BP7 / (PScycle)

            txtMeanBP1.Text = Format(Mean_BP1, "#0.###")    '************* print out *********
            txtMeanBP2.Text = Format(Mean_BP2, "#0.###")
            txtMeanBP3.Text = Format(Mean_BP3, "#0.###")
            txtMeanBP4.Text = Format(Mean_BP4, "#0.###")
            'txtMeanBP5.Text = Format(Mean_BP5, "#0.###")
            'txtMeanBP6.Text = Format(Mean_BP6, "#0.###")

        Else                                              'erase the final BP(x) points When there were no progress in PS
            graphBP1MSE.PlotPoint(BP(1), valueY, Color.White, 3, 3)                'erase erroneous points
            graphBP2MSE.PlotPoint(BP(2), valueY, Color.White, 3, 3)
            graphBP3MSE.PlotPoint(BP(3), valueY, Color.White, 3, 3)
            graphBP4MSE.PlotPoint(BP(4), valueY, Color.White, 3, 3)
            'graphBP5MSE.PlotPoint(BP(5), valueY, Color.White, 3, 3)
            'graphBP6MSE.PlotPoint(BP(6), valueY, Color.White, 3, 3)
        End If

        My.Application.DoEvents()
    End Sub

    'prepared by Dr. Kohjitani
    '********************* parameters used *************************************************
    Dim SHIFT_JIS As System.Text.Encoding = System.Text.Encoding.GetEncoding("Shift_JIS")
    Dim reader As FileIO.TextFieldParser = Nothing
    Public count As Long                    'count the number of data points
    Public countSelect As Long
    Public totNpoint As Long          'total number of cell models
    Dim input_expdata As String
    Dim str As String()                               'data array

    Private Sub cbCsvFileList_SelectedIndexChanged(ByVal sender As System.Object, ByVal e As System.EventArgs) Handles cbCsvFileList.SelectedIndexChanged
        If cbCsvFileList.SelectedIndex <> -1 Then                  ' item is selected
            Dim item As ComboBoxControl

            'Load時に追加したオブジェクトの中から選択中のものを取得
            item = DirectCast(cbCsvFileList.SelectedItem, ComboBoxControl)
            input_expdata = item.FullPath
            'TextBox1.Text = input_expdata
        End If

        cbselectCSVfile.Visible = False             'hide the command button 'cbexpData'
        cbPlot.Visible = True                   'show the command button 'cbStart''
    End Sub


    '********** コマンドボタン　Select csvFile *****************************************
    '*************  Select csvFile using 'ButtonAcquire_Click()' and  'cbCsvFileList_SelectedIndexChanged()' ****************

    Private Sub cbselectCSVfile_Click(sender As Object, e As EventArgs) Handles cbselectCSVfile.Click
        Dim path As String
        path = Application.StartupPath()            '実行ファイルのDirectoryを取得
        Dim di As New System.IO.DirectoryInfo(path)
        Dim files_List As System.IO.FileInfo() = di.GetFiles("*.csv", System.IO.SearchOption.AllDirectories)   'list all csv file

        Dim filePropaties As ComboBoxControl

        For Each strFile As System.IO.FileInfo In files_List                       'list up all csv files
            filePropaties = New ComboBoxControl(strFile.FullName, strFile.Name)
            cbCsvFileList.Items.Add(filePropaties)
        Next
        cbPlot.Visible = True
    End Sub


    Private Sub cbClear_Click(sender As Object, e As EventArgs)
        graphpsVm.Clear()
        graphMSE.Clear()

        graphpsVm.DrawAxis(IntervalX8, IntervalY1, "", 9)
        graphMSE.DrawAxis(IntervalX3, IntervalY2, "", 8)
        graphMSE.PlotLine(MinX3, crtMSEdepthA, MaxX3, crtMSEdepthA, Color.Chocolate, 1)             'zero hilizontal level of SQS = 1 mV^2
        graphMSE.PlotLine(MinX3, crtMSEdepthB, MaxX3, crtMSEdepthB, Color.Magenta, 1)             'zero hilizontal level of SQS = 1 mV^2

    End Sub

    'Public gRND As Random = New Random()
    Public valueRND1 As Double
    Public valueRND2 As Double
    Public valueRND3 As Double

    'Public N As Long
    Public previousNP As Integer = 1     'previous No. of parameter
    Public rdNP(7) As Long                     'randamized No. of parameter

    Private Sub RandomizeSequence(ByRef previousNP As Integer)     'determine rdNP(I)
        Select Case previousNP

            Case Is = 4     '*444444444444444444444444444444444444444*  1, 2, 3 being remained 

                valueRND1 = gRND.NextDouble()            '第一段階　　RND(1) によってL(1)を決定する

                '*************** the flow in the First Cascade ****************************
                If valueRND1 <= 1 / 3 Then                      '>>>>>>>>  oldNを除く３個の可能性から
                    rdNP(1) = 1

                    valueRND2 = gRND.NextDouble()                        '続いて、RND(2) によってL(2), L(3)を決定する
                    If valueRND2 <= 1 / 3 Then                           '++++++ 1/3
                        rdNP(2) = 2

                        valueRND3 = gRND.NextDouble()                                   'RND(1) によってL(1)を決定する
                        If valueRND3 <= 1 / 2 Then                              '@@@@@  1/2         ' 4, 1, 2, 3   あるいは、4, 1, 3, 2
                            rdNP(3) = 3
                            rdNP(4) = 4
                        Else
                            rdNP(3) = 4
                            rdNP(4) = 3
                        End If

                    ElseIf valueRND2 <= 2 / 3 Then                        '++++++ 2 / 3
                        rdNP(2) = 3

                        valueRND3 = gRND.NextDouble()                                   'RND(1) によってL(1)を決定する
                        If valueRND3 <= 1 / 2 Then                              '@@@@@  1/2         ' 4, 1, 2, 3   あるいは、4, 1, 3, 2
                            rdNP(3) = 2
                            rdNP(4) = 4
                        Else
                            rdNP(3) = 4
                            rdNP(4) = 2
                        End If

                    ElseIf valueRND2 <= 3 / 3 Then                       '++++++ 2 / 3
                        rdNP(2) = 4

                        valueRND3 = gRND.NextDouble()                                   'RND(1) によってL(1)を決定する
                        If valueRND3 <= 1 / 2 Then                              '@@@@@  1/2         ' 4, 1, 2, 3   あるいは、4, 1, 3, 2
                            rdNP(3) = 2
                            rdNP(4) = 3
                        Else
                            rdNP(3) = 3
                            rdNP(4) = 2
                        End If
                        'Else
                        '    Dim x = valueRND2
                    End If


                    '*************** the flow in the second Cascade ****************************
                ElseIf valueRND1 <= 2 / 3 Then                      '>>>>>>>>  oldNを除く３個の可能性から
                    rdNP(1) = 2

                    valueRND2 = gRND.NextDouble()                        '続いて、RND(2) によってL(2), L(3)を決定する
                    If valueRND2 <= 1 / 3 Then                           '++++++ 1/3
                        rdNP(2) = 3

                        valueRND3 = gRND.NextDouble()                                   'RND(1) によってL(1)を決定する
                        If valueRND3 <= 1 / 2 Then                              '@@@@@  1/2         ' 4, 1, 2, 3   あるいは、4, 1, 3, 2
                            rdNP(3) = 1
                            rdNP(4) = 4
                        Else
                            rdNP(3) = 4
                            rdNP(4) = 1
                        End If

                    ElseIf valueRND2 <= 2 / 3 Then                        '++++++ 2 / 3
                        rdNP(2) = 4

                        valueRND3 = gRND.NextDouble()                                   'RND(1) によってL(1)を決定する
                        If valueRND3 <= 1 / 2 Then                              '@@@@@  1/2         ' 4, 1, 2, 3   あるいは、4, 1, 3, 2
                            rdNP(3) = 1
                            rdNP(4) = 3
                        Else
                            rdNP(3) = 3
                            rdNP(4) = 1
                        End If

                    ElseIf valueRND2 <= 3 / 3 Then                       '++++++ 2 / 3
                        rdNP(2) = 1

                        valueRND3 = gRND.NextDouble()                                   'RND(1) によってL(1)を決定する
                        If valueRND3 <= 1 / 2 Then                              '@@@@@  1/2         ' 4, 1, 2, 3   あるいは、4, 1, 3, 2
                            rdNP(3) = 3
                            rdNP(4) = 4
                        Else
                            rdNP(3) = 4
                            rdNP(4) = 3
                        End If

                    End If


                    '*************** the flow in the third Cascade ****************************
                ElseIf valueRND1 <= 3 / 3 Then                      '>>>>>>>>  oldNを除く３個の可能性から
                    rdNP(1) = 3

                    valueRND2 = gRND.NextDouble()                        '続いて、RND(2) によってL(2), L(3)を決定する
                    If valueRND2 <= 1 / 3 Then                           '++++++ 1/3
                        rdNP(2) = 4

                        valueRND3 = gRND.NextDouble()                                   'RND(1) によってL(1)を決定する
                        If valueRND3 <= 1 / 2 Then                              '@@@@@  1/2         ' 4, 1, 2, 3   あるいは、4, 1, 3, 2
                            rdNP(3) = 1
                            rdNP(4) = 2
                        Else
                            rdNP(3) = 2
                            rdNP(4) = 1
                        End If

                    ElseIf valueRND2 <= 2 / 3 Then                        '++++++ 2 / 3
                        rdNP(2) = 1

                        valueRND3 = gRND.NextDouble()                                   'RND(1) によってL(1)を決定する
                        If valueRND3 <= 1 / 2 Then                              '@@@@@  1/2         ' 4, 1, 2, 3   あるいは、4, 1, 3, 2
                            rdNP(3) = 2
                            rdNP(4) = 4
                        Else
                            rdNP(3) = 4
                            rdNP(4) = 2
                        End If

                    ElseIf valueRND2 <= 3 / 3 Then                       '++++++ 2 / 3
                        rdNP(2) = 2

                        valueRND3 = gRND.NextDouble()                                   'RND(1) によってL(1)を決定する
                        If valueRND3 <= 1 / 2 Then                              '@@@@@  1/2         ' 4, 1, 2, 3   あるいは、4, 1, 3, 2
                            rdNP(3) = 1
                            rdNP(4) = 4
                        Else
                            rdNP(3) = 4
                            rdNP(4) = 1
                        End If

                    End If

                End If

                'txt1st.Text = Format(rdNP(1), "#0")
                'txt2nd.Text = Format(rdNP(2), "#0")
                'txt3rd.Text = Format(rdNP(3), "#0")
                'txt4th.Text = Format(rdNP(4), "#0")

                'My.Application.DoEvents()
                'End If


                '****************** oldNP() = 1 *************************************************
            Case Is = 1           '*1111111111111111111111111111111111111111111**  1, 2, 3 being remained 

                valueRND1 = gRND.NextDouble()            '第一段階　　RND(1) によってL(1)を決定する

                '*************** the flow in the First Cascade ****************************
                If valueRND1 <= 1 / 3 Then                      '>>>>>>>>  oldNを除く３個の可能性から
                    rdNP(1) = 2

                    valueRND2 = gRND.NextDouble()                        '続いて、RND(2) によってL(2), L(3)を決定する
                    If valueRND2 <= 1 / 3 Then                           '++++++ 1/3
                        rdNP(2) = 3

                        valueRND3 = gRND.NextDouble()                                   'RND(1) によってL(1)を決定する
                        If valueRND3 <= 1 / 2 Then                              '@@@@@  1/2         ' 4, 1, 2, 3   あるいは、4, 1, 3, 2
                            rdNP(3) = 1
                            rdNP(4) = 4
                        Else
                            rdNP(3) = 4
                            rdNP(4) = 1
                        End If

                    ElseIf valueRND2 <= 2 / 3 Then                        '++++++ 2 / 3
                        rdNP(2) = 4

                        valueRND3 = gRND.NextDouble()                                   'RND(1) によってL(1)を決定する
                        If valueRND3 <= 1 / 2 Then                              '@@@@@  1/2         ' 4, 1, 2, 3   あるいは、4, 1, 3, 2
                            rdNP(3) = 1
                            rdNP(4) = 3
                        Else
                            rdNP(3) = 3
                            rdNP(4) = 1
                        End If

                    ElseIf valueRND2 <= 3 / 3 Then                       '++++++ 2 / 3
                        rdNP(2) = 1

                        valueRND3 = gRND.NextDouble()                                   'RND(1) によってL(1)を決定する
                        If valueRND3 <= 1 / 2 Then                              '@@@@@  1/2         ' 4, 1, 2, 3   あるいは、4, 1, 3, 2
                            rdNP(3) = 3
                            rdNP(4) = 4
                        Else
                            rdNP(3) = 4
                            rdNP(4) = 3
                        End If

                    End If


                    '*************** the flow in the second Cascade ****************************
                ElseIf valueRND1 <= 2 / 3 Then                      '>>>>>>>>  oldNを除く３個の可能性から
                    rdNP(1) = 3

                    valueRND2 = gRND.NextDouble()                        '続いて、RND(2) によってL(2), L(3)を決定する
                    If valueRND2 <= 1 / 3 Then                           '++++++ 1/3
                        rdNP(2) = 4

                        valueRND3 = gRND.NextDouble()                                   'RND(1) によってL(1)を決定する
                        If valueRND3 <= 1 / 2 Then                              '@@@@@  1/2         ' 4, 1, 2, 3   あるいは、4, 1, 3, 2
                            rdNP(3) = 1
                            rdNP(4) = 2
                        Else
                            rdNP(3) = 2
                            rdNP(4) = 1
                        End If

                    ElseIf valueRND2 <= 2 / 3 Then                        '++++++ 2 / 3
                        rdNP(2) = 1

                        valueRND3 = gRND.NextDouble()                                   'RND(1) によってL(1)を決定する
                        If valueRND3 <= 1 / 2 Then                              '@@@@@  1/2         ' 4, 1, 2, 3   あるいは、4, 1, 3, 2
                            rdNP(3) = 2
                            rdNP(4) = 4
                        Else
                            rdNP(3) = 4
                            rdNP(4) = 2
                        End If

                    ElseIf valueRND2 <= 3 / 3 Then                       '++++++ 2 / 3
                        rdNP(2) = 2

                        valueRND3 = gRND.NextDouble()                                   'RND(1) によってL(1)を決定する
                        If valueRND3 <= 1 / 2 Then                              '@@@@@  1/2         ' 4, 1, 2, 3   あるいは、4, 1, 3, 2
                            rdNP(3) = 1
                            rdNP(4) = 4
                        Else
                            rdNP(3) = 4
                            rdNP(4) = 1
                        End If

                    End If


                    '*************** the flow in the third Cascade ****************************

                ElseIf valueRND1 <= 3 / 3 Then                      '>>>>>>>>  oldNを除く３個の可能性から
                    rdNP(1) = 4

                    valueRND2 = gRND.NextDouble()                        '続いて、RND(2) によってL(2), L(3)を決定する
                    If valueRND2 <= 1 / 3 Then                           '++++++ 1/3
                        rdNP(2) = 1

                        valueRND3 = gRND.NextDouble()                                   'RND(1) によってL(1)を決定する
                        If valueRND3 <= 1 / 2 Then                              '@@@@@  1/2         ' 4, 1, 2, 3   あるいは、4, 1, 3, 2
                            rdNP(3) = 2
                            rdNP(4) = 3
                        Else
                            rdNP(3) = 3
                            rdNP(4) = 2
                        End If

                    ElseIf valueRND2 <= 2 / 3 Then                        '++++++ 2 / 3
                        rdNP(2) = 2

                        valueRND3 = gRND.NextDouble()                                   'RND(1) によってL(1)を決定する
                        If valueRND3 <= 1 / 2 Then                              '@@@@@  1/2         ' 4, 1, 2, 3   あるいは、4, 1, 3, 2
                            rdNP(3) = 1
                            rdNP(4) = 3
                        Else
                            rdNP(3) = 3
                            rdNP(4) = 1
                        End If

                    ElseIf valueRND2 <= 3 / 3 Then                       '++++++ 2 / 3
                        rdNP(2) = 3

                        valueRND3 = gRND.NextDouble()                                   'RND(1) によってL(1)を決定する
                        If valueRND3 <= 1 / 2 Then                              '@@@@@  1/2         ' 4, 1, 2, 3   あるいは、4, 1, 3, 2
                            rdNP(3) = 1
                            rdNP(4) = 2
                        Else
                            rdNP(3) = 2
                            rdNP(4) = 1
                        End If

                    End If

                End If

                'txt1st.Text = Format(rdNP(1), "#0")
                'txt2nd.Text = Format(rdNP(2), "#0")
                'txt3rd.Text = Format(rdNP(3), "#0")
                'txt4th.Text = Format(rdNP(4), "#0")

                'My.Application.DoEvents()

                'cbStart.Visible = True
                'End If


            Case Is = 2                 ''*222222222222222222222222222222222222222222*  1, 2, 3 being remained 

                valueRND1 = gRND.NextDouble()            '第一段階　　RND(1) によってL(1)を決定する

                '*************** the flow in the First Cascade ****************************
                If valueRND1 <= 1 / 3 Then                      '>>>>>>>>  oldNを除く３個の可能性から
                    rdNP(1) = 3

                    valueRND2 = gRND.NextDouble()                        '続いて、RND(2) によってL(2), L(3)を決定する
                    If valueRND2 <= 1 / 3 Then                           '++++++ 1/3
                        rdNP(2) = 4

                        valueRND3 = gRND.NextDouble()                                   'RND(1) によってL(1)を決定する
                        If valueRND3 <= 1 / 2 Then                              '@@@@@  1/2         ' 4, 1, 2, 3   あるいは、4, 1, 3, 2
                            rdNP(3) = 1
                            rdNP(4) = 2
                        Else
                            rdNP(3) = 2
                            rdNP(4) = 1
                        End If

                    ElseIf valueRND2 <= 2 / 3 Then                        '++++++ 2 / 3
                        rdNP(2) = 1

                        valueRND3 = gRND.NextDouble()                                   'RND(1) によってL(1)を決定する
                        If valueRND3 <= 1 / 2 Then                              '@@@@@  1/2         ' 4, 1, 2, 3   あるいは、4, 1, 3, 2
                            rdNP(3) = 2
                            rdNP(4) = 4
                        Else
                            rdNP(3) = 4
                            rdNP(4) = 2
                        End If

                    ElseIf valueRND2 <= 3 / 3 Then                       '++++++ 2 / 3
                        rdNP(2) = 2

                        valueRND3 = gRND.NextDouble()                                   'RND(1) によってL(1)を決定する
                        If valueRND3 <= 1 / 2 Then                              '@@@@@  1/2         ' 4, 1, 2, 3   あるいは、4, 1, 3, 2
                            rdNP(3) = 1
                            rdNP(4) = 4
                        Else
                            rdNP(3) = 4
                            rdNP(4) = 1
                        End If
                        'Else
                        '    Dim x = valueRND2
                    End If


                    '*************** the flow in the second Cascade ****************************
                ElseIf valueRND1 <= 2 / 3 Then                      '>>>>>>>>  oldNを除く３個の可能性から
                    rdNP(1) = 4

                    valueRND2 = gRND.NextDouble()                        '続いて、RND(2) によってL(2), L(3)を決定する
                    If valueRND2 <= 1 / 3 Then                           '++++++ 1/3
                        rdNP(2) = 1

                        valueRND3 = gRND.NextDouble()                                   'RND(1) によってL(1)を決定する
                        If valueRND3 <= 1 / 2 Then                              '@@@@@  1/2         ' 4, 1, 2, 3   あるいは、4, 1, 3, 2
                            rdNP(3) = 2
                            rdNP(4) = 3
                        Else
                            rdNP(3) = 3
                            rdNP(4) = 2
                        End If

                    ElseIf valueRND2 <= 2 / 3 Then                        '++++++ 2 / 3
                        rdNP(2) = 2

                        valueRND3 = gRND.NextDouble()                                   'RND(1) によってL(1)を決定する
                        If valueRND3 <= 1 / 2 Then                              '@@@@@  1/2         ' 4, 1, 2, 3   あるいは、4, 1, 3, 2
                            rdNP(3) = 1
                            rdNP(4) = 3
                        Else
                            rdNP(3) = 3
                            rdNP(4) = 1
                        End If

                    ElseIf valueRND2 <= 3 / 3 Then                       '++++++ 2 / 3
                        rdNP(2) = 3

                        valueRND3 = gRND.NextDouble()                                   'RND(1) によってL(1)を決定する
                        If valueRND3 <= 1 / 2 Then                              '@@@@@  1/2         ' 4, 1, 2, 3   あるいは、4, 1, 3, 2
                            rdNP(3) = 1
                            rdNP(4) = 2
                        Else
                            rdNP(3) = 2
                            rdNP(4) = 1
                        End If

                    End If


                    '*************** the flow in the third Cascade ****************************
                ElseIf valueRND1 <= 3 / 3 Then                      '>>>>>>>>  oldNを除く３個の可能性から
                    rdNP(1) = 1

                    valueRND2 = gRND.NextDouble()                        '続いて、RND(2) によってL(2), L(3)を決定する
                    If valueRND2 <= 1 / 3 Then                           '++++++ 1/3
                        rdNP(2) = 2

                        valueRND3 = gRND.NextDouble()                                   'RND(1) によってL(1)を決定する
                        If valueRND3 <= 1 / 2 Then                              '@@@@@  1/2         ' 4, 1, 2, 3   あるいは、4, 1, 3, 2
                            rdNP(3) = 3
                            rdNP(4) = 4
                        Else
                            rdNP(3) = 4
                            rdNP(4) = 3
                        End If

                    ElseIf valueRND2 <= 2 / 3 Then                        '++++++ 2 / 3
                        rdNP(2) = 3

                        valueRND3 = gRND.NextDouble()                                   'RND(1) によってL(1)を決定する
                        If valueRND3 <= 1 / 2 Then                              '@@@@@  1/2         ' 4, 1, 2, 3   あるいは、4, 1, 3, 2
                            rdNP(3) = 2
                            rdNP(4) = 4
                        Else
                            rdNP(3) = 4
                            rdNP(4) = 2
                        End If

                    ElseIf valueRND2 <= 3 / 3 Then                       '++++++ 2 / 3
                        rdNP(2) = 4

                        valueRND3 = gRND.NextDouble()                                   'RND(1) によってL(1)を決定する
                        If valueRND3 <= 1 / 2 Then                              '@@@@@  1/2         ' 4, 1, 2, 3   あるいは、4, 1, 3, 2
                            rdNP(3) = 2
                            rdNP(4) = 3
                        Else
                            rdNP(3) = 3
                            rdNP(4) = 2
                        End If

                    End If

                End If

                'txt1st.Text = Format(rdNP(1), "#0")
                'txt2nd.Text = Format(rdNP(2), "#0")
                'txt3rd.Text = Format(rdNP(3), "#0")
                'txt4th.Text = Format(rdNP(4), "#0")

                'My.Application.DoEvents()
                'End If


            Case Is = 3            'Then        '*3333333333333333333333333333333333333333333333333  1, 2, 3 being remained 

                valueRND1 = gRND.NextDouble()            '第一段階　　RND(1) によってL(1)を決定する

                '*************** the flow in the First Cascade ****************************
                If valueRND1 <= 1 / 3 Then                      '>>>>>>>>  oldNを除く３個の可能性から
                    rdNP(1) = 4

                    valueRND2 = gRND.NextDouble()                        '続いて、RND(2) によってL(2), L(3)を決定する
                    If valueRND2 <= 1 / 3 Then                           '++++++ 1/3
                        rdNP(2) = 1

                        valueRND3 = gRND.NextDouble()                                   'RND(1) によってL(1)を決定する
                        If valueRND3 <= 1 / 2 Then                              '@@@@@  1/2         ' 4, 1, 2, 3   あるいは、4, 1, 3, 2
                            rdNP(3) = 2
                            rdNP(4) = 3
                        Else
                            rdNP(3) = 3
                            rdNP(4) = 2
                        End If

                    ElseIf valueRND2 <= 2 / 3 Then                        '++++++ 2 / 3
                        rdNP(2) = 2

                        valueRND3 = gRND.NextDouble()                                   'RND(1) によってL(1)を決定する
                        If valueRND3 <= 1 / 2 Then                              '@@@@@  1/2         ' 4, 1, 2, 3   あるいは、4, 1, 3, 2
                            rdNP(3) = 1
                            rdNP(4) = 3
                        Else
                            rdNP(3) = 3
                            rdNP(4) = 1
                        End If

                    ElseIf valueRND2 <= 3 / 3 Then                       '++++++ 2 / 3
                        rdNP(2) = 3

                        valueRND3 = gRND.NextDouble()                                   'RND(1) によってL(1)を決定する
                        If valueRND3 <= 1 / 2 Then                              '@@@@@  1/2         ' 4, 1, 2, 3   あるいは、4, 1, 3, 2
                            rdNP(3) = 1
                            rdNP(4) = 2
                        Else
                            rdNP(3) = 2
                            rdNP(4) = 1
                        End If
                        'Else
                        '    Dim x = valueRND2
                    End If


                    '*************** the flow in the second Cascade ****************************
                ElseIf valueRND1 <= 2 / 3 Then                      '>>>>>>>>  oldNを除く３個の可能性から
                    rdNP(1) = 1

                    valueRND2 = gRND.NextDouble()                        '続いて、RND(2) によってL(2), L(3)を決定する
                    If valueRND2 <= 1 / 3 Then                           '++++++ 1/3
                        rdNP(2) = 2

                        valueRND3 = gRND.NextDouble()                                   'RND(1) によってL(1)を決定する
                        If valueRND3 <= 1 / 2 Then                              '@@@@@  1/2         ' 4, 1, 2, 3   あるいは、4, 1, 3, 2
                            rdNP(3) = 3
                            rdNP(4) = 4
                        Else
                            rdNP(3) = 4
                            rdNP(4) = 3
                        End If

                    ElseIf valueRND2 <= 2 / 3 Then                        '++++++ 2 / 3
                        rdNP(2) = 3

                        valueRND3 = gRND.NextDouble()                                   'RND(1) によってL(1)を決定する
                        If valueRND3 <= 1 / 2 Then                              '@@@@@  1/2         ' 4, 1, 2, 3   あるいは、4, 1, 3, 2
                            rdNP(3) = 2
                            rdNP(4) = 4
                        Else
                            rdNP(3) = 4
                            rdNP(4) = 2
                        End If

                    ElseIf valueRND2 <= 3 / 3 Then                       '++++++ 2 / 3
                        rdNP(2) = 4

                        valueRND3 = gRND.NextDouble()                                   'RND(1) によってL(1)を決定する
                        If valueRND3 <= 1 / 2 Then                              '@@@@@  1/2         ' 4, 1, 2, 3   あるいは、4, 1, 3, 2
                            rdNP(3) = 2
                            rdNP(4) = 3
                        Else
                            rdNP(3) = 3
                            rdNP(4) = 2
                        End If

                    End If


                    '*************** the flow in the third Cascade ****************************
                ElseIf valueRND1 <= 3 / 3 Then                      '>>>>>>>>  oldNを除く３個の可能性から
                    rdNP(1) = 2

                    valueRND2 = gRND.NextDouble()                        '続いて、RND(2) によってL(2), L(3)を決定する
                    If valueRND2 <= 1 / 3 Then                           '++++++ 1/3
                        rdNP(2) = 3

                        valueRND3 = gRND.NextDouble()                                   'RND(1) によってL(1)を決定する
                        If valueRND3 <= 1 / 2 Then                              '@@@@@  1/2         ' 4, 1, 2, 3   あるいは、4, 1, 3, 2
                            rdNP(3) = 1
                            rdNP(4) = 4
                        Else
                            rdNP(3) = 4
                            rdNP(4) = 1
                        End If

                    ElseIf valueRND2 <= 2 / 3 Then                        '++++++ 2 / 3
                        rdNP(2) = 4

                        valueRND3 = gRND.NextDouble()                                   'RND(1) によってL(1)を決定する
                        If valueRND3 <= 1 / 2 Then                              '@@@@@  1/2         ' 4, 1, 2, 3   あるいは、4, 1, 3, 2
                            rdNP(3) = 1
                            rdNP(4) = 3
                        Else
                            rdNP(3) = 3
                            rdNP(4) = 1
                        End If

                    ElseIf valueRND2 <= 3 / 3 Then                       '++++++ 2 / 3
                        rdNP(2) = 1

                        valueRND3 = gRND.NextDouble()                                   'RND(1) によってL(1)を決定する
                        If valueRND3 <= 1 / 2 Then                              '@@@@@  1/2         ' 4, 1, 2, 3   あるいは、4, 1, 3, 2
                            rdNP(3) = 3
                            rdNP(4) = 4
                        Else
                            rdNP(3) = 4
                            rdNP(4) = 3
                        End If

                    End If

                End If

        End Select

        txt1st.Text = Format(rdNP(1), "#0")
        txt2nd.Text = Format(rdNP(2), "#0")
        txt3rd.Text = Format(rdNP(3), "#0")
        txt4th.Text = Format(rdNP(4), "#0")
        My.Application.DoEvents()

        'txtoldX.Text = Format(rdNP(4), "#0")

        'previousNP = NP(4)
        previousNP = rdNP(4)                        'renew the previousNP

    End Sub


    Private Sub cbEnd_Click(sender As Object, e As EventArgs) Handles cbEnd.Click
        End
    End Sub

End Class
```
